# Lysine-*R*2HGylation identified as a post-translational modification in *R*2HG-elevated cancers

**DOI:** 10.1101/2025.10.03.680013

**Authors:** Qian Zhao, Peiwu Huang, Changhao Yuan, Peng Xia, Qiang-Shuai Gu, Tonghua Yang, Xiang David Li, Ruijun Tian, Dan Yang

## Abstract

*R*-2-Hydroxyglutarate (*R*2HG), an oncometabolite predominantly produced by mutated isocitrate dehydrogenase 1/2 (IDH1/2) in various cancers, is known to drive cancer progression through noncovalent inhibition of α-ketoglutarate (*α*KG)-dependent enzymes. In this work, we propose an alternative mechanism wherein *R*2HG contributes to cancer development via covalent modification of biologically critical lysines, a process termed lysine-*R*2HGylation (K_*R*2HG_). We designed and synthesized *R*2HG-mimicking probes, demonstrating their effectiveness in facilitating K_*R*2HG_ target profiling and site mapping. We identified K_*R*2HG_ as a previously unrecognized post-translational modification, confirmed its C5-linkage on GSTP1(K209), and demonstrated that SIRT5 functions as a deacylase for GSTP1-K_*R*2HG_ *in vitro*. Furthermore, we found that *R*2HG slightly but significantly inhibits the enzymatic activity of GSTP1 through K_*R*2HG_ and dramatically suppresses monocyte differentiation via this catalytically important lysine modification. Our findings provide an alternative perspective on the role of *R*2HG in leukemia progression and offer a practical tool for the clinical investigation of *R*2HG-elevated cancers.

## Introduction

*R*-2-Hydroxyglutarate (*R*2HG) is an oncometabolite that promotes leukemogenesis in both *in vitro* and *in vivo* models^1–3^. Under normal physiological conditions, *R*2HG is regarded as a minor byproduct present at low levels (ten to hundreds of micromolar)^4–6^. Pathologically, particularly in cancer, *R*2HG levels can surge to tens^6–11^ or even hundreds^12^ of millimolar. This accumulation primarily stems from mutated isocitrate dehydrogenase 1/2 (IDH1/2)^13–16^—key enzymes in the tricarboxylic acid (TCA) cycle that normally catalyze the conversion of isocitrate to *α*-ketoglutarate (*α*KG)^13^. Upon acquiring gain-of-function mutations, most commonly at residues IDH1^R132^ and IDH2^R172^, these enzymes cease *α*KG production and instead convert *α*KG to *R*2HG. These neomorphic mutations are prevalent in multiple cancers, most notably gliomas^14^ and leukemia^14–16^, where they account for 60−90% and 10−40% of all known mutations^14^, respectively.

As the key mediator of oncogenicity induced by IDH1/2 mutants (IDH1/2^mut^), *R*2HG has been shown to drive cancer progression^1,2^ through multiple mechanisms, including metabolic and energetic reprogramming^1,17–23^, epigenetic and signaling desregulation^18,23–28^, and modulation of the tumor microenvironment and immunity^17,29–34^. Most of these proposed mechanisms involve the competitive inhibition of *α*KG-dependent enzymes^28^, including *α*KGDH^17^, SDH, FH^20^, BCAT1-2^21^, ATP5B^19^, TET^18,24^, JMJD2A^27^, JHDMs^27,28^, FTO, ALKBH5^23^, PHD^28^, C-P4H I-III, and Plod 1-3^30^, among others. Owing to its structural similarity to *α*KG, *R*2HG can bind to the *α*KG-binding pockets of these enzymes, inhibiting their activity, disrupting normal function, and thereby promoting carcinogenesis^28^. However, case-specific inconsistencies persist in these proposed models^19,23,35^, and no therapies targeting these mechanisms have yet shown clinical efficacy. Furthermore, given the severe side effects of FDA-approved IDH1/2^mut^-inhibitors^36,37^, particularly in acute myeloid leukemia (AML)^38,39^, there is a need for alternative clinical targets. Ideally, such targets would be more downstream effectors, which may disrupt the established cellular homeostasis^14,17^ to a lesser extent, thereby minimizing adverse effects^38,39^.

Over the past few decades, metabolites have emerged as covalent modulators of reactive amino acid residues in proteins, as exemplified by studies on oncometabolites such as succinate and fumarate^40^. These metabolites influence biological processes through post-translational modifications (PTMs): succinate modifies lysines, while fumarate targets cysteines, for instance^41,42^. Given the low stoichiometry of most, if not all PTMs, enrichment of modified peptides has been deemed necessary to map modification sites, prompting the development of various capture strategies^43^. For example, leveraging changes in reactivity upon cysteine PTMs, chemoselective probes have been widely used to rapidly capture cysteine modifications, including redox PTMs and fatty acylations^43^.

In small lysine acylations, antibody-assisted enrichment has helped the discovery and study of many PTMs, including malonylation^44^, succinylation^41^, glutarylation^45^, and lactylation^46^. Alternatively, bioorthogonal PTM analogs can mimic natural PTM donors and incorporate competitively into potential modification sites, a strategy successfully applied in studies of small acylation^47–49^ and long-chain fatty lipidations^50^. Building on the recent discovery of diverse small lysine acylations^50^, we hypothesized that *R*2HG may contribute to oncogenesis through the modification of functional proteins. Guided by the recent use of alkyne analogs in lysine acylation studies^48,49^, we designed and evaluated a series of 2HG mimics, using them to identify potential *R*2HG-modification targets. Using this probe-assisted approach, we aim to elucidate the biological role of *R*2HG as a PTM. This work could help identify potential druggable targets for malignancies harboring IDH1/2^mut^ and potentially resolve existing controversies in *R*2HG studies^19,23,35^.

Here, we show an alternative mechanism wherein *R*2HG contributes to cancer development via covalent modification of biologically critical lysines, a process termed lysine-*R*2HGylation (K_*R*2HG_). We designed and synthesized *R*2HG-mimicking probes, demonstrating their effectiveness in facilitating K_*R*2HG_ target profiling and site mapping. We identified K_*R*2HG_ as an unrecognized PTM, confirmed its C5-linkage on GSTP1(K209), and demonstrated that SIRT5 functions as a deacylase for GSTP1-K_*R*2HG_ *in vitro*. Furthermore, we found that R2HG slightly but significantly inhibits the enzymatic activity of GSTP1 through K_*R*2HG_ and dramatically suppresses monocyte differentiation via this catalytically important lysine modification. Our findings provide a not-yet studied perspective on the role of *R*2HG in leukemia progression and offer a useful tool for the clinical investigation of *R*2HG-elevated cancers.

## Results

### Design and preparation of *R*2HG probes

Based on the structure of 2HG, we designed and prepared several probes with an alkyne group appended at distinct positions (Figure 1A and Supplementary Schemes 1–6). In these probes, we esterified carboxylic acid groups to enhance cell permeability and incorporated a propargyl group to enable fluorescence visualization and biotin-based enrichment via click chemistry. Given the unresolved enantiomer-specific effects of *R*2HG, we separated the enantiomers and diastereomers of the probes for comparative analysis (Supplementary Schemes 1–6 and Supplementary Figures 1–3).

**Figure 1.**
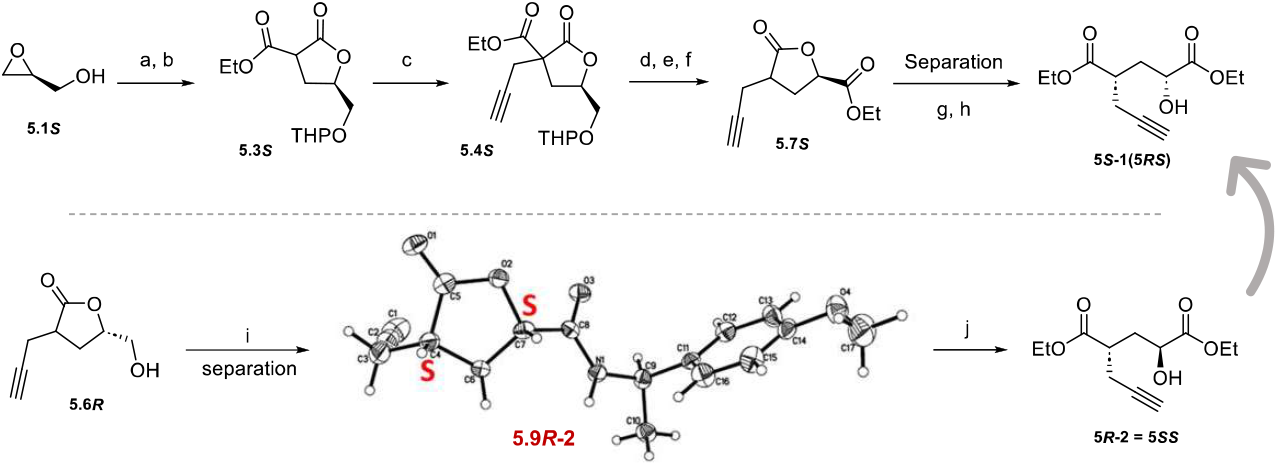
Design and evaluation of synthetic *R*2HG probes. A. *R*2HG mimics were prepared by installing a functional alkyne handle at different positions of *R*2HG, in ester forms to increase cell permeability. Enantiomers and diastereomers were separated, and representative candidates are drawn for each probe series. B. An experimental scheme for evaluating probe similarity to *R*2HG during metabolic labeling, calculated as the intensity difference between probe-labeled and *R*2HG-competed groups. C. Comparison of different probes based on competitive metabolic labeling (Supplementary Figure 9). Probe 5*RS* was selected for further applications with the highest *R*2HG equivalence. D. Metabolic labeling of THP-1 monocytes with probes followed by fluorophore addition via CuAAC enabled in-gel visualization of potential K_*R*2HG_ targets. Competitive labeling by either endogenous (dox-induced IDH1/2^mut^) or exogenous *R*2HG diethyl ester during cotreatments demonstrated the functional similarity between probe 5*RS* and the metabolite *R*2HG. Dox: doxycycline; FL, fluorescence; CB: Coomassie blue staining.

In Series 1 probes, we replaced the hydroxyl O–H moiety of 2HG with an amide N–H bond to preserve potential hydrogen-bonding interactions (Supplementary Scheme 1). However, given the chemical instability of Series 1 probes (Supplementary Figure 4), we extended the alkyne moiety to generate Series 2 probes (Figure 1A, Supplementary Scheme 2), which exhibited enhanced labeling intensity following storage (Supplementary Figure 5). In Probe 3, we used a tertiary alcohol to replace the secondary alcohol of 2HG to prevent potential oxidation in cells (Supplementary Scheme 3). In Probes 4 and 5, we installed propargyl handles at positions farther from the hydroxyl group while preserving its original configuration to better mimic the native structure of 2HG (Supplementary Schemes 4–5).

Finally, we determined the relative configurations of Probe Series 3–5 via chiral derivatization (Scheme 1, Supplementary Schemes 3–6 and Supplementary Figures 1–3), with diastereomeric and enantiomeric purities confirmed using chiral HPLC (Supplementary Information).

### Probes exhibit differential capacity to label *R*2HG substrates

Given the established biological significance of *R*2HG in leukemogenesis, we first tested the probes in several leukemia cell lines, including THP-1, NB-4, OCI-AML3, U937, HL-60, and Jurkat E6-1. Among these suspension cell lines, THP-1, NB-4, and OCI-AML3 showed high labeling efficiency, while others exhibited weak or no labeling (Supplementary Figure 5). We also tested cells from other tissues and species; only human cord blood cells and differentiated THP-1 macrophages exhibited comparable labeling intensity, indicating cell type-dependent *R*2HG probe labeling (Supplementary Figure 6).

A rapid assessment of Series 2 probes with different ester modifications in THP-1 cells revealed that ethyl ester derivatives performed best (Supplementary Figure 7); we therefore used ethyl esters in preparing other probes to optimize labeling efficiency. Among positively labeled cell lines, distinct labeling patterns were observed, likely reflecting differences in protein expression across these cells. For example, fluorescence imaging of gels revealed two distinct bands at ~50 kDa in THP-1 cell lysates, whereas NB-4 and OCI-AML3 lysates showed prominent bands between 30– 40 kDa (Supplementary Figure 5).

Moreover, metabolic labeling intensity did not correlate significantly with protein expression levels, indicating target specificity during metabolic incorporation of the probes. In THP-1 cells, for instance, the 40 kDa protein band was more intense than the 50 kDa band on Coomassie staining but exhibited weaker fluorescence signals (Supplementary Figure 8A). In comparison to other lysine acylation analogs, the 2HG probe exhibited a distinct labeling pattern, and metabolic incorporation was competitively inhibited only by *R*2HG, not by other structurally similar compounds (Supplementary Figures 8B and 9).

Collectively, these results indicate that metabolic labeling with synthetic 2HG probes is *R*2HG-specific, protein-selective, and cell-type-dependent.

### Probe 5*RS* is optimal for experimental applications

To identify the optimal probe for mimicking *R*2HG in metabolic labeling, we treated THP-1 cells—a human M5 acute myeloid leukemia cell line—with various probes in the presence or absence of *R*2HG diethyl ester (used as a competitive inhibitor). Greater structural similarity between a probe and *R*2HG leads to more effective displacement of probe-bound sites by *R*2HG ester during metabolic labeling, manifested as a larger reduction in labeling intensity between the probe-only and competition groups (Figure 1B). Among all tested probes, 5*RS*—with an *R*-configured hydroxyl group and *S*-configured alkyne moiety—most effectively mimicked *R*2HG in labeling assays (Figure 1C).

This specificity and selectivity were further validated in 5*RS* labeling competition assays: both endogenous and exogenous *R*2HG effectively displaced 5*RS* (Figure 1D), whereas other metabolites—including αKG and glutarate—did not (Supplementary Figure 8B). These preliminary tests identified probe 5*RS* as the most robust 2HG mimic for applications in *R*2HGylation target profiling and site mapping.

### Application of 5*RS* in profiling and site-mapping of *R*2HGylated proteins

To optimize labeling conditions for subsequent assays, we evaluated the dose- and time-dependency of probe 5*RS* in human monocytic THP-1 cells (Supplementary Figure 10), identifying 0.5 mM 5*RS* treatment for 6 hours as optimal. Using a strategy similar to that for metabolic labeling (Figure 1B), we employed probe 5*RS* to profile potential *R*2HGylated proteins and used *R*2HG diethyl ester displacement to highlight the most probable targets (Figure 2A). Preliminary comparison of probe-enriched proteins with DMSO-treated controls identified >3,000 and ~1,000 candidates, respectively, indicating that basal (untreated) *R*2HGylation signals are too low to meaningfully contribute to the comparison (Supplementary Figure 11A).

**Figure 2.**
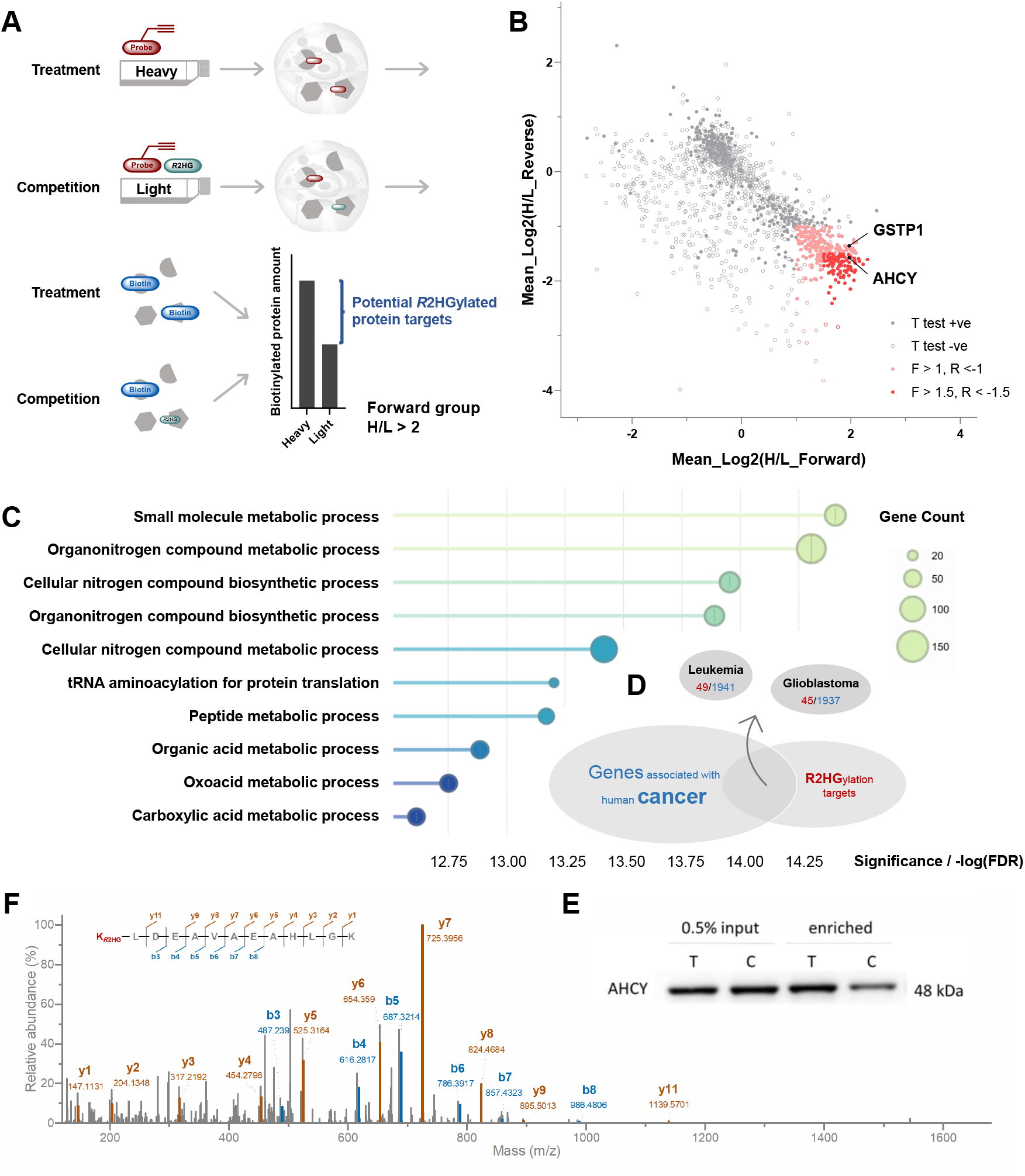
Application of probe 5*RS* in quantitative profiling and site mapping of potential K_*R*2HG_ targets. A. Experimental scheme for sample preparation. B. SILAC quantification of *R*2HG-competed targets from 5*RS*-profiled proteins highlighted potential K_*R*2HG_ targets, with log2(fold change) thresholds set at _±_1.0 and _±_1.5, as labeled in pink and red, respectively. Two profiled targets, GSTP1 and AHCY, are further confirmed via WB (Supplementary Figure 12 and Figure 2E). Results are the means of three independent experiments. Candidates passing one sample t-test (p < 0.05) in either forward or reverse group are noted as filled circles, which counts 271 among the 311 enriched targets. C. Functional clustering of *R*2HG-competed targets from 5*RS-*enriched proteins. Gene ontology of significantly enriched targets (fold change >2) done with STRING (V12.0) highlighted metabolic pathway interruption by excess *R*2HG. D. Around 40−50 out of the top 271 *R*2HGylation targets are reported as genes associated with human leukemia and glioblastoma. E. Validation of AHCY enrichment by synthetic probe 5*RS*. F. An *N*-terminal K_*R*2HG_ was identified with high ion coverage from immunoprecipitated AHCY in dox-induced HEK293T cells co-transfected with IDH1^R132H^. Chemical formula for K_*R*2HG_: C5H6O4, with exact mass = 130.0266.

As outlined in Figure 2A, in a forward SILAC workflow, THP-1 cells were cultured with heavy or light amino acids and labeled with probe 5*RS* in the absence or presence of excess *R*2HG diethyl ester, respectively. Collected protein lysates were quantified, mixed in equal amounts, conjugated to azido-biotin using click chemistry, and affinity-enriched via streptavidin pull-down. Peptides from on-bead digestion were then processed for LC-MS/MS analysis. Quantification results revealed that 311 of 1,586 identified proteins were enriched with at least 2-fold *R*2HG displacement, 271 of which were statistically significant based on a one-sample t-test (Figure 2B and Supplementary Table).

Gene ontology (GO) analysis of potential *R*2HGylation targets suggested a key role for *R*2HG in regulating metabolism-related biological processes (Figure 2C and Supplementary Table). Comparison of these targets with reported cancer-associated genes highlighted ~40–50 proteins potentially involved in leukemia or glioblastoma (Figure 2D and Supplementary Table). From GO biological process clustering, two candidates—GSTP1 and AHCY—appeared in 8/10 and 6/10 of the top 10 GO categories, respectively, and are linked to leukemia/glioblastoma-associated genes. Their protein-specific metabolic labeling and *R*2HG displacement were further confirmed via immunoblotting (Figure 2E and Supplementary Figure 12).

We next mapped lysine-*R*2HGylation (K_*R*2HG_) sites in these potential candidates. Selected *R*2HG targets were overexpressed alongside IDH1/2 mutants in 293T cells for immunoprecipitation (IP)-assisted site mapping (Supplementary Figure 13). Among preliminary spectra, we identified K_*R*2HG_ in AHCY with robust ion coverage and in GSTP1 with acceptable ion coverage (Figure 2F and Supplementary Figure 14). Collectively, these results demonstrate the utility of probe 5*RS* for profiling and site-mapping *R*2HGylated proteins.

### Lysine-*R*2HGylation is an unrecognized PTM

To validate the discovery and obtain high-quality mass spectrometry (MS) spectra, we performed peptide fractionation and MS parallel reaction monitoring (PRM). Given clinically observed acute myeloid leukemia (AML) cases harboring IDH mutations, we selected the human monocytic cell line THP-1 (FAB M5) as a model system and used lentiviral transduction to generate stable THP-1 cell lines expressing IDH1^R132H^ or IDH2^R172K^ (Supplementary Figure 15). Following confirmation that IDH^mut^ was expressed in the correct subcellular compartment (Supplementary Figure 16), IDH^mut^-expressing THP-1 cells were lysed, digested, and subjected to targeted LC-MS/MS analysis using predefined *m/z* values. Across multiple experiments, we consistently detected GSTP1-K_*R*2HG_ in lysates from IDH1^R132H^-expressing THP-1 cells, with corresponding tandem MS spectra, MS^1^ peak envelopes, and LC-extracted ion chromatograms (XICs) shown in Figure 3A and Supplementary Figure 17.

**Figure 3.**
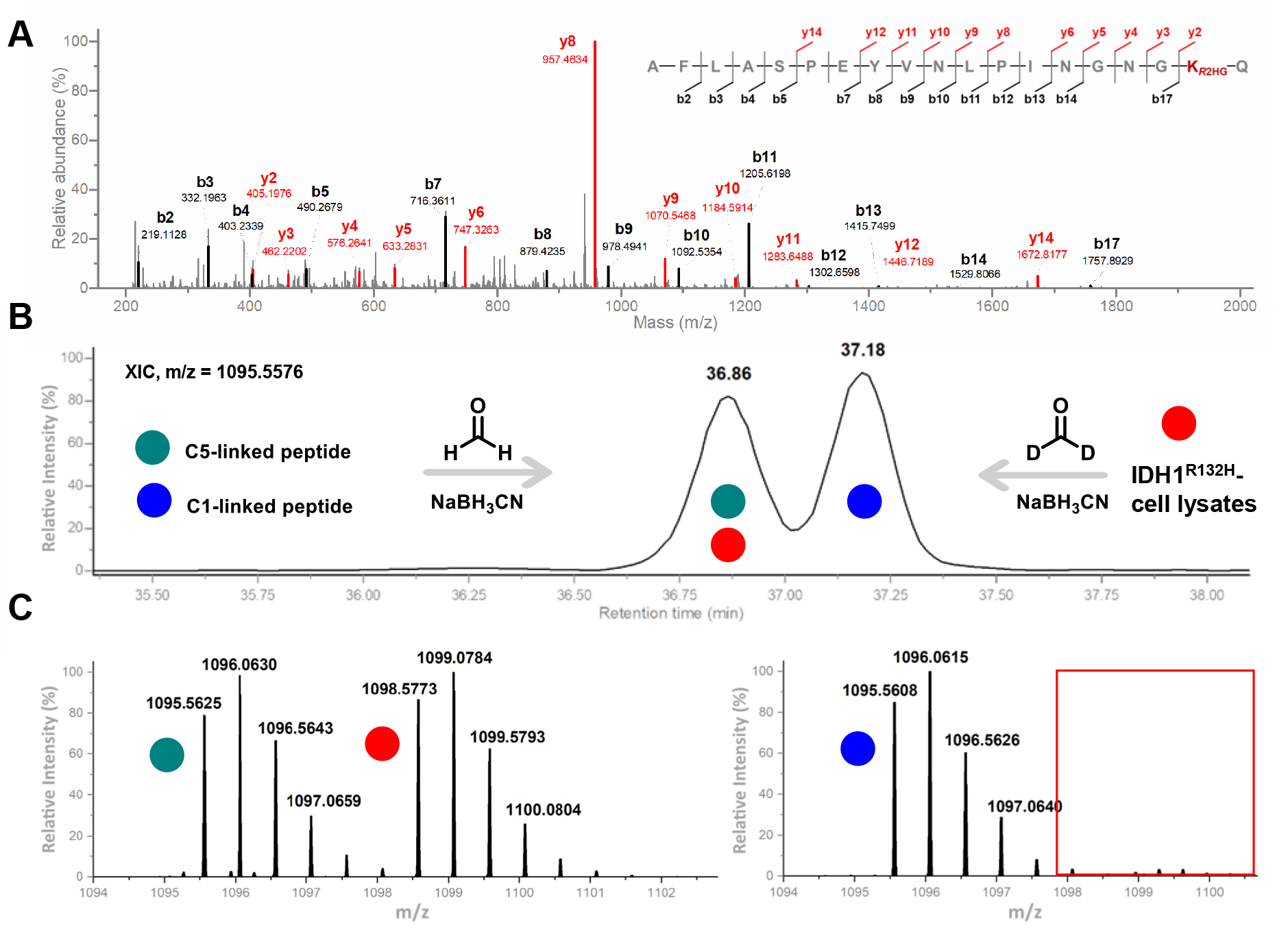
Identification and confirmation of K_*R*2HG_ structure. A. MS2 spectrum of identified K_*R*2HG_ on GSTP1 from IDH1^R132H^-expressed THP-1 monocytes. B. The two LC peaks of the coeluted peptide mixture correspond to dimethylated peptides with C5-(earlier) and C1-*R*2HGylated lysine (later), as proven in a separate trial with different isotopes (Supplementary Figure 19). C. Representative MS1 spectra of each peak in B, which validates the coelution of a heavily dimethylated peptide from THP-1 cells with the lightly labeled synthetic peptide with a C5-K_*R*2HG_.

### Confirmation of the K_*R*2HG_ linkage structure

To determine whether lysine is acylated via the C1 or C5 position of *R*2HG, we synthesized two peptides bearing each modification; however, they were indistinguishable via MS^2^ comparison (Supplementary Figure 18). Noting their subtly different chromatographic behavior, we reductively dimethylated the two peptides with distinct isotopes and confirmed that the C5-*R*2HGylated peptide exhibited a shorter elution time (Supplementary Figure 19).

We then isotopically labeled peptides from solid-phase peptide synthesis (SPPS) and IDH1^R132H^-expressing cells with distinct isotopes and co-eluted them under optimized chromatographic conditions. Co-elution of heavily labeled cellular peptides and lightly labeled SPPS peptides in the earlier-eluting LC peak indicated a C5-linked structure for K_*R*2HG_ *in cellulo* (Figure 3B–C). This assignment was confirmed by their corresponding MS^1^ (Figure 3C) and MS^2^ spectra (Supplementary Figures 20–22).

### SIRT5 functions as the deacylase for lysine *R*2HGylation

During metabolic labeling of THP-1 cells with probe 5*RS*, we observed time-dependent decay of the signal, which indicated the dynamic nature of lysine-*R*2HGylation (Supplementary Figure 10). Given the reported deacylase activity of histone deacetylases (HDACs)^51^, we treated cells with various HDAC inhibitors before 5*RS* labeling to identify a potential deacylase for K_*R*2HG_. Labeling results revealed a slight enhancement in fluorescence upon treatment with the sirtuin (SIRT) inhibitor nicotinamide (Supplementary Figure 23), suggesting sirtuins as candidate K_*R*2HG_-deacetylating enzymes.

To pinpoint the specific sirtuin responsible for K_*R*2HG_ deacylation, we incubated a synthetic K-*R*2HGylated GSTP1 *C*-terminal peptide with recombinant SIRT1–7 under *in vitro* conditions reported in previous studies^48,52^. This assay identified SIRT5 as the sole deacylase for K_*R*2HG_ and confirmed the critical role of an additional *C*-terminal residue in substrate peptides for SIRT5 enzymatic activity (Figure 4A–B and Supplementary Figure 24).

Furthermore, we synthesized the 19-amino-acid *C*-terminal fragment of GSTP1 bearing K_*R*2HG_ and incubated it with SIRT5 over varying durations. Using a pan-antibody against K_*R*2HG_ (Supplementary Figures 25–26), we demonstrated time-dependent deacylation of K_*R*2HG_ by SIRT5 (Figure 4C), further confirming SIRT5 as a deacylase for K_*R*2HG_.

**Figure 4.**
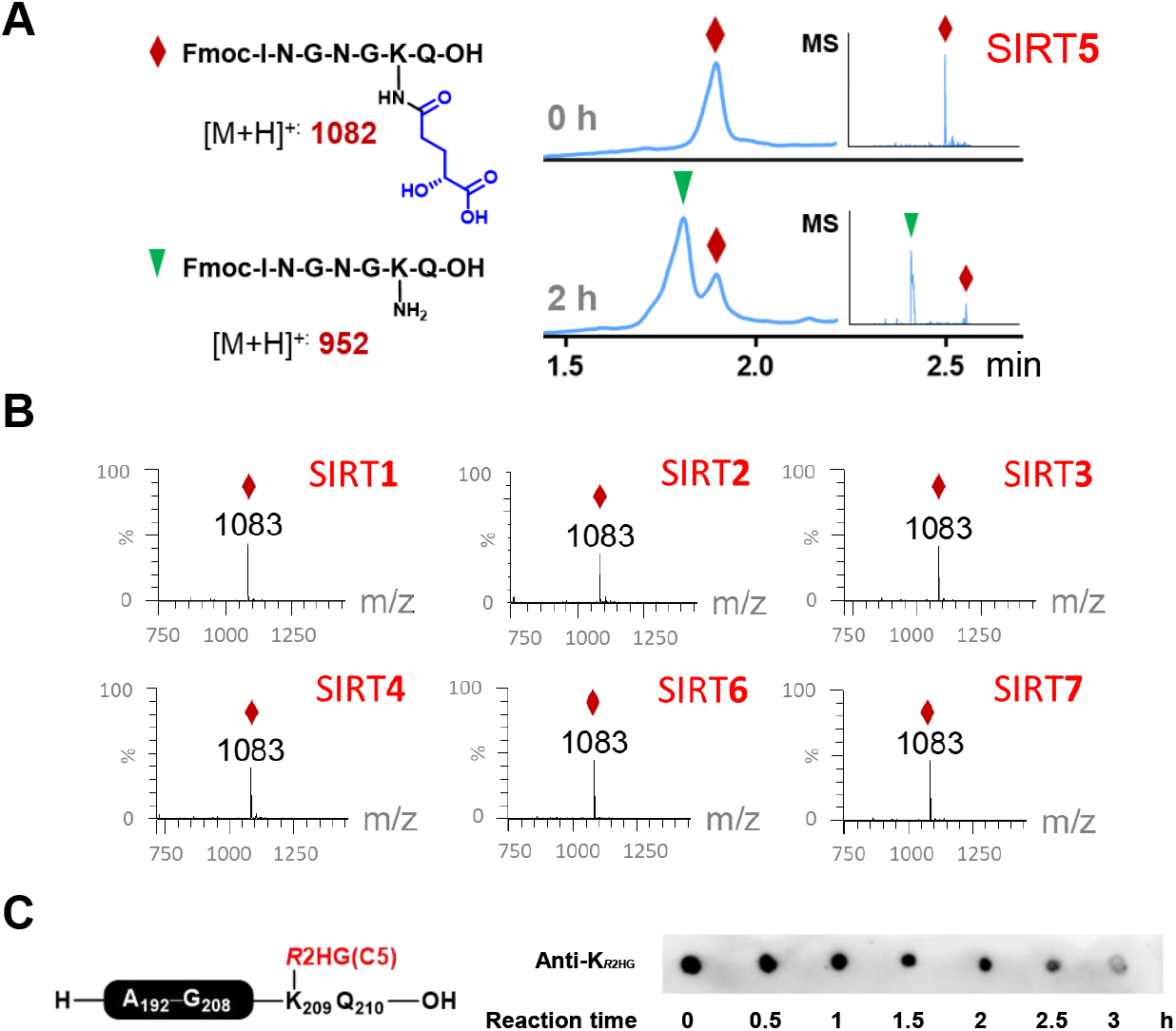
SIRT5 is an eraser for K_*R*2HG_. A. A UV-visible GSTP1 fragment with K_*R*2HG_ can be deacylated by SIRT5, as indicated by the time-dependently increased UPLC-MS signals of unmodified peptide; B. SIRT5 is the only deacylase for K_*R*2HG_ among SIRT1-7. C. The dot blot with pan-K_*R*2HG_ antibody proves the time-dependent deacylase activity of SIRT5 towards K_*R*2HG_ on the identified GSTP1 *C*-terminus (A192-Q210).

### *R*2HGylation of the *C*-terminal region suppresses GSTP1 enzymatic activity

Following the identification of K_*R*2HG_ on the *C*-terminal region of GSTP1, we sought to investigate how this modification influences GSTP1 function. GSTP1 is a well-characterized detoxification enzyme that protects cells from mutagens and carcinogens^51^, and its polymorphisms have been linked to increased cancer risk^52,53^. As a model for functional studies, GSTP1 exhibits measurable activity in conjugating glutathione (GSH) with xenobiotics such as 1-chloro-2,4-dinitrobenzene (CDNB).

We prepared site-specifically K209-*R*2HGylated GSTP1 *in vitro* using intein-mediated protein ligation and assessed its enzymatic activity (Figure 5A and Supplementary Figure 27). Compared to a similarly prepared but unmodified GSTP1 control, K209-*R*2HGylated GSTP1 showed a slight yet significant reduction in catalytic activity (Figure 5B). Both modified and unmodified full-length GSTP1 variants were more active than GSTP1(1–169), a truncated variant lacking the C-terminal peptide (Figure 5B). The *C*-terminal region of GSTP1 is known to form its substrate-binding site^54^, which supports our observation that both *C*-terminal truncation and K209-*R*2HGylation negatively regulate GSTP1 enzymatic activity.

**Figure 5.**
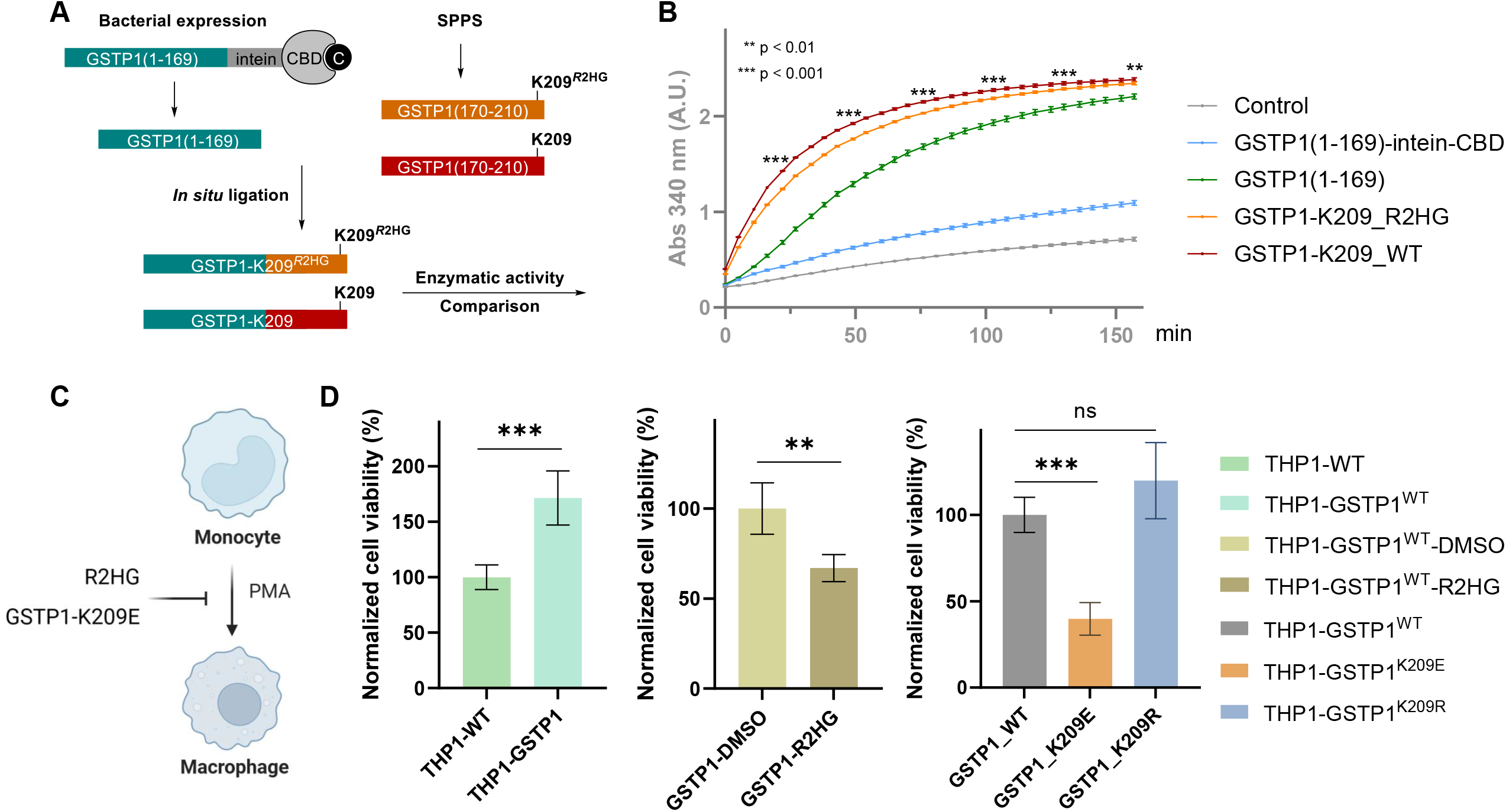
Effect of GSTP1-K209 modifications on its enzymatic activity and cellular fate. A−B. *R*2HGylation on GSTP1-K209 suppresses its enzymatic activity *in vitro*. A. Experimental scheme for intein-mediated protein ligation of K209-modified/unmodified GSTP1. B. Activity comparison of truncated (1 - 169), K209-*R*2HGylated, and K209-unmodified GSTP1 in catalyzing GSH-CDNB conjugation. A.U.: arbitrary unit. C-D. *R*2HG and *R*2HGylation mimic on GSTP1-K209 suppresses THP1 monocyte differentiation. Wt/mutant-GSTP1-expressing THP1 monocytes were treated with 0.1 mM *R*2HG diethyl ester and PMA for 2 days, and the viability of adherent macrophages was measured. D. Left: GSTP1 overexpression promotes THP1 differentiation; middle: *R*2HG inhibits PMA-induced differentiation of wt-GSTP1-expressing THP-1; right: *R*2HGylation-mimicking K209E suppresses the differentiation of mutant-expressing THP1 cells, while the wt-mimicking K209R does not cause a significant change. Data were acquired in triplicates, with t-test values labeled as: ^***^: p < 0.01, ^**^: p < 0.05.

### *R*2HGylation-mimicking GSTP1-K209 mutation suppresses monocyte differentiation

As a cancer-associated marker, GSTP1 polymorphism has been reported to influence leukemogenesis by reducing its detoxification capacity^53^. Given our observation of K_*R*2HG_ in THP-1 monocytes, we next investigated the biological impact of GSTP1-K209_*R*2HG_ on cellular fate. To this end, we generated stable THP-1 cell lines overexpressing wild-type GSTP1 or mutant variants: K209E (to mimic the negatively charged *R*2HGylated lysine) and K209R (to mimic a non-modifiable lysine). Compared to wild-type controls, GSTP1-overexpressing THP-1 cells exhibited enhanced differentiation following 2 days of PMA stimulation, indicating a promotive role for GSTP1 in monocyte-to-macrophage differentiation (Figure 5C and D).

*R*2HG is known to drive leukemogenesis, as it induces growth factor independence, impairs erythroid differentiation *in vitro*^1^, and promotes hyperleukocytosis and leukemia onset *in vivo*^3^. Under mild *R*2HG treatment (0.1 mM), GSTP1-expressing THP-1 cells showed suppressed differentiation upon PMA stimulation (Figure 5D), with no impairment of cell proliferation (Supplementary Figure 28A). This supports that *R*2HG contributes to leukemogenesis, at least in part, by inhibiting monocyte-to-macrophage differentiation. Notably, this differentiation inhibition was not observed in wild-type THP-1 cells under the same mild *R*2HG treatment (Supplementary Figure 28B), indicating that GSTP1 mediates *R*2HG’s suppressive effect on monocyte differentiation.

Furthermore, compared to THP-1 cells overexpressing wild-type GSTP1, those overexpressing GSTP1^K209E^ exhibited significantly reduced differentiation, whereas GSTP1^K209R^-overexpressing cells were largely unaffected (Figure 5D). These results highlight the functional importance of K209 in GSTP1 during monocyte differentiation. Collectively, our findings demonstrate that *R*2HG inhibits THP-1 monocyte differentiation in a GSTP1-dependent manner, likely via covalent *R*2HGylation of GSTP1 at lysine 209—a residue critical for both its enzymatic activity and its role in regulating cell differentiation.

## Discussion

In this study, we proposed an alternative mechanism by which the oncometabolite *R*2HG influences cancer biology: through the covalent modification of cellular proteins, a process we term *R*2HGylation. By synthesizing a panel of enantio- and diastereomerically pure *R*2HG-mimicking probes and applying them in metabolic labeling experiments, we successfully profiled candidate K_*R*2HG_-modified proteins using quantitative proteomics. Among the probe-enriched target proteins, GSTP1 and AHCY enabled us to validate K_*R*2HG_ as a previously unrecognized PTM, with further confirmation of the acylation structure via co-elution using isotopically labeled synthetic peptides.

Notably, even under pathological conditions associated with IDH mutations—where intracellular *R*2HG levels are elevated—K_*R*2HG_ occurs at low stoichiometry, making direct site mapping from total cell lysates particularly challenging. In our LC-MS/MS analyses of 0.5 mM *R*2HG-treated or IDH-mutant-expressing THP-1 cell lysates, we did not detect significant *R*2HGylation. This limitation likely stems from the constraints of data-dependent acquisition (DDA), where MS^2^ spectra are acquired only for the most abundant peptides within a given MS^1^ scan window— potentially excluding low-abundance modified peptides.

To overcome this, our chemical probes helped highlight the most probable *R*2HGylated proteins; when coupled with immunoprecipitation of overexpressed targets, this approach effectively reduced sample complexity and facilitated the identification of K_*R*2HG_. Once the likely modified peptide sequence was identified, we further enhanced detection—even from initially low-quality spectra—by applying targeted MS strategies to enrich and reanalyze the MS^1^ parent ion, enabling improved fragmentation and confident site localization.

Using a synthetic peptide bearing the known K_*R*2HG_ modification, we identified SIRT5 as its deacylase. During characterization of SIRT5 as a K_*R*2HG_ eraser, we observed that deacylase activity was inhibited when the substrate peptide lacked its *C*-terminal glutamine (Q), leaving the K_*R*2HG_ moiety at the peptide terminus. This specificity aligns with SIRT5’s enzymatic mechanism: SIRT5 relies on Arg105 to recognize negatively charged acyl chains^55^, and the proximal *C*-terminal carboxylic acid on the *R*2HGylated lysine may disrupt this interaction by introducing a competing negative charge, thereby impairing substrate binding.

To investigate the acylation mechanism, we explored whether K_*R*2HG_ proceeds via a CoA-dependent pathway, as observed for many acylations. However, attempts to synthesize *R*2HG-C(5)-CoA by activating the C(5) carboxylic acid yielded only cyclic *R*2HG, not the CoA adduct— likely due to spontaneous intramolecular cyclization of the activated C(5) moiety. This instability suggests *R*2HG-CoA is not a viable intermediate, making enzyme-catalyzed acylation via a CoA intermediate unlikely.

In this study, we identified K_*R*2HG_ on GSTP1 at K209. Following confirmation of the acylation structure, we developed pan-K_*R*2HG_ antibodies, which we successfully applied to enrich K_*R*2HG_ sites in human cord blood samples. By combining probe- and antibody-assisted approaches, we are expanding our analyses to include additional candidates, with future work aimed at clarifying the deposition mechanism and deepening our understanding of this PTM’s regulation.

For functional studies, we compared the enzymatic activity of K209-modified and unmodified GSTP1, finding that K_*R*2HG_ induces a slight yet significant suppression of activity. In GSTP1, K209 forms ionic interactions with E113 and hydrogen bonds with I108/Y112^56^, which help restrict the flexible *C*-terminal loop to create a defined substrate-binding pocket. *R*2HGylation of K209 is expected to impair these ionic interactions and potentially disrupt hydrogen bonding due to increased residue size, thereby reducing catalytic activity. However, the small molecular size of our chosen substrate (CDNB) may make it less sensitive to the distorted binding pocket, resulting in subtle enzymatic inhibition. Future studies will evaluate larger carcinogens, which require more space in the catalytic pocket for GSH-mediated detoxification and may thus exhibit greater enzymatic impairment by the *C*-terminal K209-*R*2HGylation.

In cellular fate experiments, our results highlighted the importance of the potential negative charge at GSTP1-K209 and suggested that *R*2HG inhibits differentiation by covalently modifying this critical lysine. Given the role of a negatively charged GSTP1-K209 in maintaining normal monocyte differentiation under appropriate stimuli, *R*2HGylation at this site may disrupt the functional substrate-binding domain, thereby suppressing differentiation, potentially via a mechanism analogous to that reported for GSTP1 polymorphisms^53,57^.

In summary, this study identifies K_*R*2HG_ as a previously unrecognized PTM in *R*2HG-elevated leukemia cells, providing insights into the cellular consequences of this oncometabolite’s accumulation (Figure 6). Given the prevalence of IDH1/2 mutations across cancers—which drive elevated intracellular *R*2HG levels—our findings reveal an alternative mechanism by which *R*2HG may contribute to tumorigenesis. By developing a set of chemically defined 2HG-mimicking probes, we enabled effective metabolic labeling, target identification, and site mapping of *R*2HGylated proteins. The successful application of these tools in both cellular models and clinical blood samples, coupled with the generation of pan-K_*R*2HG_ antibodies, establishes a versatile platform for uncovering previously uncharacterized aspects of *R*2HG biology. These advances not only expand our understanding of metabolic regulation in cancer but also open potential avenues for diagnostic and therapeutic development in 2HG-associated pathologies.

**Figure 6.**
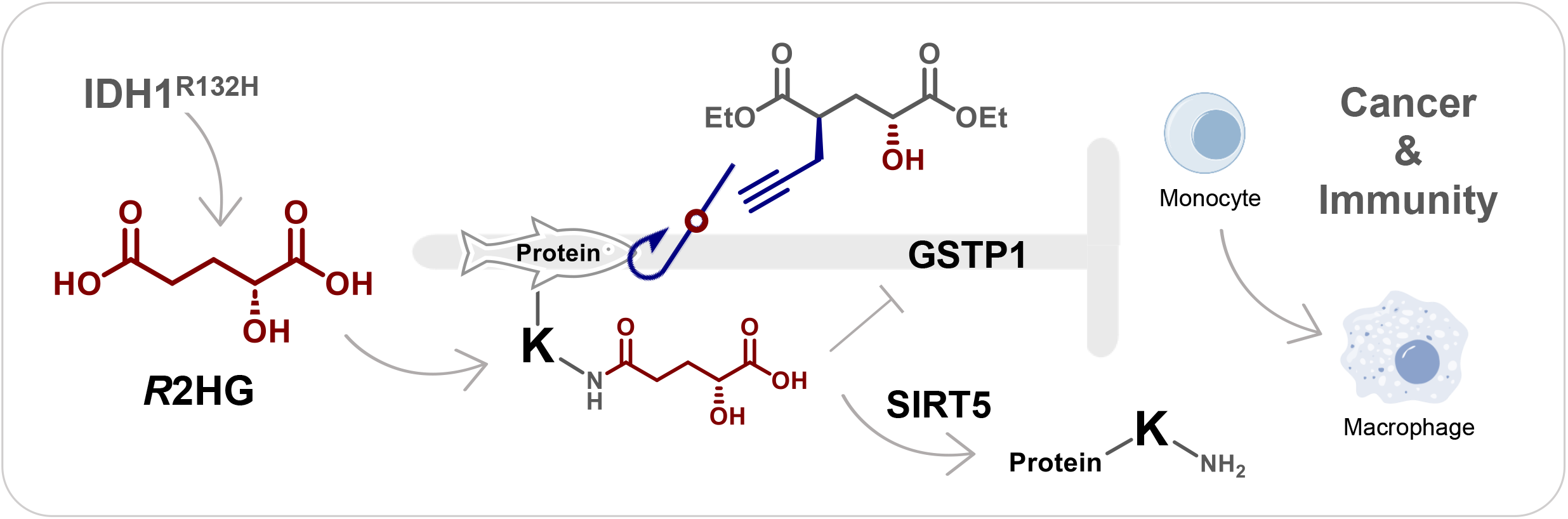
Schematic of the proposed mechanism showing how R2HG impairs monocyte differentiation via KR2HG, a previously unrecognized PTM identified and characterized through probe-assisted approach.

## Methods

Detailed methods sections are provided in the Supplementary Information, including experimental procedures and characterization data.

## Data availability

The mass spectrometry proteomics data have been deposited to the ProteomeXchange Consortium via the PRIDE^58^ partner repository with the dataset identifiers PXD052406 (target identification via SILAC), PXD052411 (PTM identification) and PXD052388 (PTM structure characterization). Other data that support the findings of this study are available from the corresponding author upon reasonable request.

## Acknowledgments

We appreciate the inspiring discussion and warm help from Dr. Xiucong Bao, Dr. Xin Li, and Mr. Hardik Shah. We thank Prof. Quan Hao for recombinant SIRT proteins, Prof. Yu-Hung Leung for cord blood cells, HUABIO company for pan-antibody generation, Ms. Xinning Hu and Dr. Sha Liu for help with experiments, and the NMR, MS, and X-ray facilities of HKU for assistance. We acknowledge financial support from the Westlake Education Foundation, The University of Hong Kong, Morningside Foundation, and the Hong Kong Research Grants Council under the Area of Excellence Scheme (AoE/P705/16).

## Author contributions

D.Y. and Q.Z. conceptualized the study and wrote the manuscript; D.Y., Q.G., Q.Z. designed, Q.Z. conducted synthesis; R.T. designed, P.H., Q.Z., T.Y. conducted proteomics analysis; D.Y., X.D.L, Q.Z. designed, Q.Z., C.Y., P.X. conducted biological studies.

## Competing interests

The authors declare no competing interests.

---

**Scheme 1.**
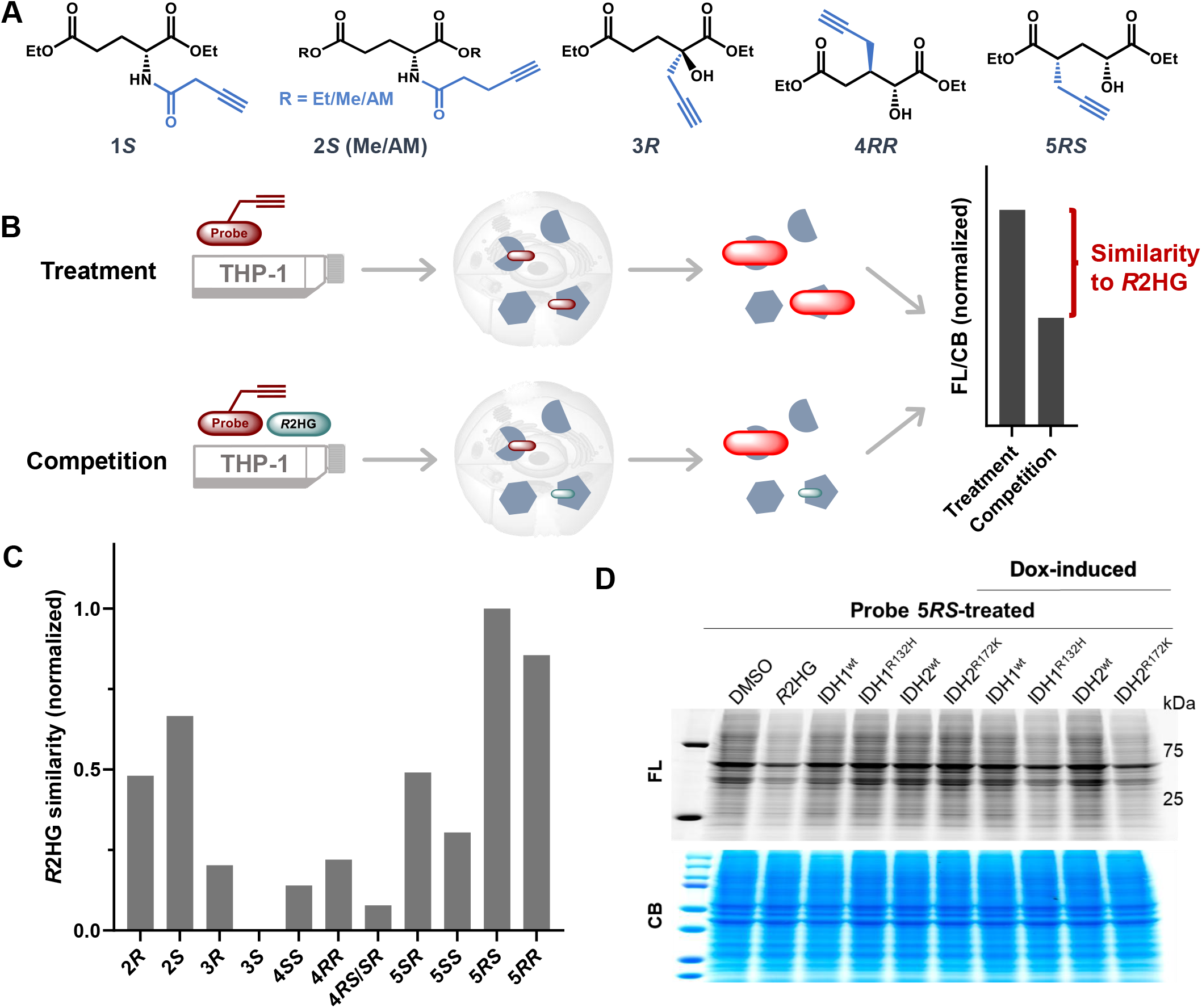
Preparation and chirality determination of probe *5RS*. Probe 5 series was prepared from chiral epoxides to control the chirality of hydroxyl carbons. The intermediate *5*.*6R* was derivatized with a chiral methylphenylamine, enabling crystallization and determination of the relative configuration of the carefully separated product *5*.*9R-2*. Further deprotection and esterification afforded probe *5R-2*, which has the same NMR pattern as 5S-2, indicating an S configuration of the propargyl carbon in probe 5S-1. Conditions: (a) 3,4-dihydropyran, p-TsOH·H_2_O, DCM, rt, 2 h, 99%; (b) diethyl malonate, NaOEt, rt to reflux, 4 h, 75%; (c) NaOEt, EtOH, propargyl bromide, rt, 40 min, 33%; (d) LiOH·H_2_O, 60 °C, 6 h; 3M HCl, to pH= 3; toluene, Dean-Stark apparatus, reflux, overnight, 39%; (e) p-TsOH·H_2_O, MeOH, reflux, 3 h, 86%; (f) CrO_3_, 1.5 M H_2_SO_4_, rt, overnight, crude yield 56%; Etl, NaHCO_3_, DMF, rt, overnight, 9% 5.7S-1; (g) H_2_SO_4_, EtOH, reflux, overnight; TESCl, imidazole, DCM, 0 °C to rt, 15 min, 50%; (h) PPTS, EtOH, rt, 2.5 h, 5S-1: 64%; (i) CrO_3_, 1.5 M H_2_SO_4_, rt, overnight; R-methoxyphenylethylamine, N-methylmorpholine, HOBt, EDCI, DMF/DCM, rt, overnight, 5.9R-2: 2%; G) CAN, MeCN/H_2_O, rt, 40 min, 44%; H_2_SO_4_, EtOH, reflux, overnight; TESCl, imidazole, DCM, 0 °C to rt, 15 min, 27%; PPTS, EtOH, rt, 2.5 h, 5R-2: 63%.

## Notes

### Competing Interest Statement

The authors have declared no competing interest.

