## Supplementary Table for "Lysine-*R*2HGylation identified as a post-translational modification in *R*2HG-elevated cancers": Supplementary Information.pdf

#### Supplementary Figures

##### Synthesis and characterization of probes

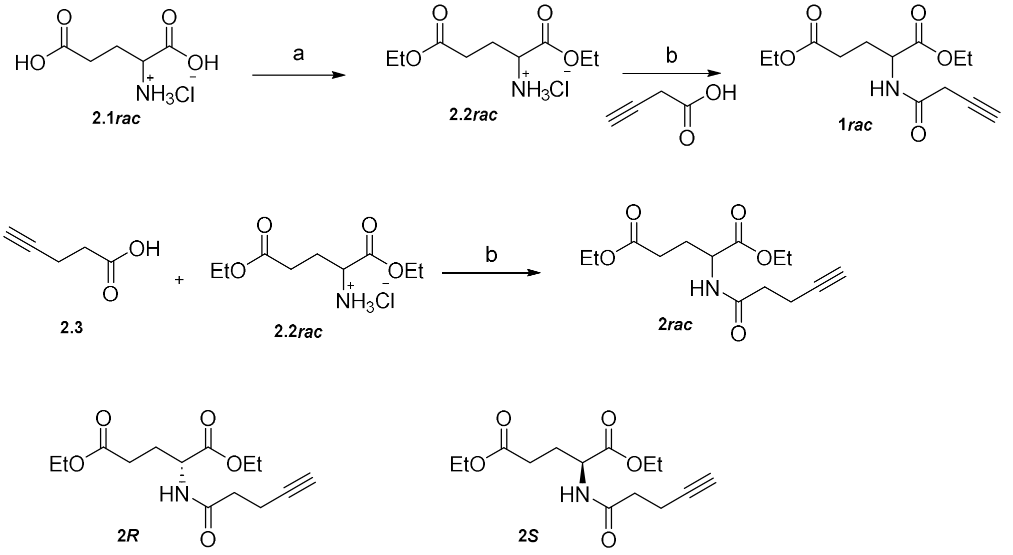

**Supplementary Scheme 1** Synthesis of probes **1*rac*, 2*rac*, 2*R***, and **2*S***. The hydroxyl moiety in 2HG is changed to an amide bond to maintain potential hydrogen bonding, and an additional carbon was added on the alkyne handle to increase probe stability. Conditions: (a) $SOCl_2$, EtOH, 0 °C to reflux, 1.5 h, 99%; (b) EDCI, HOBt, DIEA, DMF, rt, 24 h, **2*rac***: 79%; **2*R***: 72%, 99% ee; **2*S***: 74%, 99% ee.

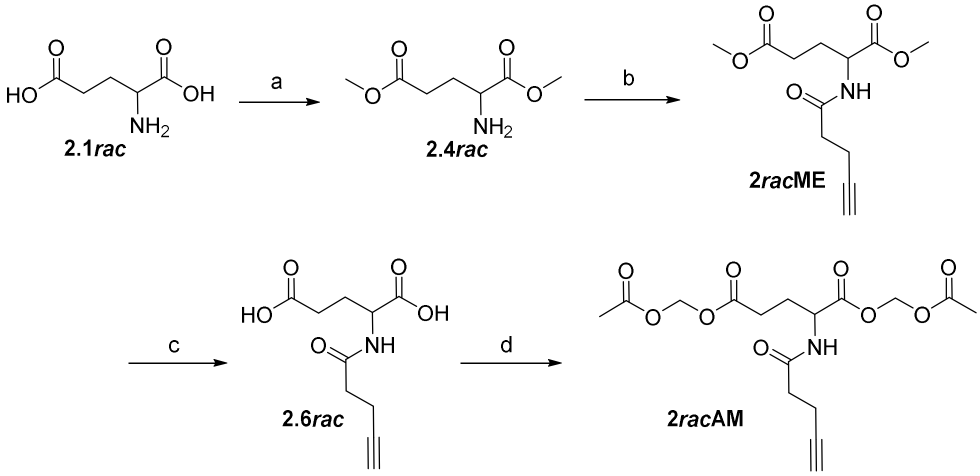

**Supplementary Scheme 2** Synthesis of methyl and acetoxymethyl (AM) esters for probe series **2**, considering possible solubility and cell permeability differences. Conditions: (a) $SOCl_2$, MeOH, 0 °C to rt, reflux, 60 min, 99% for **2.4*R*/*S*/*rac***; (b) 4-pentynoic acid, EDCI, HOBt, DIEA, rt, overnight, **2*R*ME**: 64%, 97% ee; **2*S*ME**:61%, 97% ee; **2*rac*ME**: 66%; (c) LiOH·$H_2O$, dioxane/$H_2O$, rt, 1 h, crude, **2.6*R***: 80%; **2.6*S***:

70%; **2.6*rac***: 56%; (d) bromomethyl acetate, DIPEA, MeCN, rt, 2 h, **2*R*AM**: 54%, 97% ee; **2*S*AM**: 43%, 97% ee; **2*rac*AM**: 74%.

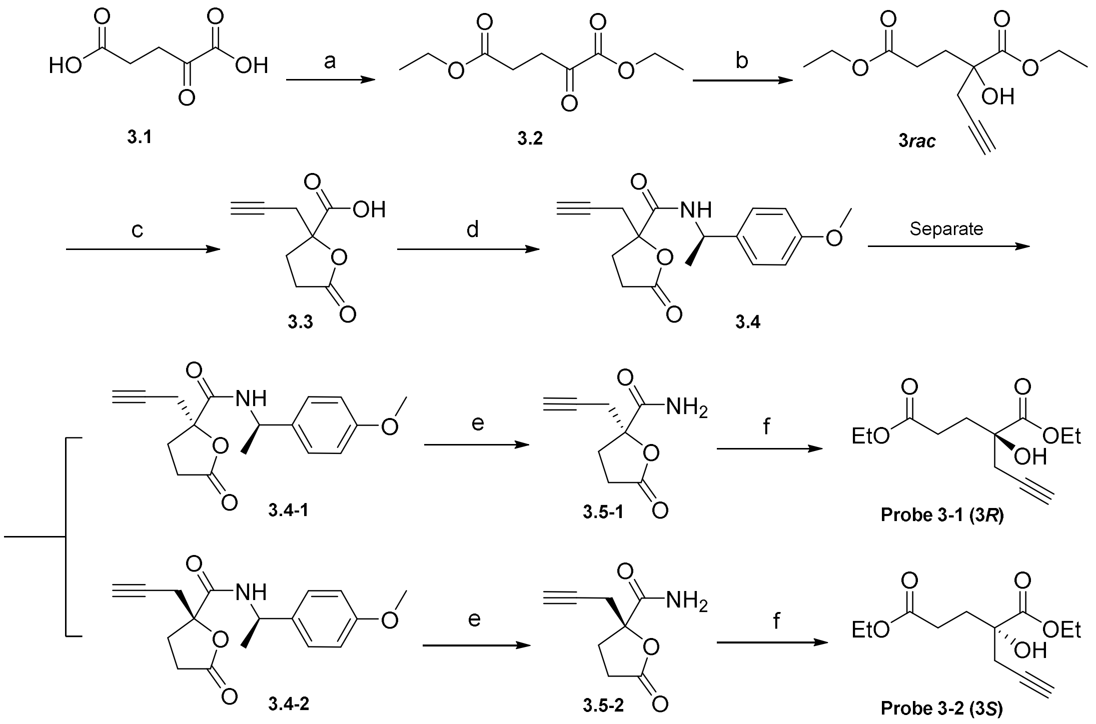

**Supplementary Scheme 3** Synthetic route of probe **3*rac*** and its enantiomers. A tertiary alcohol is used to replace the secondary alcohol of 2HG to suppress oxidative reactions in cells. Conditions: (a) AcCl, EtOH, rt, overnight, 90%; (b) propargyl bromide, Al, $HgCl_2$, THF, −78°C, 2 h, 85%; (c) LiOH, THF /$H_2O$, rt, overnight, then 6 M HCl, 91%; (d) *N*-methylmorpholine, HOBt, EDCI, DCM/DMF, rt, overnight, **3.4-1**: 33%; **3.4-2**: 33%; (e) CAN, CAN/$H_2O$, rt, 40 min, **3.5-1**: 40%; **3.5-2**: 39%; (f). $H_2SO_4$, EtOH, 80 °C, overnight, **3-1**: 30%, 98% ee; **3-2**: 37%, 99% ee.

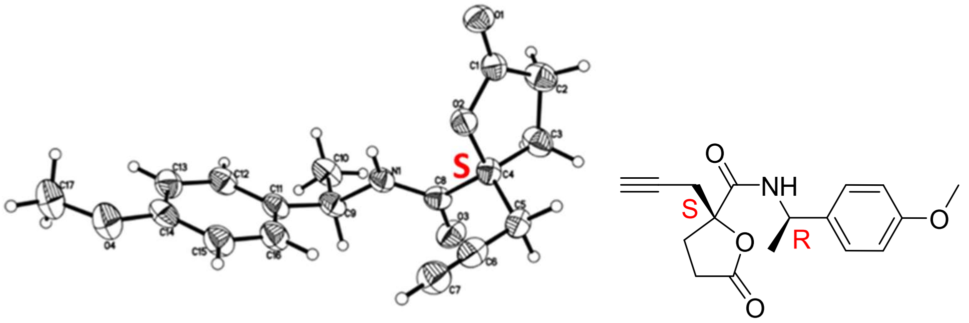

**Supplementary Figure 1** Relative configuration of **3.4-2** determined by X-ray crystallographic analysis. The absolute configuration of the quaternary carbon in **3.4-2** was assigned as ***S***, considering the ***R***-configuration of the chiral auxiliary.

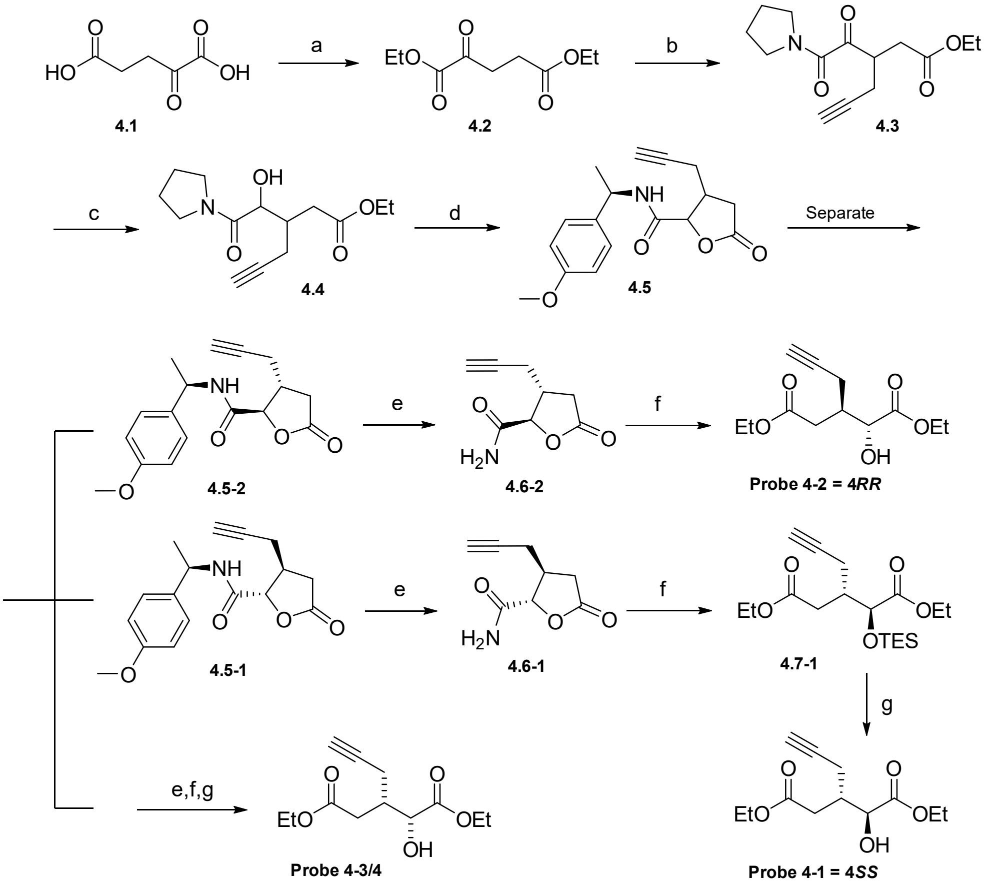

**Supplementary Scheme 4** Synthetic route for probe **4** enantiomers and diastereomers. A propargyl group is installed on the carbon near the hydroxyl group, the original configuration of which is kept to better mimic the original structure of 2HG. Conditions: (a) AcCl, EtOH, rt, overnight, 99%; (b) propargyl bromide

(1−1.2 equiv), freshly prepared LDA (1 equiv), −78 ºC to rt or entirely −78 °C; (c) $NaBH_4$, 0 °C, 40 min, 60%, dr = 1:1; (d) 1 M HCl in EtOH, reflux, overnight, crude 70%; then *R*-methoxyphenylethylamine, *N*-methylmorpholine, HOBt, EDCI, DMF/DCM, rt, overnight, 11% **4.5-1** + 9% **4.5-2** + 6% **4.5-3/4** + 30% mixture; (e) CAN, MeCN/$H_2O$, rt, 40 min, **4.6-1**: 28%, **4.6-2**: 35%, **4.6-3/4**: 40%; (f) $H_2SO_4$, EtOH, reflux, overnight; then TESCl, imidazole, DCM, 0 °C to rt, 15 min; (g) PPTS, EtOH, rt, 2.5 h. Overall yields of steps e and f: **4-1**: 37%, **4-2**: 39%, **4-3/4**: 28%.

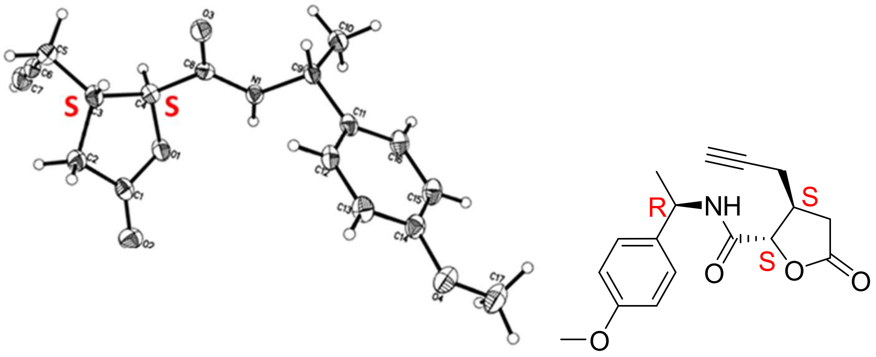

**Supplementary Figure 2** Relative configuration of **4.5-1** determined by X-ray crystallographic analysis. The absolute configurations of the two chiral centers in **4.5-1** were both assigned as ***S***, considering the ***R***-configuration of the chiral auxiliary.

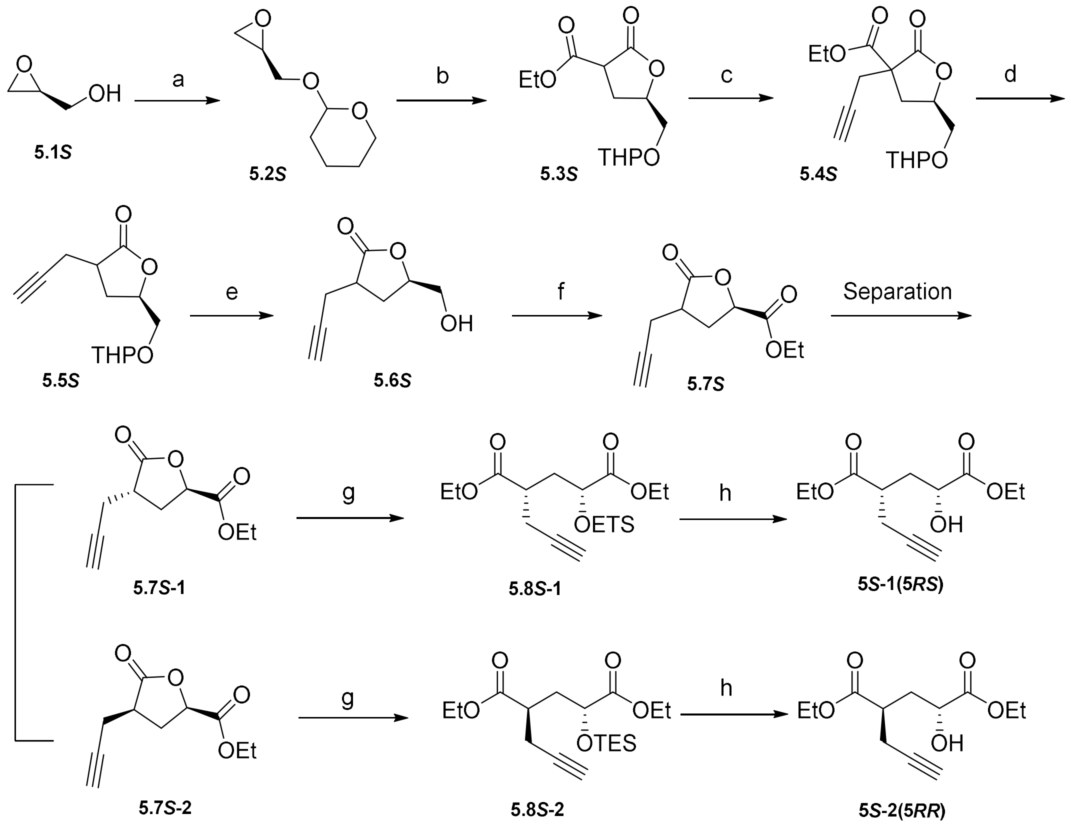

**Supplementary Scheme 5** Synthetic route for probe **5** diastereomers with *R*-hydroxyl group. The propargyl handle of probe **5** was installed in a position further away from the hydroxyl group to mimic the original structure of 2HG. Conditions: (a) 3,4-dihydropyran, *p*-TsOH·$H_2O$, DCM, rt, 2 h, 99%; (b) diethyl malonate, NaOEt, rt to reflux, 4 h, 75%; (c) NaOEt, EtOH, propargyl bromide, rt, 40 min, 33%; (d) LiOH·$H_2O$, 60 °C, 6 h; 3 M HCl, to pH = 3; toluene, Dean-Stark apparatus, reflux, overnight, 39%; (e) *p*-TsOH·$H_2O$, MeOH, reflux, 3 h, 86%; (f) $CrO_3$, 1.5 M $H_2SO_4$, rt, overnight, crude yield 56%; EtI, $NaHCO_3$, DMF, rt, overnight, 9% **5.7*S*-1** + 12% **5.7*S*-2** + 11% mixture; (g) $H_2SO_4$, EtOH, reflux, overnight; TESCl, imidazole, DCM, 0 °C to rt, 15 min, **5.8*S*-1**: 50%; **5.8*S*-2**: 63%; (h) PPTS, EtOH, rt, 2.5 h, **5*S*-1**: 64%; **5*S*-2**: 73%. Diastereomers with *S*-hydroxyl group were prepared similarly from *R*-glycidol, with yields listed below: (a) **5.2*R***, 74%; (b) **5.3*R***, 69%; (c) **5.4*R***, 86%; (d) **5.5*R***, 64%; (e) **5.6*R***, 90%; (f) 38%; 5% **5.7*R*-1** + 15% **5.7*R*-1** + 10% mixture; (g) **5.8*R*-1**, 44%; **5.8*R*-1**, 33%; (h) **5*R*-1**: 35%; **5*R*-2**: 50%.

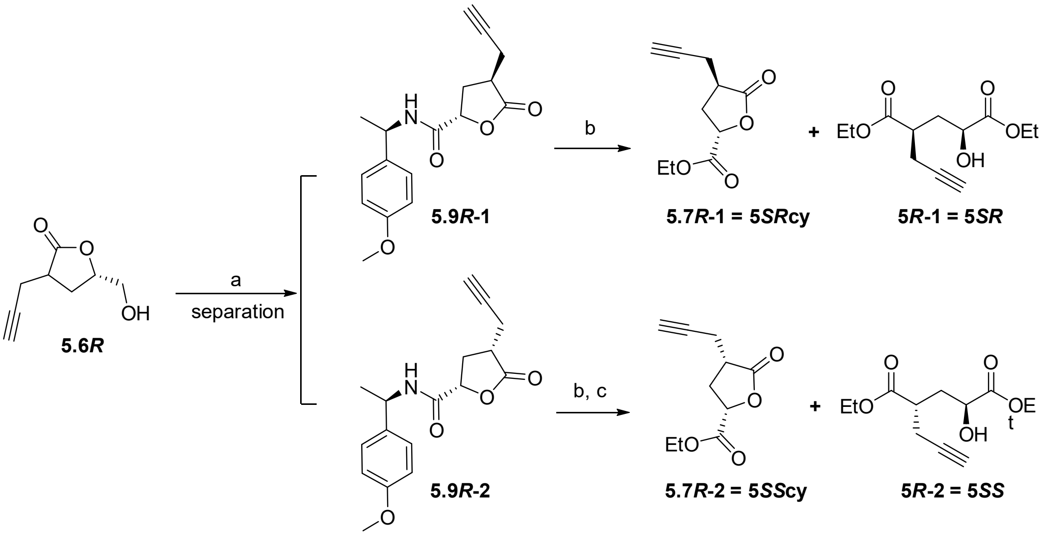

**Supplementary Scheme 6** Synthesis of diastereomer derivatives for assignments of probe chiral centers. Conditions: (a) $CrO_3$, 1.5 M $H_2SO_4$, rt, overnight; *R*-methoxyphenylethylamine, *N*-methylmorpholine, HOBt, EDCI, DMF/DCM, rt, overnight, 5% **5.9*R*-1** + 2% **5.9*R*-2** + 17% mixture; (b) CAN, MeCN/$H_2O$, rt, 40 min, 60% and 44% from **5.9*R*-1** and **5.9*R*-2**; $H_2SO_4$, EtOH, reflux, overnight, **5.7*R*-1**: 23%, **5*R*-1**: 37%; (c) TESCl, imidazole, DCM, 0 °C to rt, 15 min, 30%, **5.7*R*-2**: 27% from b; PPTS, EtOH, rt, 2.5 h, **5*R*-2**: 63%.

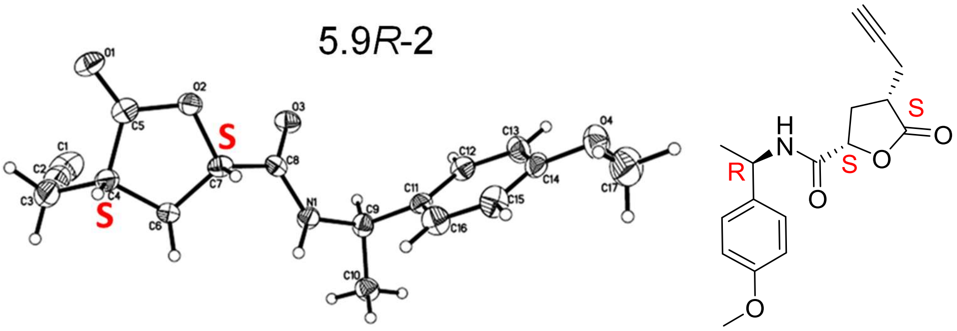

**Supplementary Figure 3** Relative configuration of **5.9*R*-2** determined by X-ray crystallographic analysis. The absolute configurations of the two chiral carbons in **5.9*R*-2** were both assigned as ***S*,** considering the ***R***-configuration of the chiral auxiliary.

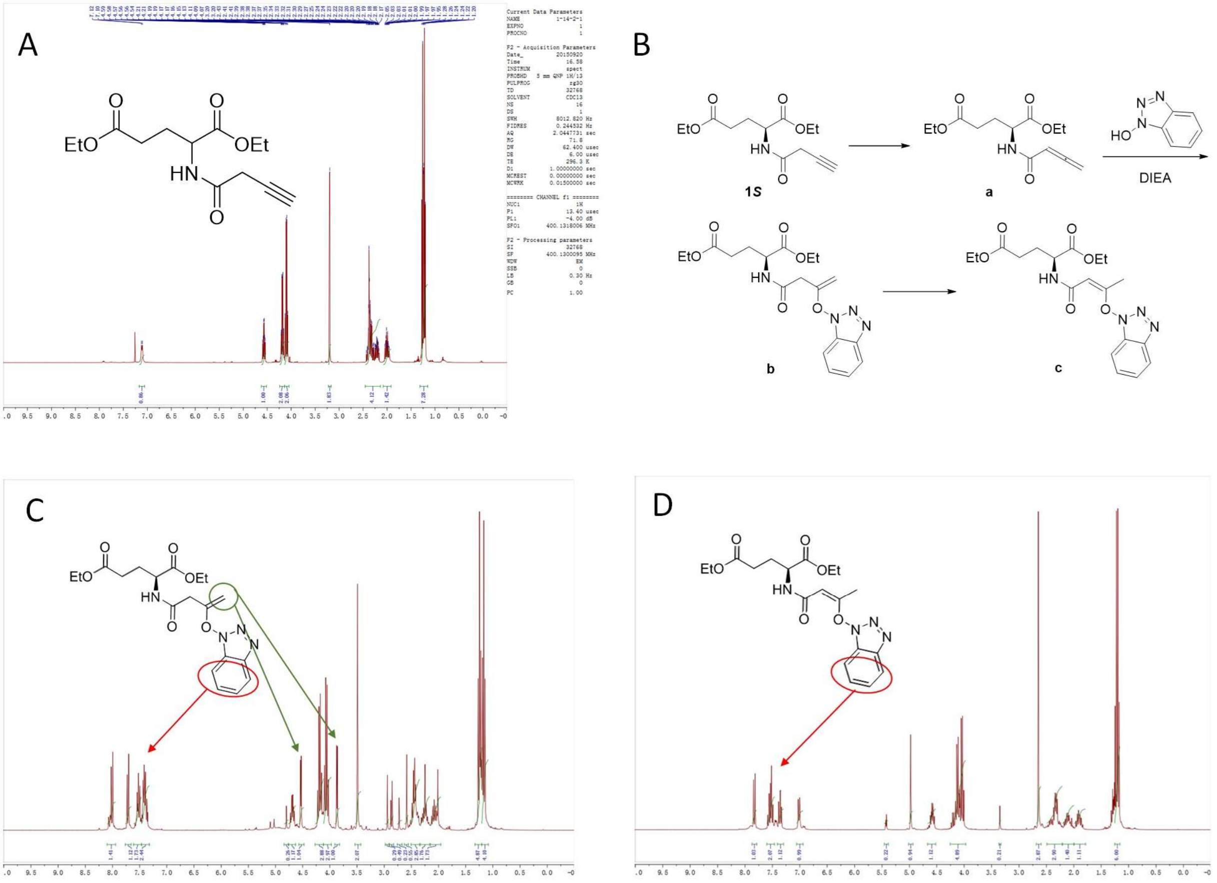

**Supplementary Figure 4** Undesired products obtained during probe 1 synthesis. A. The desired probe 1. B. proposed structure of impurities. C-D. NMR of the proposed side products. Probe 1 was prepared following a similar synthetic route to probe 2.

#### Evaluation of probes in metabolic labeling

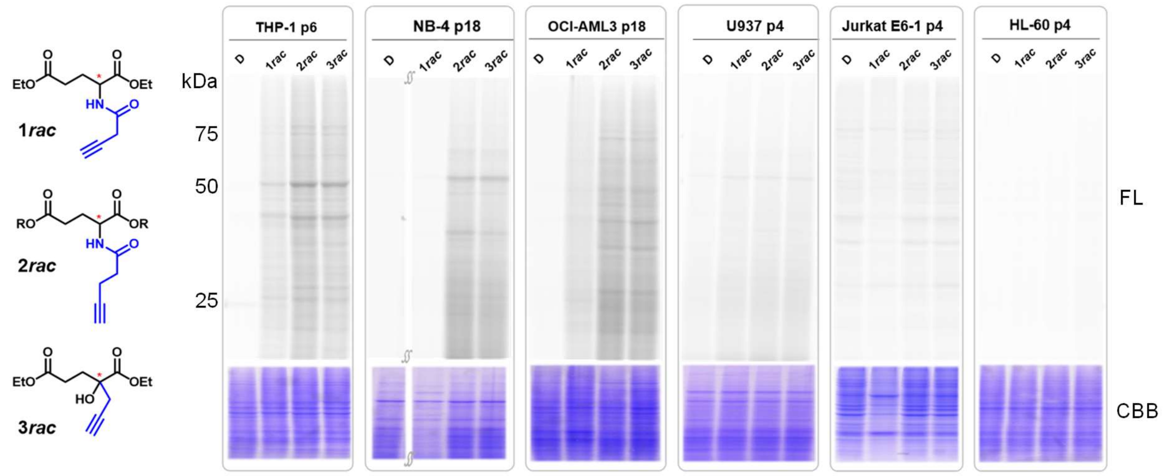

**Supplementary Figure 5** Metabolic labeling results of probes **1–3*rac*** in leukemia cell lines. Probe 1 is unstable after storage and gave decayed labeling results. Cells were cultured according to the manufacturer's recommendation and adjusted to 4–6x$10^5$ cells/mL before metabolic labeling assay with 0.5 mM probes for 4 h. THP-1, NB-4, and OCI-AML3 cells were efficiently labeled by probes, while others were tested negative under the given conditions. Left: probe structure with chiral centers indicated by red asterisks. D: DMSO control. 'p' represents the passage number of the cells. R in 2*rac* are ethyl groups.

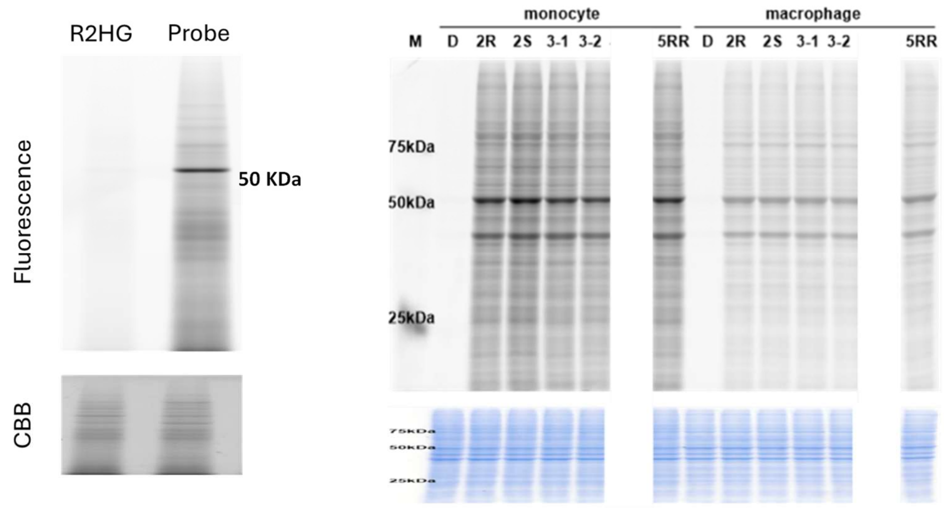

**Supplementary Figure 6** Metabolic labeling results in cord blood cells (left) and THP1 macrophages (right). THP1 macrophages were differentiated with PMA stimulation for 3 days before metabolic labeling.

THP1 cells were treated with 0.5 mM probes or *R*2HG diethyl ester for 4 h before sample collection, and cord blood cells were labeled with 0.25 mM probe for 6 h.

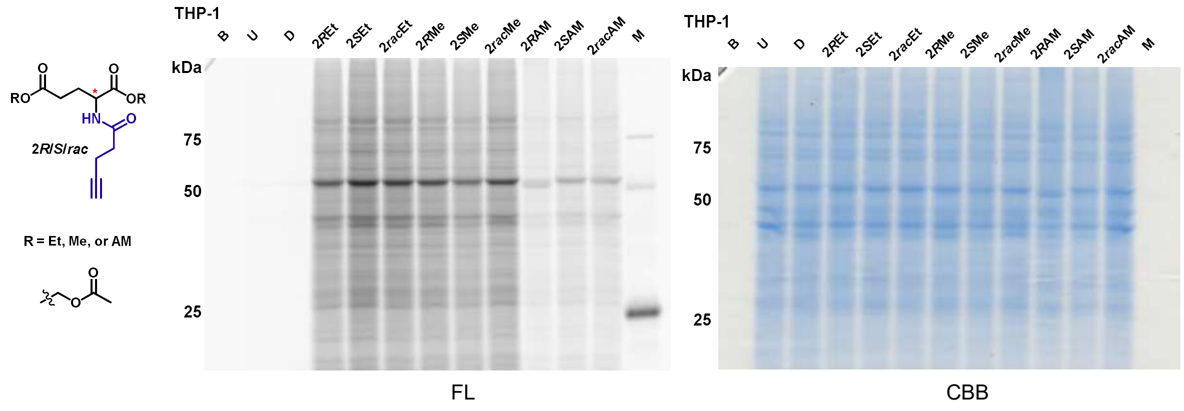

**Supplementary Figure 7** Ethyl and methyl esters label better than acetoxymethyl esters in THP-1 cells. Cells were treated with 0.5 mM probes for 4 h before sample collection. Cells were treated with 0.5 mM probes for 4 h. B: blank. U: untreated. D: DMSO control group. M: protein size marker. Et: ethyl ester. Me: methyl ester. AM: acetoxymethyl ester.

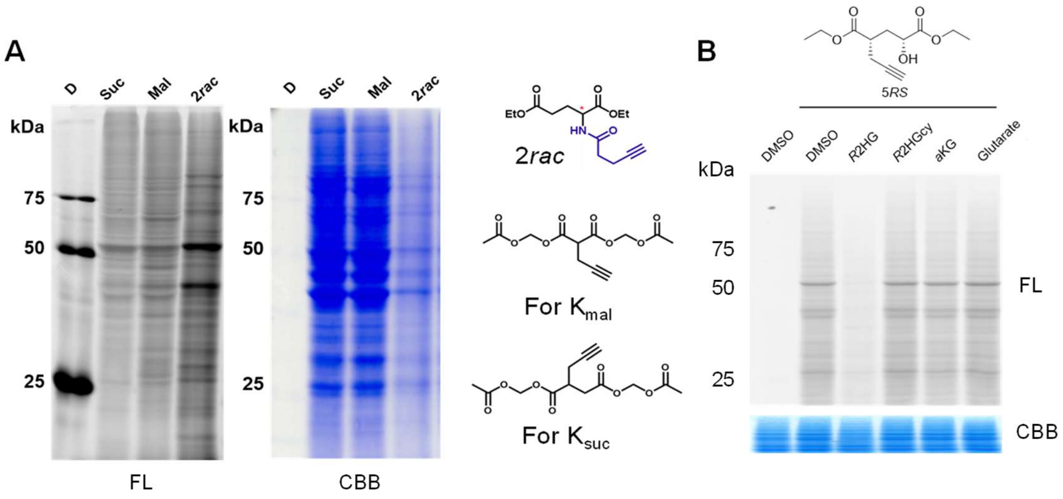

**Supplementary Figure 8** Probes are selective for *R*2HG compared to other metabolites. **A**. Comparison of metabolic labeling patterns with different probes. **B**. Competitive metabolic labeling of probe **5*RS*** in THP-1 cells. THP-1 cells were treated with 0.5 mM probes for 4 h in the presence or absence of 5 mM competitors before sample collection. *R*2HGcy: cyclic *R*2HG ethyl ester.

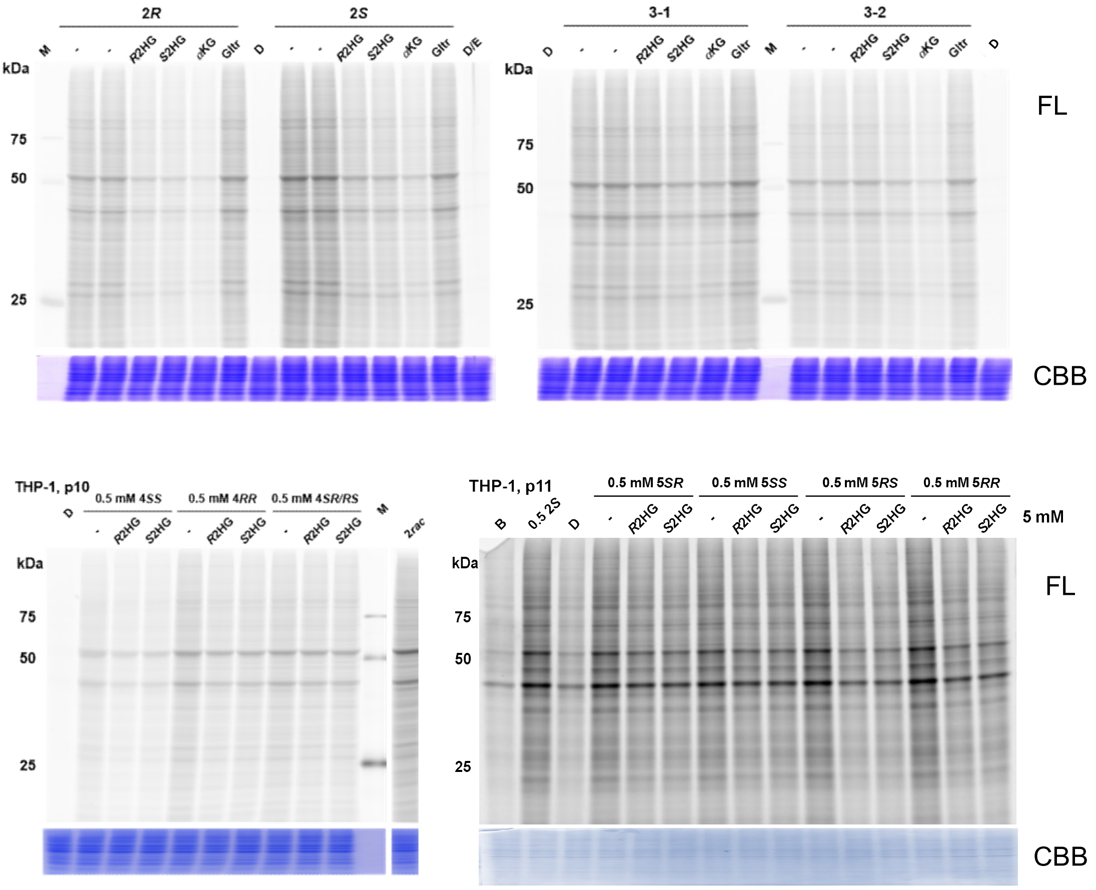

**Supplementary Figure 9** Metabolic labeling of THP-1 cells by various synthetic probes in the presence or absence of potential competitors under the same conditions as in Supplementary Figure 8B. Fluorescence/protein staining values were normalized to DMSO controls (upper) or the value of probe **2*S*** (lower) for Figure 2C. Gltr: diethyl glutarate.

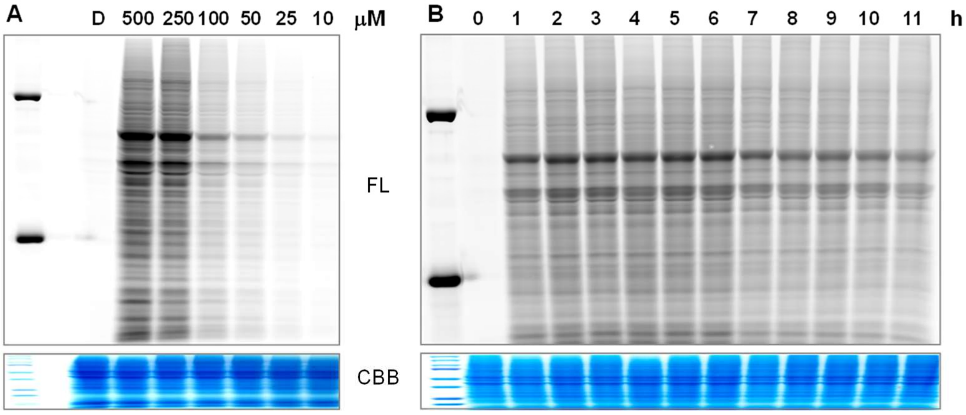

**Supplementary Figure 10** Dose- and time-dependent labeling of THP-1 cells with probe **5*RS***. **A**. Cells were treated with probe **5*RS*** at the indicated concentrations for 4 h before sample collection and visualization. **B.** Cells were treated with probe **5*RS*** at 0.25 mM for the indicated time before sample collection and visualization.

**Quantitative proteomics to profile potential *R*2HGylation targets**

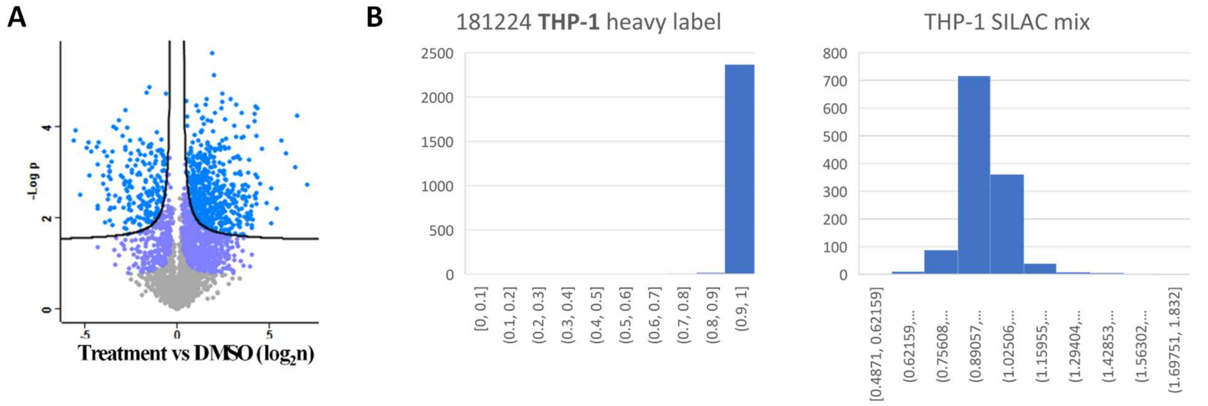

**Supplementary Figure 11** Preliminary tests before SILAC-based replaceable *R*2HGylation target profiling. A. Probe-enriched proteins with DMSO-treated controls gave >3000 and ~1000 candidates, respectively, indicating that basal (untreated) *R*2HGylation signals are too low to contribute meaningfully to the comparison. B. Sufficient isotope labeling of SILAC cells (left) and the equal mix for SILAC sample preparation (right). The distribution of heavily labeled peptides and the H/L ratio in the SILAC mixture are shown. Peptide numbers are plotted versus H/(H+L) and the normalized H/L ratio, respectively. H/(H+L) = (H/L)/[(H/L)+1].

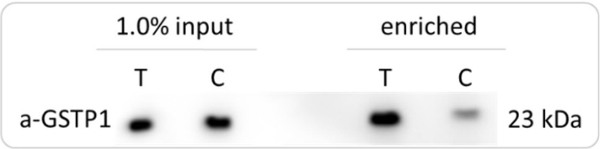

**Supplementary Figure 12** Validation of GSTP1 enrichment by synthetic probe **5*RS***. THP-1 cells were treated with 0.5 mM probe **5*RS*** in the presence (C) or absence (T) 5 mM *R*2HG for 4 h, then lysed and clicked with biotin linker for enrichment and immunoblotting.

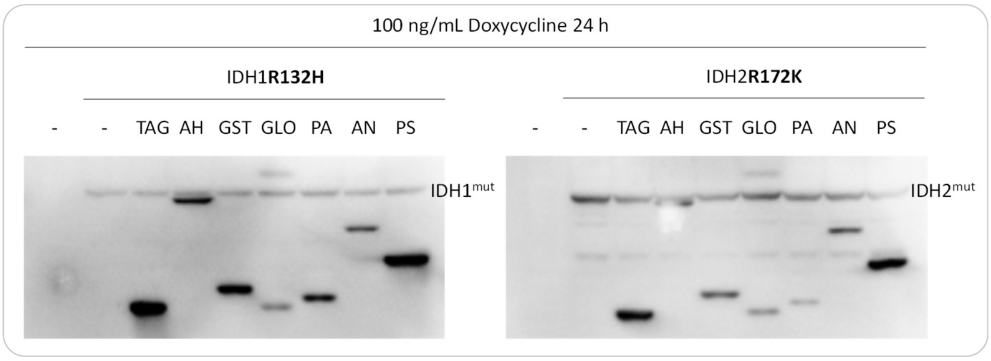

**Supplementary Figure 13** Immunoblot confirmation of the co-expression of potential FLAG-targets and FLAG-IDH1/2$^{mut}$ in HEK293T cells. Targets are (from left to right, in gene name): TAGLN2, AHCY, GSTP1, GLO1, PARK7, ANXA1, PSME1.

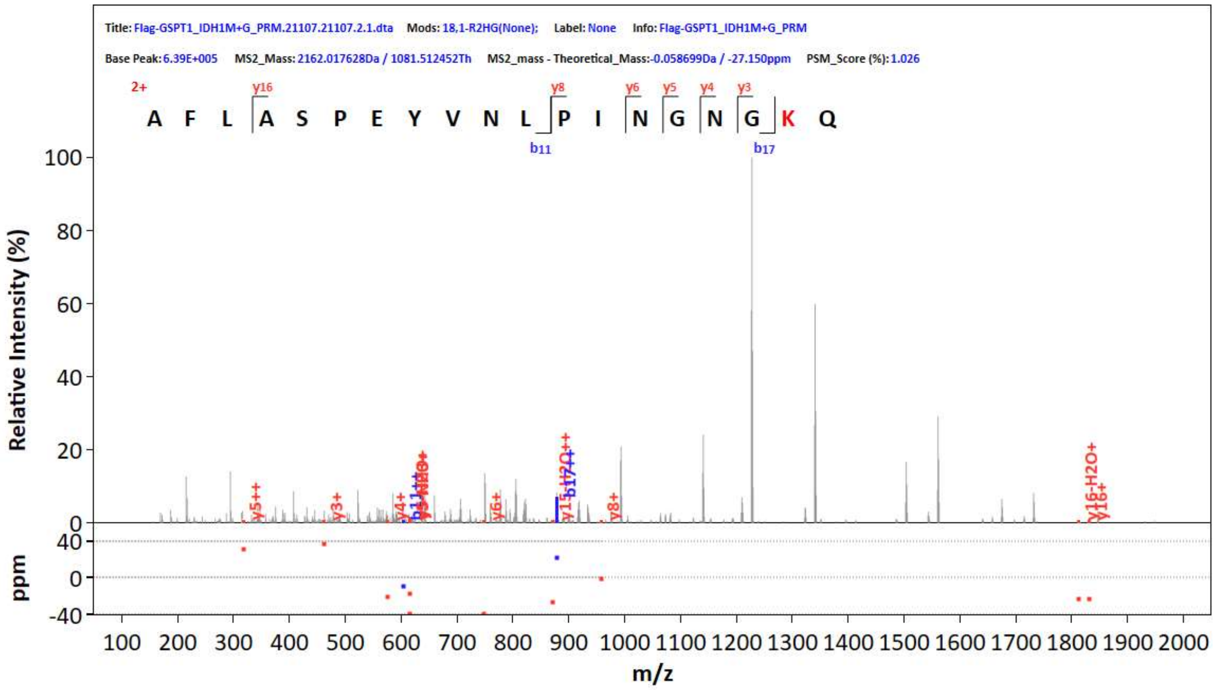

**Supplementary Figure 14** Identified $K_{R2HG}$ on immune-enriched FLAG-GSTP1 from IDH1$^{R132H}$-293T cells in preliminary screening. The parent ion m/z was set as the MS$^1$ value in PRM-MS$^2$ acquisition, which gave the spectra in Figure 3A.

**Establishment of IDH1/2-transduced cells and $K_{R2HG}$ characterization**

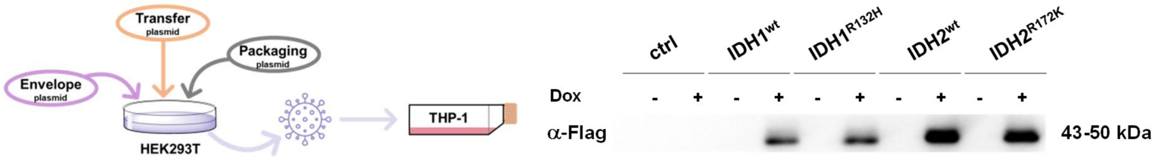

**Supplementary Figure 15** Preparation of stable IDH$^{mut}$-THP1 cells and dox-induced expression of FLAG-IDH1$^{wt}$, FLAG-IDH1$^{R132H}$, FLAG-IDH2$^{wt}$, and FLAG-IDH2$^{R172K}$ in control and transduced THP-1 cells. Cells were induced with 100 ng/mL dox for 24 h before protein harvest.

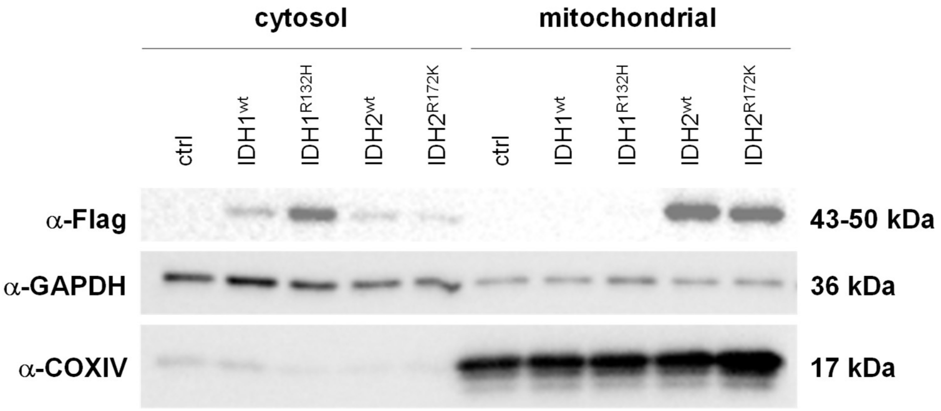

**Supplementary Figure 16** FLAG-IDH1$^{wt}$ and FLAG-IDH1$^{R132H}$ are localized in the cytoplasm. FLAG-IDH2$^{wt}$ and FLAG-IDH2$^{R172K}$ are expressed in mitochondria. Cells were induced with 100 ng/mL dox for 24 h before analysis.

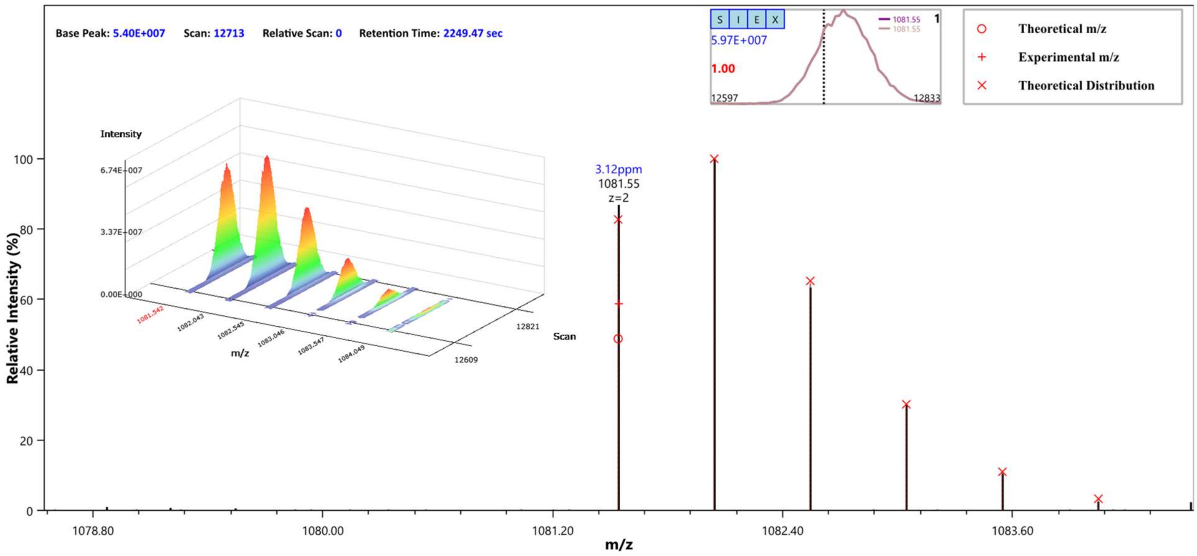

**Supplementary Figure 17** LC elution and MS[1] envelope of identified $K_{R2HG}$ peptide. Left: extracted ion current (XIC) of MS[1] isotopic peaks with parallel reaction monitoring (PRM) target defined as *m/z* = 1081.542. Upper right: LC-MS[1] integration of the targeted GSTP1 peptide with scan number = 12597−12833. Lower right: extracted MS[1] envelope of *R*2HGylated GSTP1 peptide.

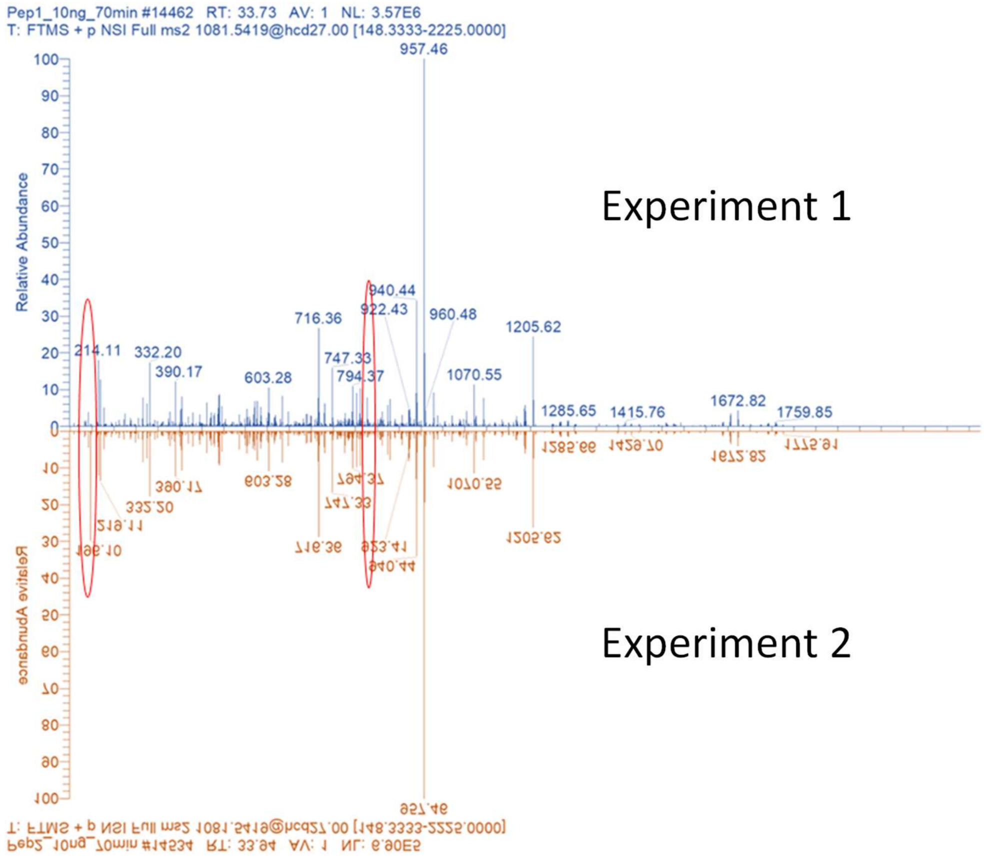

**Supplementary Figure 18** MS$^2$ spectra of C5- and C1-*R*2HGylated peptides of GSTP1 (AFLASPEYVNLPINGNGK$^{R2HG}$Q) are not distinguishable. Distinct bands are circled.

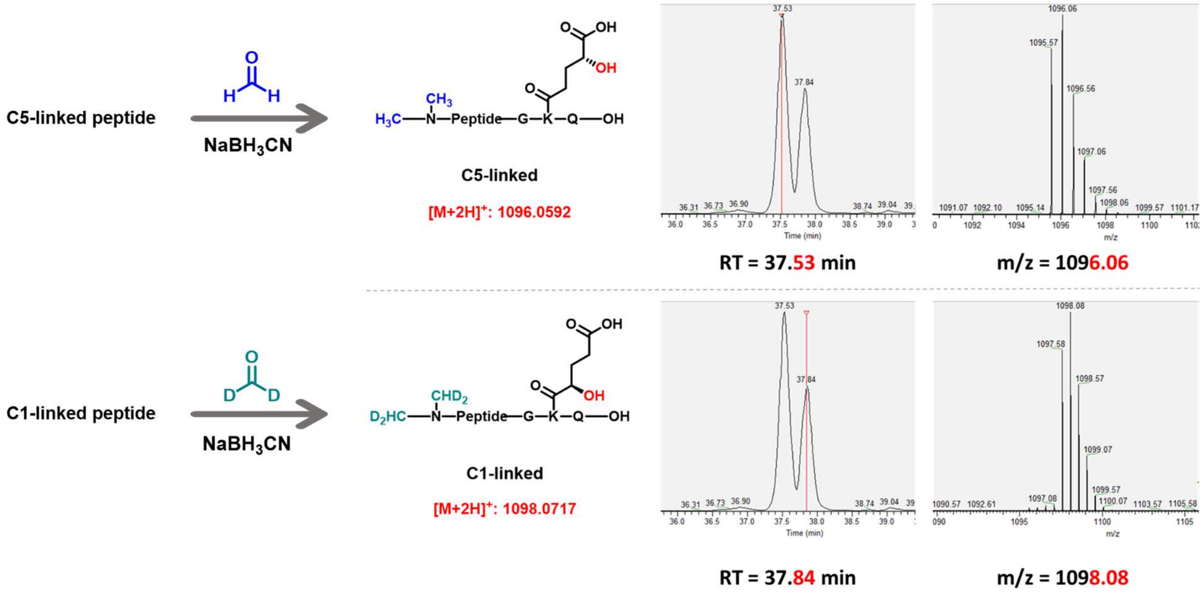

**Supplementary Figure 19** C5-*R*2HGylated peptide eluted earlier compared to C1-*R*2HGylated peptide after reductive dimethylation. The sequence of the peptide is AFLASPEYVNLPINGNGK$^{R2HG}$Q. C5-linked peptide: *m/z* [calcd] = 1096.0592, found 1096.0631, RT = 37.53 min; C1-linked peptide: *m/z* [calcd] = 1098.0717, found 1098.0760, RT = 37.84 min. Data was analysed with XCalibur software.

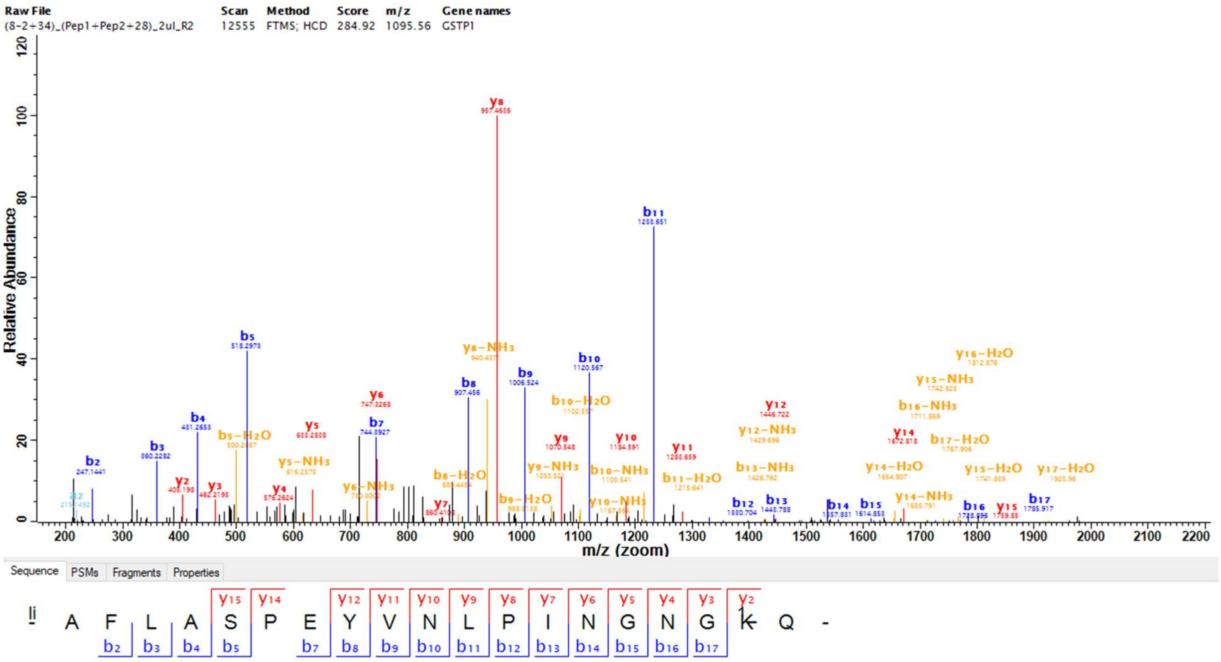

**Supplementary Figure 20** A representative MS$^2$ spectrum for a C5-linked synthetic peptide, light-labeled.

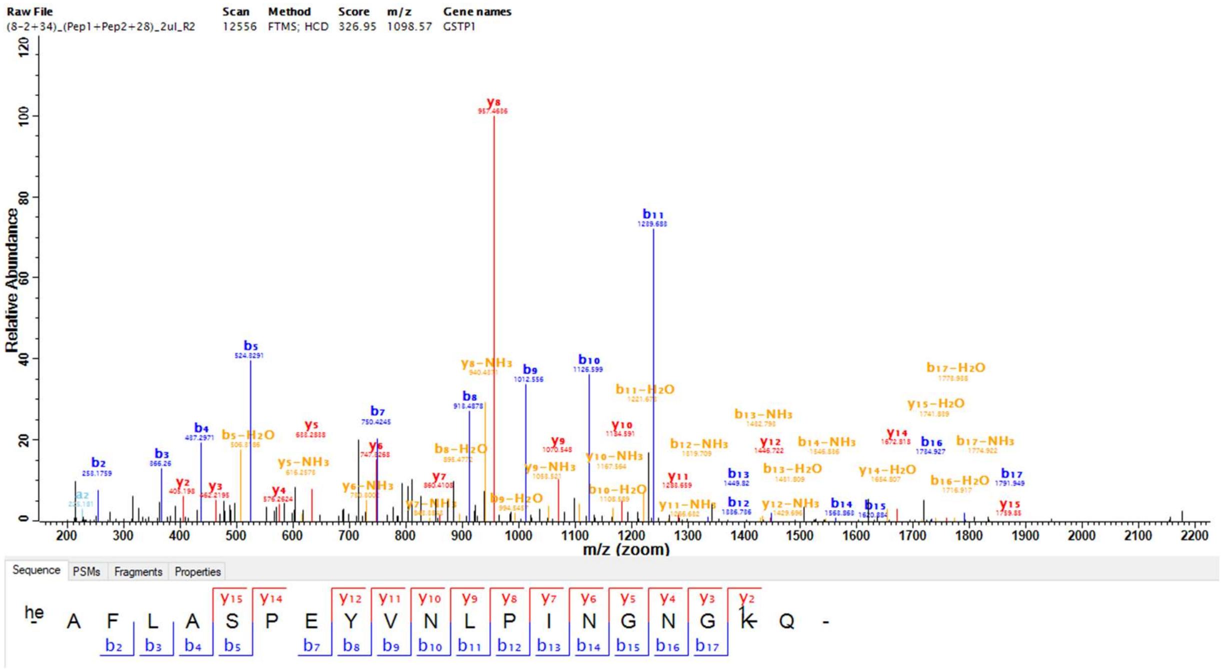

**Supplementary Figure 21** A representative MS$^2$ spectrum for a C5-linked peptide from cell lysate, heavy-labeled.

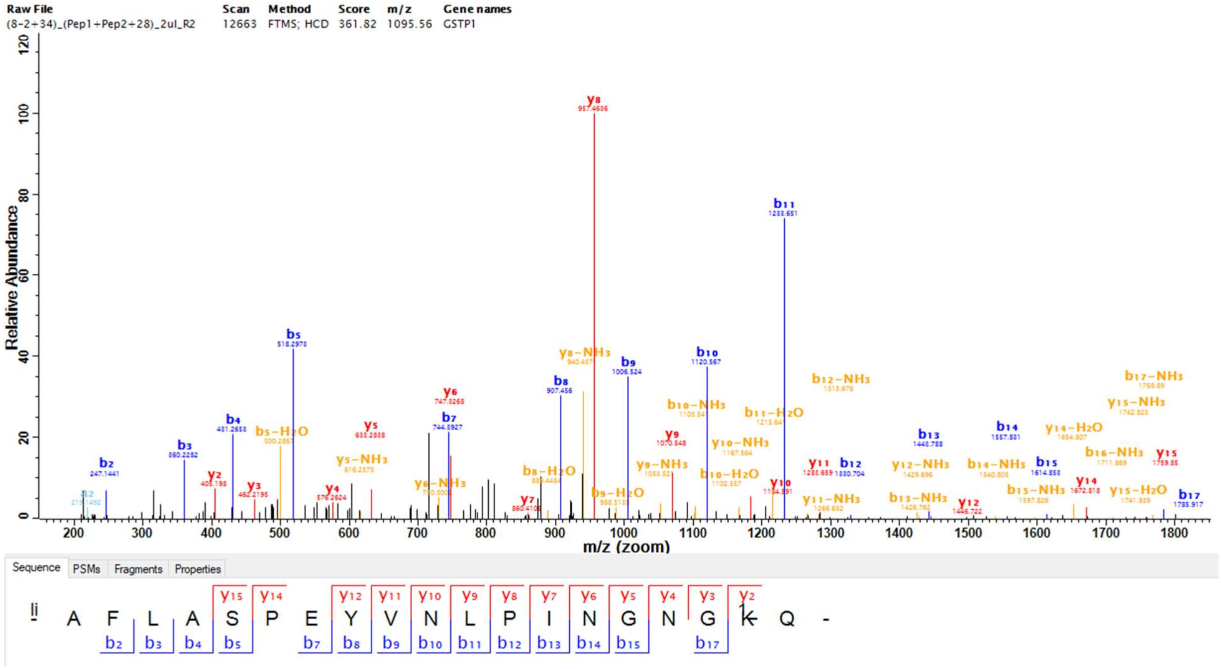

**Supplementary Figure 22** A representative MS$^2$ spectrum for a synthetic C1-linked peptide, light-labeled.

**SIRT5 act as a $K_{R2HG}$ deacylase**

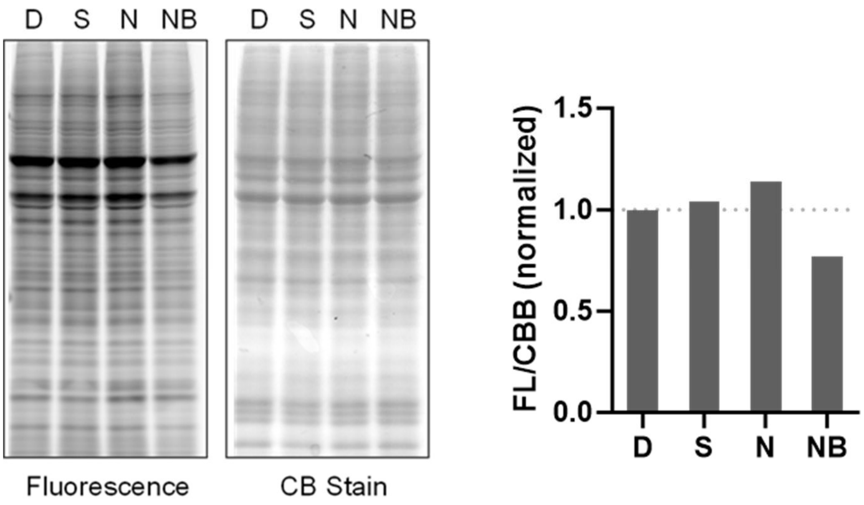

**Supplementary Figure 23** The sirtuin inhibitor nicotinamide arrests probe removal after metabolic labeling. Left: Cells were treated with 25 μM vorinostat (S, specific for class I and II HDACs), 5 mM nicotinamide (N, specific for sirtuins), 5 mM NaBut (NB, specific for class I and IIa HDACs), or same volume of DMSO (D) for 10 h before metabolic labeling with 0.5 mM **5*RS*** in the presence of HDAC inhibitors for 4 h; right: comparison of quantification results, fluorescence intensities are normalized to CBB stain intensity, and further normalized to DMSO group.

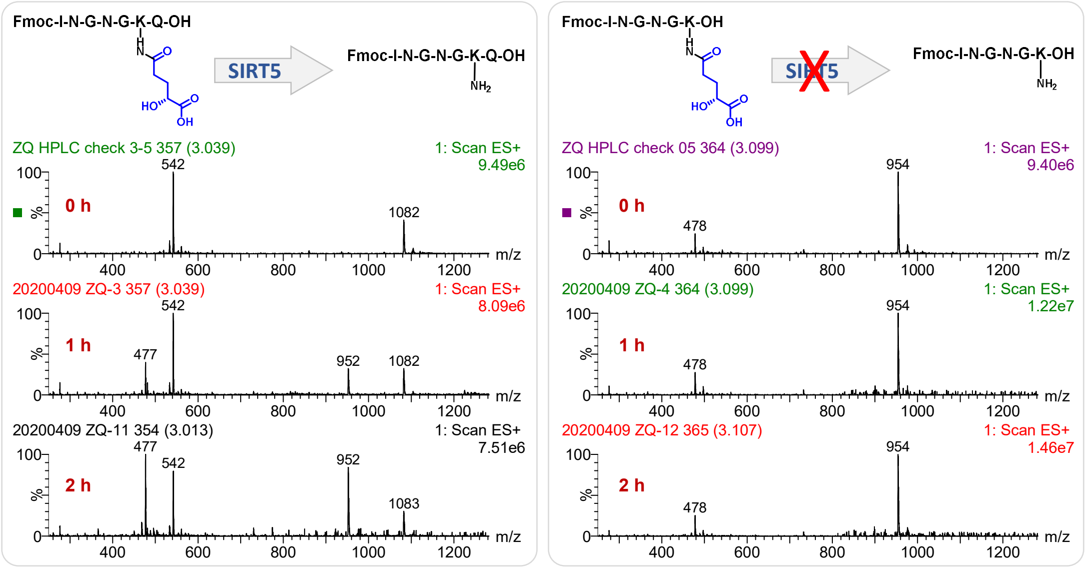

**Supplementary Figure 24** *C*-Terminal glutamine (Q) of GSTP1 peptide is necessary for SIRT5's de-*R*2HGylation activity.

**Pan-antibody for $K_{R2HG}$**

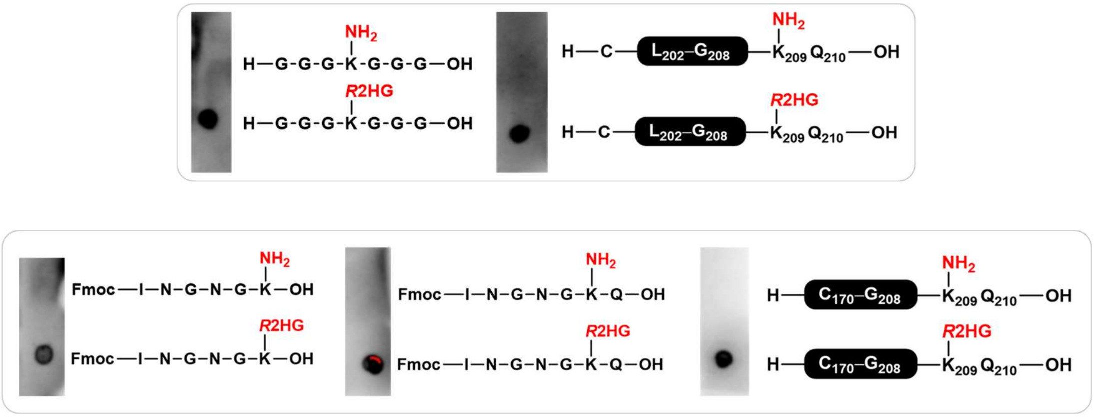

**Supplementary Figure 25** The pan-antibody is selective towards *R*2HGylated peptides compared to unmodified counterparts.

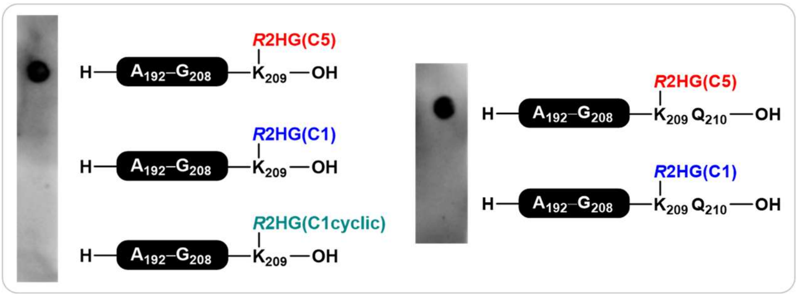

**Supplementary Figure 26** The pan-antibody is specific to C5-*R*2HGylated peptides compared to C1-*R*2HGyalation.

**Evaluation of K$_{R2HG}$ in GSTP1**

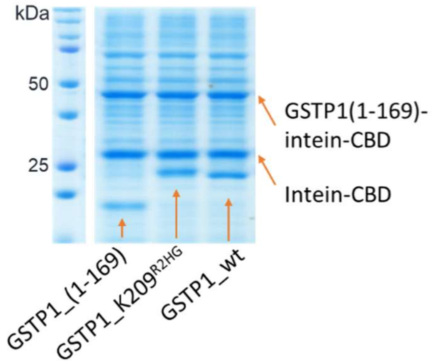

**Supplementary Figure 27** SDS-PAGE of peptide-ligated proteins in the presence of cleaved intein-CBD fragments and inactive conjugates.

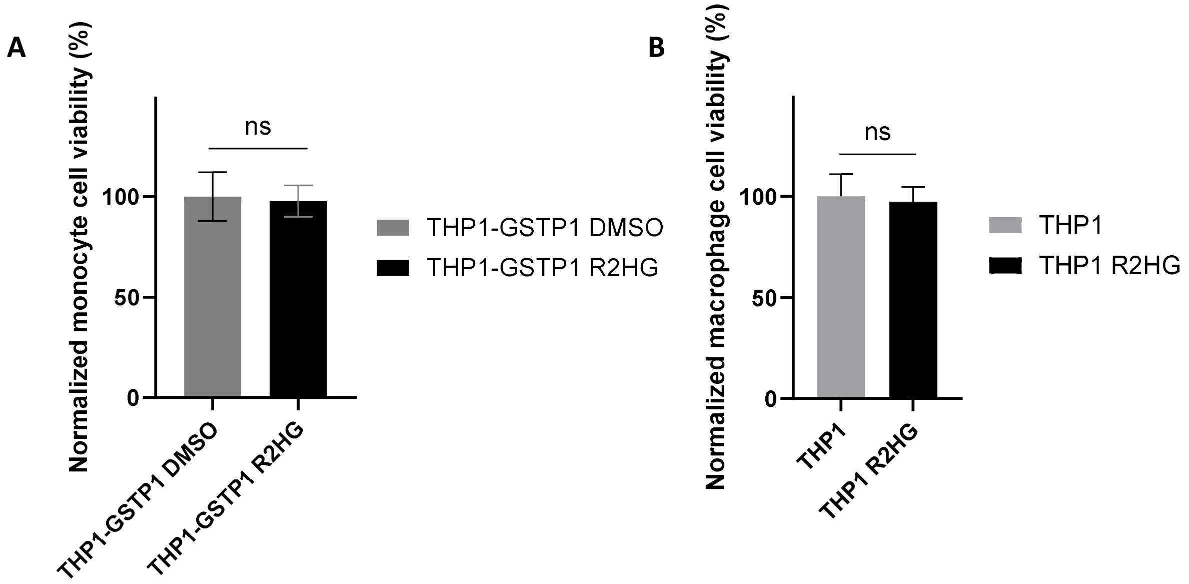

**Supplementary Figure 28** *R*2HG (0.1 mM) does not impair the proliferation of wt-GSTP1-expressing THP1 cells (A) or the differentiation of wt THP1 cells (B).

**Biological methods**

**In-gel visualization of metabolically labeled proteins**

For suspension cells pre-seeded at a density of $5\times10^5$ cells/mL, the probe in 1/4 volume of cell suspension was freshly prepared via sonication, added into the cell suspension with gentle swirling to make a final concentration of 0.5 mM if not specified. For adherent cells pre-seeded at 70–80% confluence, the old medium was half removed, followed by the addition of freshly prepared probe-supplemented medium of the same volume to give a final concentration of 0.5 mM probe. Treatment was done in a humidified incubator at 37 °C, under 5% $CO_2$, for 4 h if not specified.

All lysis steps were done on ice. Suspension cells were collected in Falcon tubes, centrifuged at a speed of 1,000 g for 5 min, transferred to 1.5 mL Eppendorf tubes, and washed with PBS three times before lysis with buffer A (1% SDS, 150 mM NaCl, 50 mM HEPES, 2 mM $MgCl_2$, 10% glycerol, pH 7.5, freshly supplemented with EDTA-free Roche protease inhibitor cocktail and Benzonase®) via vortex. Cell lysates were centrifuged at 6,000 g for 15 min at 4 °C. The supernatant was collected in new Eppendorf tubes as clear lysates.

Clear cell lysates were diluted to 1 mg/mL with buffer A, and reacted with 0.1 mM rhodamine azide, 1 mM TCEP, 0.1 mM TBTA, and 1 mM $CuSO_4$ via lid-addition (an operation of adding reagents to lids of EP tubes and then inverting to start the reaction simultaneously). After incubation at room temperature for 1.5 h with continuous shaking, the mixture was precipitated with ice-cold acetone at −20 °C overnight.

Protein precipitates were collected via centrifugation at 6,000 g for 15 min at 4 °C, followed by careful washing with ice-cold methanol twice. Air-dried pellets were re-suspended in LDS/SDS loading buffer supplemented with 50 mM DTT, heated at 85 °C for 8 min, and loaded onto an SDS gel (protein ladder 1/10 diluted to reduce fluorescence interference). SDS-PAGE was done at 80 V for 20 min and then 120–150 V for about 1 h using 10% or 12% polyacrylamide gels.

After electrophoresis, the SDS gels were cropped, washed twice with Milli Q water, and fixed in fixation buffer (10% AcOH + 40 % $H_2O$ + 50 % MeOH) for 15 min with gentle shaking. Then, the gel was rinsed twice with $H_2O$ and scanned with Typhoon Scanner at 532 nm excitation and 572 nm emission, within energy levels of 500–900 photomultiplier tube (PMT) voltage. After scanning, the gels were stained with stain buffer (1% Coomassie blue in fixation buffer) for 2 h and destained with fixation buffer twice for 10 min and 2 h, respectively. The gels were then soaked in water overnight to recover their native size and scanned for protein loading control.

Fluorescence intensity (FL) and protein loading intensity (CB) were quantified using ImageJ. FL from DMSO control was subtracted from labeled groups and then divided by the corresponding CB value. To make results consistent among different gels, (FL-FL(D))/CB were normalized to either DMSO groups (results with probe series **2**–**4**) or probe **2*S*** groups (results with probe series **5**, considering the slightly higher background signal with new chemicals and platforms) and plotted for comparison.

**MS-based proteomic quantification**

THP-1 cells were cultured in a T75 flask at a density of $5x10^5$ cells/mL in RPMI 1640 medium supplemented with 10 % FBS and 1% penicillin/streptomycin. For SILAC experiments, plain SILAC media containing dialyzed heat-inactivated FBS, L-Arg (unlabeled or 13C6, 15N4), and L-Lys (unlabeled or 13C6, 15N2) were applied until satisfactory isotopic labeling before experiments. After recovering overnight, cells were treated with 0.5 mM probe for 4 h at a density of $4x10^5$ cells/mL. After metabolic labeling, cells were collected in Falcon tubes via centrifugation, then transferred to 1.5 mL Eppendorf tubes and washed with cold PBS three times. The obtained cell pellet was then lysed on ice with lysis buffer A freshly supplemented with protease inhibitor cocktail and Benzonase$^{@}$, followed by lysate clearance via centrifugation at 6,000 g for 15 min. No less than 2 mg of proteins were harvested and used in affinity enrichment.

Cleared lysates were adjusted to 1 mg/mL after BCA measurement of concentration, followed by click chemistry with biotin azide (AZO or DADPS, 100 μM) in the presence of TCEP (1 mM), TBTA (100 μM), and $CuSO_4$ (1 mM). The mixture was rotated at rt for 1.5 h, avoiding light, followed by precipitation with 8 mL ice-cold acetone for each 2 mg input.

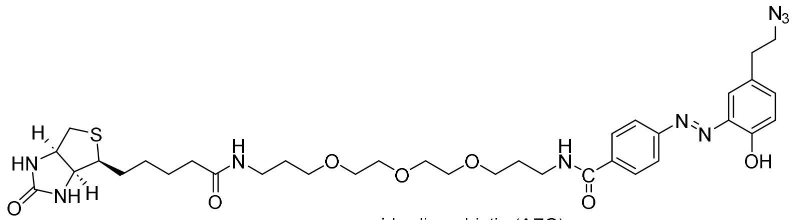

azido-diazo-biotin (AZO)

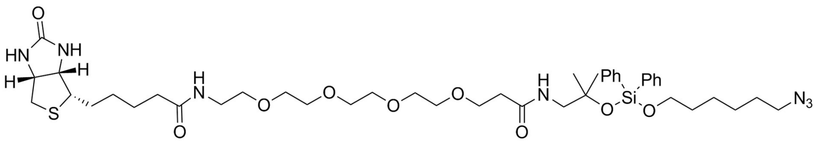

DADPS linker

For each 2 mg input, proteins were then pelleted at 3500 g for 8 min at 10 °C and washed with 7 mL ice-cold methanol twice. Pellets were air-dried for 10 min and dissolved in 2 mL PBS with 3% SDS and 6 M urea. Protein solutions were then diluted with 12 mL PBS and transferred to 50 mL Falcon tubes containing 200 μL PBS-washed beads. The mixture was then rotated at rt for 1.5 h. After that, beads were transferred into Bio-spin columns, washed with 0.2 % SDS in PBS, 0.1 % SDS in PBS with 6 M urea, and 0.05 % SDS in 0.25 M aqueous $NH_4HCO_3$ solutions five times, respectively. Alternatively, for biotin resin from GE company, binding was done in 20 mM Tris with 100 mM $Na_2HPO_4$, 150 mM NaCl, 1.6 M urea, and 0.1% SDS at pH = 7.5, followed by three times washing with 100 mM Tris with 0.1% SDS, and 6 M urea at pH = 7.5.

Proteins on streptavidin beads were suspended in PBS with 6 M urea, reduced with 10 mM TCEP at 65 ºC for 30 min, and alkylated with 20 mM IAA for 30 min at rt in the dark. The resulting resin was washed with PBS, followed by the addition of 2 μg trypsin solution in 100 μL 25 mM ammonium bicarbonate. Released peptides were desalted with StageTip and vacuum concentrated for MS analysis. For trials with AZO tag, peptides on resin were cleaved with 50 μL 25 mM $Na_2S_2O_4$, 0.05% SDS in 0.25 M ABC aqueous solution twice, followed by a StageTip desalting step. For DADPS linker-enriched peptides, the cleavage step was done with a 5% FA aqueous solution. Peptides from either linker were then concentrated under vacuum before LC-MS/MS analysis.

The obtained peptides were then dissolved in 0.1% formic acid (mobile phase A) and separated chromatographically on an Easy-nLC 1000 system (Thermo Fisher Scientific) equipped with a home-made analytical column (100 μm inner diameter silica-fused capillary tube [Molex Polymicro] packed with 20 cm of 1.9 μm/120 Å C18-AQ beads [Dr. Maisch]). Peptides were eluted at a constant flow rate of 250 nl/min using an effective linear gradient from 7% to 35% mobile phase B (0.1% formic acid in acetonitrile) over 120 min. Separated peptides by online nano LC were analyzed on an Orbitrap Fusion Tribrid mass spectrometer (Thermo Fisher Scientific) via nano-spray ionization. The survey scans covering *m/z* range of 350 to 1,550 were performed in the Orbitrap analyzer using resolution of 120,000 (at *m/z* 200), maximum injection time (MIT) of 100 ms, and automatic gain control (AGC) target value of $2 \times 10^5$. The tandem MS spectra were acquired using a data-dependent acquisition (DDA) method at Top Speed mode with 3 s cycles and a dynamic exclusion duration of 30 s. Precursors were isolated through the quadrupole using a 1.6 Da window, and then fragmented by HCD using normalized collision energy (NCE) of 30%. Fragment ions were scanned in the Orbitrap analyzer at a resolution of 15,000, MIT of 45 ms, and AGC target value of $5 \times 10^4$.

DDA files for quantification were searched against the Uniprot Human protein database using MaxQuant (v1.6.2.3)[1] and pFind (v3.1.5)[2]. *R*2HG was added as a dynamic modification, with other settings as default.

**K$_{R2HG}$ identification from IDH1/2$^{mut}$ cells**

THP-1 cells stably expressing FLAG-tagged IDH1/2 and their mutants were generated via lentiviral transduction. Briefly, pSLIK-IDH1-FLAG, pSLIK-IDH1-R132H-FLAG, pSLIK-IDH2-FLAG, and pSLIK-IDH2-R172K-FLAG were acquired from Addgene as a gift from Christian Metallo (Addgene plasmid # 66802; #66803; # 66806; # 66807). HEK293T cells were co-transfected with acquired plasmids along with 3rd generation lentiviral packaging plasmids pMDLg/pRRE and pRSV-Rev, as well as the envelope plasmid pMD2.G, as provided by Prof. Tian's group, to produce lentivirus. After transfection for 10 hours, the medium containing Lipofectamine 3000 reagents and plasmids was replaced with fresh medium, and the cells were cultured for an additional 24 hours to allow for virus production. The medium containing lentiviral particles was collected 48 hours later and filtered with a 0.45 μm sterile filter to infect THP-1 cells for 3 days. Infected cells were recovered 24 hours before being selected with 350 μg/mL hygromycin for ten days. Protein expression was checked by inducing transfected cells using 0.1 μg/mL doxycycline hyclate (dox, Sigma) for 24–48 hours. Cell lysates were then processed under similar conditions for MS-based quantification and analysis.

**Solid-phase peptide synthesis for control peptides**

The 2-chlorotrityl chloride resin (100 mg, loading: 1 mmol/g) was swollen in 1 mL dry DCM for 20 min in a 5 mL disposable syringe equipped with a porous polypropylene disc at the bottom. A solution of Fmoc-Gln(trt)-OH (122 mg, 0.2 mmol) and DIEA (70 μL, 0.4 mmol) in DCM was added, and the reaction vessel was shaken on the vortex at room temperature for 1 h. The resin was washed with DMF, then merged in a solution of DCM/MeOH (17:2:1, v/v/v, 1 mL) for 20 min and washed with DMF.

The Fmoc deprotection was carried out using 1 mL of 20% piperidine in DMF at room temperature for 20 min. The resin was then washed with DMF. For the coupling step, a solution of Fmoc amino acid (4 equiv), HATU (4 equiv), and DIPEA (8 equiv) in DMF was gently agitated on a vortex with the resin at room temperature for 1 h. The resin was then washed with DMF. The procedures were repeated until the coupling of the last amino acid before further deprotection of the *N*-terminal Fmoc group. Lysine of the R2HGylation site was protected with alloc, with other reactive residues protected by tBu-, Trt-, Pbf-, and Boc-groups if not specified.

The resin with Fmoc-protected peptides was dried under vacuum to calculate the yield based on increased weight and peptide molecular weight. Then, the resin with the peptide was swollen in DCM for 20 min and charged with argon. $PhSiH_3$ (0.16 mL, 1.3 mmol) in dry DCM (1 mL) was added to the resin by syringe, followed by the addition of a solution of $Pd(PPh_3)_4$ (10 mg, 8.7 μmol) in dry DCM (1 mL). After 1 h, the resin was washed with DCM and re-deprotected with the above procedures to remove Alloc on the lysine residue.

The deprotected peptide was resuspended in dry DCM, followed by the addition of *R*2HG C5- or C1-donor (2 equiv) and DIPEA (4 equiv). The reaction was vortexed overnight and then washed with DCM.

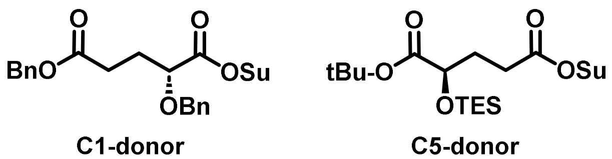

Peptides were subjected to 1 mL of 20% piperidine in DMF at room temperature for 20 min. After Fmoc deprotection, resins were washed with DMF and DCM, followed by the addition of 1 mL of acidic cleavage cocktail of DCM/AcOH/trifluoroethanol (8/1/1, v/v/v) for 1 h twice. Following filtration, peptides in the resulting cleavage solutions were precipitated in ice-cold ether, pelleted via centrifugation, washed with ether, purified via preparative-HPLC, further deprotected under standardized conditions, and purified.

**Reductive demethylation of endogenous and exogenous peptides**

For dimethyl labeling, 0.1 mL peptide solution (1 mg/mL) was reacted with 50 μL of 4% “light” formaldehyde and “heavy” formaldehyde ($CD_2O$), respectively, in 0.9 mL phosphate buffer containing 0.2 mL 50 mM $NaH_2PO_4{\cdot}2H_2O$ and 0.7 mL 50 mM $Na_2HPO_4{\cdot}12H_2O$. The resulting solution was treated with 50 μL 0.6 M sodium cyanoborohydride ($NaCNBH_3$) and incubated at rt. for 1 h. The reaction was quenched by adding 16 μL 1% ammonia and 8 μL 5% formic acid. All peptides were individually analyzed for concentration adjustment based on their XIC integration intensity, after which the “light” and “heavy” samples were combined for coelution and LC-MS-based structural confirmation.

**MS-based peptide identification and characterization**

For PTM structural determination, synthetic peptides were reductively demethylated as reported in the literature[3]. Peptides were dissolved in 0.1% formic acid (mobile phase A) and separated chromatographically on an Easy-nLC 1000 system (Thermo Fisher Scientific) equipped with a home-made analytical column (100 μm inner diameter silica-fused capillary tube [Molex Polymicro] packed with 20 cm of 1.9 μm/120 Å C18-AQ beads [Dr. Maisch]). Peptides were eluted at a constant flow rate of 250 nl/min using an effective linear gradient from 7% to 35% mobile phase B (0.1% formic acid in acetonitrile) over 120 min. Separated peptides by online nano LC were analyzed on an Orbitrap Fusion Tribrid mass spectrometer (Thermo Fisher Scientific) via nano-spray ionization. Under PRM scanning mode, the *m/z*s of the desired dimethylated synthetic and target *R*2HGylated GSTP1 peptide were programmed for scanning. The PRM acquisition method was developed as described in a previous study[1] by combining a survey scan followed by up to 20 PRM scans (for IPed samples) of precursor ions. The survey scans were collected from *m/z* 400 to 1,300 at a resolution of 60,000 (at *m/z* 200), MIT of 100 ms, and AGC target of $2 \times 10^5$. Target precursors were then isolated through a window of 1.6 Da, followed by fragmentation at NCE of 30%. The product ions within the *m/z* range of 200-2,000 were scanned with an Orbitrap resolution of 60,000, MIT of 150 ms, and AGC target of $1 \times 10^5$.

PRM LC-MS[1]for $K_{R2HG}$ confirmation was generated from Xcalibur (v4.0.27.10, Thermo), through base peak selection with targeted *m/z* range, with mass tolerance of 5 ppm and mass precision at 4 decimal points.

**SIRT5 deacylation assay**

The enzymatic activities of human SIRT1−7 were measured by LC-MS. SIRT (5 mM for 1−3 and 5−7, 2.5 mM for 4) was incubated with 500 mM synthetic peptide in 50 mM Tris (pH 8.0) buffer containing 150 mM NaCl, 2 mM $NAD^+$, 1 mM DTT (and salmon sperm DNA for SIRT7) at 37 °C for 1−2 h. The reactions were stopped by adding 1/3 (v/v) of 20% TFA. Samples were then desalted with Stage-Tips[2] and analyzed by LC-MS.

**Intein-mediated protein ligation for GSTP1 synthesis**

The coding sequence of *N*-terminal GSTP1 was PCR-amplified from GSTP1 cDNA clone (Sino Bio) and recombined to double-enzyme digested pTXB1 backbone (NheI, SapI). The sequence of GSTP1(1-169)-intein-CBD was confirmed via Sanger sequencing.

Primers:

pTXB1-GSTP1-F: AGAAGGAGATATACATATGCCGCCCTACACCGTG

pTXB1-GSTP1-R: TGCATCTCCCGTGATGCAGCCAGGGGCTAGGAC

The plasmid coding truncated GSTP1 in-frame with intein and chitin-binding domain (CBD) was transformed into Rosetta (DE3) *E. coli* cells. Bacteria cultured to $OD_{600}$ = 0.6 were cooled and cultured in the presence of 1 mM IPTG at 16 °C for 16 h. Precipitated cell pellet was lysed with 20 mM HEPES (supplemented with 0.5 M KCl, 1 mM EDTA, and 0.1% v/v Triton X-100 or Tween20, adjusted to pH = 7.4, freshly added Roche protease inhibitor cocktail) via sonication, cleared via centrifugation, and affinity enriched by pre-washed chitin resin (NEB). The cleavage was done in the same buffer without Triton X-100, in the presence of 200 mM MESNA, at room temperature for 72 h.

The peptides for ligation were prepared similarly to those described for PTM identification control peptides. Ligation was done *in situ* at a concentration of 1 mM upon cleavage of expressed GSTP1_1-169.

**GSTP1 enzymatic assay (CDNB-GSH conjugation)**

The GSTP1 enzymatic assay was performed in 0.2 M sodium phosphate buffer at pH = 6.5. To the 50 µL buffer were added 5 µL of 20 mM freshly prepared GSH in deionized water and 5 µL of *in situ*-ligated enzyme solution, followed by the addition of 40 µL phosphate buffer containing 5 µL 20 mM CDNB. The 96-well plate was monitored spectrophotometrically by an increase in absorbance at 340 nm with shaking.

**THP1 viability and differentiation assay**

For *R*2HG's effect on THP1 proliferation, cells were seeded into 96-well plates at a density of $2\times10^4$ cells per well and incubated with 0.1 mM *R*2HG diethyl esters for 48 hours. After that, the cell viability was measured with the Cell Counting Kit-8 (CCK-8) assay. For differentiation assays, THP-1 cells were treated with 10 ng/mL phorbol 12-myristate 13-acetate (PMA) for 48 hours to induce differentiation. Suspension monocytes were washed away, and adherent macrophages were assayed with the CCK-8 kit. For *R*2HG-impaired differentiation, cells were co-treated with 0.1 mM *R*2HG diethyl ester. The results were collected in triplicate and processed with GraphPad Prism.

**Synthesis and Characterization**

All reagents and solvents were used as received from commercial sources unless otherwise specified below. Anhydrous tetrahydrofuran (THF), diethyl ether ($Et_2O$), toluene, and dichloromethane (DCM) were collected from a PureSolv MD Solvent Purification System made by Innovative Technology. Anhydrous triethylamine ($Et_3N$) and acetonitrile (MeCN) were distilled from calcium hydride (CaH) before use. All anhydrous reactions were carried out in oven-dried flasks under an Ar atmosphere unless otherwise noted. Air- and moisture-sensitive reagents were manipulated via syringes through rubber septa. Reactions were monitored by analytical thin-layer chromatography on glass silica gel plates (Merck silica gel 60, 0.25 mm layer thickness) with fluorescent indicator from Merck KGaA. After developing, TLC plates were either illuminated with a UV lamp (254 nm) or stained in an appropriate manner. Staining solutions were made by following standard recipes. Flash chromatography was carried out on silica gel 60 (particle size of 0.040–0.063 mm, from various commercial sources) and eluted with solvents specified.

Melting points were measured by Melting Point SMP1 from Stuart Scientific Co., LTD. Nuclear magnetic resonance (NMR) spectra were recorded in deuterated solvents on Bruker Avance Fourier Transform Spectrometers (300 MHz, 400 MHz, and 500 MHz for proton, 75 MHz, 100 MHz, and 125 MHz for carbon, respectively) at room temperature. NMR spectra were normalized by calibrating appropriate peaks of solvents employed (7.26 ppm for $^1H$ NMR, 77.16 ppm for $^{13}C$ NMR in $CDCl_3$; 3.34 ppm for $^1H$ NMR, 49.86 ppm for $^{13}C$ NMR in $CD_3OD$), and reported in ppm. High-resolution mass spectra (MS) were recorded on a Thermo Scientific DFS High Resolution Magnetic Sector MS. Enantiomeric excesses were measured through normal phase high performance liquid chromatography (HPLC) with CHIRALCEL® AD-H and OF columns. Optical rotations were recorded on a Bellingham Stanley ADP440+ polarimeter. X-ray diffraction data were collected on Bruker D8 VENTURE FIXED-CHI PHOTON 100 CMOS.

**Probe series 2**

Considering the instability of the butynoic acid moiety in probe **1** series, we improved the structure to probes **2*R***, **2*S***, and **2*rac*** via amide coupling with the stable 4-pentynoic acid **2.3** (Scheme 4.2), which provided the desired products in satisfactory yields, with enantiopurity of both **2*R*** and **2*S*** confirmed by chiral HPLC.

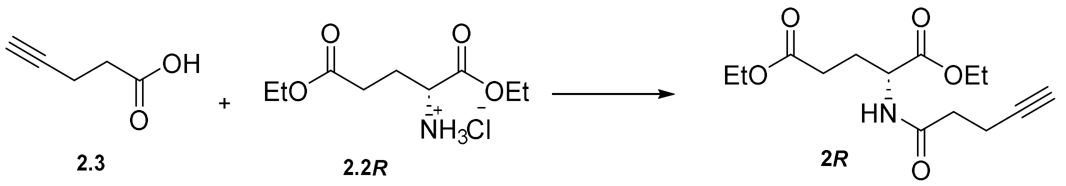

To a solution of 4- pentynoic acid (98.1 mg, 1 mmol) and diethyl *R*-glutamate **2.2*R*** (479.4 mg, 2 mmol) in anhydrous DMF (3.6 mL) were added EDCI (421.7 mg, 2.2 mmol), HOBt (297.3 mg, 2.2 mmol), and DIEA (696.7 µL, 4 mmol) sequentially under an Ar atmosphere. The reaction mixture was stirred for 24 h under an Ar atmosphere. Upon completion, the reaction mixture was diluted with DCM (to 30 mL), washed with 3M HCl, saturated $NaHCO_3$, water, and brine, and dried with anhydrous $Na_2SO_4$. The mixture was filtered and concentrated under reduced pressure. The resulting residue was azeotroped with *n*-hexane to get rid of residual DMF. Purification by column chromatography using 1/2 (v/v) EtOAc/*n*-hexane afforded **2*R*** as a yellowish liquid (204 mg, 72% yield). **2*S*** and **2*rac*** were similarly prepared, with yields of 74% and 79%, respectively.

**2*R***

Analytical TLC, EtOAc/*n*-hexane = 1/1 (v/v), $R_f$ = 0.4; $^1$H NMR (400 MHz, $CDCl_3$) δ 6.66 (d, *J* = 7.9 Hz, 1H), 4.53 (td, *J* = 8.0, 5.1 Hz, 1H), 4.17 – 4.07 (m, 2H), 4.04 (q, *J* = 7.2 Hz, 2H), 2.52 – 2.20 (m, 6H), 2.18 – 2.05 (m, 1H), 1.97 – 1.83 (m, 2H), 1.19 (t, *J* = 6.9 Hz, 3H), 1.16 (t, *J* = 6.8 Hz, 3H); $^{13}$C NMR (100 MHz, $CDCl_3$) δ 172.8, 171.9, 171.0, 82.8, 69.3, 61.6, 60.6, 51.6, 35.0, 30.2, 27.3, 14.7, 14.1, 14.0; HRMS (EI) for $C_{14}H_{21}O_5N_1$ [$M^+$]: calcd 283.1420, found 283.1423; HPLC: CHIRALPAK AD-H, *n*-hexane/*i*-PrOH = 90/10, 1.0 mL/min, $t_{R,1}$ = 18.280 min (**2*R***, major peak), $t_{R,2}$ = 21.250 min; 98% ee.

**2*S***

Yellowish liquid; analytical TLC, EtOAc/*n*-hexane = 1/1 (v/v), $R_f$ = 0.4; $^1$H NMR (400 MHz, $CDCl_3$) δ 6.56 (d, *J* = 7.9 Hz, 1H), 4.56 (td, *J* = 7.9, 5.0 Hz, 1H), 4.19 – 4.10 (m, 2H), 4.07 (q, *J* = 7.1 Hz, 2H), 2.51 – 2.43 (m, 2H), 2.43 – 2.25 (m, 4H), 2.21 – 2.08 (m, 1H), 1.99 – 1.87 (m, 2H), 1.28 – 1.13 (m, 6H); $^{13}$C NMR (100 MHz, $CDCl_3$) δ 172.9, 171.9, 171.0, 82.8, 69.4, 61.7, 60.7, 51.7, 35.1, 30.3, 27.4, 14.7, 14.2, 14.1; HRMS (EI) for $C_{14}H_{21}O_5N_1$ [$M^+$]: calcd 283.1420, found 283.1413; HPLC: CHIRALPAK AD-H, *n*-hexane/*i*-PrOH = 90/10, 1.0 mL/min, $t_{R,1}$ = 18.877 min, $t_{R,2}$ = 20.775 min (**2*S***, major peak); 98% ee; $[\alpha]_D^{22}$ +7.1 (*c* 5.6, $CHCl_3$).

**2rac**

Yellowish liquid; analytical TLC, EtOAc/*n*-hexane = 1/1 (v/v), $R_f$ = 0.4; $^1$H NMR (400 MHz, $CDCl_3$) δ 6.73 (d, *J* = 7.9 Hz, 1H), 4.49 (td, *J* = 8.1, 5.1 Hz, 1H), 4.07 (q, *J* = 7.1 Hz, 2H), 4.00 (q, *J* = 7.1 Hz, 2H), 2.52 – 2.17 (m, 6H) , 2.15 – 1.99 (m, 1H), 1.97 – 1.77 (m, 2H), 1.20 – 1.09 (m, 6H); $^{13}$C NMR (100 MHz, $CDCl_3$) δ 172.7, 171.8, 171.0, 82.7, 69.3, 61.4, 60.5, 51.6, 34.9, 30.2, 27.2, 14.6, 14.01, 13.96; HRMS (EI) for $C_{14}H_{21}O_5N_1$ [$M^+$]: calcd 283.1420, found 283.1409.

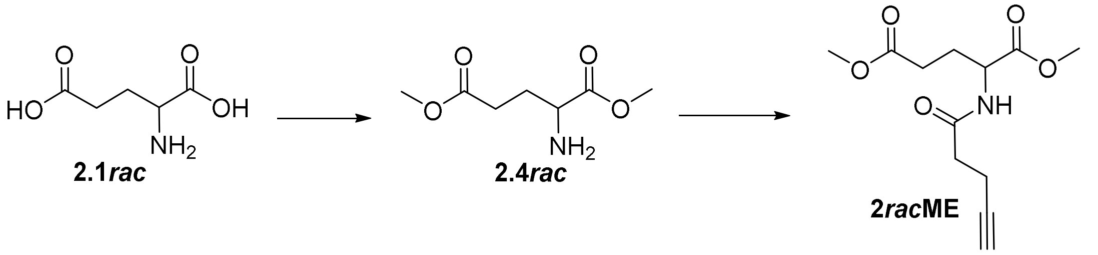

To a stirred solution of **2.1*rac*** (4.000 g, 27.2 mmol) in MeOH (39 mL) at 0 °C was added $SOCl_2$ (4.68 mL, 64.2 mmol) dropwise under Ar atmosphere. The mixture was stirred at rt for 30 min, then refluxed for 30 min. Residues were concentrated under vacuo and azeotroped with benzene to give product **2.4*rac*** as a white solid (4.700 g, 99% yield). **2.4*R*** and **2.4*S*** were prepared similarly with a yield of 99% and 99%, respectively.

To a solution of **2.4*rac*** (423.3 mg, 2.4 mmol) and 4-pentynoic acid (98.1 mg, 1 mmol) in anhydrous DMF (3.6 mL) were subsequently added EDCI (421.7 mg, 2.6 mmol), HOBt (297.3 mg, 2.6 mmol), and DIPEA (670 μL, 4.8 mmol) under an Ar atmosphere. The mixture was stirred at rt under an Ar atmosphere overnight. The resulting mixture was diluted with EtOAc, washed with 0.3 M HCl, saturated $NaHCO_3$ aqueous solution, water, and brine. The combined organic layer was dried over anhydrous $NaSO_4$ and concentrated under vacuo. Product was purified with column chromatography using 1/1 (v/v) EtOAc/*n*-hexane to give **2*rac*ME** as a colourless oil (168.5 mg, 66% yield). **2*R*ME** and **2*S*ME** were similarly prepared, with yields of 64% and 61%, respectively.

**2*R*ME**

Colorless sticky liquid; analytical TLC, $R_f$ = 0.3 (1/1 v/v EtOAc/*n*-hexane); $^1$H NMR (400 MHz, $CDCl_3$) δ 6.52 (d, *J* = 7.8 Hz, 1H), 4.61 (td, *J* = 7.9, 5.1 Hz, 1H), 3.71 (s, 3H), 3.63 (s, 3H), 2.57 – 2.28 (m, 6H), 2.24 – 2.12 (m, 1H), 2.03 – 1.90 (m, 2H); $^{13}$C NMR (100 MHz, $CDCl_3$) δ 173.4, 172.4, 171.0, 82.8, 69.5, 52.6,

51.9, 51.7, 35.1, 30.1, 27.4, 14.8; HRMS (EI) for $C_{12}H_{17}O_5N_1$ $[M]^+$: calcd 255.1107, found 255.1108; HPLC: CHIRALPAK AD-H, *n*-hexane/*i*-PrOH = 83/17, 1.0 mL /min, $t_{R,1}$ = 14.934 min (**2*R*ME**, major peak), $t_{R,2}$ = 17.115 min. 97% ee.

**2*S*ME**

Colorless sticky liquid; analytical TLC, $R_f$ = 0.3 (1/1 v/v EtOAc/*n*-hexane); $^1$H NMR (400 MHz, $CDCl_3$) δ 6.47 (d, *J* = 7.7 Hz, 1H), 4.62 (td, *J* = 7.9, 5.1 Hz, 1H), 3.73 (s, 3H), 3.65 (s, 3H), 2.60 – 2.27 (m, 6H), 2.27 – 2.10 (m, 1H), 2.05 – 1.86 (m, 2H); $^{13}$C NMR (100 MHz, $CDCl_3$) δ 173.4, 172.4, 171.0, 82.8, 69.5, 52.7, 52.0, 51.7, 35.2, 30.1, 27.4, 14.8; HRMS (EI) for $C_{12}H_{17}O_5N_1$ $[M]^+$: calcd 255.1107, found 255.1105; HPLC: CHIRALPAK AD-H, *n*-hexane/*i*-PrOH = 83/17, 1.0 mL/min, $t_{R,1}$ = 14.351 min, $t_{R,2}$ = 17.037 min (**2*S*ME**, major peak); 97% ee.

**2*rac*ME**

Analytical TLC, $R_f$ = 0.3 (1/1 v/v EtOAc/*n*-hexane); $^1$H NMR (400 MHz, $CDCl_3$) δ 6.54 (d, *J* = 7.9 Hz, 1H), 4.60 (td, *J* = 7.9, 5.1 Hz, 1H), 3.70 (s, 3H), 3.63 (s, 3H), 2.58 – 2.28 (m, 6H), 2.25 – 2.11 (m, 1H), 2.03 – 1.88 (m, 2H); $^{13}$C NMR (100 MHz, $CDCl_3$) δ 173.3, 172.4, 171.0, 82.8, 69.5, 52.6, 51.9, 51.6, 35.1, 30.0, 27.3, 14.8; HRMS (EI) for $C_{12}H_{17}O_5N_1$ $[M]^+$: calcd 255.1107, found 255.1107.

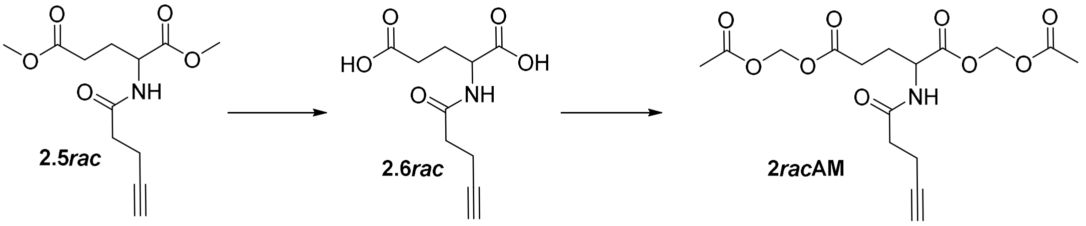

To a solution of **2.5*rac*** (335.4 mg, 1.3 mmol) in dioxane/$H_2O$ (2.9/2.9 mL) was added LiOH·$H_2O$ (161.2 mg, 3.1 mmol). The mixture was stirred at rt for 1 h, then quenched with 3 M HCl to pH = 2–3. The mixture was then extracted with EtOAc. The combined organic layer was washed with brine, dried over anhydrous $NaSO_4$ and concentrated under vacuo to afford crude **2.6*rac*** as a white sticky solid (165.4 mg, 56% yield). **2.6*R*** and **2.6*S*** were prepared similarly, with yields of 80% and 70%, respectively.

Crude diacid **2.6*rac*** (145.1 mg, 0.64 mmol) was dissolved in anhydrous MeCN (1 mL) under an Ar atmosphere. To the stirred solution were then added DIPEA (317.9 µL, 1.6 mmol) and bromomethyl acetate (157.5 µL, 1.4 mmol). The mixture was stirred under an Ar atmosphere at rt for 2 h, then diluted in EtOAc and washed with $H_2O$. The organic layer was concentrated under vacuo and purified with column chromatography using 1/1 (v/v) EtOAc/*n*-hexane to afford **2*rac*AM** as a transparent sticky liquid (175.8 mg, 74% yield). **2*R*AM** and **2*S*AM** were similarly prepared, with yields of 54% and 43%, respectively.

**2*R*AM**

Colorless sticky liquid; analytical TLC, $R_f$ = 0.5 (1/1 v/v EtOAc/*n*-hexane); $^1$H NMR (400 MHz, $CDCl_3$) δ 6.46 (d, *J* = 7.8 Hz, 1H), 5.83 – 5.73 (m, 2H), 5.72 (s, 2H), 4.68 (td, *J* = 8.0, 5.0 Hz, 1H), 2.60 – 2.38 (m, 6H), 2.27 – 2.12 (m, 1H), 2.12 (s, 6H), 2.07 – 1.94 (m, 2H); $^{13}$C NMR (100 MHz, $CDCl_3$) δ 171.5, 171.1, 170.6, 169.7, 169.5, 82.7, 79.7, 79.3, 69.6, 51.4, 35.0, 29.8, 26.6, 20.7, 20.6, 14.7; HRMS (ESI) for $C_{16}H_{21}O_9N_1$ $[M]^+$: calcd 371.1216, found 371.1217; HPLC: CHIRALPAK AD-H, *n*-hexane/*i*-PrOH = 70/30, 1.0 mL/min, $t_{R,1}$ = 12.190 min (**2*R*AM**, major peak), $t_{R,2}$ = 14.504 min; 97% ee.

**2*S*AM**

Colorless sticky liquid; analytical TLC, $R_f$ = 0.5 (1/1 v/v EtOAc/*n*-hexane); $^1$H NMR (400 MHz, $CDCl_3$) δ 6.40 (d, *J* = 7.8 Hz, 1H), 5.81 – 5.71 (m, 2H), 5.71 (s, 2H), 4.67 (td, *J* = 8.0, 5.1 Hz, 1H), 2.58 – 2.36 (m, 6H), 2.27 – 2.14 (m, 1H), 2.10 (s, 6H), 2.07 – 1.92 (m, 2H); $^{13}$C NMR (100 MHz, $CDCl_3$) δ 171.6, 171.1, 170.7, 169.8, 169.5, 82.8, 79.8, 79.4, 69.7, 51.5, 35.1, 29.9, 26.7, 20.8, 20.7, 14.8; HRMS (ESI) for $C_{16}H_{21}O_9N_1$ $[M]^+$: calcd 371.1216, found 371.1213; HPLC: CHIRALPAK AD-H, *n*-hexane/*i*-PrOH = 70/30, 1.0 mL/min, $t_{R,1}$ = 12.188 min, $t_{R,2}$ = 14.483 min (**2*S*AM**, major peak); 97% ee.

**2*rac*AM**

Analytical TLC, $R_f$ = 0.5 (1/1 v/v EtOAc/*n*-hexane); $^1$H NMR (400 MHz, $CDCl_3$) δ 6.47 (d, *J* = 7.8 Hz, 1H), 5.79 – 5.70 (m, 2H), 5.69 (s, 2H), 4.65 (td, *J* = 8.1, 5.1 Hz, 1H), 2.57 – 2.35 (m, 6H), 2.23 – 2.13 (m, 1H), 2.09 (s, 6H), 2.04 – 1.91 (m, 2H); $^{13}$C NMR (100 MHz, $CDCl_3$) δ 171.6, 171.2, 170.6, 169.8, 169.5, 82.8, 79.8, 79.4, 69.7, 51.4, 35.1, 29.9, 26.7, 20.8, 20.7, 14.8; HRMS (ESI) for $C_{16}H_{21}O_9N_1$ $[M]^+$: calcd 371.1216, found 371.1214.

**Probes series 3**

Diethyl *a*-ketoglutarate (αKG) **3.2** was prepared from 2-ketoglutaric acid **3.1** under acidic conditions and was propargylated in the presence of aluminium and mercury chloride to afford the racemic **3*rac***. To separate enantiomers, probe **3*rac*** was hydrolyzed to a lactone acid and derivatized with a chiral auxiliary *R*-4-methoxyphenylethylamine to facilitate the separation via diastereomers of **3.3**. The absolute configuration of the quaternary carbon in **3.4-2**, which was the second component eluted during repetitive column chromatography purification of **3.4**, was found to be of *S* form, suggesting that probe **3*S*** was synthesized after oxidative deprotection and acidic esterification. The configuration of the probe generated from **3.4-2** was then assigned as *R* accordingly.

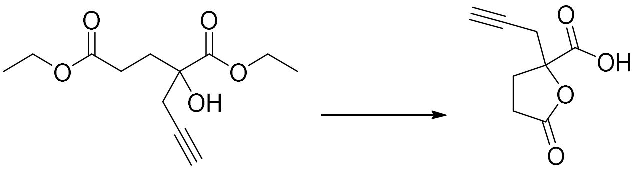

**3*rac*** **3.3**

To a solution of compound **3*rac*** (1.000 g, 4.1 mmol) in THF (17.5 mL) was added a solution of LiOH (384.8 mg, 9 mmol) in water (1.75 mL). The reaction was stirred overnight, and then 6 M HCl (15 mL) was added. The reaction mixture was stirred overnight, evaporated, azeotroped with acetonitrile, and purified with column chromatography using 1/1 (v/v) DCM and acetonitrile to afford **3.6** as a yellowish solid (627 mg, 91% yield).

**3.3**

Yellowish solid, m.p. 70–75 °C; analytical TLC, MeOH/DCM = 1/9 (v/v), 0.5% AcOH, $R_f$ = 0.4; $^1$H NMR (400 MHz, $CDCl_3$) δ 6.74 (s, 1H), 2.99 – 2.78 (m, 2H), 2.67 – 2.59 (m, 2H), 2.53 – 2.42 (m, 1H), 2.42 – 2.32 (m, 2H); $^{13}$C NMR (100 MHz, $CDCl_3$) δ 177.1, 172.2, 85.0, 78.8, 72.8, 30.7, 28.6, 27.6; HRMS (EI) for $C_8H_8O_4$ $[M]^+$: calcd 168.0423, found 168.0429.

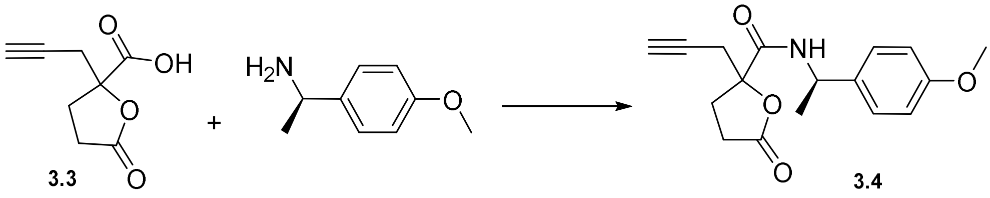

**3.3** **3.4**

To a stirred solution of **3.3** (168.15 mg, 1 mmol) and *R*-4-methoxyphenylethylamine (118 μL, 0.8 mmol) in DMF (1.06 mL) were added *N*-methylmorpholine (1.06 mL, 11.9 mmol), HOBt (191.6 mg, 1.53 mmol), and EDCI (216.1 mg, 1.35 mmol) sequentially. The reaction was stirred under an Ar atmosphere at room temperature overnight. Then the reaction mixture was diluted with ethyl acetate, washed with saturated

$NaHCO_3$ solution, water, and brine. The combined organic layer was dried with anhydrous $NaSO_4$ and concentrated under vacuo. The two diastereomers were separated with column chromatography using EtOAc/*n*-hexane (1/1 v/v) to afford **3.4-1** (99 mg, 33% yield) and **3.4-2** (99 mg, 33% yield) as white solids (1:1 dr).

**3.4-1**

M.p. 130–138 °C; analytical TLC, EtOAc/*n*-hexane = 1/1 (v/v), $R_f$ = 0.3; $^1$H NMR (400 MHz, $CDCl_3$) δ 7.25 – 7.16 (m, 2H), 6.91 – 6.83 (m, 2H), 6.80 (d, *J* = 8.2 Hz, 1H), 5.05 (p, *J* = 6.9 Hz, 1H), 3.78 (s, 3H), 2.86 (d, *J* = 2.6 Hz, 2H), 2.78 – 2.62 (m, 1H), 2.55 – 2.34 (m, 3H), 2.11 (t, *J* = 2.6 Hz, 1H), 1.49 (d, *J* = 6.9 Hz, 3H); $^{13}$C NMR (100 MHz, $CDCl_3$) δ 175.4, 169.6, 159.0, 134.6, 127.3, 114.2, 85.5, 77.6, 72.3, 55.3, 48.6, 29.8, 28.7, 28.4, 21.7; HRMS (EI) for $C_{17}H_{19}O_4N$ $[M]^+$: calcd 301.1314, found 301.1309.

**3.4-2**

M.p. 132–143 °C; analytical TLC, EtOAc/*n*-hexane = 1/1 (v/v), $R_f$ = 0.2; $^1$H NMR (400 MHz, $CDCl_3$) δ 7.27 – 7.19 (m, 2H), 6.92 – 6.80 (m, 3H), 5.05 (p, *J* = 7.1 Hz, 1H), 3.78 (s, 3H), 2.85 – 2.69 (m, 3H), 2.65 – 2.51 (m, 2H), 2.50 – 2.39 (m, 1H), 2.06 – 1.99 (m, 1H), 1.48 (d, *J* = 7.0 Hz, 3H); $^{13}$C NMR (100 MHz, CD $Cl_3$) δ 175.5, 169.6, 158.9, 134.5, 127.4, 114.0, 85.5, 77.4, 72.3, 55.2, 48.4, 29.7, 28.6, 28.4, 21.5; HRMS (EI) for $C_{17}H_{19}O_4N$ $[M]^+$: calcd 301.1314, found 301.1311.

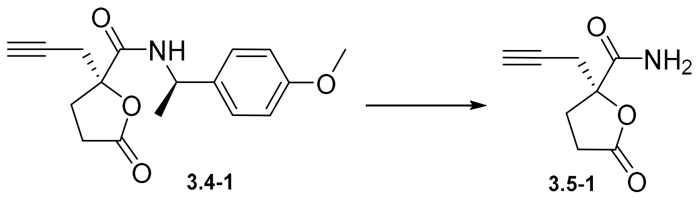

To a solution of **3.4-1** (30.1 mg, 0.1 mmol) in MeCN (3.49 mL) and $H_2O$ (0.93 mL) was added ceric ammonium nitrate (272.6 mg, 0.5 mmol). The reaction mixture was stirred at rt for 40 min, then diluted with $H_2O$ and extracted with EtOAc. The combined organic layer was washed with saturated $NaHCO_3$ aqueous solution and water, dried with anhydrous $Na_2SO_4$, and concentrated under vacuo. The reaction mixture was then purified with column chromatography using 1/1 (v/v) EtOAc/*n*-hexane to afford **3.5-1** as a yellowish liquid (6.7 mg, 40% yield). **3.5-2** was prepared from **3.4-2** similarly, with a yield of 39%.

**3.5-1**

Analytical TLC, EtOAc/*n*-hexane = 1/1 (v/v), $R_f$ = 0.1; $^1$H NMR (300 MHz, $CDCl_3$) δ 6.61 (s, 1H), 6.05 (s, 1H), 2.85 (d, *J* = 2.7 Hz, 2H), 2.82 – 2.67 (m, 1H), 2.68 – 2.30 (m, 3H), 2.11 (t, *J* = 2.7 Hz, 1H); $^{13}$C NMR (100 MHz, $CDCl_3$) δ 175.5, 173.8, 85.4, 77.3, 72.6, 29.7, 28.58, 28.55; HRMS (EI) for $C_8H_9O_3N$ $[M]^+$: calcd 167.0582, found 167.0578.

**3.5-2**

Yellowish liquid; analytical TLC, EtOAc/*n*-hexane = 1/1 (v/v), $R_f$ = 0.1; $^1$H NMR (300 MHz, $CDCl_3$) δ 6.59 (s, 1H), 6.04 (s, 1H), 2.85 (dd, *J* = 2.7, 0.7 Hz, 2H), 2.80 – 2.70 (m, 1H), 2.69 – 2.28 (m, 3H), 2.11 (t, *J* = 2.6 Hz, 1H); $^{13}$C NMR (100 MHz, $CDCl_3$) δ 175.5, 173.7, 85.4, 77.4, 72.5, 29.7, 28.57, 28.55; HRMS (EI) for $C_8H_9O_3N$ $[M]^+$: calcd 167.0582, found 167.0574.

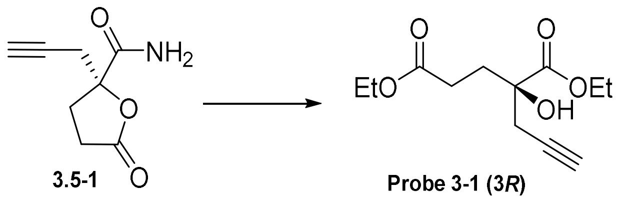

To a solution of compound **3.5-1** (14.7 mg, 0.088 mmol) in EtOH (0.5 mL) was added one drop of 98% $H_2SO_4$. The reaction mixture was stirred at 80 °C overnight and then poured into a cold saturated aqueous $NaHCO_3$ solution. The mixture was then extracted with EtOAc, dried with anhydrous $NaSO_4$, and concentrated under vacuo. The product **3** was purified with column chromatography using 1/2 (v/v) EtOAc/*n*-hexane to afford probe **3-1** as a yellowish liquid (6.4 mg, 30% yield, 99% ee). Probe **3-2** was similarly prepared from **3.5-2**, with a yield of 37% and 99% ee.

**3-1 (3*R*)**

Analytical TLC, EtOAc/*n*-hexane = 1/1 (v/v), $R_f$ = 0.6; $^1$H NMR (400 MHz, $CDCl_3$) δ 4.34 – 4.17 (m, 2H), 4.16 – 3.93 (m, 2H), 3.55 (s, 1H), 2.70 – 2.52 (m, 2H), 2.48 (dad, *J* = 15.8, 9.2, 6.4 Hz, 1H), 2.28 – 2.13 (m, 1H), 2.09 (ddd, *J* = 9.7, 5.8, 4.7 Hz, 2H), 2.05 – 2.01 (t, *J* = 2.6 Hz, 1H), 1.30 (t, *J* = 7.1 Hz, 3H), 1.23 (t, *J* = 7.1 Hz, 3H); $^{13}$C NMR (100 MHz, $CDCl_3$) δ 174.4, 173.1, 78.7, 75.7, 71.6, 62.6, 60.7, 33.0, 30.2, 28.9, 14.29, 14.27; HRMS (EI) for $C_{12}H_{18}O_5$ $[M]^+$: calcd 242.1154, found 242.1159; HPLC: CHIRALPAK

AD-H, *n*-hexane/*i*-PrOH = 90/10, 1.0 mL/min, $t_{R,1}$ = 9.500 min (**3*R***, major peak), $t_{R,2}$ = 12.154 min; 98% ee.

**3-2 (3*S*)**

Yellowish liquid; analytical TLC, EtOAc/*n*-hexane = 1/1 (v/v), $R_f$ = 0.6; $^1$H NMR (400 MHz, $CDCl_3$) δ 4.34 – 4.14 (m, 2H), 4.08 (q, *J* = 7.1 Hz, 2H), 3.56 (s, 1H), 2.66 – 2.50 (m, 2H), 2.46 (ddd, *J* = 15.8, 9.2, 6.4 Hz, 1H), 2.18 (ddd, *J* = 15.9, 9.9, 5.8 Hz, 1H), 2.11 – 2.04 (m, 2H), 2.02 (t, *J* = 2.6 Hz, 1H), 1.28 (t, *J* = 7.1 Hz, 3H), 1.21 (t, *J* = 7.1 Hz, 3H); $^{13}$C NMR (100 MHz, $CDCl_3$) δ 174.3, 173.0, 78.6, 75.6, 71.6, 62.5, 60.7, 32.9, 30.2, 28.9, 14.24, 14.21; HRMS (EI) for $C_{12}H_{18}O_5$ $[M]^+$: calcd 242.1154, found 242.1158; HPLC: CHIRALPAK AD-H, *n*-hexane/*i*-PrOH = 90/10, 1.0 mL/min, $t_{R,1}$ = 9.583 min, $t_{R,2}$ = 12.125 min (**3*S***, major peak); 99% ee.

**Probes series 4**

Diethyl *α*-ketoglutarate **4.2** and pyrrolidine were refluxed in toluene, and freshly reacted with propargyl bromide to afford a C1-amidated αKG alkyne **4.3**. The reduction of **4.3** with $NaBH_4$ generated **4.4**, which was hydrolysed and derivatized with chiral auxiliary *R*-4-methoxyphenylethylamine to afford **4.5** as diastereomers. The first component after column chromatography, **4.5-1**, was crystallized and analysed by X-ray crystallography, with both chiral centers determined to be *S* from its relative configuration. Therefore, after oxidative deprotection, acidic esterification, and further purification, diethyl ester **4-1** should be **4*SS***. The second component, after **4.5-1** isolation, **4.5-2**, provided **4.6-2** and **4-2** that share the same NMR patterns as **4.6-1** and **4-1**, respectively, suggesting an absolute configuration of **4*RR*** for **4-2**, which was purified via derivatization with TESCl. The separation of **4.5-3/4** was much more difficult, and probes **4-3/4** were prepared as a mixture.

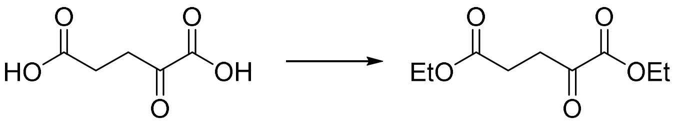

**4.1** **4.2**

To a stirred solution of **4.1** (14.610 g, 100 mmol) in EtOH (420 mL) was added AcCl (0.71 mL, 10 mmol). The reaction was stirred under an Ar atmosphere overnight at rt, then concentrated under vacuo to afford **4.2** as a colorless liquid (20.000 g, 99% yield).

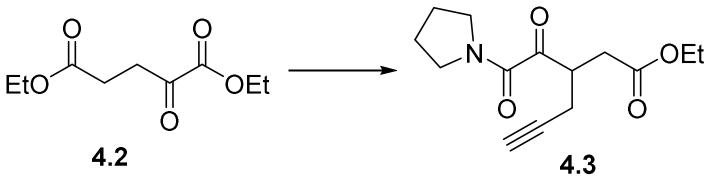

To a mixture of diethyl αKG **4.2** (4.900 g, 24.2 mmol) and pyrrolidine (3.92 mL, 48.4 mmol) in toluene (72 mL) was added *p*-TsOH·$H_2O$ (228 mg, 12.1 mmol). The mixture was refluxed with a Dean-Stark apparatus overnight under an Ar atmosphere, then concentrated under vacuo. The residue was dissolved in $Et_2O$, filtered over a celite pad, and then concentrated. The crude product was dissolved in anhydrous MeCN (120 mL), followed by the addition of propargyl bromide (5.35 mL, 48.4 mmol). The resulting mixture was stirred at 70℃ under an Ar atmosphere for 2 d. Then $H_2O$ (18 mL) was added, and the mixture was further refluxed for 2 h. After cooling to rt, the mixture was concentrated under vacuo, dissolved in EtOAc, washed with $H_2O$ brine, and dried over anhydrous $NaSO_4$. The combined organic layer was concentrated under vacuo and purified with column chromatography using 1/5-1/1 (v/v) EtOAc/*n*-hexane to afford **4.3** as a yellowish liquid (642 mg, 10% yield).

**4.3**

Analytical TLC, EtOAc/*n*-hexane = 1/1 (v/v), $R_f$ = 0.6; $^1$H NMR (400 MHz, $CDCl_3$) δ 4.10 (q, *J* = 7.1 Hz, 2H), 3.73 (tt, *J* = 7.8, 5.3 Hz, 1H), 3.64 (dd, *J* = 6.9, 5.8 Hz, 2H), 3.50 (dd, *J* = 7.3, 5.9 Hz, 2H), 2.98 – 2.73 (m, 2H), 2.73 – 2.53 (m, 2H), 2.02 (t, *J* = 2.7 Hz, 1H), 1.96 – 1.81 (m, 4H), 1.23 (t, *J* = 7.1 Hz, 3H); $^{13}$C NMR (100 MHz, $CDCl_3$) δ 199.5, 172.1, 162.3, 80.5, 71.2, 61.0, 47.4, 46.4, 42.6, 34.1, 26.4, 23.8, 19.9, 14.3; HRMS (EI) for $C_{14}H_{19}O_4N$ $[M]^+$: calcd 265.1314, found 265.1310.

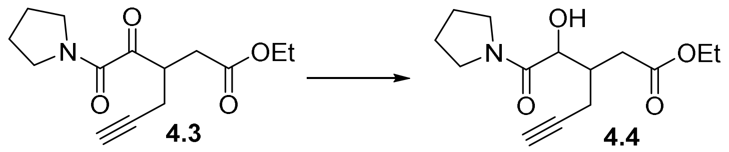

To a stirred solution of **4.3** (50.9 mg, 0.2 mmol) in EtOH (2 mL) was added $NaBH_4$ (7.6 mg, 1 mmol) at 0℃ under an Ar atmosphere. The mixture was stirred at 0℃ for 40 min, then quenched with saturated $NH_4Cl$ aqueous solution and extracted with EtOAc. The combined organic layer was dried over anhydrous $Na_2SO_4$, concentrated under vacuo, and purified with column chromatography using 1/1 (v/v) EtOAc/*n*-hexane to afford **4.4** as a colorless liquid (32 mg, 60% yield, dr = 1:1).

**4.4**

Analytical TLC, EtOAc/*n*-hexane = 1/1 (v/v), $R_f$ = 0.4, a mixture of 2 diastereomers and 2 enantiomers, dr = 1: 1; $^1$H NMR (500 MHz, $CDCl_3$) δ 4.55 (d, *J* = 6.4 Hz, 1H), 4.41 (d, *J* = 5.6 Hz, 1H), 4.14 (q, *J* = 7.2 Hz, 2H), 4.08 (q, *J* = 7.2 Hz, 2H), 3.67 (t, *J* = 7.2 Hz, 2H), 3.63 – 3.47 (m, 6H), 3.48 – 3.35 (m, 2H), 2.79 – 2.62 (m, 2H), 2.48 – 2.34 (m, 5H), 2.25 – 2.11 (m, 3H), 2.07 – 1.97 (m, 3H), 1.96 (t, *J* = 2.7 Hz, 1H), 1.95 – 1.83 (m, 6H), 1.27 (t, *J* = 7.2, 1.0 Hz, 3H), 1.23 (t, *J* = 7.1, 1.0 Hz, 3H); $^{13}$C NMR (125 MHz, $CDCl_3$) δ 173.2, 172.6, 171.32, 171.27, 82.6, 82.3, 70.2, 70.0, 69.5, 69.4, 60.71, 60.68, 46.6, 46.5, 46.4, 46.1, 37.1, 36.9, 34.5, 32.6, 26.3, 26.2, 23.91, 23.90, 21.6, 17.3, 14.34, 14.29; HRMS (EI) for $C_{14}H_{21}O_4N$ $[M]^+$: calcd 267.1471, found 267.1468.

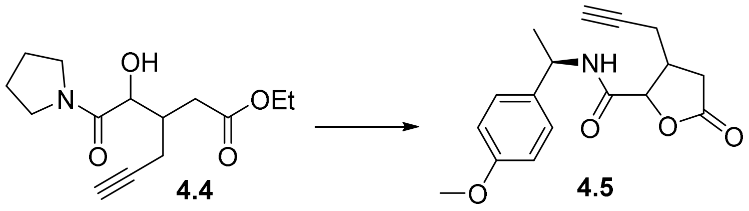

To a stirred solution of **4.4** (43.4 mg, 0.16 mmol) in MeCN was added 1 M HCl (16.3 mL). The reaction was refluxed overnight, then quenched with saturated $NaHCO_3$ solution, extracted with EtOAc, and concentrated under vacuo to afford a crude lactone acid (70%).

The resulting residue (103 mg, 0.6 mmol) was dissolved in DMF/DCM (0.8 mL/3.1 mL), followed by the addition of *R*-4-methoxyphenylethylamine (89 μL, 0.5 mmol), *N*-methylmorpholine (0.8 mL, 7 mmol), HOBt (144 mg, 0.9 mmol), and EDCI (162 mg, 0.8 mmol) sequentially. The reaction mixture was stirred at rt under an Ar atmosphere overnight, then diluted in EtOAc, washed with saturated $NaHCO_3$ solution, brine, dried over anhydrous $Na_2SO_4$, concentrated under vacuo, and separated via column chromatography using gradient 1/10–1/2 (v/v) EtOAc/*n*-hexane to afford **4.5** as a mixture of white solid (101 mg, 56% yield, including **4.5-1**: 28%, **4.5-2**: 35%, and **4.5-3/4**: 40%).

**4.5-1**

White solid, m.p. 140–155 °C; analytical TLC, EtOAc/*n*-hexane = 1/3 (v/v), $R_f$ = 0.3; $^1$H NMR (500 MHz, $CDCl_3$) δ 7.24 – 7.15 (m, 2H), 6.92 – 6.82 (m, 2H), 6.55 (d, *J* = 8.1 Hz, 1H), 5.07 (p, *J* = 7.1 Hz, 1H), 4.67 (d, *J* = 7.1 Hz, 1H), 3.79 (s, 3H), 2.79 – 2.68 (m, 2H), 2.68 – 2.55 (m, 3H), 2.10 (t, *J* = 2.5 Hz, 1H), 1.50 (d, *J* = 6.9 Hz, 3H); $^{13}$C NMR (125 MHz, $CDCl_3$) δ 174.4, 167.9, 159.2, 134.6, 127.4, 114.3, 79.9, 79.3,

72.5, 55.5, 48.4, 38.1, 32.9, 21.9, 21.8; HRMS (EI) for $C_{17}H_{19}O_4N$ $[M]^+$: calcd 301.1314, found 301.1315.

**4.5-2**

White solid, m.p. 138–153 °C; analytical TLC, EtOAc/*n*-hexane = 1/3 (v/v), $R_f$ = 0.3; $^1$H NMR (500 MHz, $CDCl_3$) δ 7.23 (d, *J* = 8.7 Hz, 2H), 6.87 (d, *J* = 8.7 Hz, 2H), 6.54 (d, *J* = 8.0 Hz, 1H), 5.06 (p, *J* = 7.1 Hz, 1H), 4.63 (d, *J* = 7.6 Hz, 1H), 3.80 (s, 3H), 2.90 – 2.79 (m, 1H), 2.79 – 2.59 (m, 4H), 2.09 (t, *J* = 2.6 Hz, 1H), 1.49 (d, *J* = 6.9 Hz, 3H); δ 174.5, 168.0, 159.2, 134.4, 127.6, 114.3, 79.8, 79.3, 72.5, 55.5, 48.4, 38.2, 32.9, 22.0, 21.7.; HRMS (EI) for $C_{17}H_{19}O_4N$ $[M]^+$: calcd 301.1314, found 301.1317.

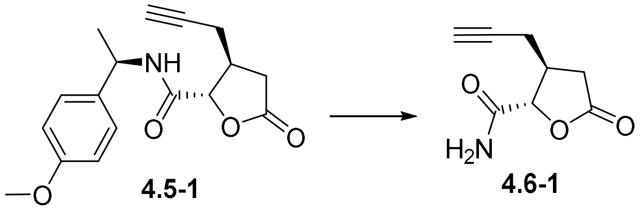

**4.5-1** **4.6-1**

To a stirred solution of **4.5-1** (108.9 mg, 0.36 mmol) in MeCN was added ceric ammonium nitrate (1.020 g, 1.8 mmol). The reaction was stirred under rt for 40 min, then diluted in water and extracted with EtOAc. The combined organic layer was washed with saturated $NaHCO_3$ aqueous solution and brine, then concentrated under vacuo and purified with column chromatography using 2/1 EtOAc/*n*-hexane (v/v) to afford **4.6-1** as a yellowish liquid (16.8 mg, 28% yield). **4.6-2** and **4.6-3/4** were prepared similarly, with yields of 35% and 40%, respectively.

**4.6-1**

Yellowish liquid; analytical TLC, EtOAc/*n*-hexane = 2/1 (v/v), $R_f$ = 0.2; $^1$H NMR (500 MHz, $CDCl_3$) δ 6.45 (s, 1H), 5.97 (s, 1H), 4.67 (d, *J* = 7.3 Hz, 1H), 2.91 – 2.78 (m, 1H), 2.79 – 2.54 (m, 4H), 2.12 (t, *J* = 2.6 Hz, 1H); $^{13}$C NMR (125 MHz, $CDCl_3$) δ 174.5, 171.7, 79.7, 79.2, 72.6, 38.1, 32.8, 22.0; HRMS (EI) for $C_8H_9O_3N$ $[M]^+$: calcd 167.0582, found 167.0580.

**4.6-2**

Yellowish liquid; analytical TLC, EtOAc/*n*-hexane = 1/1 (v/v), $R_f$ = 0.1; $^1$H NMR (500 MHz, $CDCl_3$) δ 6.42 (s, 1H), 5.88 (s, 1H), 4.67 (d, *J* = 7.3 Hz, 1H), 2.90 – 2.79 (m, 1H), 2.79 – 2.54 (m, 4H), 2.12 (t, *J* = 2.6 Hz, 1H); $^{13}$C NMR (125 MHz, $CDCl_3$) δ 174.4, 171.6, 79.7, 79.2, 72.6, 38.1, 32.8, 22.0; HRMS (EI) for

$C_8H_9O_3N$ [M]$^+$: calcd 167.0582, found 167.0583.

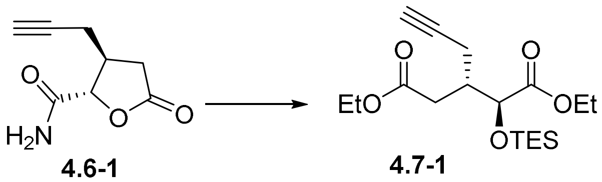

**4.6-1** **4.7-1**

To a stirred solution of **4.6-1** (14 mg, 0.08 mmol) in EtOH (1 mL) was added concentrated $H_2SO_4$ (1 drop). The reaction was refluxed overnight, then diluted in water and extracted with EtOAc. The combined organic layer was washed with saturated $NaHCO_3$ aqueous solution and brine, dried over anhydrous $Na_2SO_4$, concentrated under vacuo, and purified with column chromatography using 1/6 EtOAc/*n*-hexane (v/v) to afford a mixture of **4-1** and **4-1*cy*** as an oil.

The product-containing oil was then dissolved in DCM (1 mL), followed by the addition of imidazole (8 mg, 0.1 mmol) and TESCl (11.7 μL, 0.09 mmol) under an Ar atmosphere. After 15 min, the reaction mixture was diluted in EtOAc, washed with water and brine, concentrated under vacuo, then purified with column chromatography using 1/20 EtOAc/*n*-hexane (v/v) to afford **4.7-1** as a colorless liquid (7 mg, 39% yield).

**4.7-1**

Analytical TLC, EtOAc/*n*-hexane = 1/3 (v/v), $R_f$ = 0.9; $^1$H NMR (500 MHz, $CDCl_3$) δ 4.49 (d, *J* = 3.4 Hz, 1H), 4.18 (q, *J* = 6.9 Hz, 2H), 4.12 (q, *J* = 7.2 Hz, 2H), 2.54 (tddd, *J* = 8.1, 6.2, 5.1, 3.4 Hz, 1H), 2.51 – 2.27 (m, 4H), 2.02 (t, *J* = 2.7 Hz, 1H), 1.29 (t, *J* = 7.2 Hz, 3H), 1.25 (t, *J* = 7.1 Hz, 3H), 0.96 (t, *J* = 7.9 Hz, 9H), 0.63 (qd, *J* = 7.9, 2.3 Hz, 6H); $^{13}$C NMR (125 MHz, $CDCl_3$) δ 172.9, 172.5, 82.0, 72.4, 70.5, 61.1, 60.6, 39.3, 33.7, 20.4, 14.3, 6.9, 4.8; HRMS (EI) for $C_{18}H_{32}O_5Si$ [M]$^+$: calcd 365.2019, found 365.2022.

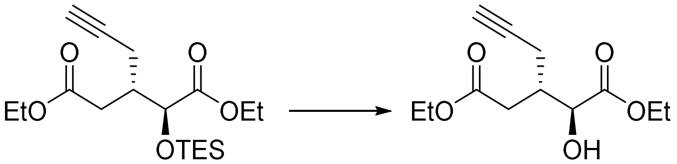

**4.7-1** **Probe 4-1 = 4*SS***

To a stirred solution of **4.7-1** (23.5 mg, 0.1 mmol) in EtOH (1 mL) was added PPTS (1.8 mg, 0.01 mmol). The reaction was stirred under rt for 2.5 h, then diluted in water and extracted with DCM. After washing with water and brine, the combined organic layer was concentrated and purified with column chromatography to afford **4-1** as a colorless liquid (9 mg, 37% yield). **4.7-2** was generated similarly to **4.7-1**, with a yield of 39%. Probe **4-3/4** was prepared similarly, with a yield of 28% from **4.5-3/4**.

**4-1 (4*SS*)**

Colorless liquid; analytical TLC, EtOAc/*n*-hexane = 1/3 (v/v), $R_f$ = 0.6; $^1$H NMR (500 MHz, $CDCl_3$) δ 4.42 (dd, *J* = 5.4, 3.3 Hz, 1H), 4.27 (qd, *J* = 7.1, 2.1 Hz, 2H), 4.12 (q, *J* = 7.2 Hz, 2H), 2.96 (d, *J* = 5.3 Hz, 1H), 2.59 (td, *J* = 7.0, 3.3 Hz, 1H), 2.47 – 2.33 (m, 4H), 2.04 (s, 1H), 1.32 (t, *J* = 7.2 Hz, 3H), 1.25 (t, *J* = 7.1 Hz, 3H); $^{13}$C NMR (125 MHz, $CDCl_3$) δ 174.4, 172.4, 81.7, 71.3, 70.6, 62.3, 60.8, 38.4, 33.7, 20.8, 14.3; HRMS (EI) for $C_{12}H_{18}O_5$ $[M]^+$: calcd 242.1154, found 242.1157.

**4-2 (4*RR*)**

Colorless liquid; analytical TLC, EtOAc/*n*-hexane = 1/3 (v/v), $R_f$ = 0.6; $^1$H NMR (500 MHz, $CDCl_3$) δ 4.42 (dd, *J* = 5.4, 3.3 Hz, 1H), 4.27 (qd, *J* = 7.2, 2.1 Hz, 2H), 4.12 (q, *J* = 7.1 Hz, 2H), 2.96 (d, *J* = 5.4 Hz, 1H), 2.58 (dt, *J* = 7.0, 3.5 Hz, 1H), 2.49 – 2.37 (m, 4H), 2.04 (t, *J* = 2.7 Hz, 1H), 1.32 (t, *J* = 7.2 Hz, 3H), 1.25 (t, *J* = 7.1 Hz, 3H); $^{13}$C NMR (125 MHz, $CDCl_3$) δ 174.4, 172.3, 81.7, 71.3, 70.6, 62.3, 60.8, 38.4, 33.6, 20.8, 14.3; HRMS (EI) for $C_{12}H_{18}O_5$ $[M]^+$: calcd 242.1154, found 242.1157.

**4-3/4 (4*RS*/*SR*)**

Colorless liquid; analytical TLC, EtOAc/*n*-hexane = 1/3 (v/v), $R_f$ = 0.6; $^1$H NMR (500 MHz, $CDCl_3$) δ 4.33 (dd, *J* = 5.3, 2.9 Hz, 1H), 4.30 – 4.24 (m, 2H), 4.16 (qd, *J* = 7.1, 2.0 Hz, 2H), 2.96 (d, *J* = 5.2 Hz, 1H), 2.65 – 2.55 (m, 3H), 2.27 (qdd, *J* = 17.0, 6.6, 2.7 Hz, 2H), 1.99 (t, *J* = 2.7 Hz, 1H), 1.32 (t, *J* = 7.1 Hz, 3H), 1.27 (t, *J* = 7.2 Hz, 3H); $^{13}$C NMR (125 MHz, $CDCl_3$) δ 174.2, 172.4, 81.7, 71.4, 70.5, 62.2, 60.8, 38.0, 35.1, 18.5, 14.3, 14.3; HRMS (EI) for $C_{12}H_{18}O_5$ $[M]^+$: calcd 242.1154, found 242.1154.

**Probes series 5**

To synthesize an enantiomerically and diastereomerically pure probe with an *R*-hydroxyl group, we applied a chiral epoxide **5.1*S*** as the starting material. After condensation of protected epoxide **5.2*S*** with diethyl malonate, a lactone ester **5.3*S*** was formed, which was propargylated under basic conditions. The resulting compound **5.4*S*** was hydrolysed, decarboxylated, and deprotected to afford lactone alcohol **5.6*S***. The alcohol was then oxidized and ethylated to afford diastereomers of lactone esters **5.7*S***, which were then esterified under acidic conditions to produce two diastereomers of probe **5**, **5*S*-1** and **5*S*-2**. Probe **5*R*-1** and **5*R*-2** were prepared from **5.1*R*** similarly. To determine the absolute configuration of the propargylated carbon, the lactone acid oxidized from **5.6*R*** was coupled to *R*-methoxyphenylethylamine to facilitate crystallization and X-ray crystallography analysis. The relative configuration of the second component after column separation, **5.9*R*-2**, was determined, indicating an *S* form at the chiral center bearing a propargyl group. **5.9*R*-2** was oxidatively deprotected and converted to diethyl ester under acidic conditions. $^{1}$H NMR showed identical spectra for **5.8*R*-2** and **5*R*-2** generated from **5.7*R*-2**, thus the configurations of the two probes synthesized from *R*-glycidol were determined as **5*SR*** and **5*SS***, respectively. The absolute configurations of probe enantiomers **5*S*-1** and **5*S*-2** with *R*-form hydroxyl group were assigned as **5*RS*** and **5*RR***, respectively.

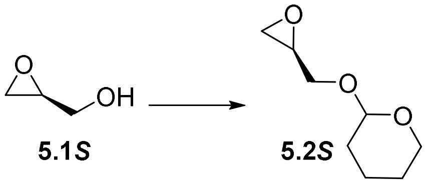

**5.1*S*** **5.2*S***

To a stirred solution of *S*-glycidol (6.64 mL, 100 mmol) in dry DCM (330 mL) were added 3,4-dihydropyran (23 mL, 500 mmol) and *p*-toluenesulfonic acid (190 mg, 1 mmol). The mixture was stirred for 3 h, and subsequently quenched with saturated aqueous $NaHCO_3$ (420 mL). After continued stirring for 10 min, the reaction mixture was extracted with DCM, dried with anhydrous $Na_2SO_4$, concentrated under vacuo, then purified with flash chromatography using 1/10–1/5 (v/v) EtOAc/*n*-hexane to give **5.2*S*** as a colourless liquid (15.800 g, 99% yield). **5.2*R*** was similarly prepared from *R*-glycidol, with a yield of 74%.

**5.2*S***

Analytical TLC, EtOAc/*n*-hexane = 1/3 (v/v), $R_f$ = 0.7, a mixture of 2 diastereomers; $^{1}$H NMR (500 MHz, $CDCl_3$) δ 4.67 (t, *J* = 3.7 Hz, 1H), 4.65 (t, *J* = 3.6 Hz, 1H), 3.95 (dd, *J* = 11.7, 3.1 Hz, 1H), 3.87 (dddd, *J* = 11.3, 8.6, 5.2, 3.1 Hz, 2H), 3.71 (qd, *J* = 11.8, 4.2 Hz, 2H), 3.55 – 3.48 (m, 2H), 3.40 (dd, *J* = 11.7, 6.4 Hz, 1H), 3.23 – 3.15 (m, 2H), 2.81 (td, *J* = 4.5, 2.6 Hz, 2H), 2.69 (dd, *J* = 5.2, 2.7 Hz, 1H), 2.60 (dd, *J* = 5.0, 2.7 Hz, 1H), 1.84 (dtd, *J* = 12.0, 9.2, 8.7, 4.7 Hz, 2H), 1.79 – 1.67 (m, 2H), 1.67 – 1.49 (m, 8H); $^{13}$C NMR (75 MHz, $CDCl_3$) δ 98.9, 98.7, 68.5, 67.3, 62.2, 62.0, 50.9, 50.6, 44.6, 44.5, 30.5, 30.6, 25.6, 25.4, 19.3, 19.2; HRMS (EI) for $C_8H_{14}O_3$ $[M]^+$: calcd 158.0943, found 158.0947.

**5.2*R***

Colorless liquid oil; analytical TLC, EtOAc/*n*-hexane = 1/3 (v/v), $R_f$ = 0.7, a mixture of 2 diastereomers; $^{1}$H NMR (400 MHz, $CDCl_3$) δ 4.63 – 4.54 (m, 2H), 3.90 – 3.73 (m, 3H), 3.69 – 3.48 (m, 2H), 3.44 (dddd,

*J* = 10.4, 5.2, 3.4, 2.0 Hz, 2H), 3.32 (dd, *J* = 11.7, 6.4 Hz, 1H), 3.10 (dddd, *J* = 7.6, 4.9, 3.9, 2.7 Hz, 2H), 2.72 (ddd, *J* = 5.2, 4.1, 2.0 Hz, 2H), 2.60 (dd, *J* = 5.2, 2.7 Hz, 1H), 2.52 (dd, *J* = 5.0, 2.7 Hz, 1H), 1.82 – 1.71 (m, 2H), 1.71 – 1.59 (m, 2H), 1.59 – 1.37 (m, 8H); $^{13}$C NMR (100 MHz, $CDCl_3$) δ 98.8, 98.7, 68.4, 67.3, 62.1, 61.9, 50.9, 50.5, 44.5, 44.4, 30.4, 30.3, 25.33, 25.31, 19.3, 19.2; HRMS (EI) for $C_8H_{14}O_3$ $[M]^+$: calcd 158.0943, found 158.0942.

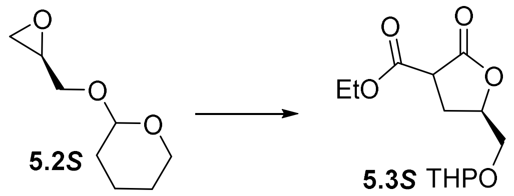

To a solution of freshly prepared NaOEt (101.3 mL, 150 mmol) were added diethyl malonate (22.9 mL, 150 mmol) and **5.2*S*** (15.800 g, 100 mmol). After refluxing for 4 h, the mixture was poured into an iced saturated $NH_4Cl$ aqueous solution and extracted with DCM. The combined organic layer was washed with brine, dried over anhydrous $NaSO_4$, and concentrated under vacuo. Purification with column chromatography using 1/3 (v/v) EtOAc/*n*-hexane gave **5.3*S*** as a colourless liquid (20.400 g, 75% yield). **5.3*R*** was prepared following a similar procedure, with a yield of 69%.

**5.3*S***

Analytical TLC, EtOAc/*n*-hexane = 1/3 (v/v), $R_f$ = 0.2, a mixture of 4 diastereomers; $^1$H NMR (400 MHz, $CDCl_3$) δ 4.66 – 4.38 (m, 2H), 4.07 (q, *J* = 7.1 Hz, 2H), 3.85 – 3.38 (m, 4H), 3.33 (dq, *J* = 11.1, 4.0 Hz, 1H), 2.58 – 2.28 (m, 2H), 1.69 – 1.47 (m, 2H), 1.47 – 1.26 (m, 4H), 1.14 (td, *J* = 7.2, 1.5 Hz, 3H); $^{13}$C NMR (100 MHz, $CDCl_3$) δ 172.0, 171.9, 171.5, 171.4, 167.9, 167.4, 98.9, 98.6, 98.4, 97.9, 77.7, 77.5, 77.4, 77.3, 68.5, 68.2, 68.0, 67.3, 62.0, 61.70, 61.66, 61.6, 61.2, 46.47, 46.45, 46.3, 30.01, 29.9, 29.9, 28.2, 27.7, 27.6, 25.0, 24.9, 19.1, 18.7, 18.5, 13.73, 13.71; HRMS (EI) for $C_{13}H_{20}O_6$ $[M]^+$: calcd 272.1260, found 272.1269.

**5.3*R***

Colorless liquid oil; analytical TLC, EtOAc/*n*-hexane = 1/3 (v/v), $R_f$ = 0.2, a mixture of 4 diastereomers; $^1$H NMR (400 MHz, $CDCl_3$) δ 4.85 – 4.54 (m, 2H), 4.28 – 4.19 (m, 2H), 4.03 – 3.40 (m, 5H), 2.79 – 2.34 (m, 2H), 1.88 – 1.64 (m, 2H), 1.64 – 1.43 (m, 4H), 1.29 (td, *J* = 7.1, 0.8 Hz, 3H); $^{13}$C NMR (100 MHz, $CDCl_3$) δ 172.6, 171.6, 168.2, 167.6, 99.6, 99.1, 99.0, 98.3, 78.0, 77.9, 77.7, 77.6, 67.0, 68.7, 68.3, 67.8, 62.7, 62.3, 62.2, 61.7, 46.9, 46.8, 46.7, 30.4, 30.4, 30.3, 28.6, 28.5, 28.2, 28.1, 25.4, 25.30, 25.28, 19.6,

19.1, 18.9, 14.2, 14.1; HRMS (EI) for $C_{13}H_{20}O_6$ $[M]^+$: calcd 272.1260, found 272.1267.

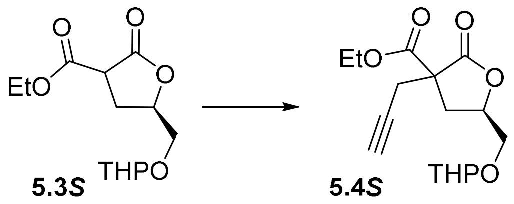

To a solution of NaOEt (65.3 mL, 119 mmol) in EtOH (14.2 mL) was slowly added **5.3*S*** (20.400 g, 75 mmol). After stirring for 1 h, a solution of propargyl bromide (10.8 mL, 142 mmol) in EtOH (14.2 mL) was slowly added. The reaction mixture was then stirred for 40 min, then diluted with water and extracted with DCM. The combined organic layer was then dried over anhydrous $Na_2SO_4$ and evaporated. The crude residue was purified with flash chromatography using 1/5 (v/v) EA/*n*-hexane to give **5.4*S*** as a colourless liquid (7.700 g, 33% yield). **5.4*R*** was prepared via a similar synthetic pathway with a yield of 86%.

**5.4*S***

Colorless liquid oil; analytical TLC, EtOAc/*n*-hexane = 1/3 (v/v), $R_f$ = 0.4, a mixture of 4 diastereomers; $^1$H NMR (400 MHz, $CDCl_3$) δ 4.77 – 4.61 (m, 1H), 4.57 – 4.48 (m, 1H), 4.25 (q, *J* = 7.1 Hz, 2H), 4.03 – 3.79 (m, 2H), 3.74 – 3.47 (m, 2H), 2.95 – 2.85 (m, 2H), 2.85 – 2.36 (m, 2H), 2.11 – 2.01 (m, 1H), 1.89 – 1.67 (m, 2H), 1.64 – 1.48 (m, 4H), 1.29 (t, *J* = 7.1 Hz, 3H); $^{13}$C NMR (100 MHz, $CDCl_3$) δ 172.9, 172.8, 168.6, 99.0, 78.8, 78.4, 77.6, 77.4, 71.85, 71.82, 71.78, 68.9, 68.3, 68.0, 67.4, 62.8, 62.7, 62.6, 62.14, 62.07, 55.1, 54.9, 54.4, 33.3, 33.2, 33.1, 30.4, 30.3, 25.3, 24.8, 24.7, 23.9, 19.2, 19.1, 14.1, 14.0; HRMS (EI) for $C_{16}H_{22}O_6$ $[M]^+$: calcd 310.1416, found 310.1421.

**5.4*R***

Colorless liquid oil; analytical TLC, EtOAc/*n*-hexane = 1/5 (v/v), $R_f$ = 0.3, a mixture of 4 diastereomers; $^1$H NMR (300 MHz, $CDCl_3$) δ 4.77 – 4.61 (m, 1H), 4.57 – 4.47 (m, 1H), 4.17 – 4.04 (m, 2H), 3.89 – 3.62 (m, 2H), 3.61 – 3.30 (m, 2H), 2.84 – 2.62 (m, 2H), 2.60 – 2.19 (m, 2H), 2.05 – 1.92 (m, 1H), 1.75 – 1.26 (m, 6H), 1.20 – 1.05 (m, 3H). $^{13}$C NMR (75 MHz, $CDCl_3$) δ 172.9, 172.8, 168.61, 168.57, 98.94, 98.90, 98.8, 78.8, 77.6, 77.4, 77.2, 72.0, 71.93, 71.90, 68.8, 68.3, 67.9, 67.3, 62.7, 62.6, 62.5, 62.0, 61.9, 60.3, 55.0, 54.8, 33.2, 33.1, 33.0, 30.33, 30.29, 25.4, 24.7, 24.6, 23.8, 21.0, 19.1, 14.2, 14.0, 13.9; HRMS (EI) for $C_{16}H_{22}O_6$ $[M]^+$: calcd 310.1416, found 310.1412.

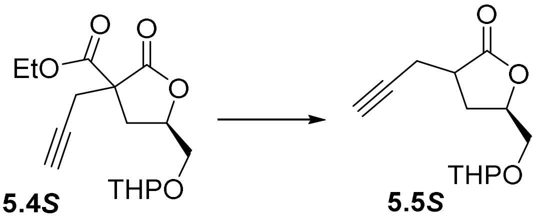

**5.4*S*** → **5.5*S***

To a solution of **5.4*S*** (7.700 g, 24.8 mmol) in DME (81.4 mL) was added 2 M LiOH aqueous solution (6.82 g in 81.4 mL water, 164 mmol). The reaction was heated to 60 °C and stirred for 6 h. After that, the organic solvent was evaporated with a rotary evaporator. The pH of the remaining aqueous phase was adjusted to 3 with 3 N HCl. The compound was extracted 3 times from the acidified mixture using EtOAc. The combined organic layer was dried over anhydrous $Na_2SO_4$ and evaporated. Concentrated residues were purified with flash chromatography using 1/1 (v/v) EtOAc to afford **5.5*S*** as a colourless liquid (2.300 g, 39% yield). **5.5*R*** was prepared similarly with a yield of 64%.

**5.5*S***

Analytical TLC, EtOAc/*n*-hexane = 1/1 (v/v), $R_f$ = 0.5, a mixture of 4 diastereomers; $^1$H NMR (500 MHz, $CDCl_3$) δ 4.72 – 4.48 (m, 2H), 3.88 – 3.65 (m, 2H), 3.66– 3.34 (m, 2H), 3.01 – 2.82 (m, 1H), 2.69 – 1.87 (m, 4H), 2.14 – 1.87 (m, 1H), 1.78 – 1.59 (m, 2H), 1.59 – 1.37 (m, 4H); $^{13}$C NMR (125 MHz, $CDCl_3$) δ 177.8, 177.7, 176.8, 176.6, 99.3, 98.9, 98.8, 98.1, 80.4, 80.30, 80.27, 80.25, 76.8, 70.63, 70.61, 70.60, 70.55, 69.3, 68.62, 68.57, 67.7, 62.4, 62.0, 61.5, 39.6, 39.4, 38.62, 38.57, 30.31, 30.26, 30.21, 30.19, 29.5, 29.42, 29.36, 29.3, 25.3, 25.20, 25.18, 20.19, 20.1, 19.52, 19.48, 19.4, 19.1, 19.0, 18.8; HRMS (EI) for $C_{13}H_{18}O_4$ $[M]^+$: calcd 238.1205, found 238.1209.

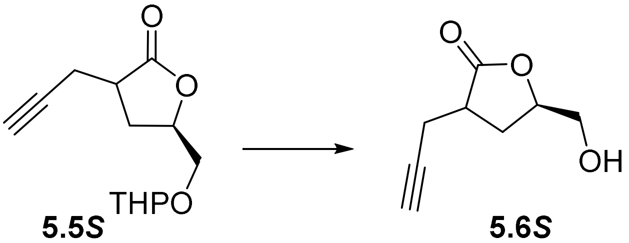

**5.5*S*** → **5.6*S***

To a stirred solution of **5.5*S*** (3.400 g, 14.3 mmol) in MeOH (143 mL) was added *p*-toluenesulfonic acid (190.2 mg, 1.1 mmol). The reaction was refluxed for 3 h and quenched with 200 μL $Et_3N$. The residue was concentrated and purified with chromatography using 1/1 (v/v) EtOAc/*n*-hexane to give **5.5*S*** as a colourless liquid (2.900 g, 86% yield). A yield of 90% was achieved for **5.5*R*** with similar synthetic processes.

**5.6*S***

Analytical TLC, EtOAc/*n*-hexane = 1/1 (v/v), $R_f = 0.3$, a mixture of 2 diastereomers, dr = 9:11; $^1$H NMR (400 MHz, $CDCl_3$) δ 4.64 – 4.43 (m, 1H), 3.87 – 3.74 (m, 1H), 3.70 – 3.48 (m, 2H), 3.00 – 2.77 (m, 1H), 2.63 – 2.18 (m, 4H), 2.07 – 1.92 (m, 1H); $^{13}$C NMR (100 MHz, $CDCl_3$) δ 178.6, 177.4, 80.3, 79.3, 79.1, 70.8, 70.7, 64.1, 63.4, 39.7, 38.7, 28.6, 28.5, 20.1, 19.3; HRMS (EI) for $C_8H_{10}O_3$ $[M]^+$: calcd 154.0630, found 154.0633.

**5.6*R***

Colorless liquid oil; analytical TLC, EtOAc/*n*-hexane = 1/1 (v/v), $R_f = 0.3$, a mixture of 2 diastereomers, dr = 1:1.5; $^1$H NMR (400 MHz, $CDCl_3$) δ 4.68 – 4.42 (m, 1H), 3.88 (ddd, *J* = 9.5, 6.5, 3.3 Hz, 1H), 3.70 – 3.53 (m, 2H), 3.08 – 2.72 (m, 1H), 2.68 – 2.23 (m, 4H), 2.18 – 1.90 (m, 1H); $^{13}$C NMR (100 MHz, $CDCl_3$) δ 178.6, 177.4, 80.3, 80.2, 79.3, 79.1, 70.83, 70.77, 64.3, 63.6, 39.9, 38.9, 28.7, 28.6, 20.3, 19.5; HRMS (EI) for $C_8H_{10}O_3$ $[M]^+$: calcd 154.0630, found 154.0632.

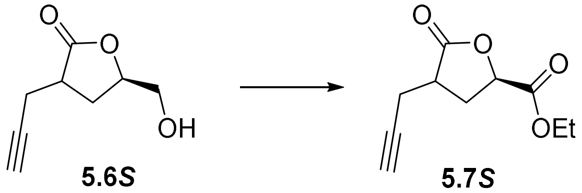

To a stirred solution of **5.6*S*** (1.890 g, 12.3 mmol) in acetone (105.8 mL) at 0 ˚C was added a solution of $CrO_3$ (4.218 g, 42 mmol) in 1.5 M $H_2SO_4$ (18.26 mL) dropwise. The mixture was stirred overnight, followed by the addition of *i*-PrOH (27.1 mL) and stirring for 1 h. The mixture was then diluted in EtOAc, washed with $H_2O$, dried over anhydrous $Na_2SO_4$, and evaporated to afford a crude acid as a white sticky solid (1.150 g, 56% yield). The crude acid from **5.6*R*** has a yield of 38%.

The crude product (1.150 g, 6.9 mmol) was dissolved in anhydrous DMF (22.15 mL), followed by the addition of $NaHCO_3$ (574.1 mg, 13.8 mmol) and EtI (2.75 mL, 69 mmol) subsequently. The mixture was stirred at rt under an Ar atmosphere for 2 days. The resulting mixture was diluted in EtOAc, washed with 1 M HCl, saturated $NaHCO_3$ aqueous solution, water, and brine. The combined organic layer was dried over anhydrous $Na_2SO_4$ and evaporated. Residues were separated with column chromatography using 1/5–1/4 (v/v) EtOAc/*n*-hexane to afford **5.7*S*-1** and **5.7*S*-2** as colourless liquids (**5.7*S*-1**, 122 mg, 9% yield;

**5.7*S*-2**, 163 mg, 12% yield). **5.7*R*-1** and **5.7*R*-2** were prepared similarly from 5.6*R* with yield of 5% and 15%, respectively.

**5.7*S*-1 (5*RS*cy)**

Analytical TLC, EtOAc/*n*-hexane = 1/3 (v/v), $R_f = 0.3$; $^1H$ NMR (500 MHz, $CDCl_3$) δ 4.97 – 4.89 (m, 1H), 4.26 (qd, *J* = 7.2, 3.2 Hz, 2H), 2.92 (tdd, *J* = 10.1, 7.1, 4.6 Hz, 1H), 2.71 – 2.49 (m, 4H), 2.04 (t, *J* = 2.7 Hz, 1H), 1.31 (t, *J* = 7.1 Hz, 3H); $^{13}C$ NMR (125 MHz, $CDCl_3$) δ 176.4, 170.0, 79.6, 74.2, 71.3, 62.3, 37.0, 31.2, 19.6, 14.2; HRMS (EI) for $C_{10}H_{12}O_4$ $[M]^+$: calcd 196.0736, found 196.0733.

**5.7*S*-2 (5*RR*cy)**

Analytical TLC, EtOAc/*n*-hexane = 1/3 (v/v), $R_f = 0.2$; $^1H$ NMR (500 MHz, $CDCl_3$) δ 4.83 (dd, *J* = 9.3, 7.1 Hz, 1H), 4.27 (q, *J* = 7.2 Hz, 2H), 2.95 – 2.65 (m, 3H), 2.53 (ddd, *J* = 17.2, 8.1, 2.6 Hz, 1H), 2.26 (ddd, *J* = 12.8, 10.0, 9.2 Hz, 1H), 2.04 (t, *J* = 2.6 Hz, 1H), 1.31 (t, *J* = 7.1 Hz, 3H); $^{13}C$ NMR (125 MHz, $CDCl_3$) δ 175.6, 169.2, 79.8, 74.3, 71.1, 62.2, 39.0, 31.2, 19.8, 14.2; HRMS (EI) for $C_{10}H_{12}O_4$ $[M]^+$: calcd 196.0736, found 196.0734; $[\alpha]_D^{22}$ −55 (*c* 0.9, $CHCl_3$).

**5.7*R*-1 (5*SR*cy)**

Colorless liquid oil; analytical TLC, EtOAc/n-hexane = 1/2 (v/v), $R_f = 0.6$; $^1H$ NMR (500 MHz, $CDCl_3$) δ 4.98 – 4.89 (m, 1H), 4.27 (qd, *J* = 7.2, 3.3 Hz, 2H), 2.93 (tdd, *J* = 10.0, 7.1, 4.6 Hz, 1H), 2.71 – 2.51 (m, 4H), 2.05 (t, *J* = 2.7 Hz, 1H), 1.32 (t, *J* = 7.2 Hz, 3H); $^{13}C$ NMR (125 MHz, $CDCl_3$) δ 176.4, 170.0, 79.6, 74.2, 71.3, 62.3, 37.0, 31.2, 19.6, 14.2; HRMS (EI) for $C_{10}H_{12}O_4$ $[M]^+$: calcd 196.0736, found 196.0738.

**5.7*R*-2 (5*SS*cy)**

Colorless liquid oil; analytical TLC, EtOAc/n-hexane = 1/2 (v/v), $R_f = 0.5$; $^1H$ NMR (500 MHz, $CDCl_3$) δ 4.84 (dd, *J* = 9.2, 7.1 Hz, 1H), 4.32 – 4.24 (m, 2H), 2.94 – 2.77 (m, 2H), 2.73 (ddd, *J* = 17.1, 4.4, 2.7 Hz, 1H), 2.54 (ddd, *J* = 17.2, 8.2, 2.6 Hz, 1H), 2.35 – 2.21 (m, 1H), 2.05 (t, *J* = 2.7 Hz, 1H), 1.32 (t, *J* = 7.1 Hz, 3H); $^{13}C$ NMR (100 MHz, $CDCl_3$) δ 175.6, 169.0, 79.6, 73.9, 70.9, 61.7, 38.5, 30.6, 19.2, 13.8; HRMS (EI) for $C_{10}H_{12}O_4$ $[M]^+$: calcd 196.0736, found 196.0736.

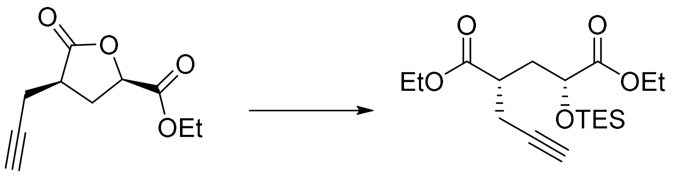

**5.7*S*-1 = 5*RS*cy** **5.8*S*-1**

To a stirred solution of **5.7*S*-1** (155.5 mg, 0.8 mmol) in EtOH (11 mL) was added concentrated $H_2SO_4$ (550 μL). The mixture was refluxed overnight, then diluted in EtOAc, washed with water, brine, dried over anhydrous $Na_2SO_4$, and evaporated. Residues were purified before the next step using column chromatography with 1/4 v/v EtOAc/*n*-hexane to afford a mixture of **5*S*-1** and **5.7*S*-1**.

The purified mixture was dissolved in anhydrous DCM (6 mL). To the stirred solution were added imidazole (53.1 mg, 0.8 mmol) and TESCl (113.5 μL, 0.9 mmol) subsequently at 0℃ under an Ar atmosphere. The mixture was stirred at rt for 15 min, then diluted in EtOAc and washed with 0.3 M HCl and brine, dried over anhydrous $Na_2SO_4$, and concentrated. Purification with column chromatography using 1/20 v/v EtOAc/*n*-hexane afforded **5.8*S*-1** as a colourless liquid (146 mg, 50% yield after 2 steps). **5.8*S*-2**, **5.8*R*-1** and **5.8*R*-2** were similarly prepared, with yield of 63%, 44%, and 33% for 2 steps, respectively.

**5.8*S*-1**

Analytical TLC, EtOAc/*n*-hexane = 1/19 (v/v), $R_f$ = 0.5; $^1$H NMR (400 MHz, $CDCl_3$) δ 4.27 (dd, *J* = 9.1, 3.7 Hz, 1H), 4.23 – 4.07 (m, 4H), 2.85 – 2.71 (m, 1H), 2.51 – 2.42 (m, 2H), 2.19 (ddd, *J* = 13.8, 10.0, 3.8 Hz, 1H), 1.98 (t, *J* = 2.7 Hz, 1H), 1.89 (ddd, *J* = 13.6, 9.1, 3.9 Hz, 1H), 1.25 (td, *J* = 7.2, 1.6 Hz, 6H), 0.92 (t, *J* = 7.9 Hz, 9H), 0.70 – 0.53 (m, 6H); $^{13}$C NMR (100 MHz, $CDCl_3$) δ 173.7, 173.4, 80.7, 70.5, 70.2, 61.0, 60.8, 40.2, 36.2, 22.1, 14.28, 14.26, 6.8, 4.6; HRMS (EI) for $C_{18}H_{32}O_5Si$ $[M]^+$: calcd 365.2019, found 365.2020.

**5.8*S*-2**

Colorless liquid oil; analytical TLC, EtOAc/*n*-hexane = 1/19 (v/v), $R_f$ = 0.5; $^1$H NMR (500 MHz, $CDCl_3$) δ 4.27 (t, *J* = 6.5 Hz, 1H), 4.16 (dt, *J* = 10.8, 6.9 Hz, 4H), 2.73 (p, *J* = 6.7 Hz, 1H), 2.50 (dd, *J* = 6.5, 2.5 Hz, 2H), 2.16 (dt, *J* = 14.2, 7.1 Hz, 1H), 2.06 – 2.00 (m, 1H), 1.98 (s, 1H), 1.26 (q, *J* = 7.4 Hz, 6H), 0.95 (t, *J* = 8.0 Hz, 9H), 0.61 (q, *J* = 7.9 Hz, 6H); $^{13}$C NMR (125 MHz, $CDCl_3$) δ 173.8, 173.2, 80.9, 70.4, 70.1, 61.1, 60.9, 40.3, 35.8, 21.3, 14.31, 14.27, 6.8, 4.7, 4.7; HRMS (EI) for $C_{18}H_{32}O_5Si$ $[M]^+$: calcd 365.2019, found 365.2021.

**5.8*R*-1**

Colorless liquid oil; analytical TLC, EtOAc/*n*-hexane = 1/19 (v/v), $R_f$ = 0.5; $^1$H NMR (500 MHz, $CDCl_3$) δ 4.29 (dd, *J* = 9.1, 3.8 Hz, 1H), 4.25 – 4.08 (m, 4H), 2.80 (ddt, J = 10.4, 6.6, 3.3 Hz, 1H), 2.49 (ddd, *J* = 6.8, 2.7, 1.6 Hz, 2H), 2.22 (ddd, *J* = 13.8, 10.0, 3.8 Hz, 1H), 2.00 (t, *J* = 2.7 Hz, 1H), 1.91 (ddd, *J* = 13.8, 9.1, 3.9 Hz, 1H), 1.28 (td, *J* = 7.1, 1.9 Hz, 6H), 0.95 (t, *J* = 7.9 Hz, 9H), 0.61 (qd, *J* = 8.0, 1.6 Hz, 6H); $^{13}$C NMR (125 MHz, $CDCl_3$) δ 173.8, 173.5, 70.5, 70.3, 61.1, 60.9, 40.3, 36.3, 22.2, 14.4, 14.3, 6.8, 4.7; HRMS (EI) for $C_{18}H_{32}O_5Si$ $[M]^+$: calcd 365.2019, found 365.2024.

**5.8*R*-2**

Colorless liquid oil; analytical TLC, EtOAc/*n*-hexane = 1/19 (v/v), $R_f$ = 0.5; $^1$H NMR (500 MHz, $CDCl_3$) δ 4.27 (dd, *J* = 7.3, 5.7 Hz, 1H), 4.21 – 4.10 (m, 4H), 2.74 (p, *J* = 6.7 Hz, 1H), 2.51 (ddd, *J* = 6.8, 2.7, 1.0 Hz, 2H), 2.21 – 2.13 (m, 1H), 2.06 – 2.00 (m, 1H), 1.99 (t, *J* = 2.6 Hz, 1H), 1.27 (q, *J* = 7.3 Hz, 6H), 0.95 (t, *J* = 7.9 Hz, 9H), 0.66 – 0.58 (m, 6H); $^{13}$C NMR (125 MHz, $CDCl_3$) δ 173.8, 173.2, 80.9, 70.4, 70.2, 61.1, 60.9, 40.3, 35.8, 21.3, 14.33, 14.30, 6.8, 4.7; HRMS (EI) for $C_{18}H_{32}O_5Si$ $[M]^+$: calcd 365.2019, found 365.2023.

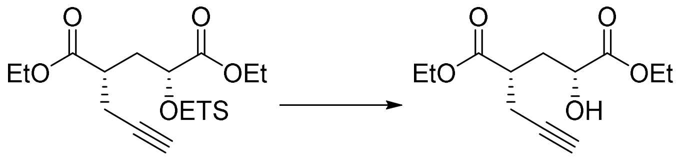

**5.8*S*-1** **5*S*-1**

To a stirred solution of **5.8*S*-1** (108.5 mg, 0.3 mmol) in EtOH (3 mL) was added PPTS (7.6 mg, 0.03 mmol). The mixture was stirred at rt for 2.5 h, then quenched with saturated $NaHCO_3$ aqueous solution and extracted with DCM. The combined organic layer was washed with water and brine, dried over anhydrous $Na_2SO_4$, and concentrated. Column chromatography using 1/3 EtOAc/*n*-hexane afforded **5*S*-1** as a colourless liquid (46.5 mg, 64% yield). **5*S*-2**, **5*R*-1** and **5*R*-2** were similarly prepared, with yield of 73%, 35% and 50%, respectively.

**5*S*-1 (5*RS*)**

Analytical TLC, EtOAc/*n*-hexane = 1/3 (v/v), $R_f$ = 0.2; $^1$H NMR (500 MHz, $CDCl_3$) δ 4.29 – 4.11 (m, 5H), 2.92 (s, 1H), 2.88 – 2.79 (m, 1H), 2.50 (qdd, *J* = 16.9, 6.8, 2.7 Hz, 2H), 2.27 (ddd, *J* = 13.7, 9.9, 3.5 Hz, 1H), 2.00 (t, *J* = 2.7 Hz, 1H), 1.86 (ddd, *J* = 13.7, 9.5, 3.8 Hz, 1H), 1.27 (dt, *J* = 14.0, 7.1 Hz, 6H); $^{13}$C NMR

(125 MHz, $CDCl_3$) δ 174.8, 173.9, 80.8, 70.5, 68.7, 62.0, 61.0, 40.5, 35.4, 21.8, 14.28, 14.26; HRMS (EI) for $C_{12}H_{18}O_5$ $[M]^+$: calcd 242.1154, found 242.1155; $[\alpha]_D^{22}$ +9.9 (*c* 1.2, $CHCl_3$).

**5*S*-2 (5*RR*)**

Colorless liquid oil; analytical TLC, EtOAc/*n*-hexane = 1/3 (v/v), $R_f$ = 0.2; $^1$H NMR (400 MHz, $CDCl_3$) δ 4.27 – 4.07 (m, 5H), 2.92 (d, *J* = 5.9 Hz, 1H), 2.81 (p, *J* = 6.7 Hz, 1H), 2.52 (dd, *J* = 6.5, 2.7 Hz, 2H), 2.18 – 2.01 (m, 2H), 2.00 (t, *J* = 2.6 Hz, 1H), 1.26 (t, *J* = 14.1 Hz, 6H); $^{13}$C NMR (100 MHz, $CDCl_3$) δ 174.6, 174.0, 80.7, 70.5, 68.7, 61.9, 60.9, 40.7, 35.0, 20.9, 14.23, 14.21; HRMS (EI) for $C_{12}H_{18}O_5$ $[M]^+$: calcd 242.1154, found 242.1154.

**5*R*-1 (5*SR*)**

Colorless liquid oil; analytical TLC, EtOAc/n-hexane = 1/2 (v/v), $R_f$ = 0.5; $^1$H NMR (500 MHz, $CDCl_3$) δ 4.29 – 4.15 (m, 5H), 2.92 – 2.79 (m, 2H), 2.62 – 2.46 (m, 2H), 2.30 (ddd, *J* = 13.7, 10.0, 3.4 Hz, 1H), 2.02 (t, *J* = 2.7 Hz, 1H), 1.89 (ddd, *J* = 14.1, 9.5, 3.9 Hz, 1H), 1.36 – 1.23 (m, 6H); $^{13}$C NMR (125 MHz, $CDCl_3$) δ 174.8, 173.9, 80.9, 70.6, 68.7, 62.1, 61.1, 40.5, 35.5, 21.9, 14.4, 14.3; HRMS (EI) for $C_{12}H_{18}O_5$ $[M]^+$: calcd 242.1154, found 242.1156.

**5*R*-2 (5*SS*)**

Colorless liquid oil; analytical TLC, EtOAc/n-hexane = 1/2 (v/v), $R_f$ = 0.5; $^1$H NMR (500 MHz, $CDCl_3$) δ 4.30 – 4.12 (m, 5H), 2.84 (p, *J* = 6.7 Hz, 1H), 2.80 (d, *J* = 5.9 Hz, 1H), 2.55 (dd, *J* = 6.5, 2.7 Hz, 2H), 2.15 (ddd, *J* = 14.2, 6.8, 4.3 Hz, 1H), 2.07 (ddd, *J* = 14.2, 9.2, 7.1 Hz, 1H), 2.02 (t, *J* = 2.7 Hz, 1H), 1.31 (t, *J* = 7.1 Hz, 3H), 1.27 (t, *J* = 7.2 Hz, 3H); $^{13}$C NMR (125 MHz, $CDCl_3$) δ 174.7, 174.1, 80.8, 70.6, 68.8, 62.1, 61.1, 40.8, 35.1, 21.0, 14.3, 14.3; HRMS (EI) for $C_{12}H_{18}O_5$ $[M]^+$: calcd 242.1154, found 242.1157.

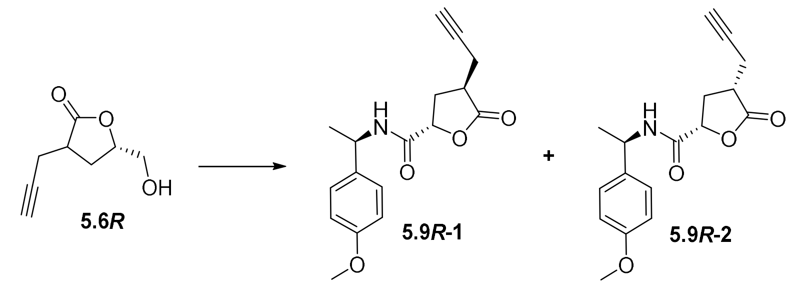

To a stirred solution of **5.6*R*** (159.9 mg, 1.04 mmol) in acetone (8.9 mL) at 0 °C was added a solution of $CrO_3$ (357 g, 3.6 mmol) in 1.5 M $H_2SO_4$ (1.54 mL) dropwise. The mixture was stirred overnight, followed by the addition of *i*-PrOH (2.3 mL) and stirring for 1 h. The mixture was then diluted in EtOAc, washed with $H_2O$, dried over anhydrous $Na_2SO_4$, and evaporated to afford a crude acid as a white sticky solid.

To a stirred solution of the resulting crude product in anhydrous DMF/DCM (0.4 mL/1.53 mL) were added *N*-methylmorpholine (0.4 mL, 3.5 mmol), HOBt (71.9 mg, 0.4 mmol), and EDCI (81.0 mg, 0.44 mmol) subsequently under an Ar atmosphere. The mixture was stirred at rt overnight, then diluted in EtOAc. The combined organic layer was washed with saturated $NaHCO_3$ aqueous solution, water, and brine. The residue was dried over anhydrous $Na_2SO_4$, concentrated under vacuo, then purified with column chromatography using 1/2 (v/v) EtOAc/*n*-hexane to afford 2 diastereomers as white solid products (**5.9*R*-1**, 16.3 mg, 5% yield; **5.9*R*-2**, 5.4 mg, 2% yield).

**5.9*R*-1**

White solid, m.p. 145–150 °C, $R_f$= 0.7 (1/1 v/v EtOAc/*n*-hexane); $^1$H NMR (500 MHz, $CDCl_3$) δ 7.18 (d, *J* = 8.6 Hz, 2H), 6.86 (d, *J* = 8.7 Hz, 2H), 6.53 (d, *J* = 8.1 Hz, 1H), 5.08 (p, *J* = 7.1 Hz, 1H), 4.92 (dd, *J* = 9.1, 3.2 Hz, 1H), 3.79 (s, 3H), 2.73 – 2.65 (m, 2H), 2.61 – 2.53 (m, 3H), 2.05 (t, *J* = 2.7 Hz, 1H), 1.49 (d, *J* = 7.0 Hz, 3H); $^{13}$C NMR (125 MHz, $CDCl_3$) δ 176.4, 168.3, 159.2, 134.6, 127.3, 114.4, 79.3, 76.0, 71.4, 55.5, 48.6, 37.4, 30.5, 21.8, 19.7; HRMS (EI) for $C_{17}H_{19}O_4N$ $[M]^+$: calcd 301.1314, found 301.1315.

**5.9*R*-2**

White solid, m.p. 136–147 °C, $R_f$= 0.6 (1/1 v/v EtOAc/*n*-hexane); $^1$H NMR (400 MHz, $CDCl_3$) δ 7.22 (d, *J* = 8.7 Hz, 2H), 6.86 (d, *J* = 8.7 Hz, 2H), 6.53 (d, *J* = 8.2 Hz, 1H), 5.12 (p, *J* = 7.2 Hz, 1H), 4.79 (dd, *J* = 10.0, 6.6 Hz, 1H), 3.79 (s, 3H), 2.94 – 2.82 (m, 2H), 2.63 – 2.52 (m, 2H), 2.24 – 2.08 (m, 1H), 1.80 (t, *J* = 2.6 Hz, 1H), 1.51 (d, *J* = 7.0 Hz, 3H); $^{13}$C NMR (100 MHz, $CDCl_3$) δ 175.8, 167.9, 159.1, 134.7, 127.5,

114.2, 79.4, 75.6, 71.5, 55.4, 48.1, 39.3, 31.3, 21.7, 19.5; HRMS (EI) for $C_{17}H_{19}O_4N$ $[M]^+$: calcd 301.1314, found 301.1313.

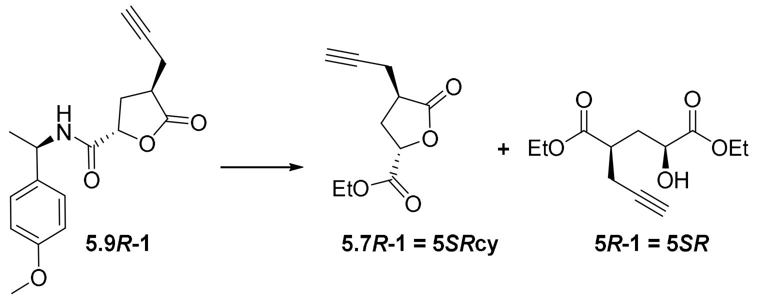

**5.9*R*-1** **5.7*R*-1 = 5*SR*cy** **5*R*-1 = 5*SR***

To a stirred solution of **5.9*R*-1** (16.3 mg, 0.054 mmol) in MeCN/$H_2O$ (2.1 mL/0.6 mL) was added CAN (164.5 mg, 0.3 mmol). The mixture was stirred at rt for 40 min, then diluted with $H_2O$ and extracted with EtOAc. The combined organic layer was washed with water and brine and dried over anhydrous $Na_2SO_4$. The residue was concentrated under vacuo and purified with column chromatography using 3/1 (v/v) EtOAc/*n*-hexane to afford deprotected lactone acid as a colourless sticky oil (5.5 mg, 60% yield, $R_f$ = 0.18 with 2/1 v/v EtOAc/*n*-hexane). 5**.9*R*-2** was similarly deprotected and purified with a yield of 44%.

The residue (3.2 mg, 0.02 mmol) was dissolved in EtOH (1 mL), followed by the addition of 1 drop of concentrated $H_2SO_4$. The mixture was refluxed overnight, diluted in EtOAc and washed with saturated $NaHCO_3$ aqueous solution and brine. The combined organic layer was dried over anhydrous $Na_2SO_4$ and evaporated. The residue was purified with column chromatography using 1/3 v/v EtOAc/*n*-hexane to afford 2 light-yellow sticky liquid products with the same NMR as **5*R*-1** (1.8 mg, 37% yield) and **5.7*R*-1** (0.9 mg, 23% yield). **5.9*R*-2** was similarly deprotected and esterified, giving same products as **5*R*-2** (63%) and **5.7*R*-2** (27%) after TESCl derivatization, separation, and deprotection.

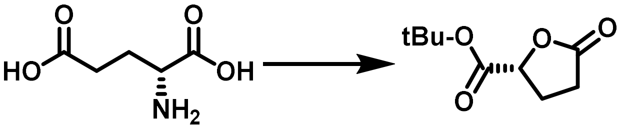

**C5-1** **C5-2**

To a stirred solution of **C5-1** (14.600 g, 100 mmol) in EtOH (420 mL) was added AcCl (0.7 mL, 10 mmol) dropwise. The reaction was stirred at rt overnight underan Ar atmosphere. The resulting mixture was concentrated under vacuum to afford a crude lactone acid.

The crude acid was dissolved in anhydrous THF (100 mL), followed by the addition of tBuOH (11.4 mL, 120 mmol), DMAP (1.200 g, 10 mmol) and DCC (24.800 g, 120 mmol) sequentially in an ice-bath. The reaction was warmed to rt and stirred overnight, then diluted in EtOAc, filtered over celite, and purified with column chromatography using 1/3 EtOAc/*n*-hexane to afford **C5-2** as a colourless liquid (13.400 g, 72% yield).

##### *R*2HG donors

### C5-2

Analytical TLC: EtOAc/*n*-hexane = 1/3, $R_f$ = 0.5; $^1$H NMR (500 MHz, $CDCl_3$) δ 4.85 – 4.77 (m, 1H), 2.66 – 2.44 (m, 3H), 2.31 – 2.21 (m, 1H), 1.49 (d, *J* = 2.4 Hz, 9H); $^{13}$C NMR (75 MHz, $CDCl_3$) δ 176.4, 169.1, 83.1, 76.3, 27.9, 26.8, 25.9; HRMS (EI) for $C_9H_{14}O_4$ $[M]^+$: calcd 186.0892, found 186.0887.

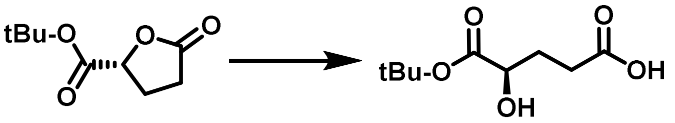

C5-2 C5-3

To a stirred solution of **C5-2** (6.700 g, 36 mmol) in dry THF (36 mL) was added 1 M KOH aqueous solution (41.5 mL, 60 mmol) in an ice bath. The reaction was stirred while warming to rt for 1 h 10 min, then concentrated, acidified with 3 N HCl to pH = 3, and extracted with EtOAc 3 times. Combined organic layer was dried over anhydrous $Na_2SO_4$, then evaporated and purified with chromatography using 1/9 MeOH/DCM containing 1% AcOH to afford **C5-3** as a thick yellowish oil (3.000 g, 42% yield).

### C5-3

Analytical TLC: MeOH/DCM = 1/9, with 0.5 % AcOH, $R_f$ = 0.3; $^1$H NMR (400 MHz, $CD_3OD$) δ 4.06 (dd, *J* = 8.1, 4.6 Hz, 1H), 2.54 – 2.31 (m, 2H), 2.18 – 1.93 (m, 1H), 1.88 – 1.67 (m, 1H), 1.49 (s, 9H); $^{13}$C NMR (100 MHz, $CD_3OD$) δ 175.4, 173.5, 81.2, 69.7, 29.2, 29.1, 26.9; HRMS (EI) for $C_9H_{16}O_5$ $[M]^+$: calcd 204.0998, found 204.0995.

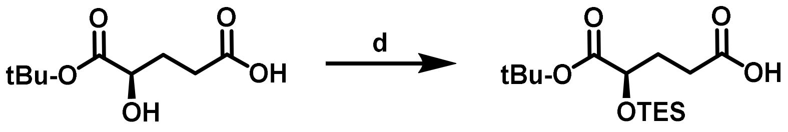

C5-3 C5-4

To a stirred slurry of **C5-3** (1.020 g, 5 mmol) in anhydrous DCM (50 mL) were added imidazole (885 mg, 13 mmol) and TESCl (1.93 mL, 12 mmol) in an ice bath. The mixture was stirred at rt for 15 min, then diluted in EtOAc, washed with 0.3 M HCl and brine, then concentrated and purified with column chromatography using 1/19 EtOAc/*n*-hexane to afford **C5-4** as a colourless liquid (1.440 g, 90% yield).

**C5-4**

Analytical TLC: EtOAc/*n*-hexane = 1/4, $R_f$ = 0.7; $^1$H NMR (500 MHz, $CDCl_3$) δ 11.14 (s, 1H), 4.12 (dd, *J* = 7.2, 4.7 Hz, 1H), 2.53 – 2.34 (m, 2H), 2.08 – 1.85 (m, 2H), 1.42 (s, 9H), 0.91 (t, *J* = 8.0 Hz, 9H), 0.58 (q, *J* = 8.0 Hz, 6H); $^{13}$C NMR (125 MHz, $CDCl_3$) δ 179.6, 172.4, 81.4, 71.1, 30.0, 29.5, 28.0, 6.7, 4.6; HRMS (EI) for $C_{15}H_{30}O_5Si$ $[M]^+$: calcd 318.1863, found 318.1868.

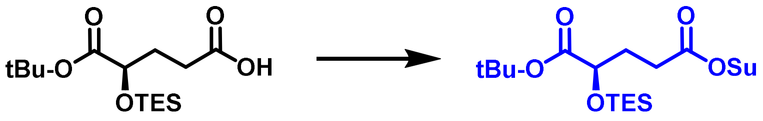

C5-4 C5-donor

To a stirred solution of **C5-4** (1.440 g, 4.5 mmol) in anhydrous DCM (25 mL) in an ice bath were added NHS (5.180 g, 45 mmol) and DCC (4.640 g, 22.5 mmol). The mixture was stirred at rt overnight, then diluted in EtOAc and filtered over Celite. Combined organic layer was concentrated and purified with column chromatography using 1/4 (v/v) EtOAc/*n*-hexane to afford **C5-donor** as a colourless liquid (1.200 g, 64 % yield).

**C5-donor**

Analytical TLC, EtOAc/*n*-hexane = 1/4, $R_f$ = 0.5; $^1$H NMR (400 MHz, $CDCl_3$) δ 4.16 (dd, *J* = 7.3, 4.6 Hz, 1H), 2.82 (s, 4H), 2.79 – 2.61 (m, 2H), 2.19 – 2.00 (m, 2H), 1.45 (s, 9H), 0.95 (t, *J* = 7.9 Hz, 9H), 0.61 (q, *J* = 7.9 Hz, 6H); $^{13}$C NMR (100 MHz, $CDCl_3$) δ 172.0, 169.2, 168.5, 81.6, 77.5, 77.2, 76.9, 70.7, 30.0, 28.1, 28.0, 26.6, 25.7, 6.8, 4.7; HRMS (EI) for $C_{19}H_{33}O_7Si$ $[M]^+$: calcd 415.2026, found 415.2032.

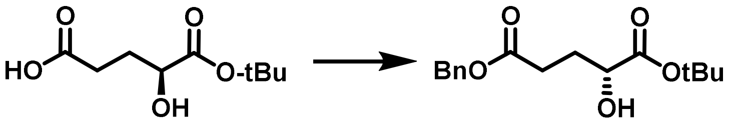

**C5-3** **C1-2**

To a stirred solution of **C5-3** (204.2 mg, 1 mmol) in anhydrous DMF (2 mL) were added $K_2CO_3$ (152 mg, 1.1 mmol) and BnBr (130 μL, 1.1 mmol). The reaction was stirred at rt overnight, then diluted in EtOAc and washed with saturated $NaHCO_3$ aqueous solution and brine. The combined organic layer was azeotroped with toluene, then purified with column chromatography using 1/4 EtOAc/*n*-hexane to afford **C1-2** as a colourless liquid (159 mg, 54% yield).

**C1-2**

Analytical TLC, EtOAc/*n*-hexane = 1/4, $R_f$ = 0.3; $^1$H NMR (400 MHz, $CDCl_3$) δ 7.40 – 7.29 (m, 5H), 5.13 (s, 2H), 4.09 (ddd, *J* = 7.6, 5.5, 4.1 Hz, 1H), 2.85 (d, *J* = 5.5 Hz, 1H), 2.61 – 2.41 (m, 2H), 2.23 – 2.05 (m, 1H), 1.96 – 1.85 (m, 1H), 1.48 (s, 9H); $^{13}$C NMR (100 MHz, $CDCl_3$) δ 174.0, 173.1, 136.0, 128.6, 128.30, 128.27, 82.8, 69.7, 66.4, 29.8, 29.6, 28.1; HRMS (EI) for $C_{16}H_{22}O_5$ $[M]^+$: calcd 294.1467, found 294.2471.

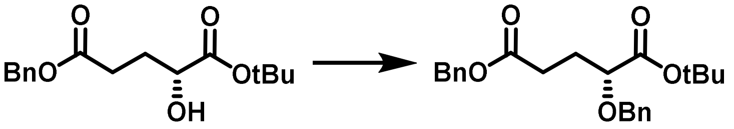

**C1-2** **C1-3**

To a stirred solution of **C1-2** (318.5 mg, 1.08 mmol) in EtOAc (2 mL) was added $Ag_2O$ (231.7 mg, 1.62 mmol), and the resulting solution was stirred at rt under Ar atmosphere for 30 min, followed by the addition of BnBr (178 μL, 1.62 mmol). The reaction was stirred for 3 d, then filtered over celite and purified with column chromatography using 1/20 EtOAc/*n*-hexane to afford **C1-3** as a colourless liquid (191 mg, 46% yield).

**C1-3**

Analytical TLC: EtOAc/*n*-hexane = 1/9, $R_f$ = 0.7; $^1$H NMR (500 MHz, $CDCl_3$) δ 7.41 – 7.25 (m, 10H), 5.09 (s, 2H), 4.70 (d, *J* = 11.4 Hz, 1H), 4.38 (d, *J* = 11.4 Hz, 1H), 3.87 (dd, *J* = 8.5, 4.4 Hz, 1H), 2.52 (t, *J* = 7.6 Hz, 2H), 2.22 – 1.99 (m, 2H), 1.49 (s, 9H); $^{13}$C NMR (125 MHz, $CDCl_3$) δ 172.9, 171.5, 137.7, 136.0,

128.68, 128.65, 128.5, 128.3, 128.2, 127.9, 81.8, 77.27, 72.33, 66.4, 30.1, 28.2, 28.1; HRMS (EI) for $C_{23}H_{28}O_5$ $[M]^+$: calcd 384.1937, found 384.1942.

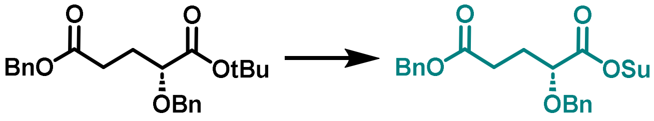

**C1-3** **C1-donor**

**C1-3** (2.660 g, 6.9 mmol) was dissolved in neat TFA (6.9 mL) at 0 °C, followed by stirring for 0.5 h. The reaction was diluted in water and extracted with EtOAc 3 times. The combined organic layer was concentrated and azeotroped with toluene to afford the crude acid for the next step.

The crude was stirred in anhydrous DCM (70 mL), followed by the addition of NHS (7.900 g, 69 mmol) and DCC (7.100 g, 34.5 mmol) in an ice bath. The mixture was stirred under an Ar atmosphere overnight, then diluted with EtOAc and filtered over Celite. The combined organic layer was concentrated and purified using column chromatography using 1/4 (v/v) EtOAc to afford **C1-donor** as a colorless liquid (2.100 g, 72% yield).

**C1-donor**

Analytical TLC, EtOAc/*n*-hexane = 1/4, $R_f$ = 0.3; $^1$H NMR (400 MHz, $CDCl_3$) δ 7.40 – 7.28 (m, 10H), 5.08 (s, 2H), 4.79 (d, *J* = 11.4 Hz, 1H), 4.46 (d, *J* = 11.4 Hz, 1H), 4.37 (dd, *J* = 7.9, 4.8 Hz, 1H), 2.87 (s, 4H), 2.66 – 2.53 (m, 2H), 2.39 – 2.16 (m, 2H); $^{13}$C NMR (100 MHz, $CDCl_3$) δ 172.3, 168.8, 167.9, 136.6, 135.9, 128.59, 128.61, 128.4, 128.29, 128.27, 74.9, 72.8, 66.4, 29.4, 28.2, 25.7. HRMS (EI) for $C_{23}H_{23}O_7N$ $[M]^+$: calcd 425.1475, found 425.1479.

**GSTP1_170-210**

LC-MS (ESI$^+$) for $C_{203}H_{324}N_{54}O_{55}S$ (GSTP1_170-210_WT): $[M+5H]^{5+}$: calcd 887.487, found 887.558; $[M+3H+2Na]^{5+}$: calcd 896.280, found 895.686; $[M+4H]^{4+}$: calcd 1109.107, found 1109.380; $[M+2H+2Na]^{4+}$: calcd 1120.098, found 1119.371; $[M+3H]^{3+}$: calcd 1478.474, found 1478.699; $[M+H+2Na]^{3+}$: calcd 1493.129, found 1491.992.

LC-MS (ESI$^+$) for $C_{208}H_{330}N_{54}O_{59}S$ (GSTP1_170-210_K209-*R*2HG): $[M+5H]^{5+}$: calcd 913.493, found 913.635; $[M+3H+2Na]^{5+}$: calcd 922.286, found 921.847; $[M+4H]^{4+}$: calcd 1141.614, found 1141.723;

[M+2H+2Na]$^{4+}$: calcd 1152.605, found 1151.882; [M+3H]$^{3+}$: calcd 1521.816, found 1522.050; [M+H+2Na]$^{3+}$: calcd 1536.471, found 1535.427.

**NMR**

**1.4**

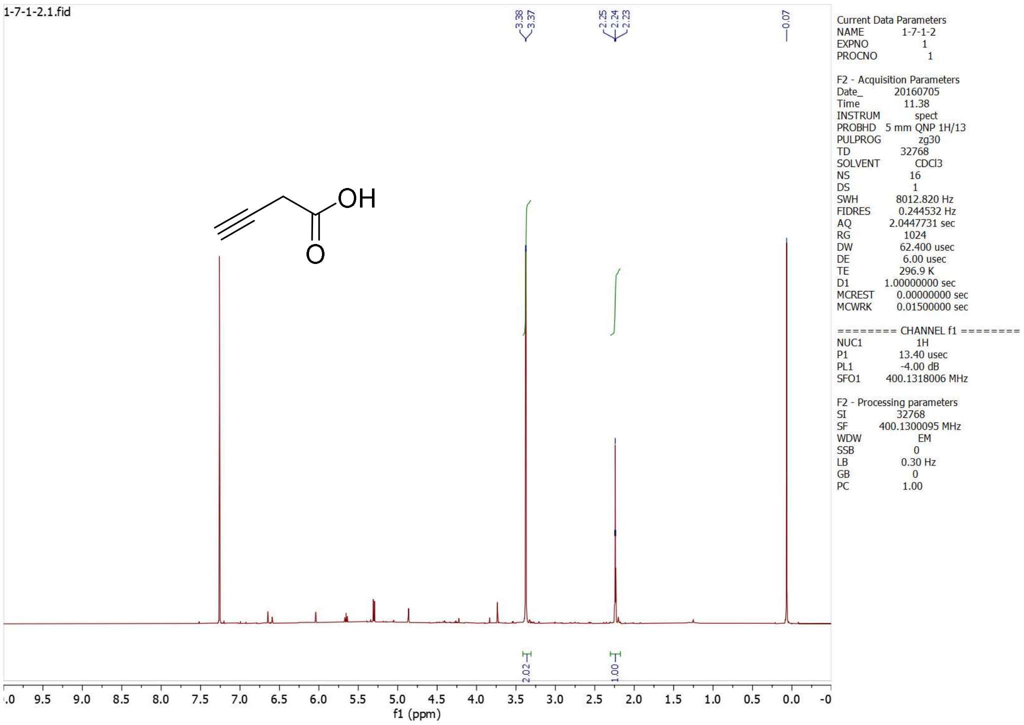

**1.2*R***

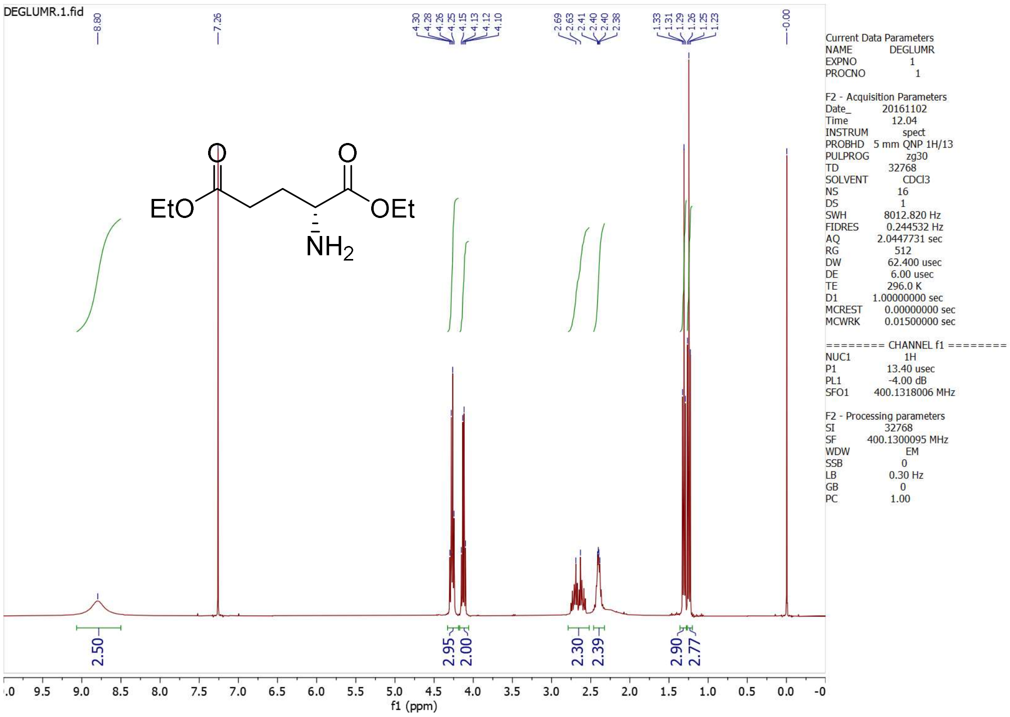

**1.2*S***

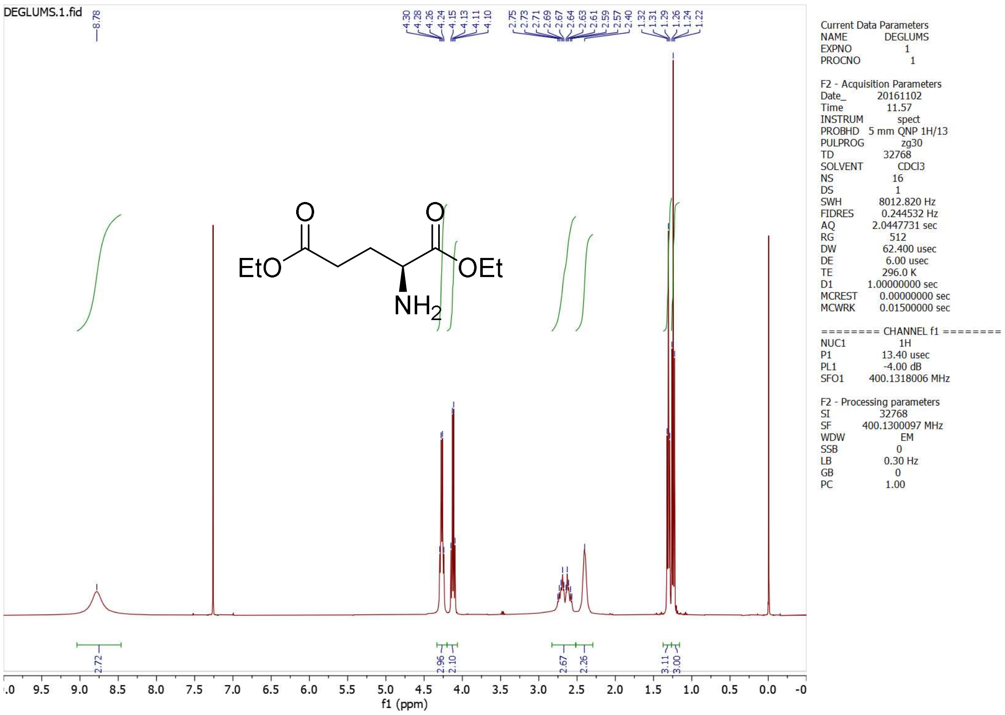

**1.2*rac***

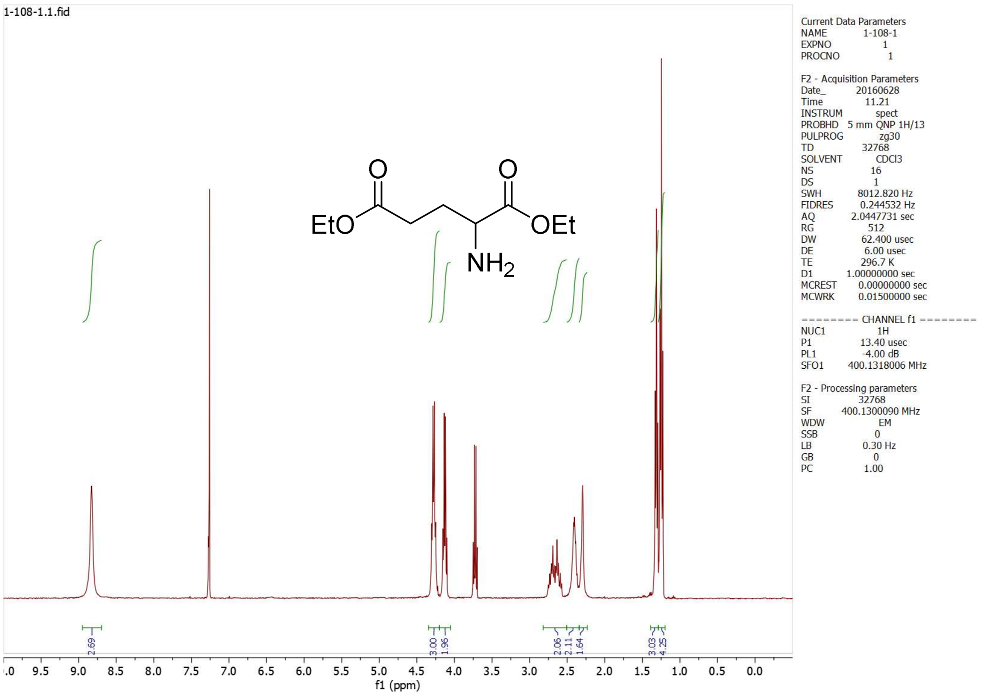
1-108-1.1.fid
EtO
O
O
OEt
NH2
Current Data Parameters
NAME 1-108-1
EXPNO 1
PROCNO 1
F2 - Acquisition Parameters
Date_ 20160628
Time 11.21
INSTRUM spect
PROBHD 5 mm QNP 1H/13
PULPROG zg30
TD 32768
SOLVENT CDCl3
NS 16
DS 1
SWH 8012.820 Hz
FIDRES 0.244532 Hz
AQ 2.0447731 sec
RG 512
DW 62.400 usec
DE 6.00 usec
TE 296.7 K
D1 1.00000000 sec
MCREST 0.00000000 sec
MCWRK 0.01500000 sec
======== CHANNEL f1 ========
NUC1 1H
P1 13.40 usec
PL1 -4.00 dB
SFO1 400.1318006 MHz
F2 - Processing parameters
SI 32768
SF 400.1300090 MHz
WDW EM
SSB 0
LB 0.30 Hz
GB 0
PC 1.00
2.69
3.00
1.96
2.06
2.11
1.64
3.03
4.25
f1 (ppm)

**1*S***

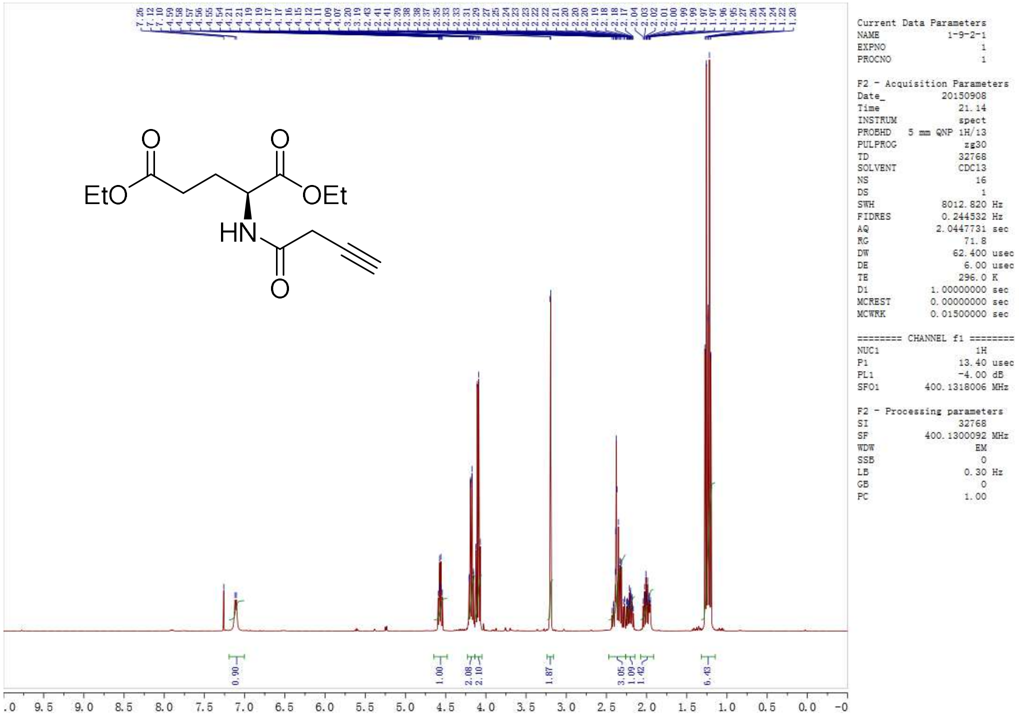
EtO
O
O
OEt
HN
O
Current Data Parameters
NAME 1-9-2-1
EXPNO 1
PROCNO 1
F2 - Acquisition Parameters
Date_ 20150908
Time 21.14
INSTRUM spect
PROBHD 5 mm QNP 1H/13
PULPROG zg30
TD 32768
SOLVENT CDCl3
NS 16
DS 1
SWH 8012.820 Hz
FIDRES 0.244532 Hz
AQ 2.0447731 sec
RG 71.8
DW 62.400 usec
DE 6.00 usec
TE 296.0 K
D1 1.00000000 sec
MCREST 0.00000000 sec
MCWRK 0.01500000 sec
======== CHANNEL f1 ========
NUC1 1H
P1 13.40 usec
PL1 -4.00 dB
SFO1 400.1318006 MHz
F2 - Processing parameters
SI 32768
SF 400.1300092 MHz
WDW EM
SSB 0
LB 0.30 Hz
GB 0
PC 1.00
0.90
1.00
2.08
2.10
1.87
3.05
1.09
1.42
6.43

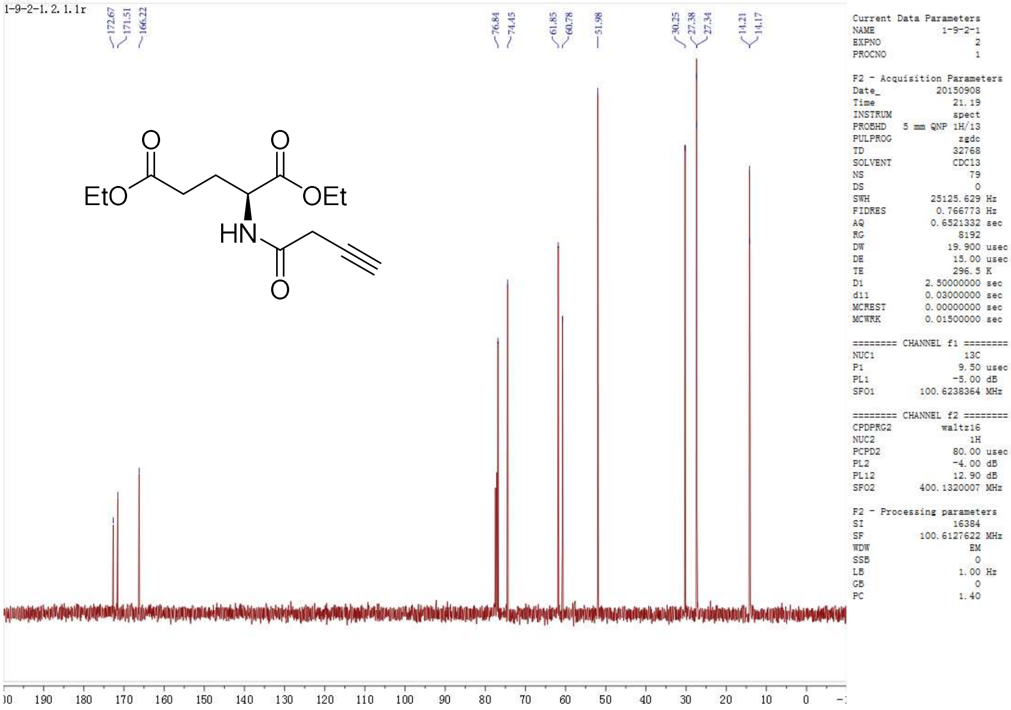

1-9-2-1.2.1.1r
172.67
171.51
166.22
76.84
74.45
61.85
60.78
51.98
30.25
27.38
27.34
14.21
14.17
EtO
OEt
HN
O
Current Data Parameters
NAME 1-9-2-1
EXPNO 2
PROCNO 1
F2 - Acquisition Parameters
Date_ 20150908
Time 21.19
INSTRUM spect
PROBHD 5 mm QNP 1H/13
PULPROG zgdc
TD 32768
SOLVENT CDCl3
NS 79
DS 0
SWH 25125.629 Hz
FIDRES 0.766773 Hz
AQ 0.6521332 sec
RG 8192
DW 19.900 usec
DE 15.00 usec
TE 296.5 K
D1 2.50000000 sec
d11 0.03000000 sec
MCREST 0.00000000 sec
MCWRK 0.01500000 sec
======== CHANNEL f1 ========
NUC1 13C
P1 9.50 usec
PL1 -5.00 dB
SFO1 100.6238364 MHz
======== CHANNEL f2 ========
CPDPRG2 waltz16
NUC2 1H
PCPD2 80.00 usec
PL2 -4.00 dB
PL12 12.90 dB
SFO2 400.1320007 MHz
F2 - Processing parameters
SI 16384
SF 100.6127622 MHz
WDW EM
SSB 0
LB 1.00 Hz
GB 0
PC 1.40
190 180 170 160 150 140 130 120 110 100 90 80 70 60 50 40 30 20 10 0

**1*R***

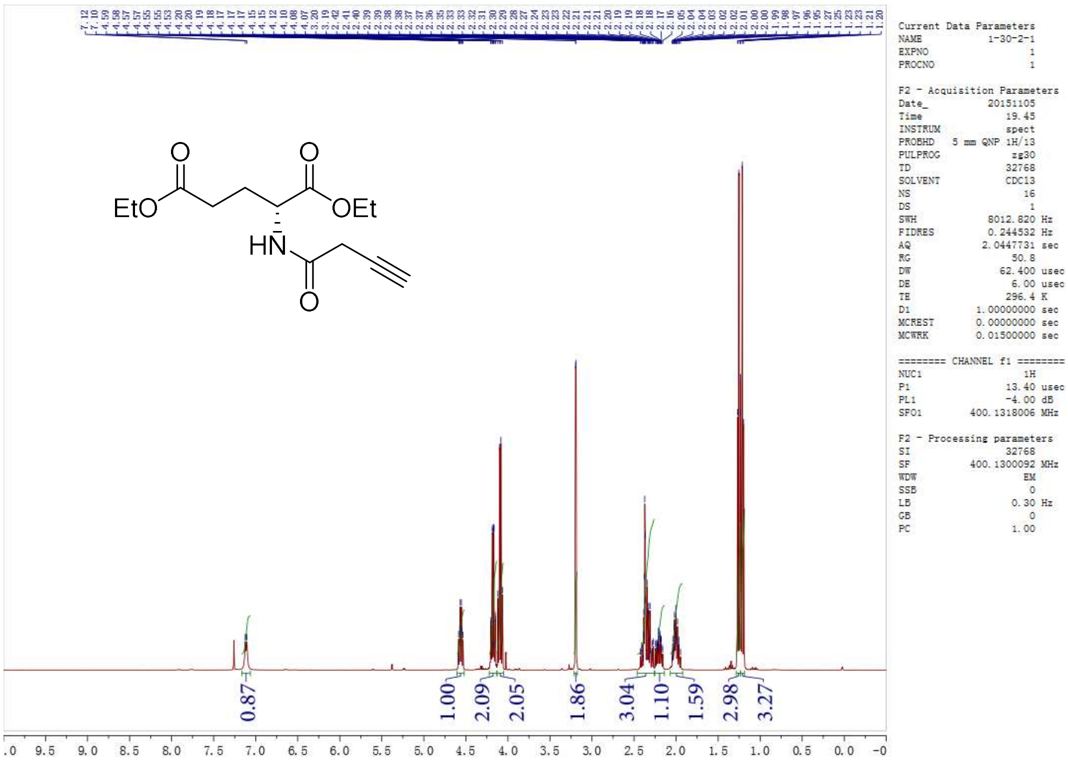

EtO
OEt
HN
O
Current Data Parameters
NAME 1-30-2-1
EXPNO 1
PROCNO 1
F2 - Acquisition Parameters
Date_ 20151105
Time 19.45
INSTRUM spect
PROBHD 5 mm QNP 1H/13
PULPROG zg30
TD 32768
SOLVENT CDCl3
NS 16
DS 1
SWH 8012.820 Hz
FIDRES 0.244532 Hz
AQ 2.0447731 sec
RG 50.8
DW 62.400 usec
DE 6.00 usec
TE 296.4 K
D1 1.00000000 sec
MCREST 0.00000000 sec
MCWRK 0.01500000 sec
======== CHANNEL f1 ========
NUC1 1H
P1 13.40 usec
PL1 -4.00 dB
SFO1 400.1318006 MHz
F2 - Processing parameters
SI 32768
SF 400.1300092 MHz
WDW EM
SSB 0
LB 0.30 Hz
GB 0
PC 1.00
0.87
1.00
2.09
2.05
1.86
3.04
1.10
1.59
2.98
3.27
9.5 9.0 8.5 8.0 7.5 7.0 6.5 6.0 5.5 5.0 4.5 4.0 3.5 3.0 2.5 2.0 1.5 1.0 0.5 0.0

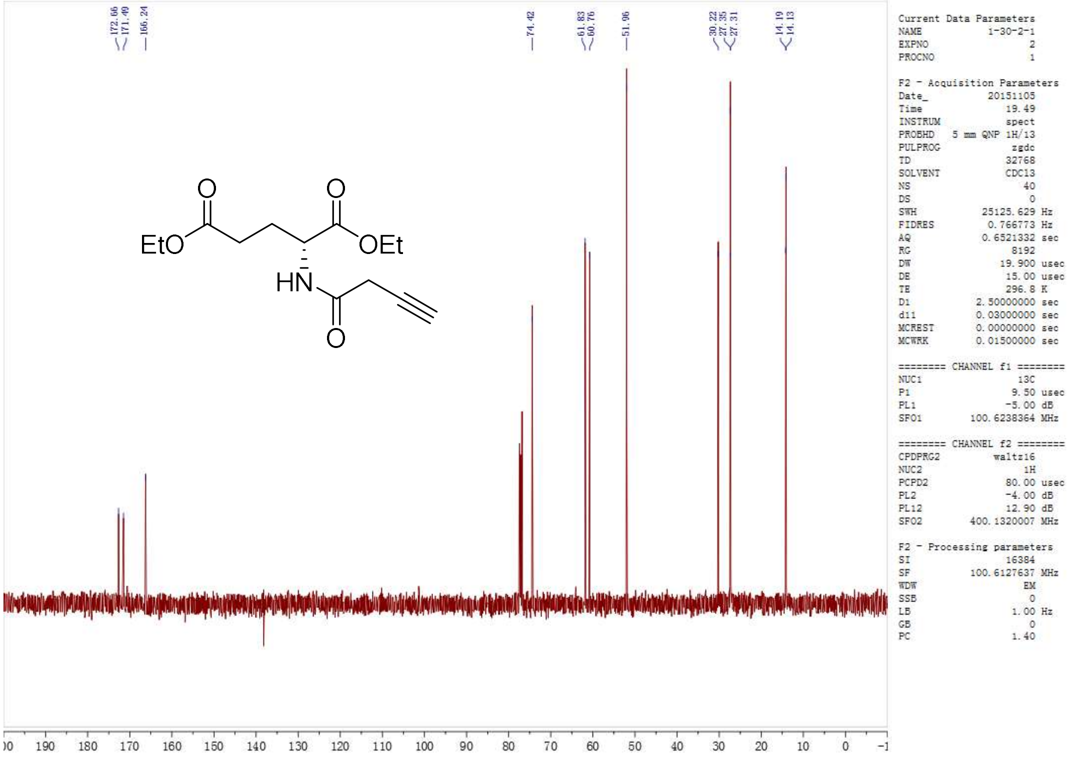
172.66
171.49
166.24
74.42
61.83
60.76
51.96
30.22
27.35
27.31
14.19
14.13
EtO
OEt
HN
O
Current Data Parameters
NAME 1-30-2-1
EXPNO 2
PROCNO 1
F2 - Acquisition Parameters
Date_ 20151105
Time 19.49
INSTRUM spect
PROBHD 5 mm QNP 1H/13
PULPROG zgdc
TD 32768
SOLVENT CDCl3
NS 40
DS 0
SWH 25125.629 Hz
FIDRES 0.766773 Hz
AQ 0.6521332 sec
RG 8192
DW 19.900 usec
DE 15.00 usec
TE 296.8 K
D1 2.50000000 sec
d11 0.03000000 sec
MCREST 0.00000000 sec
MCWRK 0.01500000 sec
======== CHANNEL f1 ========
NUC1 13C
P1 9.50 usec
PL1 -5.00 dB
SFO1 100.6238364 MHz
======== CHANNEL f2 ========
CPDPRG2 waltz16
NUC2 1H
PCPD2 80.00 usec
PL2 -4.00 dB
PL12 12.90 dB
SFO2 400.1320007 MHz
F2 - Processing parameters
SI 16384
SF 100.6127637 MHz
WDW EM
SSB 0
LB 1.00 Hz
GB 0
PC 1.40
190 180 170 160 150 140 130 120 110 100 90 80 70 60 50 40 30 20 10 0

**1rac**

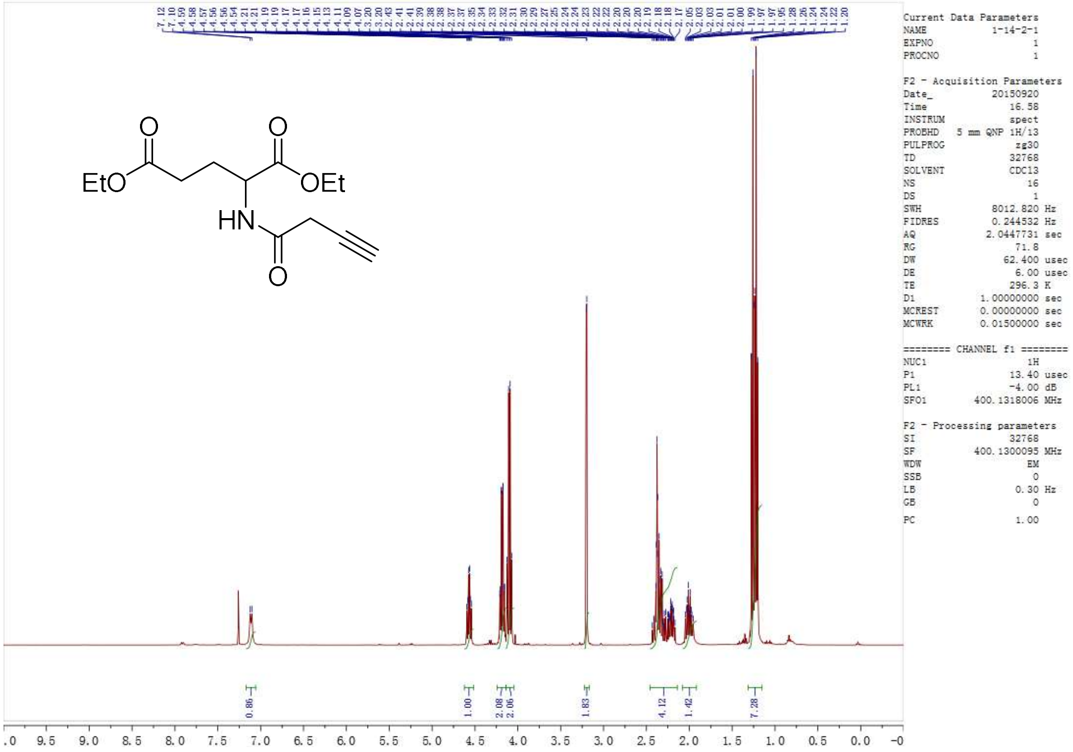
7.12 7.10 4.59 4.58 4.57 4.56 4.56 4.54 4.21 4.21 4.19 4.19 4.17 4.17 4.16 4.15 4.13 4.11 4.09 4.07 3.20 3.20 2.43 2.41 2.41 2.39 2.38 2.38 2.37 2.37 2.35 2.34 2.33 2.32 2.31 2.30 2.29 2.27 2.25 2.24 2.24 2.23 2.22 2.22 2.20 2.20 2.20 2.19 2.18 2.18 2.17 2.05 2.03 2.03 2.01 2.01 2.00 1.99 1.97 1.97 1.95 1.28 1.26 1.24 1.24 1.22 1.20
EtO
OEt
HN
O
Current Data Parameters
NAME 1-14-2-1
EXPNO 1
PROCNO 1
F2 - Acquisition Parameters
Date_ 20150920
Time 16.58
INSTRUM spect
PROBHD 5 mm QNP 1H/13
PULPROG zg30
TD 32768
SOLVENT CDCl3
NS 16
DS 1
SWH 8012.820 Hz
FIDRES 0.244532 Hz
AQ 2.0447731 sec
RG 71.8
DW 62.400 usec
DE 6.00 usec
TE 296.3 K
D1 1.00000000 sec
MCREST 0.00000000 sec
MCWRK 0.01500000 sec
======== CHANNEL f1 ========
NUC1 1H
P1 13.40 usec
PL1 -4.00 dB
SFO1 400.1318006 MHz
F2 - Processing parameters
SI 32768
SF 400.1300095 MHz
WDW EM
SSB 0
LB 0.30 Hz
GB 0
PC 1.00
0.86
1.00
2.08
2.06
1.83
4.12
1.42
7.28
9.5 9.0 8.5 8.0 7.5 7.0 6.5 6.0 5.5 5.0 4.5 4.0 3.5 3.0 2.5 2.0 1.5 1.0 0.5 0.0

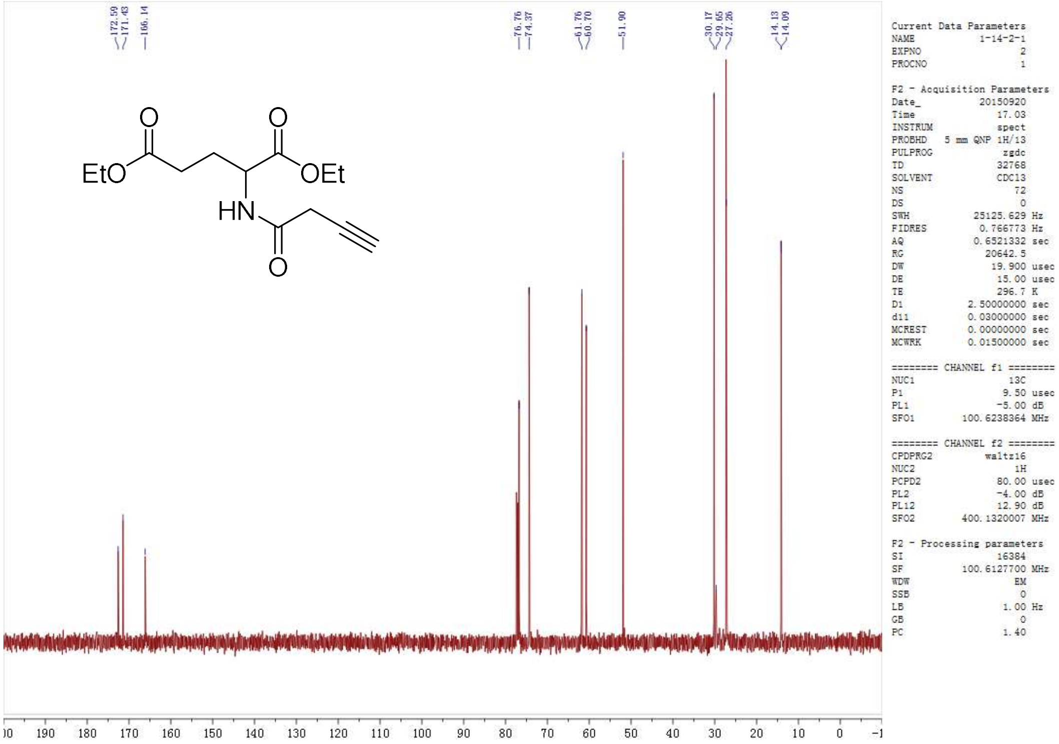

172.59
171.43
166.14
76.76
74.37
61.76
60.70
51.90
30.17
29.65
27.26
14.13
14.09
EtO
OEt
HN
O
Current Data Parameters
NAME 1-14-2-1
EXPNO 2
PROCNO 1
F2 - Acquisition Parameters
Date_ 20150920
Time 17.03
INSTRUM spect
PROBHD 5 mm QNP 1H/13
PULPROG zgdc
TD 32768
SOLVENT CDCl3
NS 72
DS 0
SWH 25125.629 Hz
FIDRES 0.766773 Hz
AQ 0.6521332 sec
RG 20642.5
DW 19.900 usec
DE 15.00 usec
TE 296.7 K
D1 2.50000000 sec
d11 0.03000000 sec
MCREST 0.00000000 sec
MCWRK 0.01500000 sec
======== CHANNEL f1 ========
NUC1 13C
P1 9.50 usec
PL1 -5.00 dB
SFO1 100.6238364 MHz
======== CHANNEL f2 ========
CPDPRG2 waltz16
NUC2 1H
PCPD2 80.00 usec
PL2 -4.00 dB
PL12 12.90 dB
SFO2 400.1320007 MHz
F2 - Processing parameters
SI 16384
SF 100.6127700 MHz
WDW EM
SSB 0
LB 1.00 Hz
GB 0
PC 1.40
190 180 170 160 150 140 130 120 110 100 90 80 70 60 50 40 30 20 10 0

**2*S***

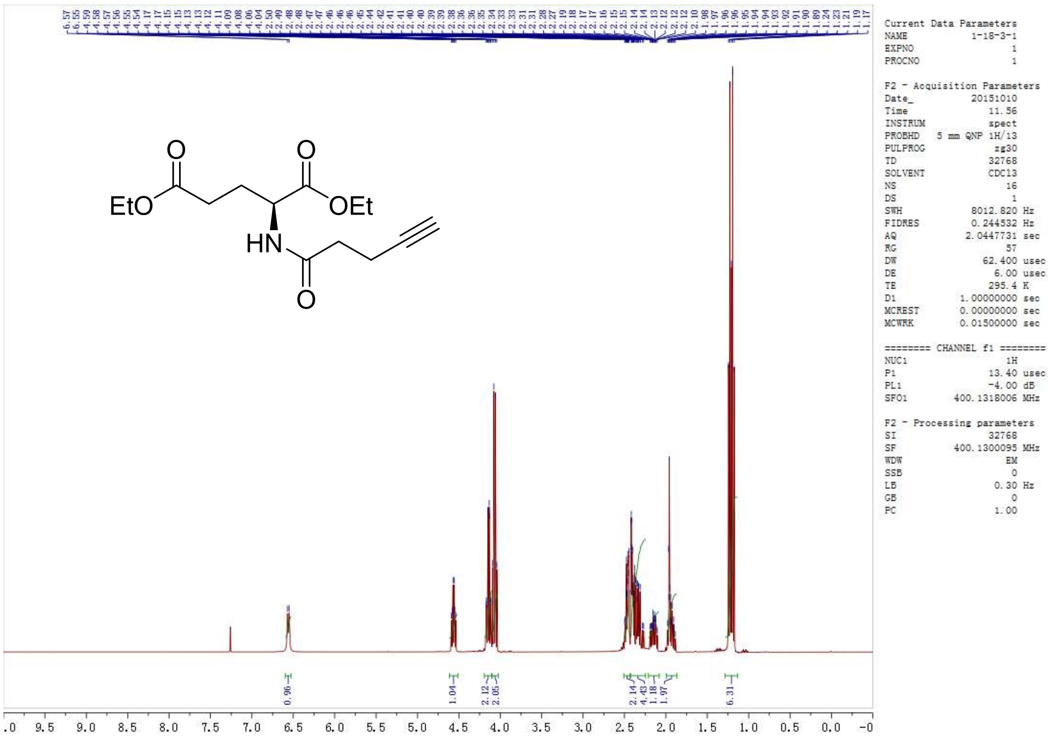

EtO
OEt
HN
O
Current Data Parameters
NAME 1-18-3-1
EXPNO 1
PROCNO 1
F2 - Acquisition Parameters
Date_ 20151010
Time 11.56
INSTRUM spect
PROBHD 5 mm QNP 1H/13
PULPROG zg30
TD 32768
SOLVENT CDCl3
NS 16
DS 1
SWH 8012.820 Hz
FIDRES 0.244532 Hz
AQ 2.0447731 sec
RG 57
DW 62.400 usec
DE 6.00 usec
TE 295.4 K
D1 1.00000000 sec
MCREST 0.00000000 sec
MCWRK 0.01500000 sec
======== CHANNEL f1 ========
NUC1 1H
P1 13.40 usec
PL1 -4.00 dB
SFO1 400.1318006 MHz
F2 - Processing parameters
SI 32768
SF 400.1300095 MHz
WDW EM
SSB 0
LB 0.30 Hz
GB 0
PC 1.00
0.96
1.04
2.12
2.05
2.14
4.43
1.18
1.97
6.31
9.5 9.0 8.5 8.0 7.5 7.0 6.5 6.0 5.5 5.0 4.5 4.0 3.5 3.0 2.5 2.0 1.5 1.0 0.5 0.0

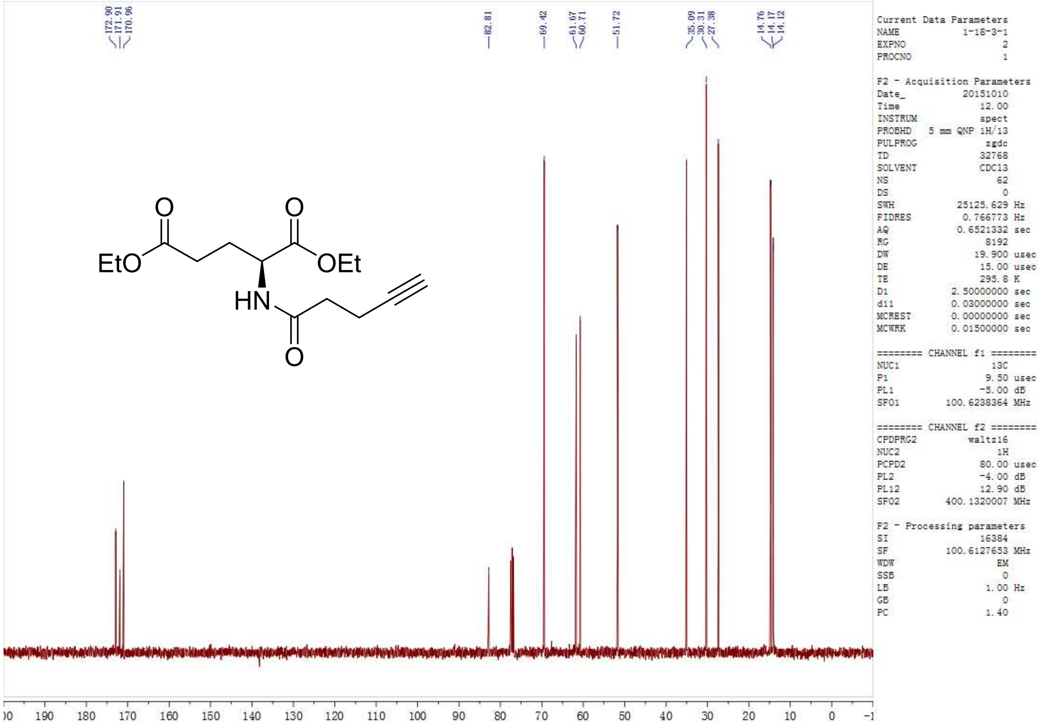

**2*R***

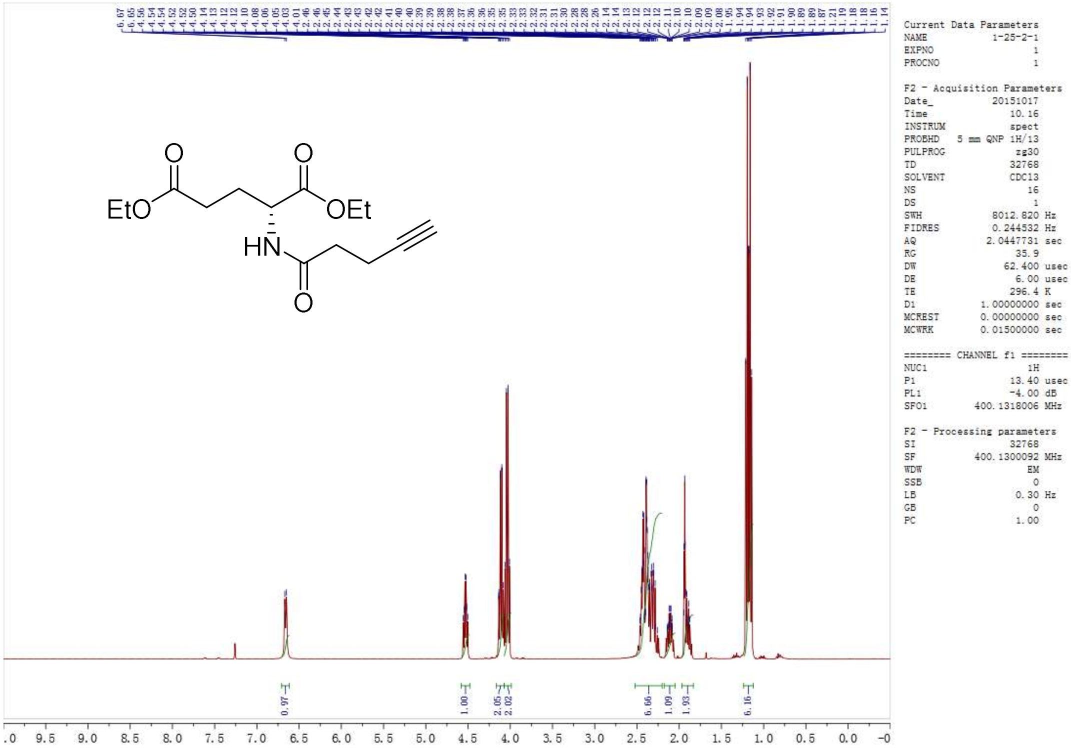

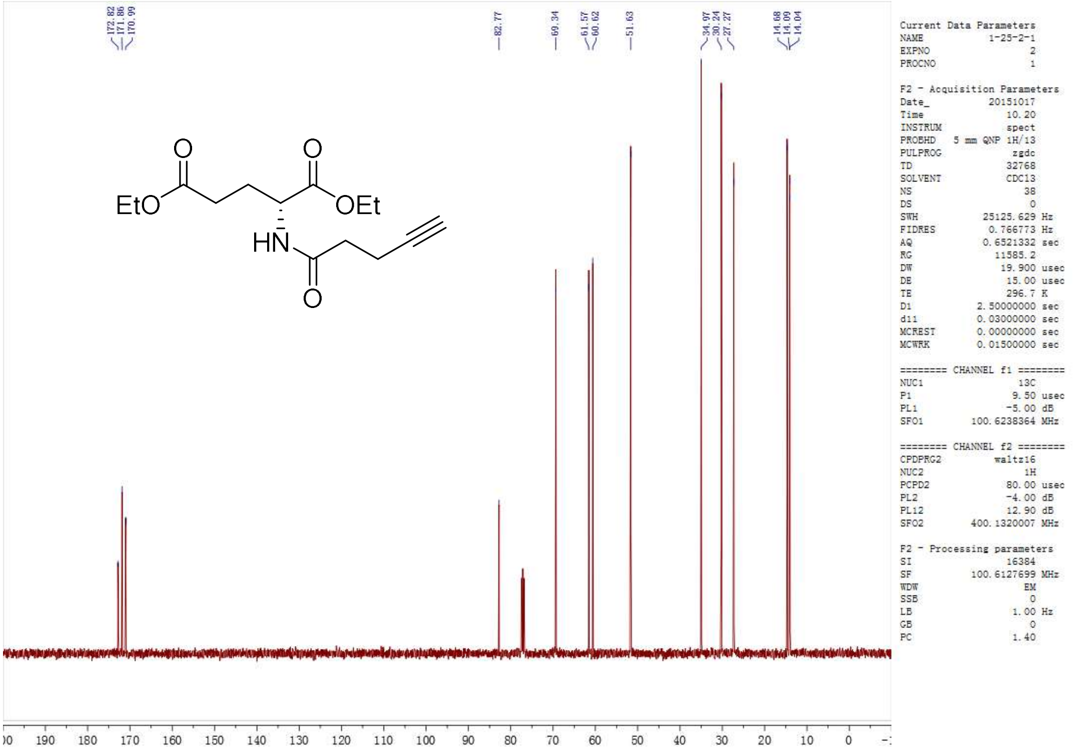
172.82
171.86
170.99
82.77
69.34
61.57
60.62
51.63
34.97
30.24
27.27
14.68
14.09
14.04
EtO
OEt
HN
Current Data Parameters
NAME 1-25-2-1
EXPNO 2
PROCNO 1
F2 - Acquisition Parameters
Date_ 20151017
Time 10.20
INSTRUM spect
PROBHD 5 mm QNP 1H/13
PULPROG zgdc
TD 32768
SOLVENT CDCl3
NS 38
DS 0
SWH 25125.629 Hz
FIDRES 0.766773 Hz
AQ 0.6521332 sec
RG 11585.2
DW 19.900 usec
DE 15.00 usec
TE 296.7 K
D1 2.50000000 sec
d11 0.03000000 sec
MCREST 0.00000000 sec
MCWRK 0.01500000 sec
======== CHANNEL f1 ========
NUC1 13C
P1 9.50 usec
PL1 -5.00 dB
SFO1 100.6238364 MHz
======== CHANNEL f2 ========
CPDPRG2 waltz16
NUC2 1H
PCPD2 80.00 usec
PL2 -4.00 dB
PL12 12.90 dB
SFO2 400.1320007 MHz
F2 - Processing parameters
SI 16384
SF 100.6127699 MHz
WDW EM
SSB 0
LB 1.00 Hz
GB 0
PC 1.40
190 180 170 160 150 140 130 120 110 100 90 80 70 60 50 40 30 20 10 0

**2rac**

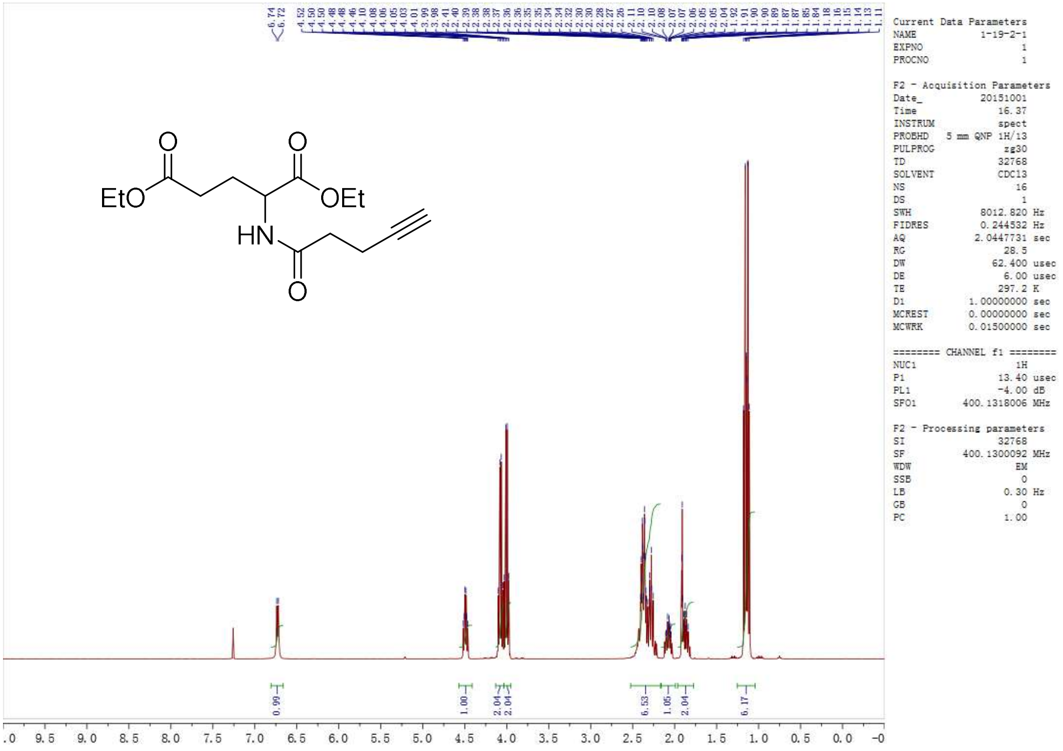
EtO
OEt
HN
Current Data Parameters
NAME 1-19-2-1
EXPNO 1
PROCNO 1
F2 - Acquisition Parameters
Date_ 20151001
Time 16.37
INSTRUM spect
PROBHD 5 mm QNP 1H/13
PULPROG zg30
TD 32768
SOLVENT CDCl3
NS 16
DS 1
SWH 8012.820 Hz
FIDRES 0.244532 Hz
AQ 2.0447731 sec
RG 28.5
DW 62.400 usec
DE 6.00 usec
TE 297.2 K
D1 1.00000000 sec
MCREST 0.00000000 sec
MCWRK 0.01500000 sec
======== CHANNEL f1 ========
NUC1 1H
P1 13.40 usec
PL1 -4.00 dB
SFO1 400.1318006 MHz
F2 - Processing parameters
SI 32768
SF 400.1300092 MHz
WDW EM
SSB 0
LB 0.30 Hz
GB 0
PC 1.00
0.99
1.00
2.04
2.04
6.53
1.05
2.04
6.17
9.5 9.0 8.5 8.0 7.5 7.0 6.5 6.0 5.5 5.0 4.5 4.0 3.5 3.0 2.5 2.0 1.5 1.0 0.5 0.0

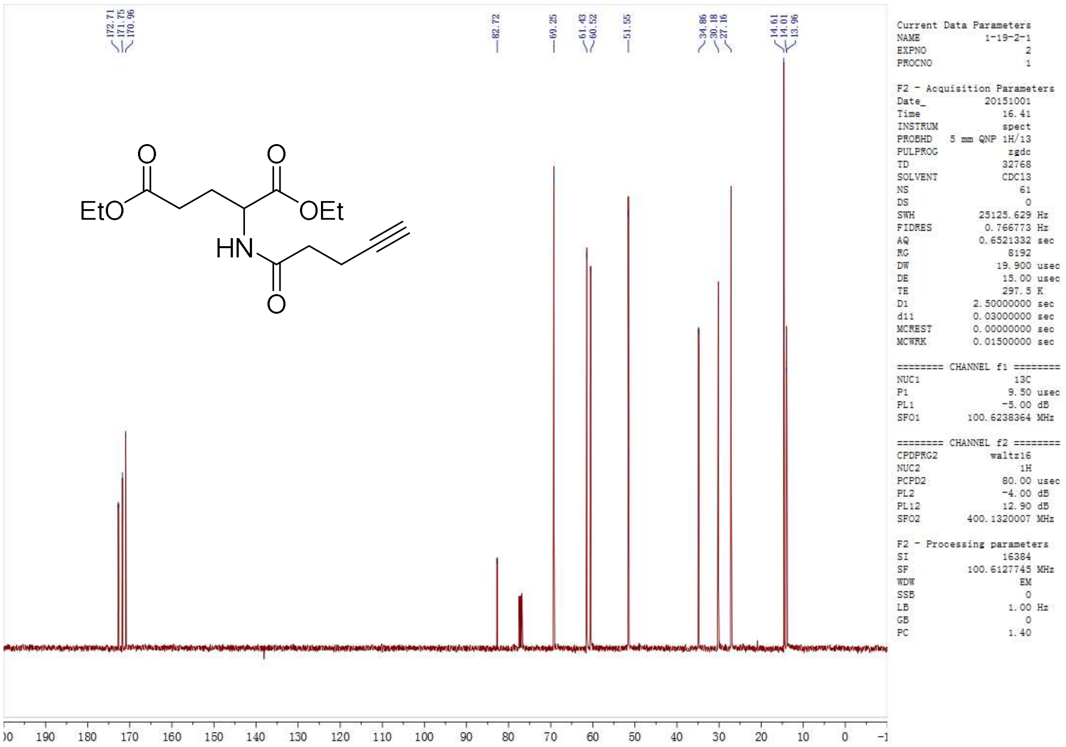
172.71
171.75
170.96
82.72
69.25
61.43
60.52
51.55
34.86
30.18
27.16
14.61
14.01
13.96
EtO
OEt
HN
O
Current Data Parameters
NAME 1-19-2-1
EXPNO 2
PROCNO 1
F2 - Acquisition Parameters
Date_ 20151001
Time 16.41
INSTRUM spect
PROBHD 5 mm QNP 1H/13
PULPROG zgdc
TD 32768
SOLVENT CDCl3
NS 61
DS 0
SWH 25125.629 Hz
FIDRES 0.766773 Hz
AQ 0.6521332 sec
RG 8192
DW 19.900 usec
DE 15.00 usec
TE 297.5 K
D1 2.50000000 sec
d11 0.03000000 sec
MCREST 0.00000000 sec
MCWRK 0.01500000 sec
======== CHANNEL f1 ========
NUC1 13C
P1 9.50 usec
PL1 -5.00 dB
SFO1 100.6238364 MHz
======== CHANNEL f2 ========
CPDPRG2 waltz16
NUC2 1H
PCPD2 80.00 usec
PL2 -4.00 dB
PL12 12.90 dB
SFO2 400.1320007 MHz
F2 - Processing parameters
SI 16384
SF 100.6127745 MHz
WDW EM
SSB 0
LB 1.00 Hz
GB 0
PC 1.40
190 180 170 160 150 140 130 120 110 100 90 80 70 60 50 40 30 20 10 0

**2*R*ME**

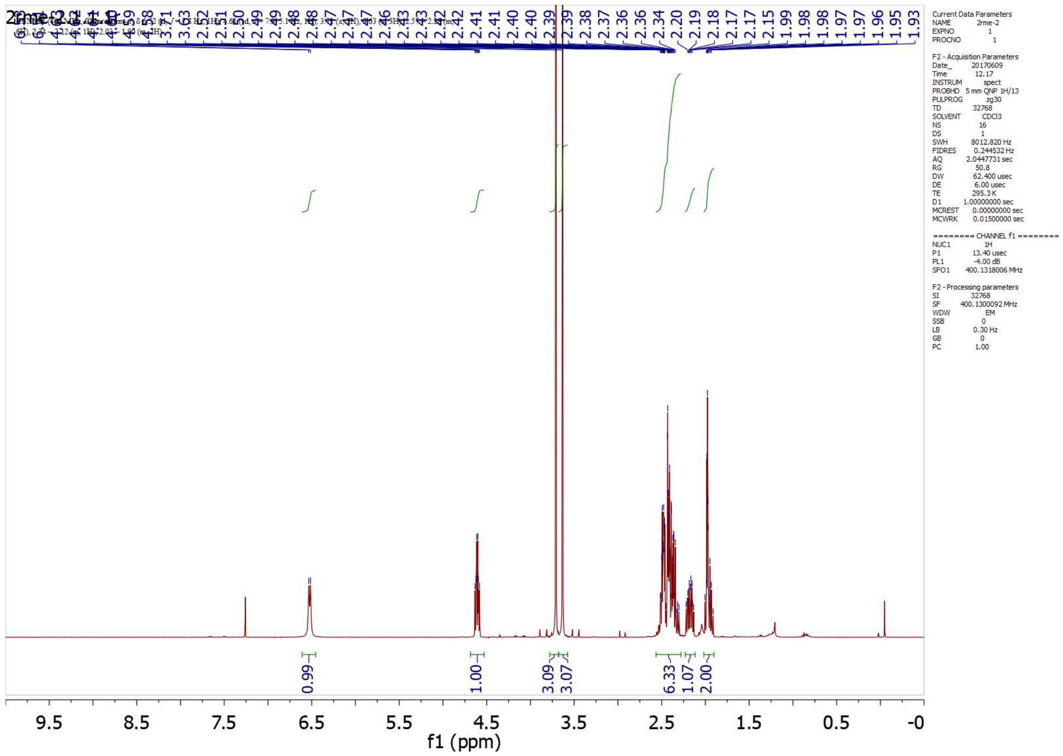
2.41
2.41
2.40
2.40
2.39
2.39
2.38
2.37
2.36
2.36
2.34
2.20
2.19
2.18
2.17
2.17
2.15
1.99
1.98
1.97
1.97
1.96
1.95
1.93
Current Data Parameters
NAME 2rme-2
EXPNO 1
PROCNO 1
F2 - Acquisition Parameters
Date_ 20170609
Time 12.17
INSTRUM spect
PROBHD 5 mm QNP 1H/13
PULPROG zg30
TD 32768
SOLVENT CDCl3
NS 16
DS 1
SWH 8012.820 Hz
FIDRES 0.244532 Hz
AQ 2.0447731 sec
RG 50.8
DW 62.400 usec
DE 6.00 usec
TE 295.3 K
D1 1.00000000 sec
MCREST 0.00000000 sec
MCWRK 0.01500000 sec
======== CHANNEL f1 ========
NUC1 1H
P1 13.40 usec
PL1 -4.00 dB
SFO1 400.1318006 MHz
F2 - Processing parameters
SI 32768
SF 400.1300092 MHz
WDW EM
SSB 0
LB 0.30 Hz
GB 0
PC 1.00
0.99
1.00
3.09
3.07
6.33
1.07
2.00
9.5
8.5
7.5
6.5
5.5
4.5
3.5
2.5
1.5
0.5
-0
f1 (ppm)

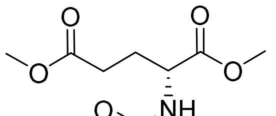
O
O
O
NH

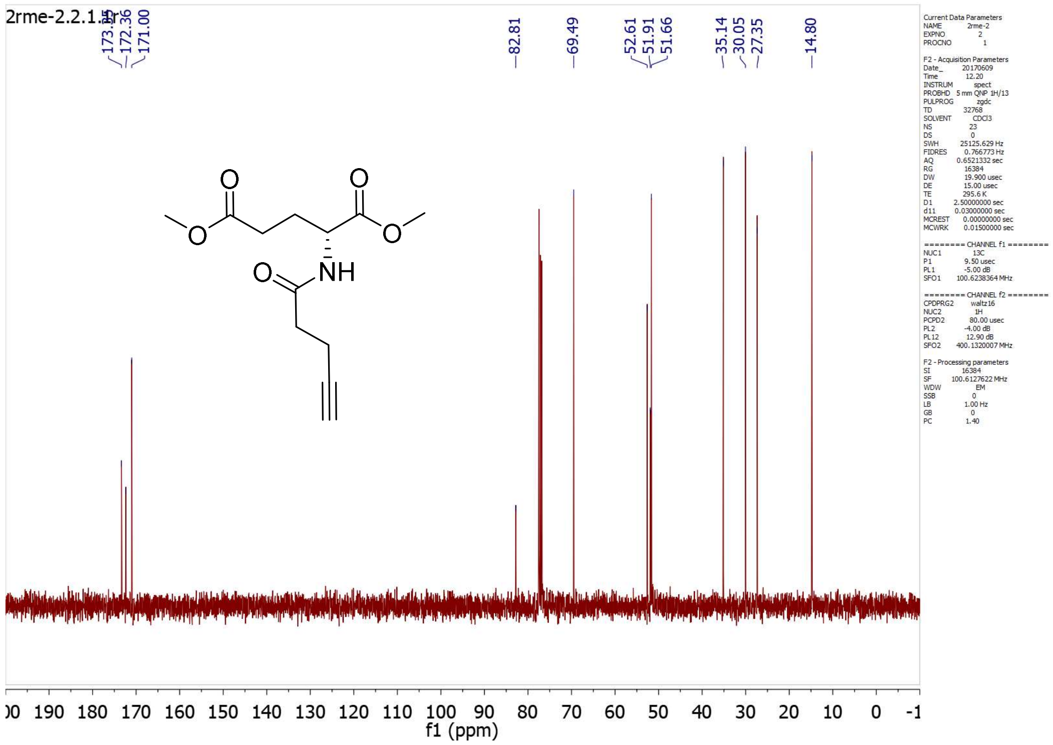
172.36
171.00
82.81
69.49
52.61
51.91
51.66
35.14
30.05
27.35
14.80
O
O
O
O
O
NH
f1 (ppm)

**2*S*ME**

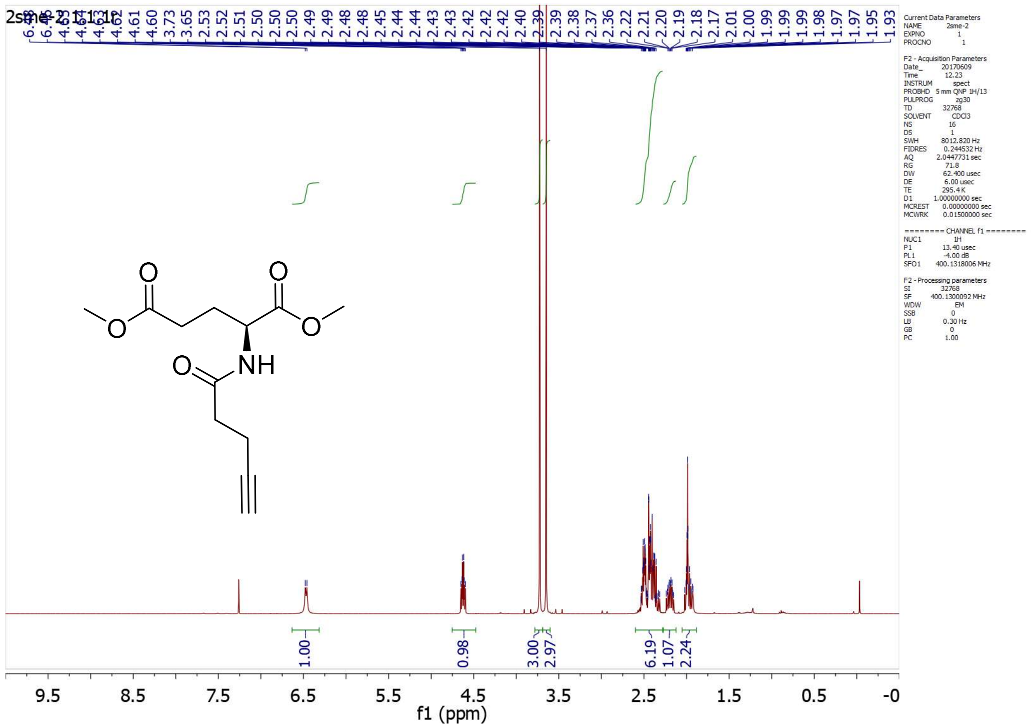
1.00
0.98
3.00
2.97
6.19
1.07
2.24
O
O
O
O
O
NH
f1 (ppm)

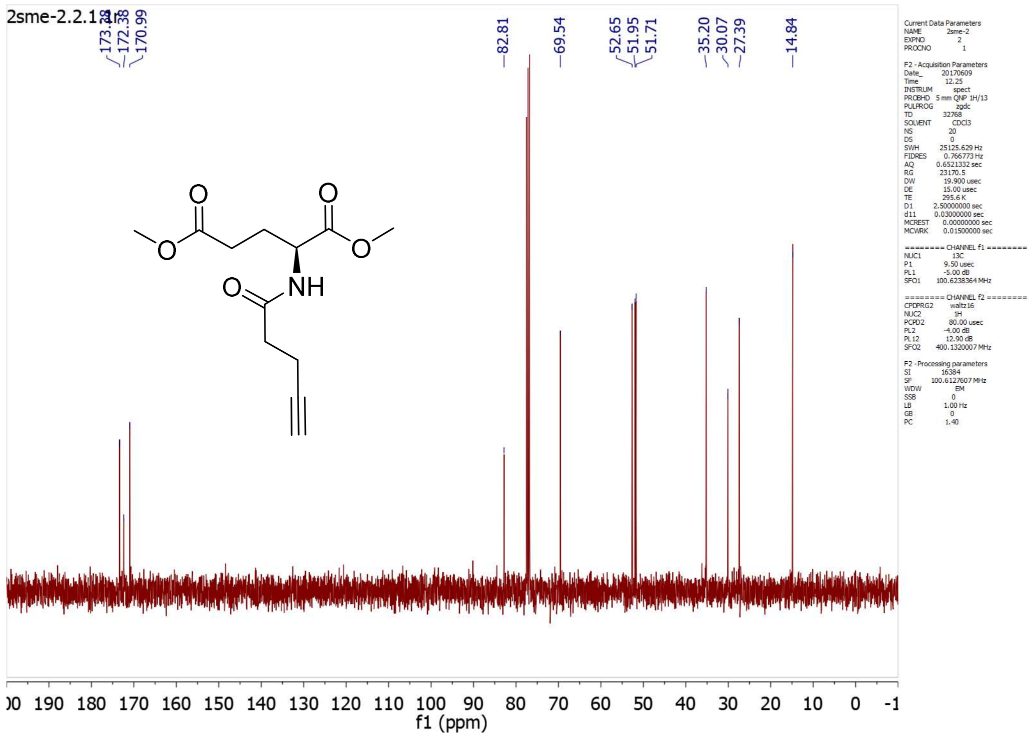

**2*rac*ME**

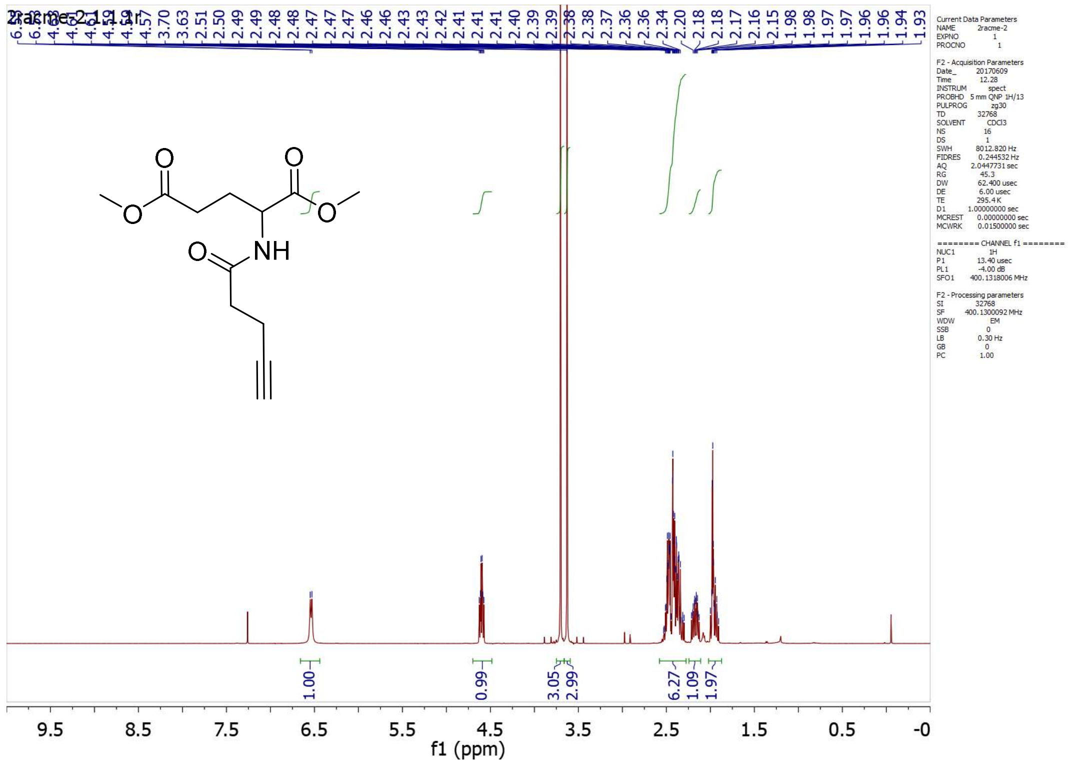

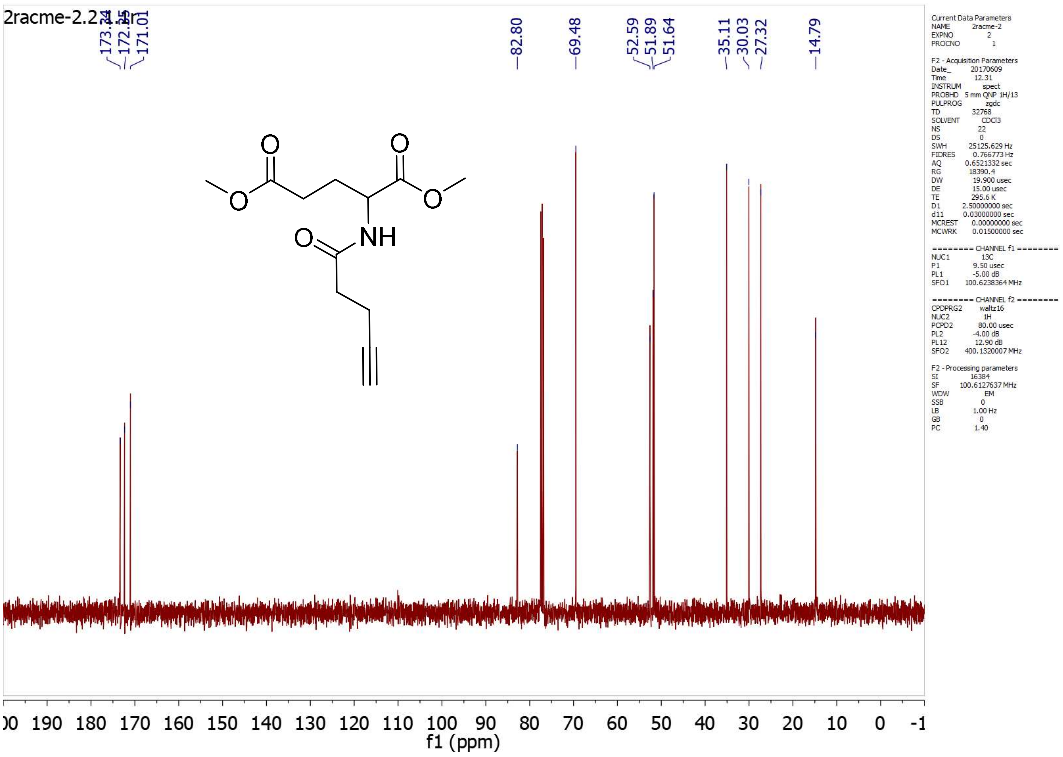
2racme-2.2.1.fid
173.24
172.25
171.01
82.80
69.48
52.59
51.89
51.64
35.11
30.03
27.32
14.79
NH
O
f1 (ppm)

**2*R*AM**

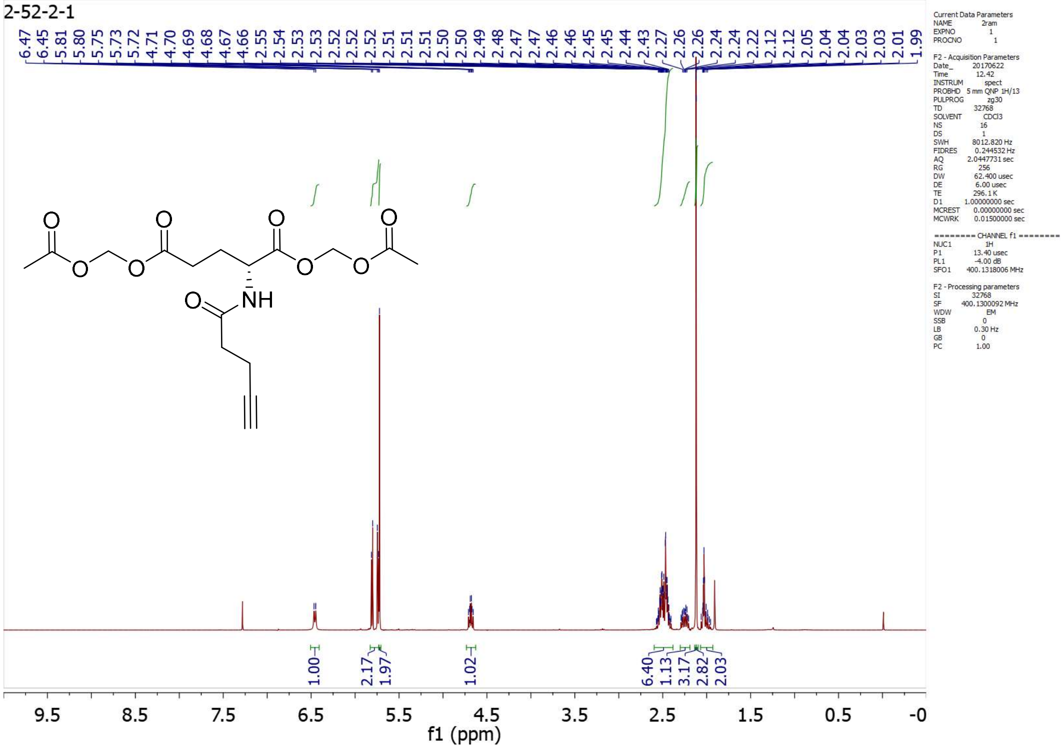
2-52-2-1
NH
O
1.00
2.17
1.97
1.02
6.40
1.13
3.17
2.82
2.03
f1 (ppm)

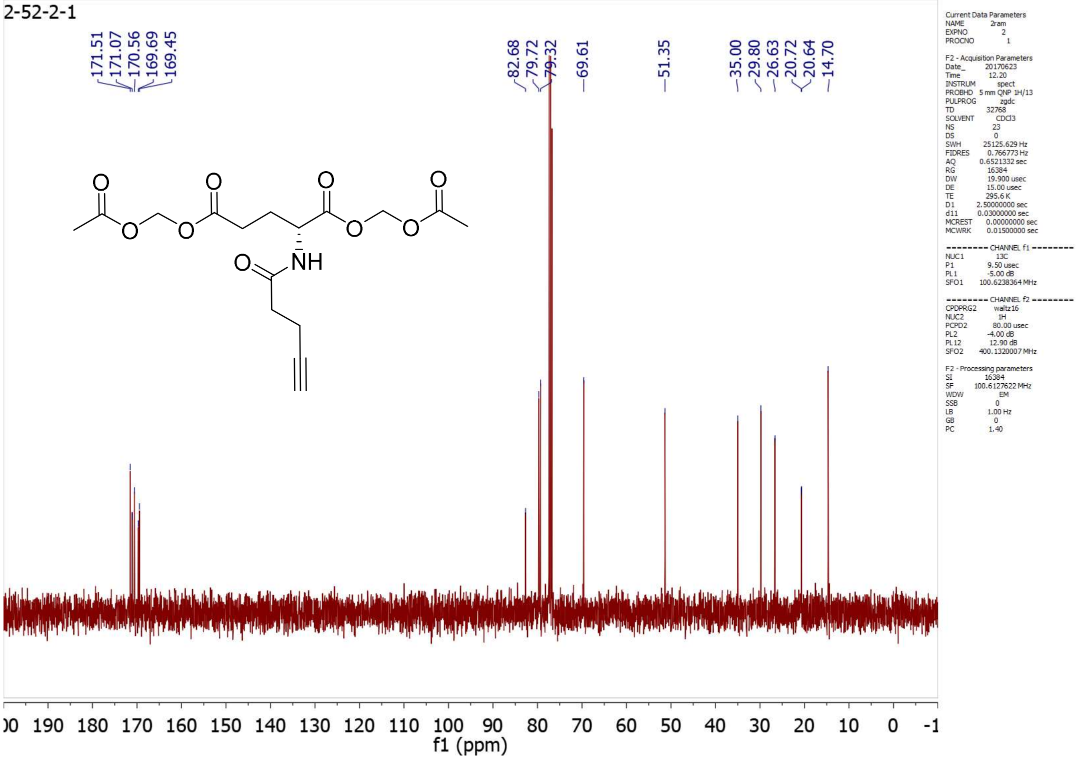
2-52-2-1
171.51
171.07
170.56
169.69
169.45
82.68
79.72
79.32
69.61
51.35
35.00
29.80
26.63
20.72
20.64
14.70
NH
f1 (ppm)

**2*S*AM**

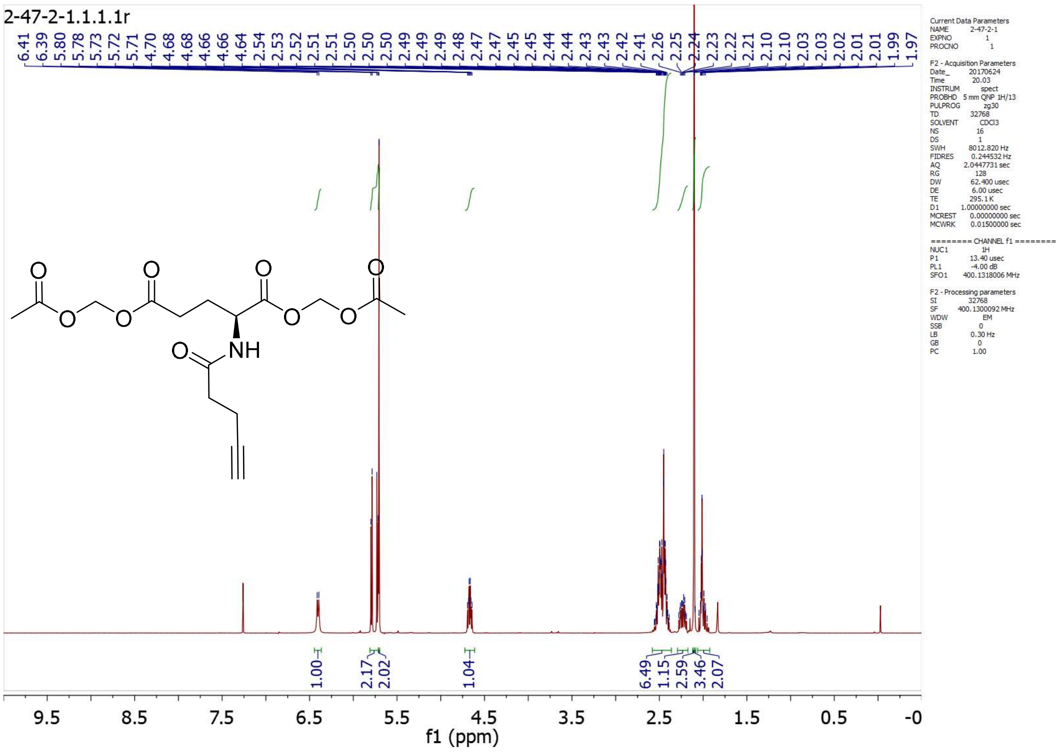
2-47-2-1.1.1.1r
6.41
6.39
5.80
5.78
5.73
5.73
5.71
4.70
4.68
4.66
4.66
4.64
2.54
2.53
2.52
2.51
2.51
2.50
2.50
2.50
2.49
2.49
2.48
2.47
2.45
2.45
2.44
2.44
2.43
2.43
2.42
2.41
2.26
2.25
2.24
2.23
2.22
2.21
2.10
2.03
2.02
2.01
1.99
1.97
NH
1.00
2.17
2.02
1.04
6.49
1.15
2.59
3.46
2.07
f1 (ppm)

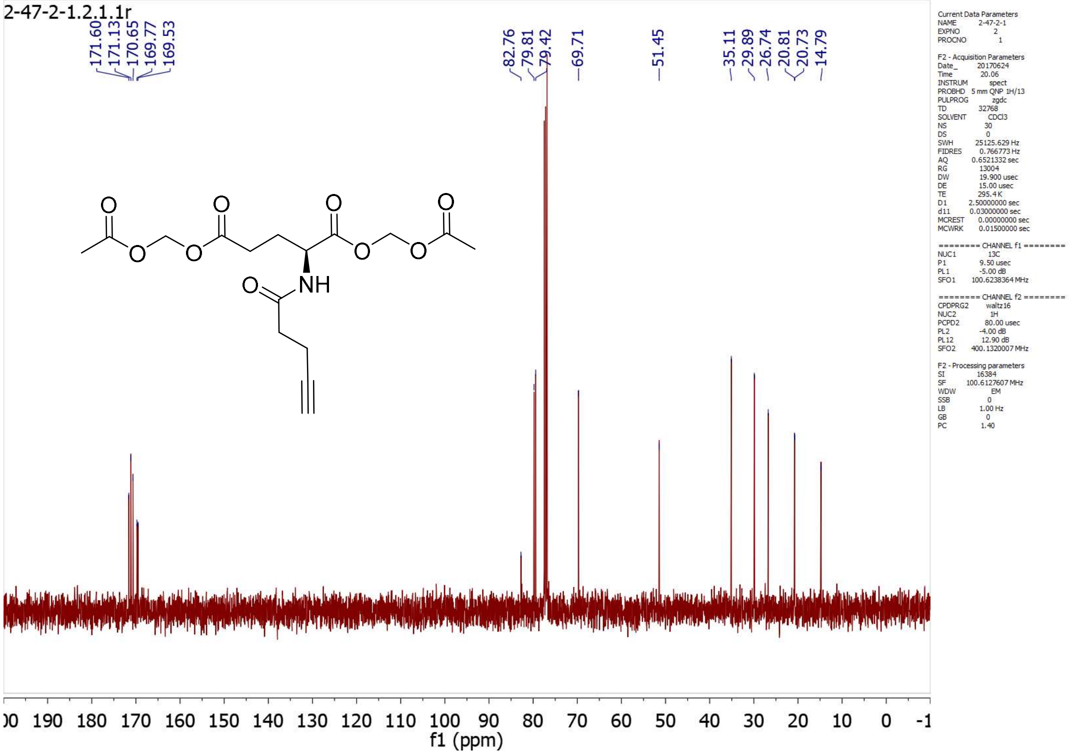
2-47-2-1.2.1.1r
171.60
171.13
170.65
169.77
169.53
82.76
79.81
79.42
69.71
51.45
35.11
29.89
26.74
20.81
20.73
14.79
NH
f1 (ppm)
Current Data Parameters
NAME 2-47-2-1
EXPNO 2
PROCNO 1
F2 - Acquisition Parameters
Date_ 20170624
Time 20.06
INSTRUM spect
PROBHD 5 mm QNP 1H/13
PULPROG zgdc
TD 32768
SOLVENT CDCl3
NS 30
DS 0
SWH 25125.629 Hz
FIDRES 0.766773 Hz
AQ 0.6521332 sec
RG 13004
DW 19.900 usec
DE 15.00 usec
TE 295.4 K
D1 2.50000000 sec
d11 0.03000000 sec
MCREST 0.00000000 sec
MCWRK 0.01500000 sec
======== CHANNEL f1 ========
NUC1 13C
P1 9.50 usec
PL1 -5.00 dB
SFO1 100.6238364 MHz
======== CHANNEL f2 ========
CPDPRG2 waltz16
NUC2 1H
PCPD2 80.00 usec
PL2 -4.00 dB
PL12 12.90 dB
SFO2 400.1320007 MHz
F2 - Processing parameters
SI 16384
SF 100.6127607 MHz
WDW EM
SSB 0
LB 1.00 Hz
GB 0
PC 1.40

**2*rac*AM**

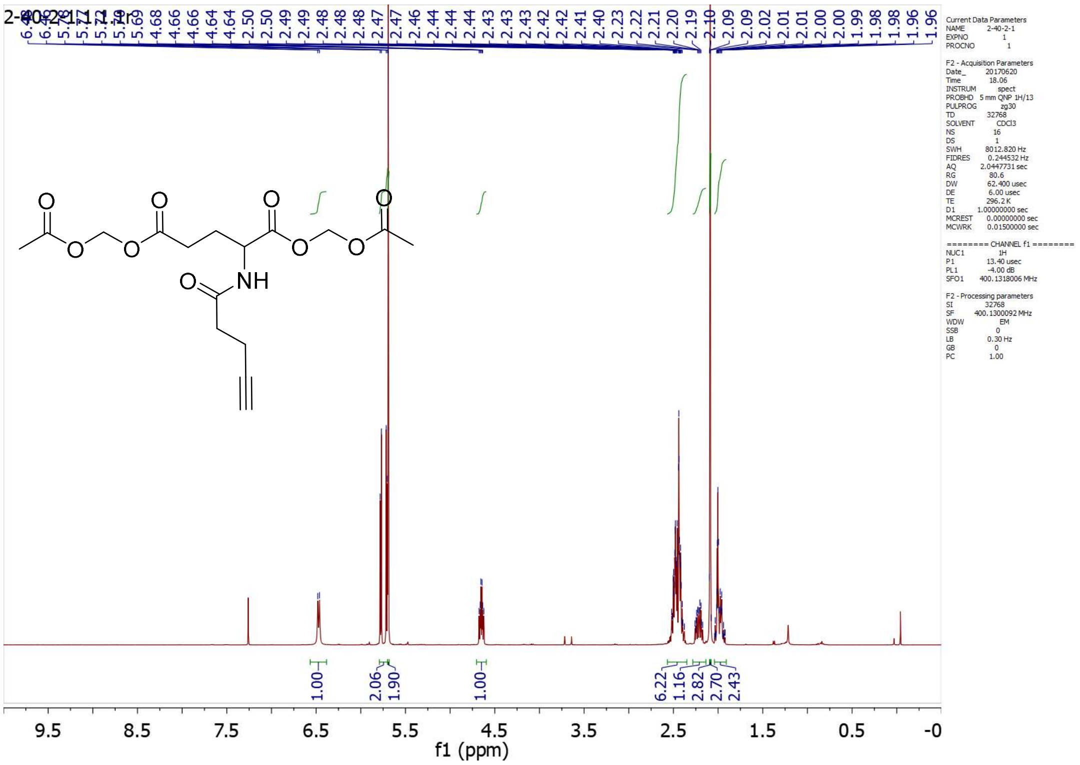
4.68
4.66
4.66
4.64
4.64
2.50
2.50
2.49
2.48
2.48
2.47
2.47
2.46
2.44
2.44
2.44
2.43
2.43
2.43
2.42
2.42
2.41
2.40
2.23
2.22
2.21
2.20
2.19
2.10
2.09
2.02
2.01
2.01
2.00
1.99
1.98
1.98
1.96
1.96
NH
1.00
2.06
1.90
1.00
6.22
1.16
2.82
2.70
2.43
f1 (ppm)
Current Data Parameters
NAME 2-40-2-1
EXPNO 1
PROCNO 1
F2 - Acquisition Parameters
Date_ 20170620
Time 18.06
INSTRUM spect
PROBHD 5 mm QNP 1H/13
PULPROG zg30
TD 32768
SOLVENT CDCl3
NS 16
DS 1
SWH 8012.820 Hz
FIDRES 0.244532 Hz
AQ 2.0447731 sec
RG 80.6
DW 62.400 usec
DE 6.00 usec
TE 296.2 K
D1 1.00000000 sec
MCREST 0.00000000 sec
MCWRK 0.01500000 sec
======== CHANNEL f1 ========
NUC1 1H
P1 13.40 usec
PL1 -4.00 dB
SFO1 400.1318006 MHz
F2 - Processing parameters
SI 32768
SF 400.1300092 MHz
WDW EM
SSB 0
LB 0.30 Hz
GB 0
PC 1.00

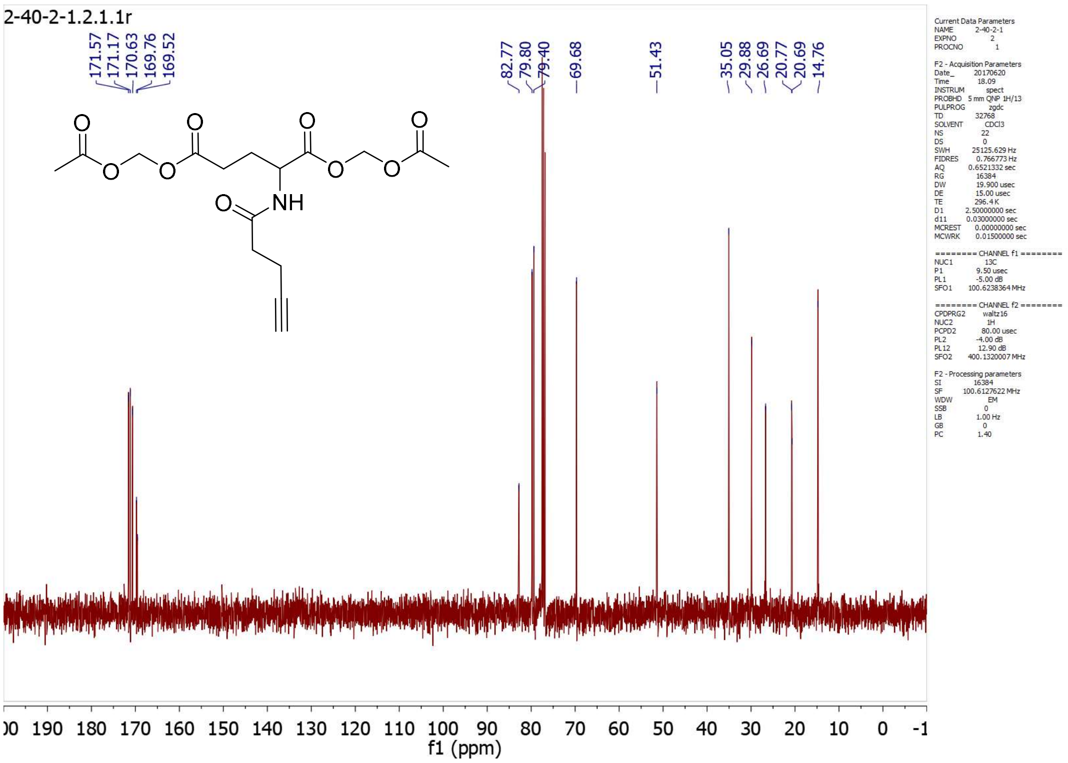
2-40-2-1.2.1.1r
171.57
171.17
170.63
169.76
169.52
82.77
79.80
79.40
69.68
51.43
35.05
29.88
26.69
20.77
20.69
14.76
f1 (ppm)

**3rac**

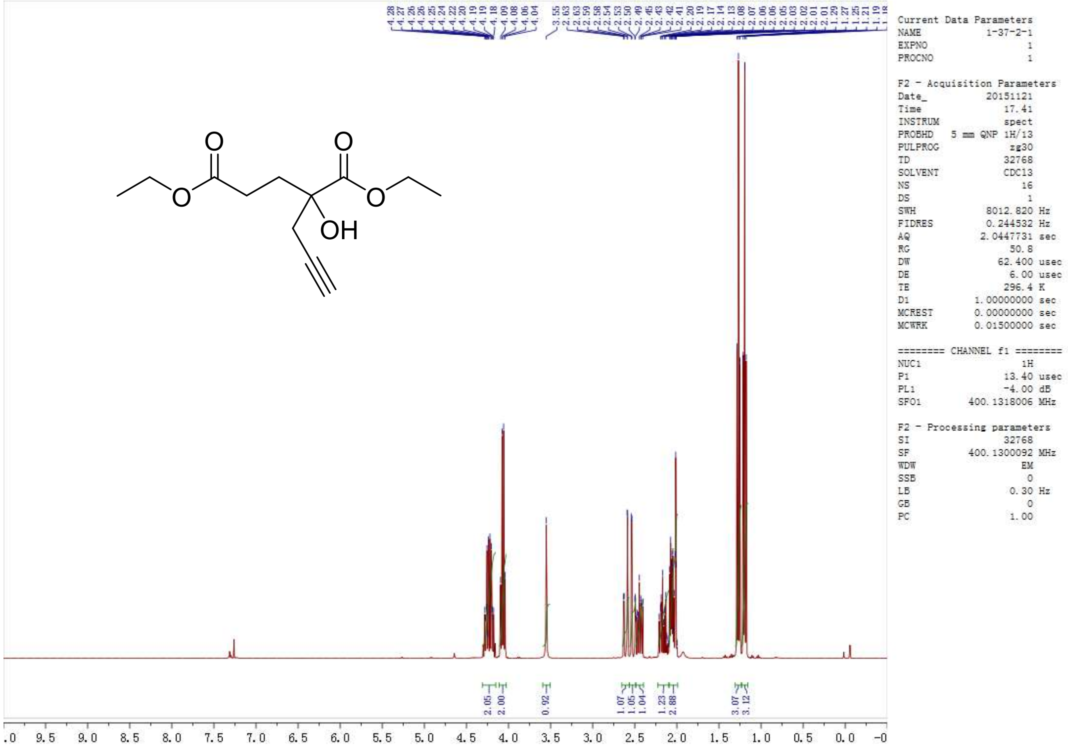

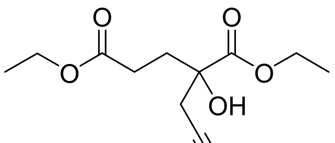

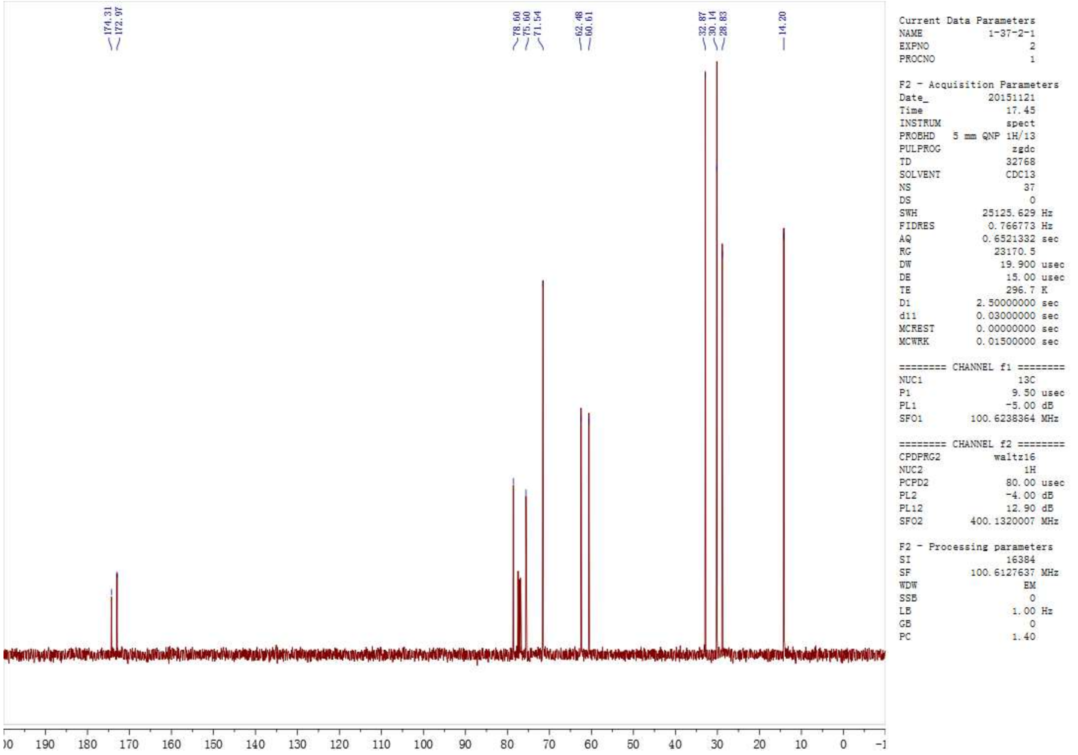
174.31
172.97
78.60
75.60
71.54
62.48
60.61
32.87
30.14
28.83
14.20
Current Data Parameters
NAME 1-37-2-1
EXPNO 2
PROCNO 1
F2 - Acquisition Parameters
Date_ 20151121
Time 17.45
INSTRUM spect
PROBHD 5 mm QNP 1H/13
PULPROG zgdc
TD 32768
SOLVENT CDCl3
NS 37
DS 0
SWH 25125.629 Hz
FIDRES 0.766773 Hz
AQ 0.6521332 sec
RG 23170.5
DW 19.900 usec
DE 15.00 usec
TE 296.7 K
D1 2.50000000 sec
d11 0.03000000 sec
MCREST 0.00000000 sec
MCWRK 0.01500000 sec
======== CHANNEL f1 ========
NUC1 13C
P1 9.50 usec
PL1 -5.00 dB
SFO1 100.6238364 MHz
======== CHANNEL f2 ========
CPDPRG2 waltz16
NUC2 1H
PCPD2 80.00 usec
PL2 -4.00 dB
PL12 12.90 dB
SFO2 400.1320007 MHz
F2 - Processing parameters
SI 16384
SF 100.6127637 MHz
WDW EM
SSB 0
LB 1.00 Hz
GB 0
PC 1.40
190 180 170 160 150 140 130 120 110 100 90 80 70 60 50 40 30 20 10 0

**3.6**

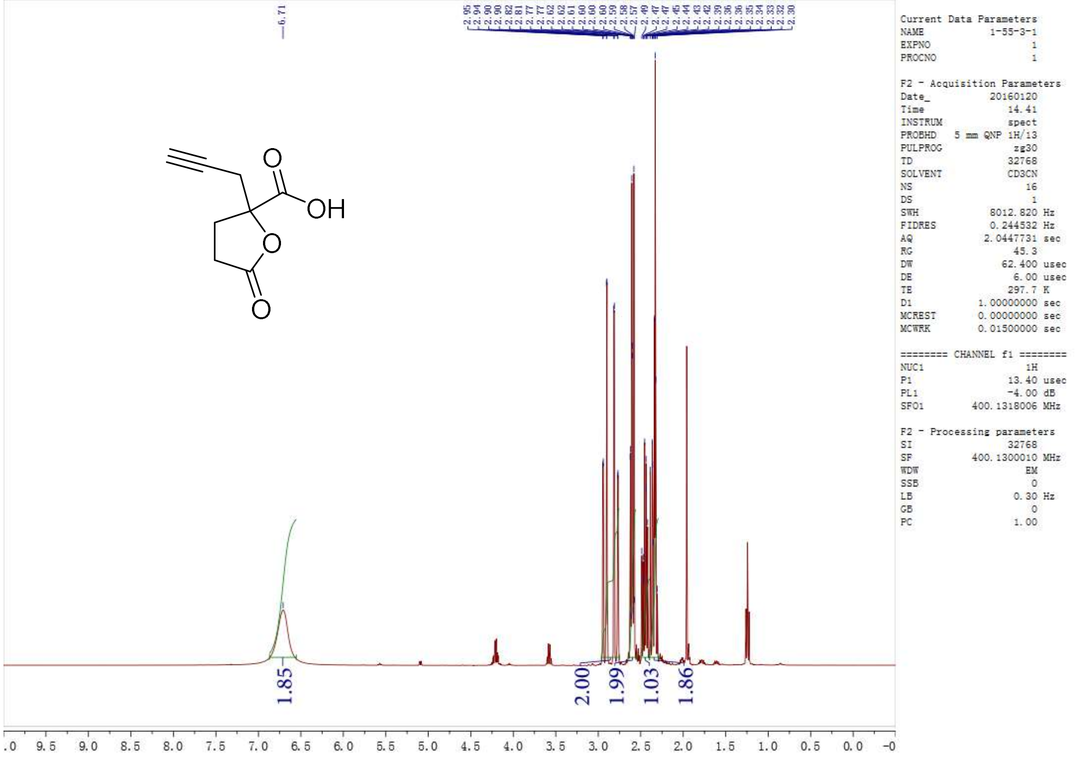
6.71
2.95 2.94 2.90 2.90 2.82 2.81 2.77 2.62 2.61 2.60 2.60 2.59 2.58 2.57 2.49 2.47 2.45 2.44 2.43 2.42 2.39 2.36 2.35 2.34 2.33 2.32 2.30
OH
O
O
O
Current Data Parameters
NAME 1-55-3-1
EXPNO 1
PROCNO 1
F2 - Acquisition Parameters
Date_ 20160120
Time 14.41
INSTRUM spect
PROBHD 5 mm QNP 1H/13
PULPROG zg30
TD 32768
SOLVENT CD3CN
NS 16
DS 1
SWH 8012.820 Hz
FIDRES 0.244532 Hz
AQ 2.0447731 sec
RG 45.3
DW 62.400 usec
DE 6.00 usec
TE 297.7 K
D1 1.00000000 sec
MCREST 0.00000000 sec
MCWRK 0.01500000 sec
======== CHANNEL f1 ========
NUC1 1H
P1 13.40 usec
PL1 -4.00 dB
SFO1 400.1318006 MHz
F2 - Processing parameters
SI 32768
SF 400.1300010 MHz
WDW EM
SSB 0
LB 0.30 Hz
GB 0
PC 1.00
1.85
2.00
1.99
1.03
1.86
9.5 9.0 8.5 8.0 7.5 7.0 6.5 6.0 5.5 5.0 4.5 4.0 3.5 3.0 2.5 2.0 1.5 1.0 0.5 0.0

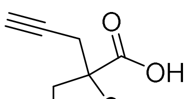
O
OH

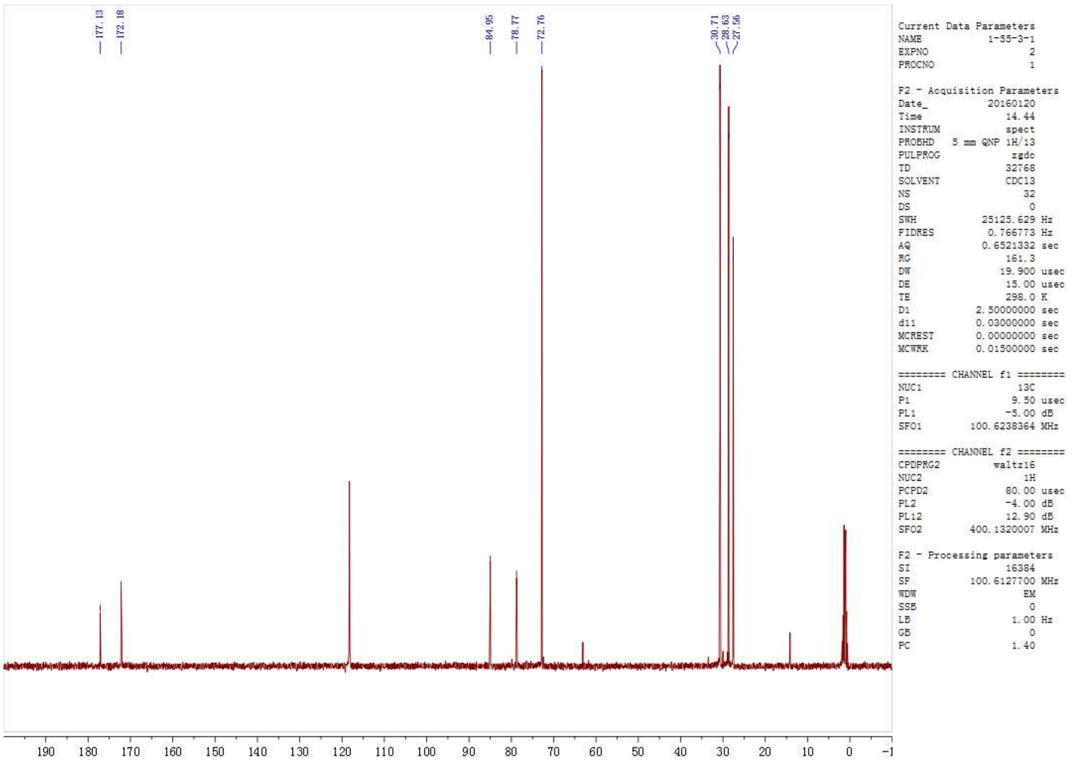
177.13
172.18
84.95
78.77
72.76
30.71
28.63
27.56
Current Data Parameters
NAME 1-55-3-1
EXPNO 2
PROCNO 1
F2 - Acquisition Parameters
Date_ 20160120
Time 14.44
INSTRUM spect
PROBHD 5 mm QNP 1H/13
PULPROG zgdc
TD 32768
SOLVENT CDCl3
NS 32
DS 0
SWH 25125.629 Hz
FIDRES 0.766773 Hz
AQ 0.6521332 sec
RG 161.3
DW 19.900 usec
DE 15.00 usec
TE 298.0 K
D1 2.50000000 sec
d11 0.03000000 sec
MCREST 0.00000000 sec
MCWRK 0.01500000 sec
======== CHANNEL f1 ========
NUC1 13C
P1 9.50 usec
PL1 -5.00 dB
SFO1 100.6238364 MHz
======== CHANNEL f2 ========
CPDPRG2 waltz16
NUC2 1H
PCPD2 80.00 usec
PL2 -4.00 dB
PL12 12.90 dB
SFO2 400.1320007 MHz
F2 - Processing parameters
SI 16384
SF 100.6127700 MHz
WDW EM
SSB 0
LB 1.00 Hz
GB 0
PC 1.40
190 180 170 160 150 140 130 120 110 100 90 80 70 60 50 40 30 20 10 0 -1

**3.19-1**

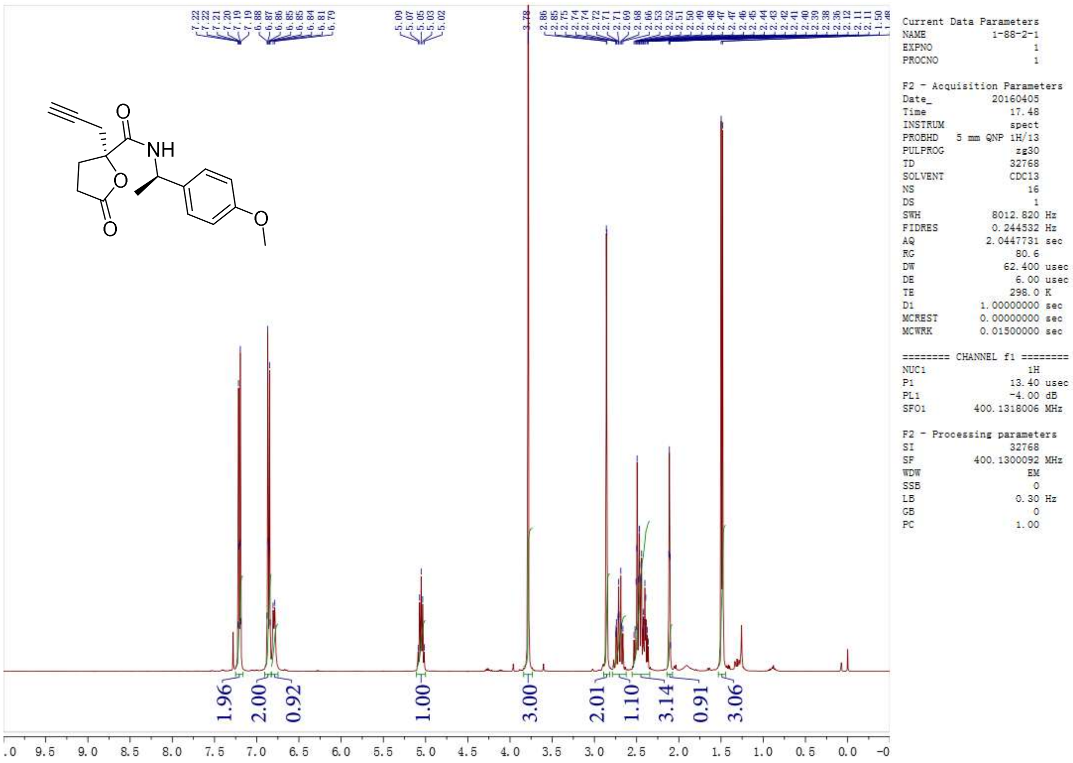
7.22
7.22
7.21
7.20
7.19
7.19
6.88
6.87
6.86
6.85
6.85
6.84
6.81
6.79
5.09
5.07
5.05
5.03
5.02
3.78
2.86
2.85
2.75
2.74
2.72
2.71
2.71
2.69
2.68
2.66
2.53
2.52
2.51
2.50
2.49
2.48
2.47
2.47
2.46
2.45
2.44
2.43
2.42
2.41
2.40
2.39
2.38
2.36
2.12
2.11
1.50
1.48
O
NH
O
O
O
Current Data Parameters
NAME 1-88-2-1
EXPNO 1
PROCNO 1
F2 - Acquisition Parameters
Date_ 20160405
Time 17.48
INSTRUM spect
PROBHD 5 mm QNP 1H/13
PULPROG zg30
TD 32768
SOLVENT CDCl3
NS 16
DS 1
SWH 8012.820 Hz
FIDRES 0.244532 Hz
AQ 2.0447731 sec
RG 80.6
DW 62.400 usec
DE 6.00 usec
TE 298.0 K
D1 1.00000000 sec
MCREST 0.00000000 sec
MCWRK 0.01500000 sec
======== CHANNEL f1 ========
NUC1 1H
P1 13.40 usec
PL1 -4.00 dB
SFO1 400.1318006 MHz
F2 - Processing parameters
SI 32768
SF 400.1300092 MHz
WDW EM
SSB 0
LB 0.30 Hz
GB 0
PC 1.00
1.96
2.00
0.92
1.00
3.00
2.01
1.10
3.14
0.91
3.06
.0 9.5 9.0 8.5 8.0 7.5 7.0 6.5 6.0 5.5 5.0 4.5 4.0 3.5 3.0 2.5 2.0 1.5 1.0 0.5 0.0 -0

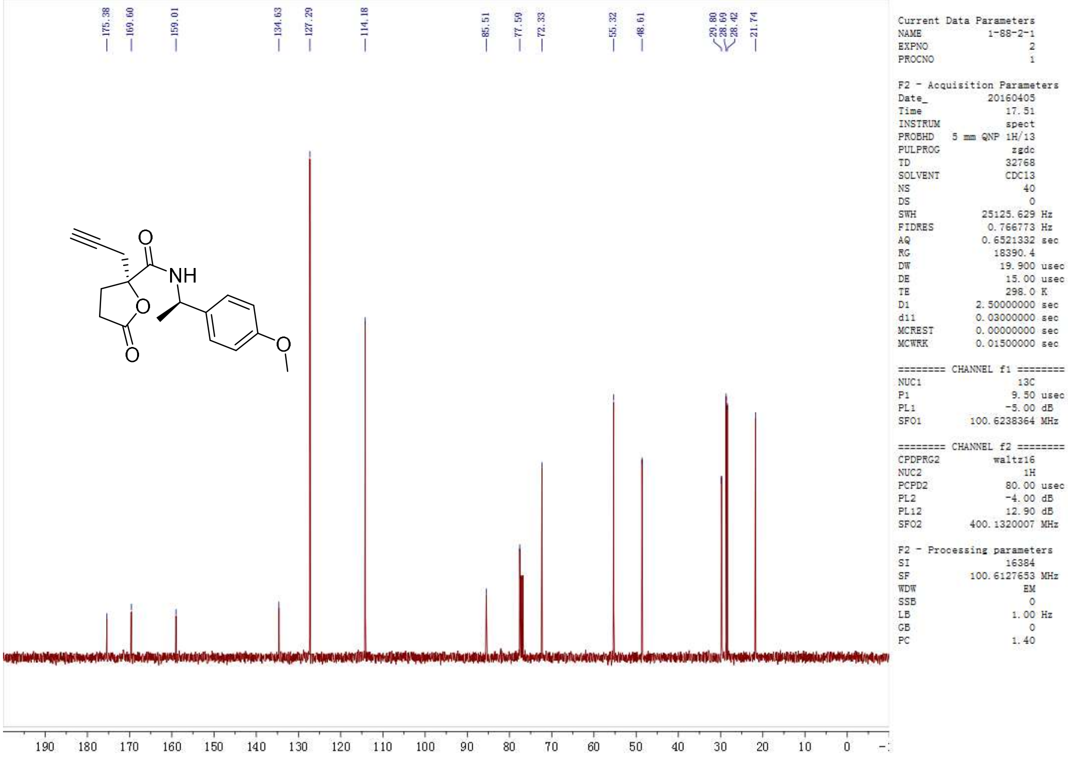
175.38
169.60
159.01
134.63
127.29
114.18
85.51
77.59
72.33
55.32
48.61
29.80
28.69
28.42
21.74
NH
O
Current Data Parameters
NAME 1-88-2-1
EXPNO 2
PROCNO 1
F2 - Acquisition Parameters
Date_ 20160405
Time 17.51
INSTRUM spect
PROBHD 5 mm QNP 1H/13
PULPROG zgdc
TD 32768
SOLVENT CDCl3
NS 40
DS 0
SWH 25125.629 Hz
FIDRES 0.766773 Hz
AQ 0.6521332 sec
RG 18390.4
DW 19.900 usec
DE 15.00 usec
TE 298.0 K
D1 2.50000000 sec
d11 0.03000000 sec
MCREST 0.00000000 sec
MCWRK 0.01500000 sec
======== CHANNEL f1 ========
NUC1 13C
P1 9.50 usec
PL1 -5.00 dB
SFO1 100.6238364 MHz
======== CHANNEL f2 ========
CPDPRG2 waltz16
NUC2 1H
PCPD2 80.00 usec
PL2 -4.00 dB
PL12 12.90 dB
SFO2 400.1320007 MHz
F2 - Processing parameters
SI 16384
SF 100.6127653 MHz
WDW EM
SSB 0
LB 1.00 Hz
GB 0
PC 1.40
190 180 170 160 150 140 130 120 110 100 90 80 70 60 50 40 30 20 10 0

**3.19-2**

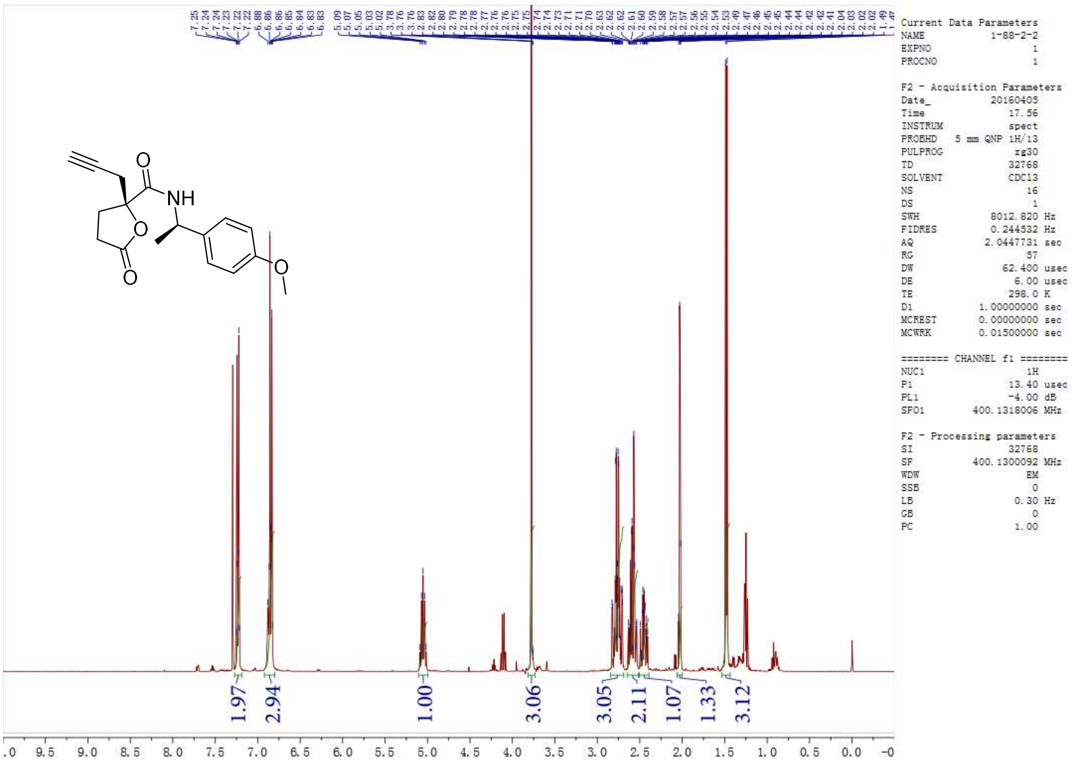
NH
O
Current Data Parameters
NAME 1-88-2-2
EXPNO 1
PROCNO 1
F2 - Acquisition Parameters
Date_ 20160405
Time 17.56
INSTRUM spect
PROBHD 5 mm QNP 1H/13
PULPROG zg30
TD 32768
SOLVENT CDCl3
NS 16
DS 1
SWH 8012.820 Hz
FIDRES 0.244532 Hz
AQ 2.0447731 sec
RG 57
DW 62.400 usec
DE 6.00 usec
TE 298.0 K
D1 1.00000000 sec
MCREST 0.00000000 sec
MCWRK 0.01500000 sec
======== CHANNEL f1 ========
NUC1 1H
P1 13.40 usec
PL1 -4.00 dB
SFO1 400.1318006 MHz
F2 - Processing parameters
SI 32768
SF 400.1300092 MHz
WDW EM
SSB 0
LB 0.30 Hz
GB 0
PC 1.00
1.97
2.94
1.00
3.06
3.05
2.11
1.07
1.33
3.12
9.5 9.0 8.5 8.0 7.5 7.0 6.5 6.0 5.5 5.0 4.5 4.0 3.5 3.0 2.5 2.0 1.5 1.0 0.5 0.0

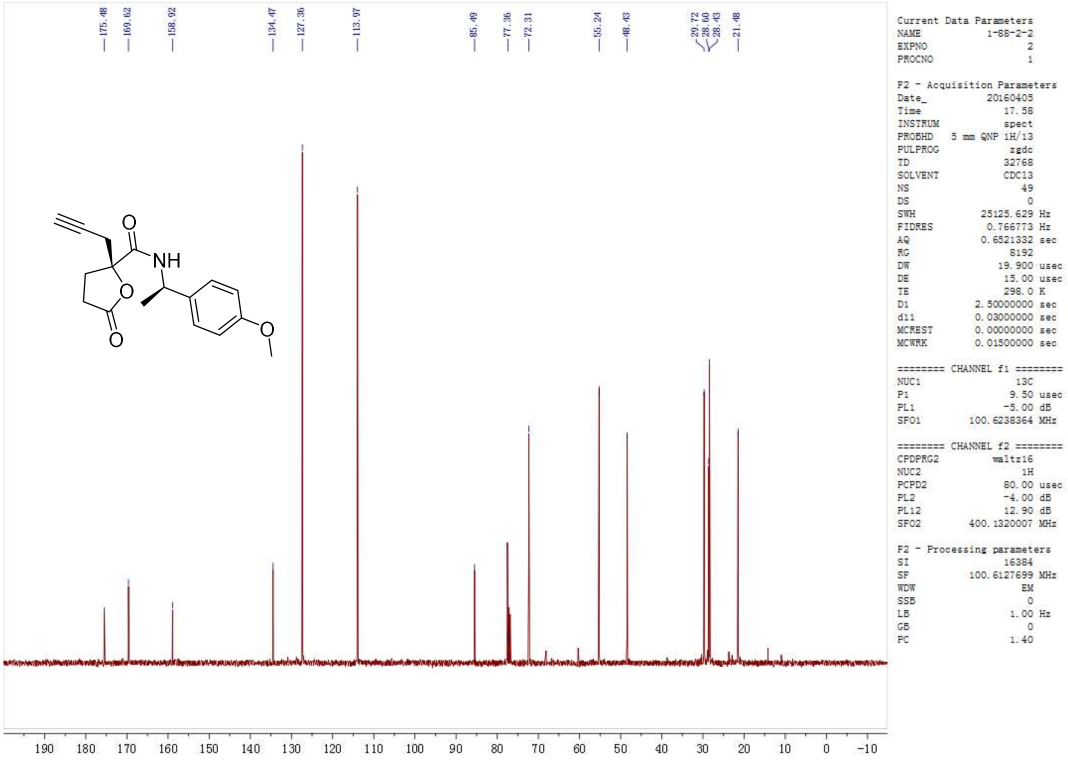

**3.20-1**

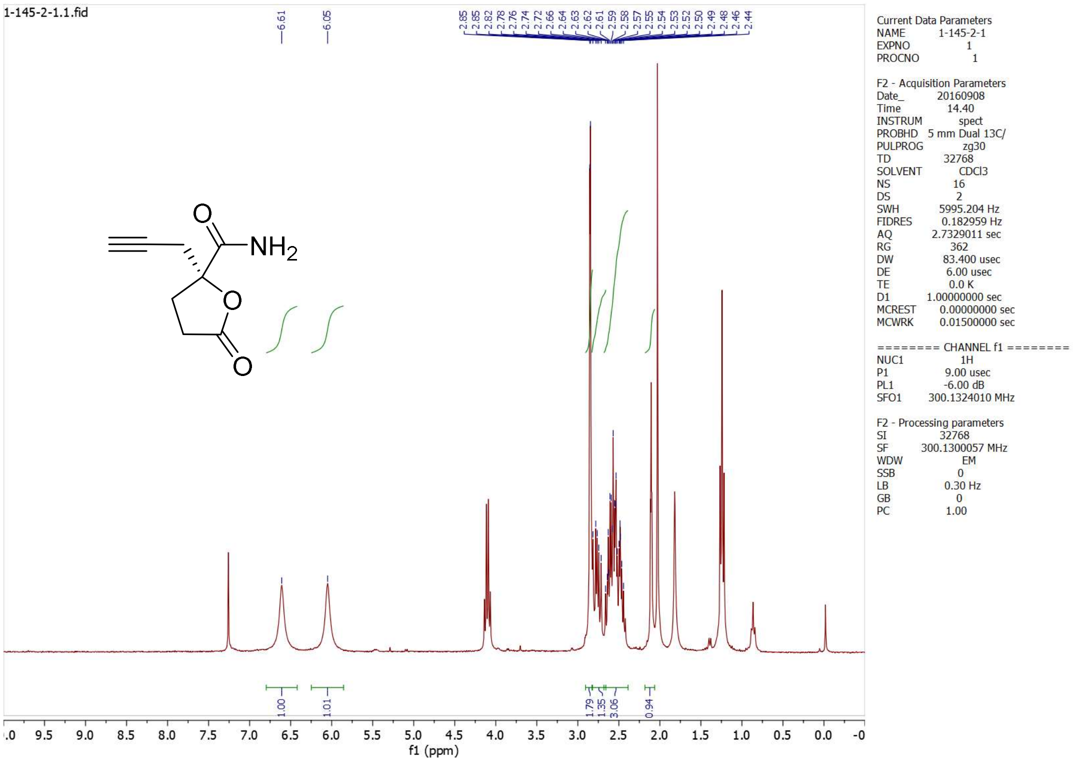

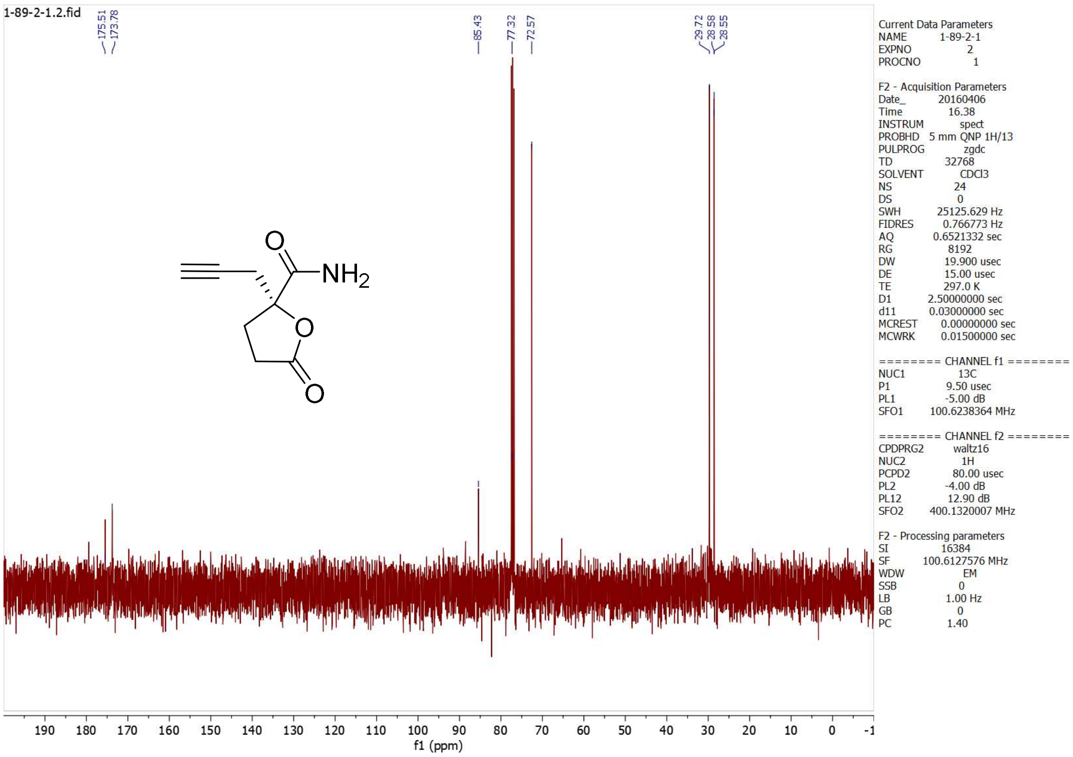
1-89-2-1.2.fid
175.51
173.78
85.43
77.32
72.57
29.72
28.58
28.55
NH2
O
Current Data Parameters
NAME 1-89-2-1
EXPNO 2
PROCNO 1
F2 - Acquisition Parameters
Date_ 20160406
Time 16.38
INSTRUM spect
PROBHD 5 mm QNP 1H/13
PULPROG zgdc
TD 32768
SOLVENT CDCl3
NS 24
DS 0
SWH 25125.629 Hz
FIDRES 0.766773 Hz
AQ 0.6521332 sec
RG 8192
DW 19.900 usec
DE 15.00 usec
TE 297.0 K
D1 2.50000000 sec
d11 0.03000000 sec
MCREST 0.00000000 sec
MCWRK 0.01500000 sec
======== CHANNEL f1 ========
NUC1 13C
P1 9.50 usec
PL1 -5.00 dB
SFO1 100.6238364 MHz
======== CHANNEL f2 ========
CPDPRG2 waltz16
NUC2 1H
PCPD2 80.00 usec
PL2 -4.00 dB
PL12 12.90 dB
SFO2 400.1320007 MHz
F2 - Processing parameters
SI 16384
SF 100.6127576 MHz
WDW EM
SSB 0
LB 1.00 Hz
GB 0
PC 1.40
f1 (ppm)

**3.20-2**

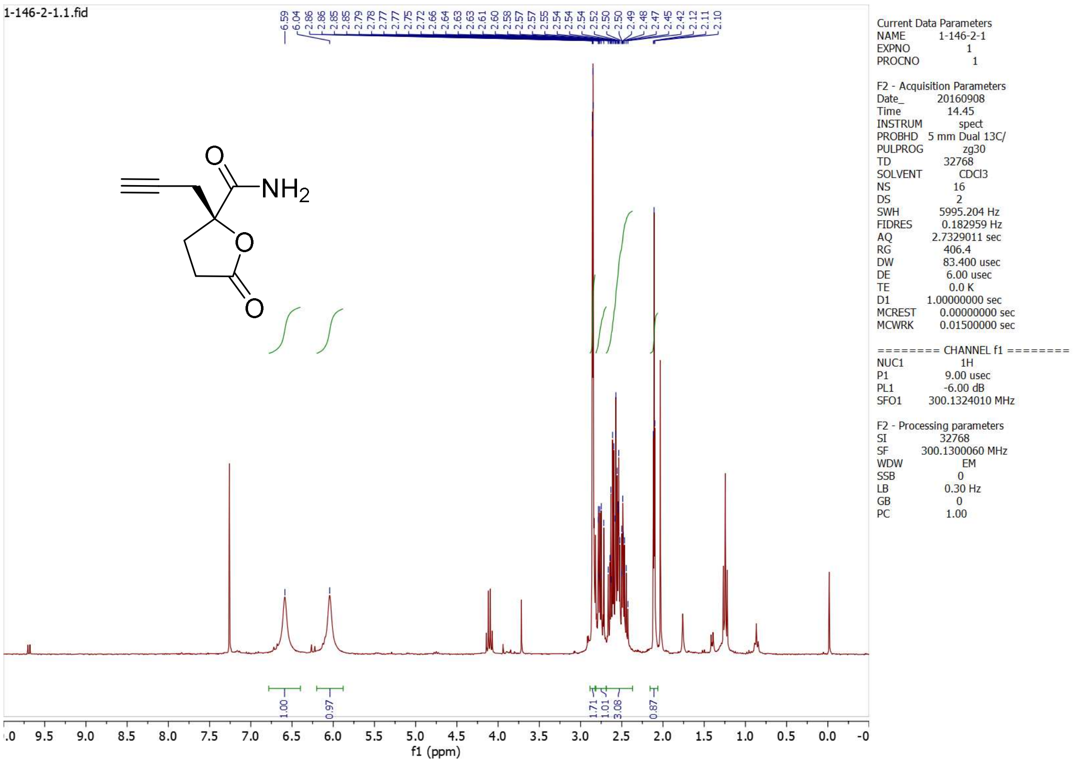
1-146-2-1.1.fid
6.59
6.04
2.86
2.86
2.85
2.85
2.79
2.78
2.77
2.75
2.72
2.66
2.64
2.63
2.63
2.61
2.60
2.58
2.57
2.57
2.55
2.54
2.54
2.54
2.52
2.50
2.50
2.49
2.48
2.47
2.45
2.42
2.12
2.11
2.10
NH2
O
1.00
0.97
1.71
1.01
3.08
0.87
Current Data Parameters
NAME 1-146-2-1
EXPNO 1
PROCNO 1
F2 - Acquisition Parameters
Date_ 20160908
Time 14.45
INSTRUM spect
PROBHD 5 mm Dual 13C/
PULPROG zg30
TD 32768
SOLVENT CDCl3
NS 16
DS 2
SWH 5995.204 Hz
FIDRES 0.182959 Hz
AQ 2.7329011 sec
RG 406.4
DW 83.400 usec
DE 6.00 usec
TE 0.0 K
D1 1.00000000 sec
MCREST 0.00000000 sec
MCWRK 0.01500000 sec
======== CHANNEL f1 ========
NUC1 1H
P1 9.00 usec
PL1 -6.00 dB
SFO1 300.1324010 MHz
F2 - Processing parameters
SI 32768
SF 300.1300060 MHz
WDW EM
SSB 0
LB 0.30 Hz
GB 0
PC 1.00
f1 (ppm)

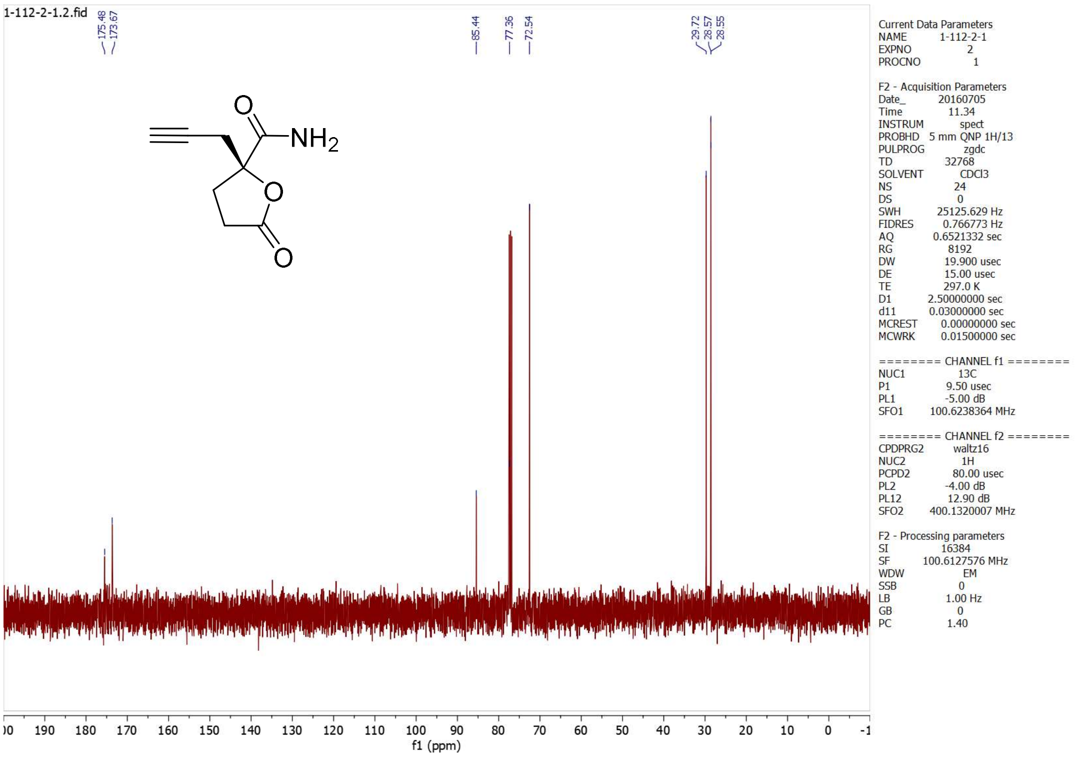
1-112-2-1.2.fid
175.48
173.67
85.44
77.36
72.54
29.72
28.57
28.55
NH2
O
Current Data Parameters
NAME 1-112-2-1
EXPNO 2
PROCNO 1
F2 - Acquisition Parameters
Date_ 20160705
Time 11.34
INSTRUM spect
PROBHD 5 mm QNP 1H/13
PULPROG zgdc
TD 32768
SOLVENT CDCl3
NS 24
DS 0
SWH 25125.629 Hz
FIDRES 0.766773 Hz
AQ 0.6521332 sec
RG 8192
DW 19.900 usec
DE 15.00 usec
TE 297.0 K
D1 2.50000000 sec
d11 0.03000000 sec
MCREST 0.00000000 sec
MCWRK 0.01500000 sec
======== CHANNEL f1 ========
NUC1 13C
P1 9.50 usec
PL1 -5.00 dB
SFO1 100.6238364 MHz
======== CHANNEL f2 ========
CPDPRG2 waltz16
NUC2 1H
PCPD2 80.00 usec
PL2 -4.00 dB
PL12 12.90 dB
SFO2 400.1320007 MHz
F2 - Processing parameters
SI 16384
SF 100.6127576 MHz
WDW EM
SSB 0
LB 1.00 Hz
GB 0
PC 1.40
f1 (ppm)

**3-1 (3*R*)**

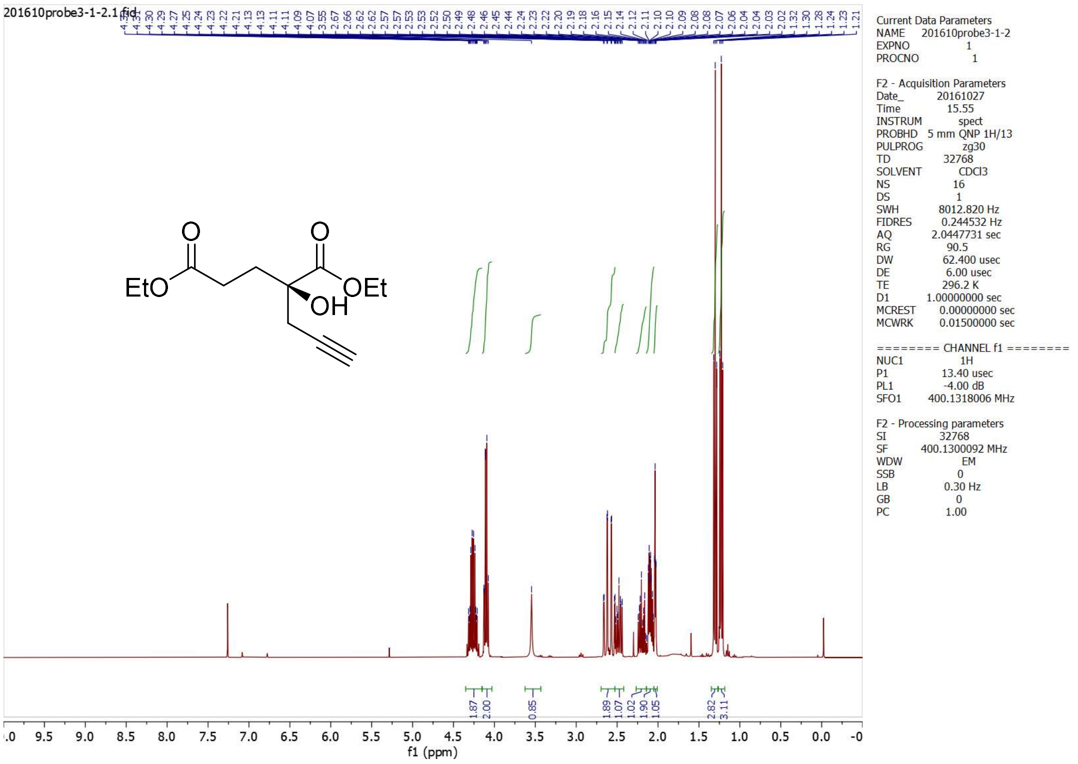
201610probe3-1-2.1.fid
EtO
OEt
OH
O
Current Data Parameters
NAME 201610probe3-1-2
EXPNO 1
PROCNO 1
F2 - Acquisition Parameters
Date_ 20161027
Time 15.55
INSTRUM spect
PROBHD 5 mm QNP 1H/13
PULPROG zg30
TD 32768
SOLVENT CDCl3
NS 16
DS 1
SWH 8012.820 Hz
FIDRES 0.244532 Hz
AQ 2.0447731 sec
RG 90.5
DW 62.400 usec
DE 6.00 usec
TE 296.2 K
D1 1.00000000 sec
MCREST 0.00000000 sec
MCWRK 0.01500000 sec
======== CHANNEL f1 ========
NUC1 1H
P1 13.40 usec
PL1 -4.00 dB
SFO1 400.1318006 MHz
F2 - Processing parameters
SI 32768
SF 400.1300092 MHz
WDW EM
SSB 0
LB 0.30 Hz
GB 0
PC 1.00
1.87
2.00
0.85
1.89
1.07
1.02
1.90
1.05
2.82
3.11
f1 (ppm)

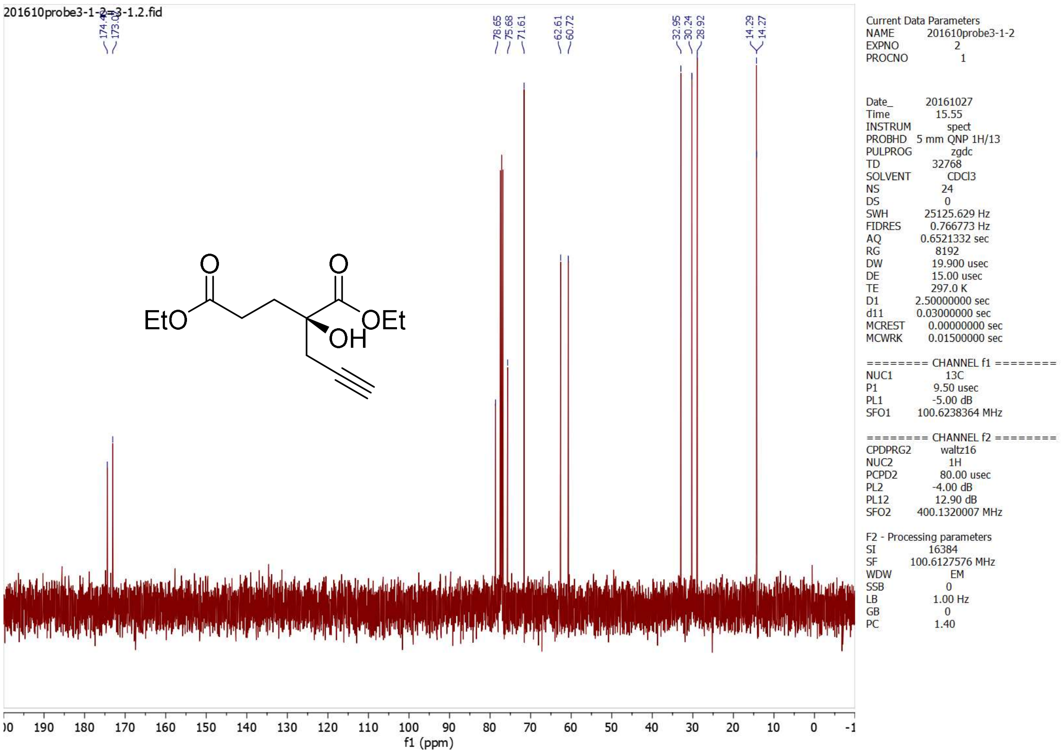
201610probe3-1-2=3-1.2.fid
78.65
75.68
71.61
62.61
60.72
32.95
30.24
28.92
14.29
14.27
EtO
O
O
OEt
OH
Current Data Parameters
NAME 201610probe3-1-2
EXPNO 2
PROCNO 1
Date_ 20161027
Time 15.55
INSTRUM spect
PROBHD 5 mm QNP 1H/13
PULPROG zgdc
TD 32768
SOLVENT CDCl3
NS 24
DS 0
SWH 25125.629 Hz
FIDRES 0.766773 Hz
AQ 0.6521332 sec
RG 8192
DW 19.900 usec
DE 15.00 usec
TE 297.0 K
D1 2.50000000 sec
d11 0.03000000 sec
MCREST 0.00000000 sec
MCWRK 0.01500000 sec
======== CHANNEL f1 ========
NUC1 13C
P1 9.50 usec
PL1 -5.00 dB
SFO1 100.6238364 MHz
======== CHANNEL f2 ========
CPDPRG2 waltz16
NUC2 1H
PCPD2 80.00 usec
PL2 -4.00 dB
PL12 12.90 dB
SFO2 400.1320007 MHz
F2 - Processing parameters
SI 16384
SF 100.6127576 MHz
WDW EM
SSB 0
LB 1.00 Hz
GB 0
PC 1.40
f1 (ppm)

**3-2 (3*S*)**

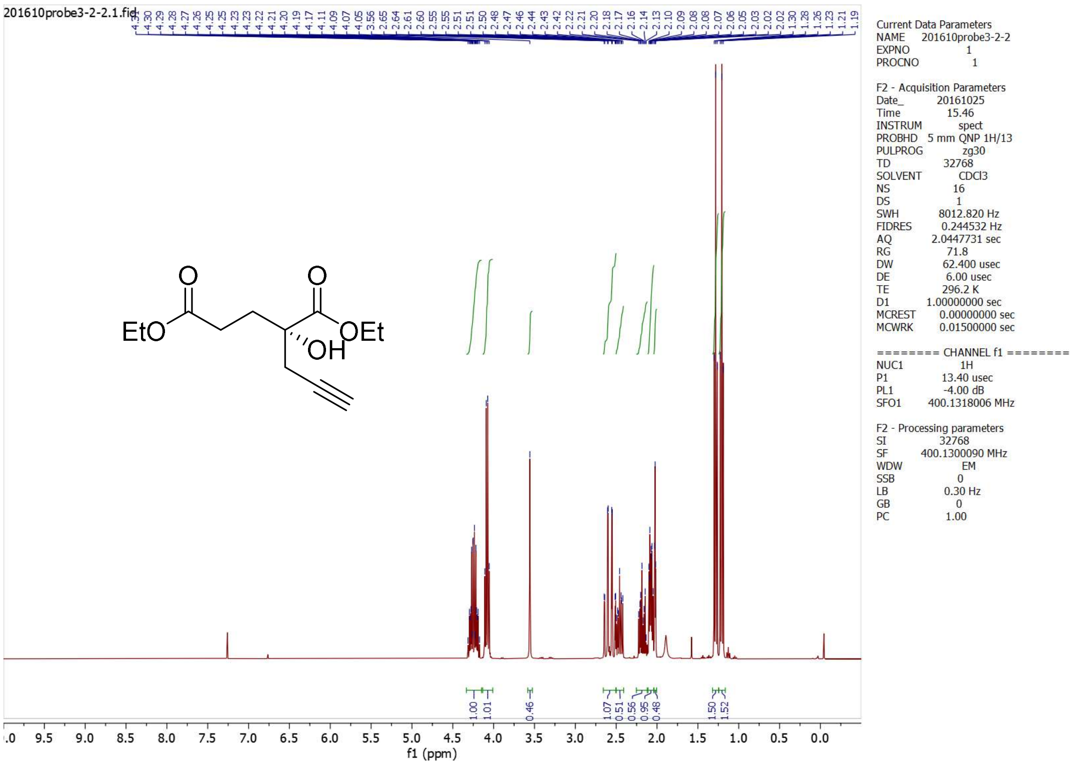
201610probe3-2-2.1.fid
EtO
O
O
OEt
OH
1.00
1.01
0.46
1.07
0.51
0.56
0.95
0.48
1.50
1.52
Current Data Parameters
NAME 201610probe3-2-2
EXPNO 1
PROCNO 1
F2 - Acquisition Parameters
Date_ 20161025
Time 15.46
INSTRUM spect
PROBHD 5 mm QNP 1H/13
PULPROG zg30
TD 32768
SOLVENT CDCl3
NS 16
DS 1
SWH 8012.820 Hz
FIDRES 0.244532 Hz
AQ 2.0447731 sec
RG 71.8
DW 62.400 usec
DE 6.00 usec
TE 296.2 K
D1 1.00000000 sec
MCREST 0.00000000 sec
MCWRK 0.01500000 sec
======== CHANNEL f1 ========
NUC1 1H
P1 13.40 usec
PL1 -4.00 dB
SFO1 400.1318006 MHz
F2 - Processing parameters
SI 32768
SF 400.1300090 MHz
WDW EM
SSB 0
LB 0.30 Hz
GB 0
PC 1.00
f1 (ppm)

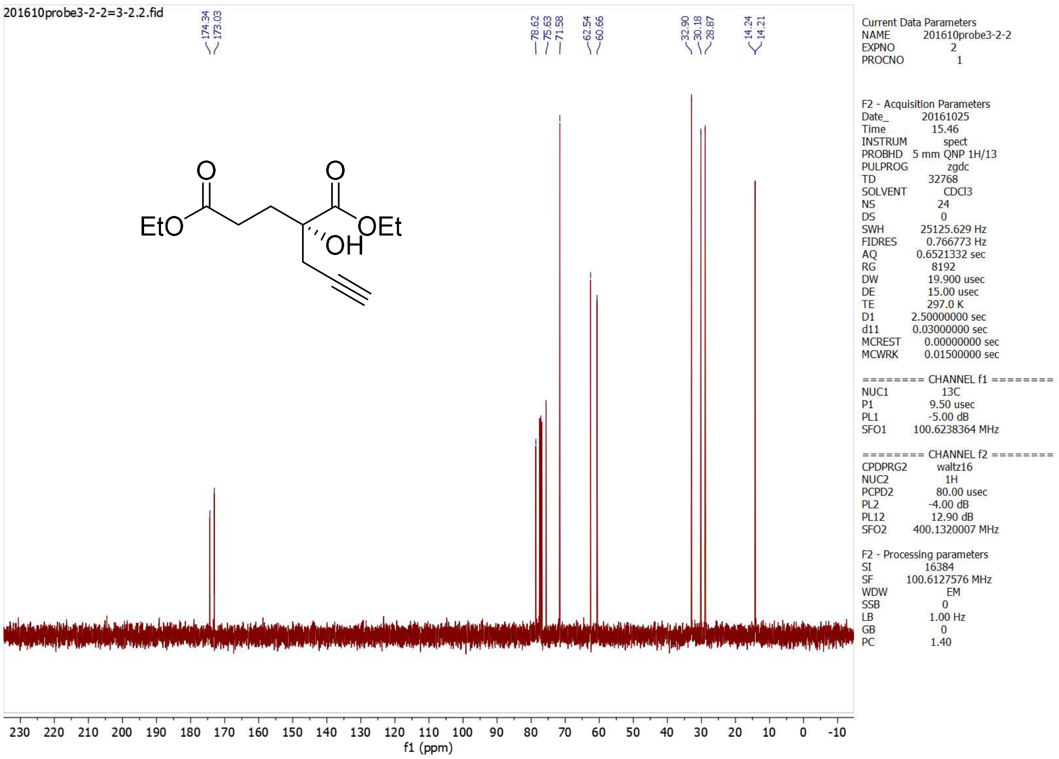

#### 4.2 diethyl $\alpha$-ketoglutarate

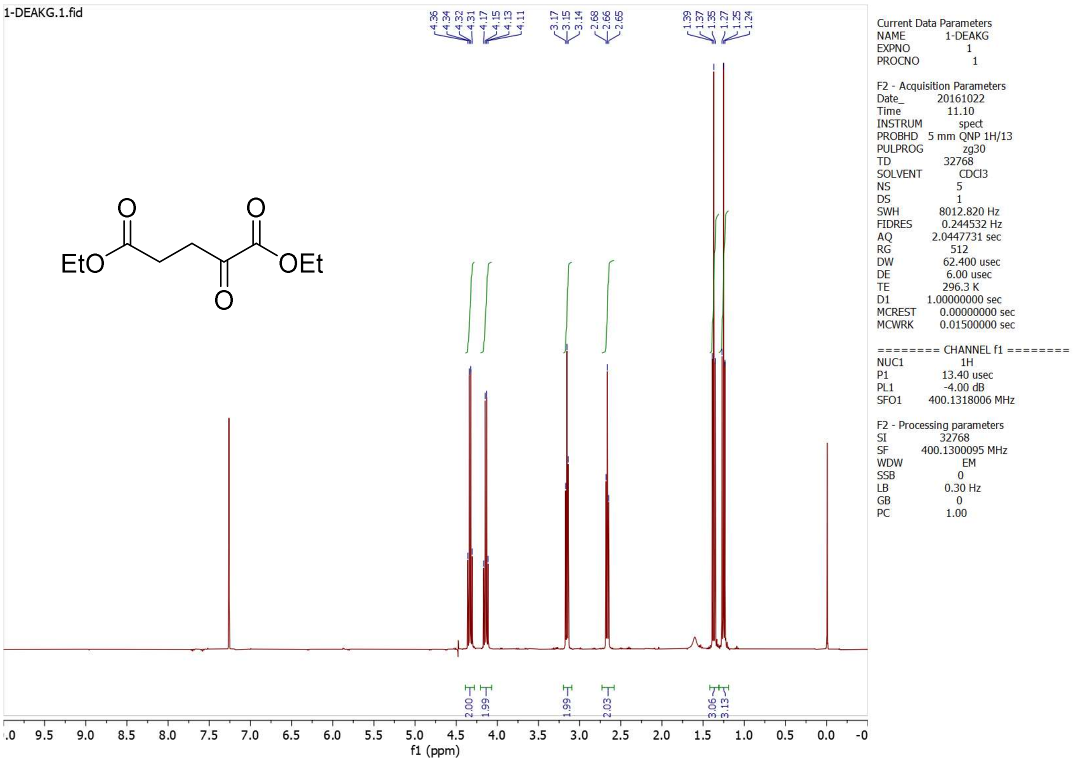

**4.10**

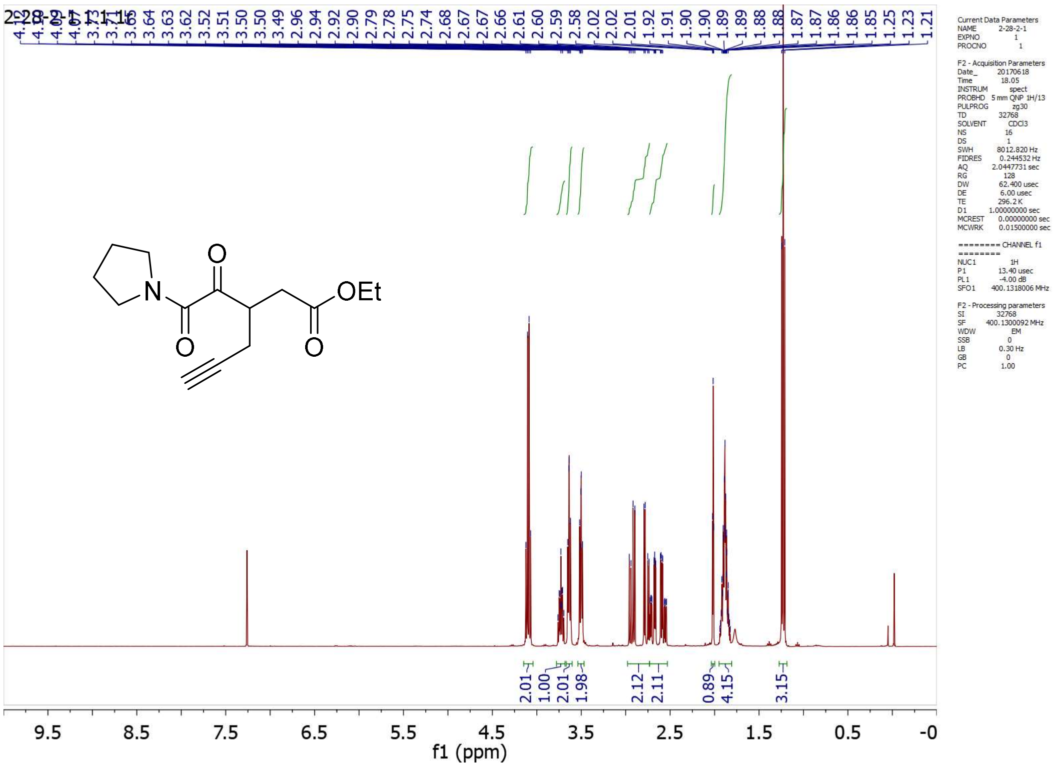

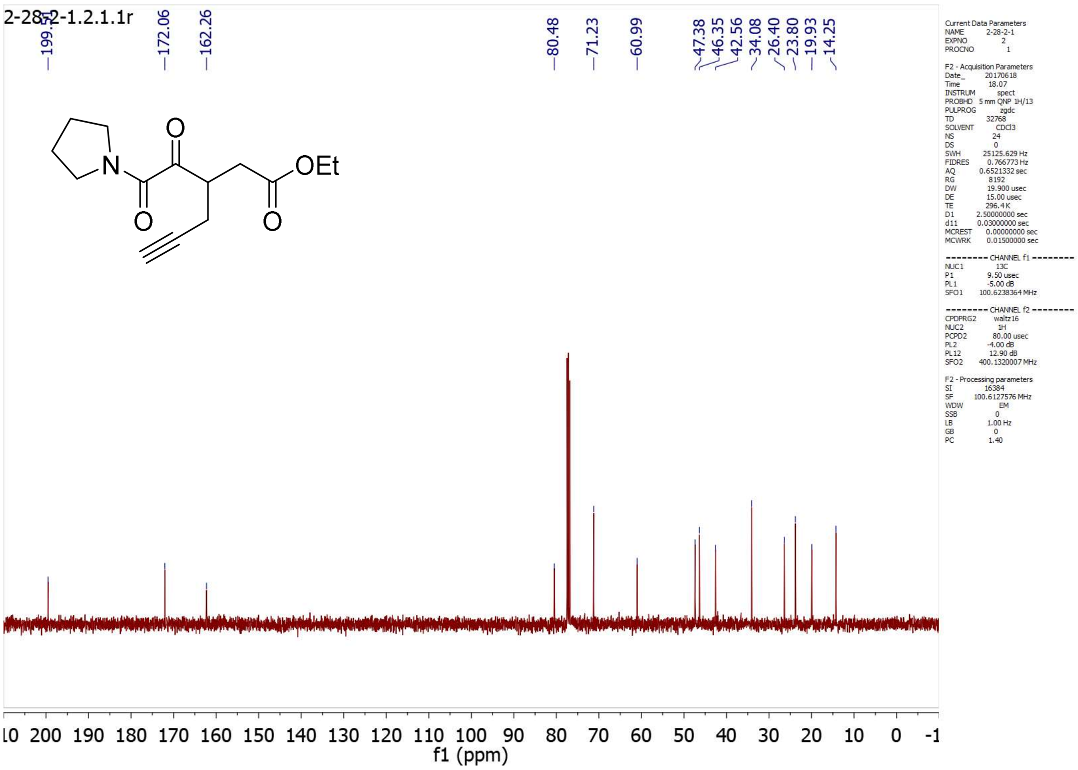

**4.11**

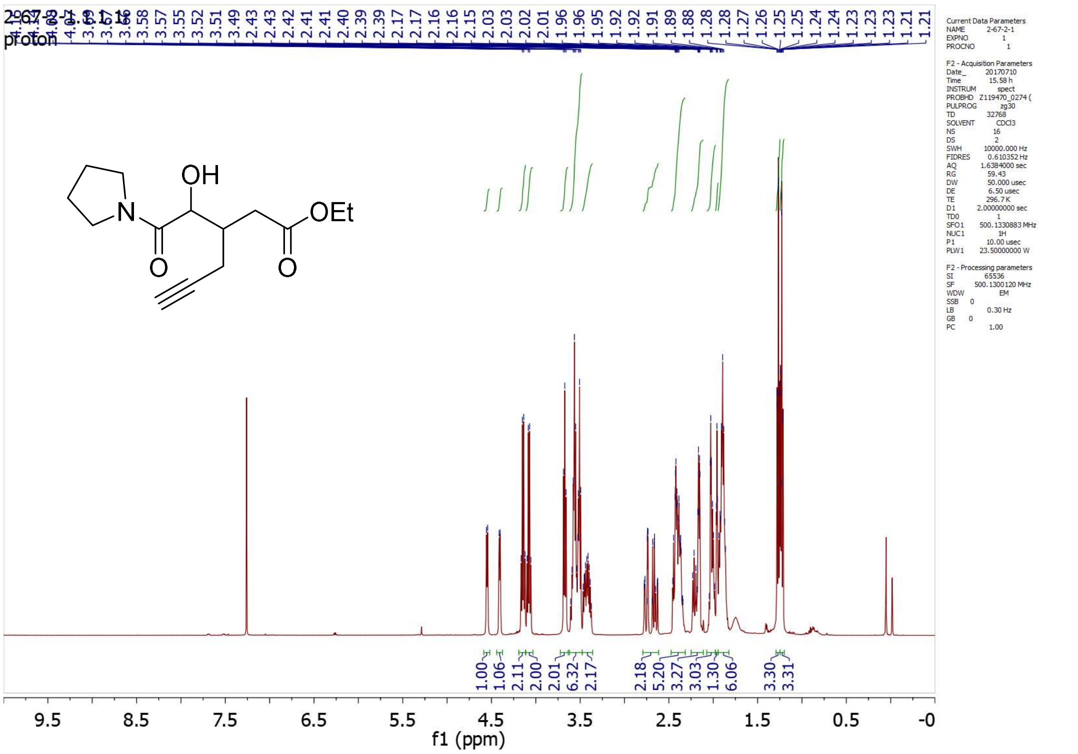

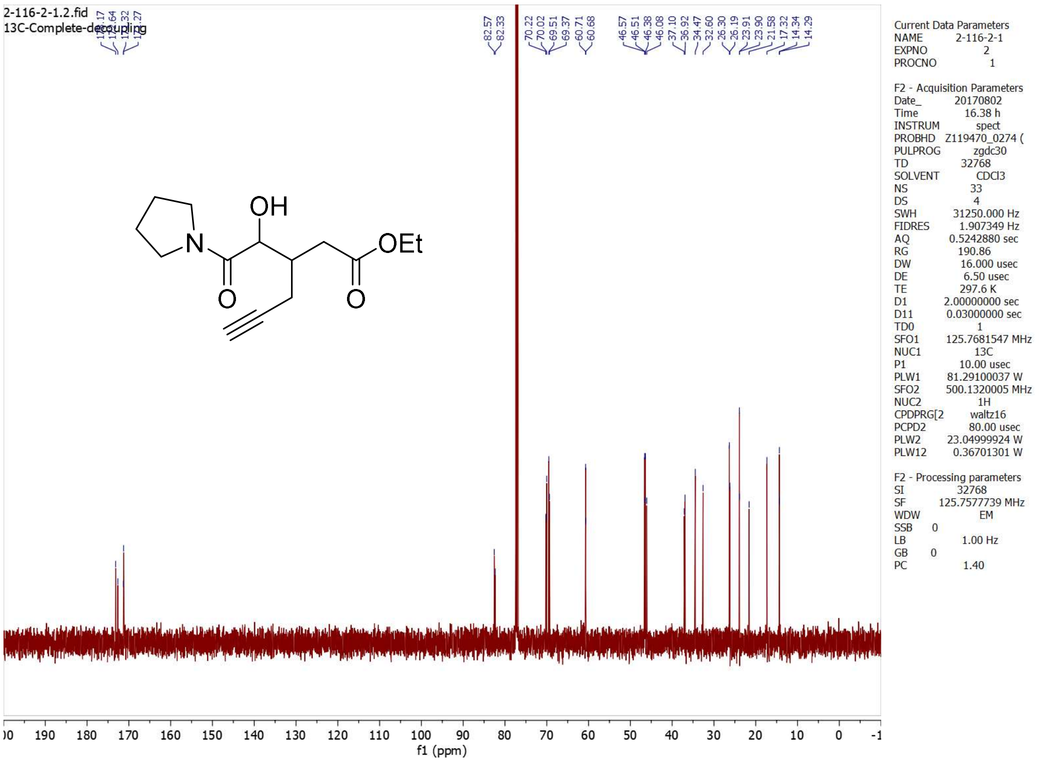
2-116-2-1.2.fid
13C-Complete-decoupling
82.57
82.33
70.22
70.02
69.51
69.37
60.71
60.68
46.57
46.51
46.38
46.08
37.10
36.92
34.47
32.60
26.30
26.19
23.91
23.90
21.58
17.32
14.34
14.29
OH
N
OEt
O
O
Current Data Parameters
NAME 2-116-2-1
EXPNO 2
PROCNO 1
F2 - Acquisition Parameters
Date_ 20170802
Time 16.38 h
INSTRUM spect
PROBHD Z119470_0274 (
PULPROG zgdc30
TD 32768
SOLVENT CDCl3
NS 33
DS 4
SWH 31250.000 Hz
FIDRES 1.907349 Hz
AQ 0.5242880 sec
RG 190.86
DW 16.000 usec
DE 6.50 usec
TE 297.6 K
D1 2.00000000 sec
D11 0.03000000 sec
TD0 1
SFO1 125.7681547 MHz
NUC1 13C
P1 10.00 usec
PLW1 81.29100037 W
SFO2 500.1320005 MHz
NUC2 1H
CPDPRG[2 waltz16
PCPD2 80.00 usec
PLW2 23.04999924 W
PLW12 0.36701301 W
F2 - Processing parameters
SI 32768
SF 125.7577739 MHz
WDW EM
SSB 0
LB 1.00 Hz
GB 0
PC 1.40
190 180 170 160 150 140 130 120 110 100 90 80 70 60 50 40 30 20 10 0 -1
f1 (ppm)

**4.12-1**

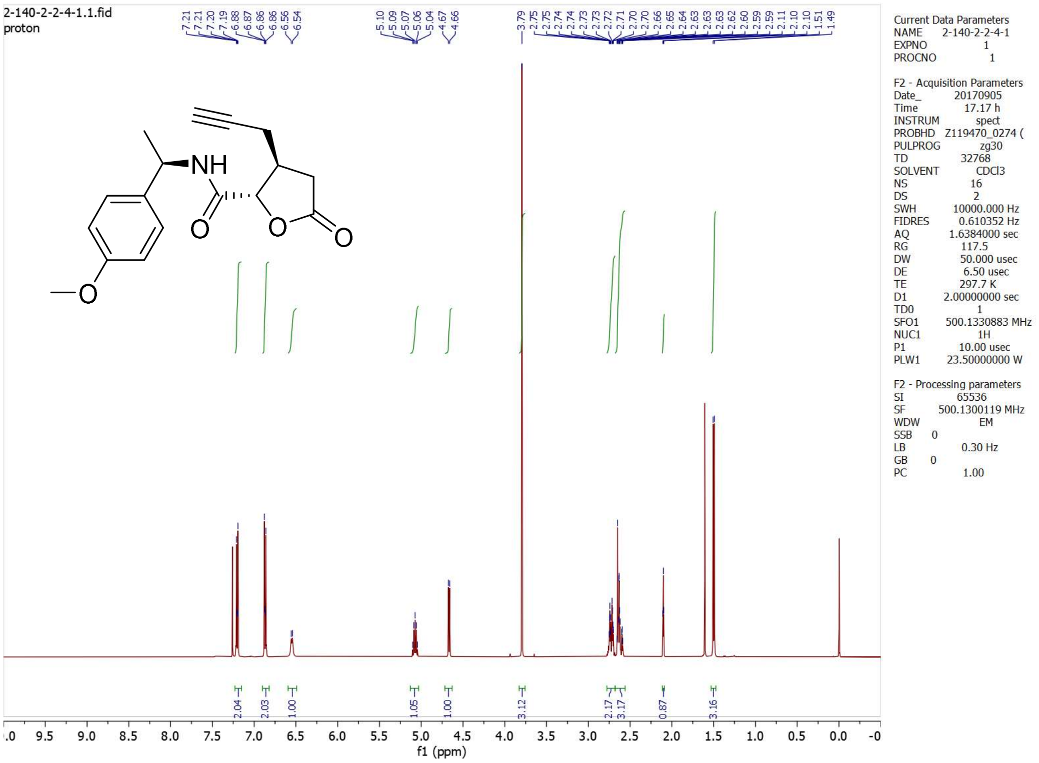
2-140-2-2-4-1.1.fid
proton
7.21
7.21
7.20
7.19
6.88
6.87
6.86
6.86
6.56
6.54
5.10
5.09
5.07
5.06
5.04
4.67
4.66
3.79
2.75
2.74
2.74
2.73
2.73
2.72
2.71
2.70
2.70
2.66
2.65
2.64
2.63
2.63
2.62
2.60
2.59
2.11
2.10
1.51
1.49
NH
O
O
O
O
Current Data Parameters
NAME 2-140-2-2-4-1
EXPNO 1
PROCNO 1
F2 - Acquisition Parameters
Date_ 20170905
Time 17.17 h
INSTRUM spect
PROBHD Z119470_0274 (
PULPROG zg30
TD 32768
SOLVENT CDCl3
NS 16
DS 2
SWH 10000.000 Hz
FIDRES 0.610352 Hz
AQ 1.6384000 sec
RG 117.5
DW 50.000 usec
DE 6.50 usec
TE 297.7 K
D1 2.00000000 sec
TD0 1
SFO1 500.1330883 MHz
NUC1 1H
P1 10.00 usec
PLW1 23.50000000 W
F2 - Processing parameters
SI 65536
SF 500.1300119 MHz
WDW EM
SSB 0
LB 0.30 Hz
GB 0
PC 1.00
2.04
2.03
1.00
1.05
1.00
3.12
2.17
3.17
0.87
3.16
9.5 9.0 8.5 8.0 7.5 7.0 6.5 6.0 5.5 5.0 4.5 4.0 3.5 3.0 2.5 2.0 1.5 1.0 0.5 0.0
f1 (ppm)

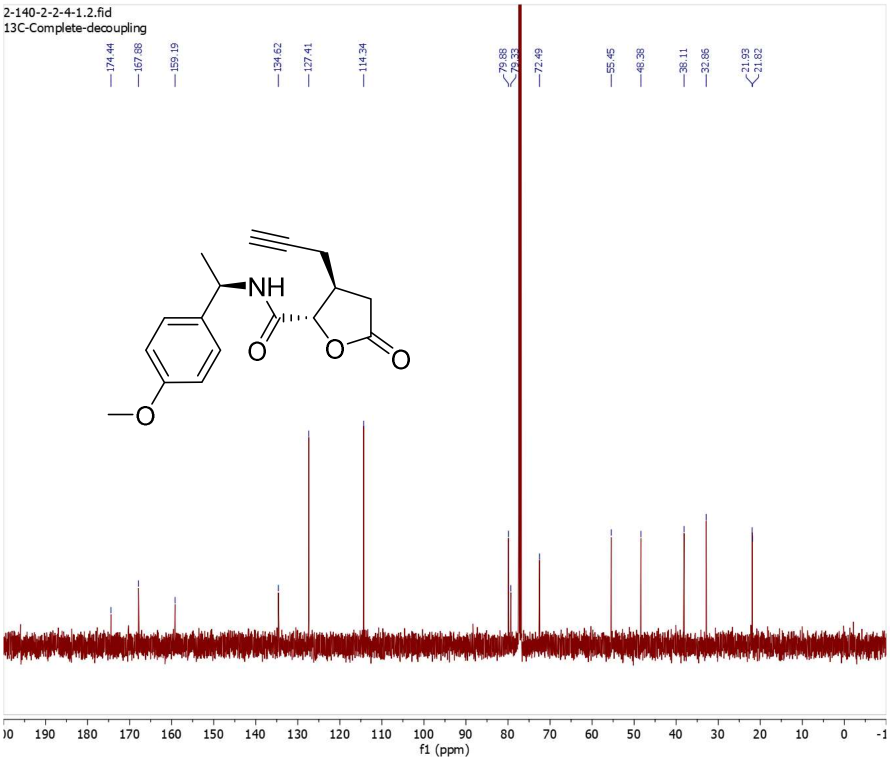

Current Data Parameters
NAME 2-140-2-2-4-1
EXPNO 2
PROCNO 1

F2 - Acquisition Parameters
Date_ 20170905
Time 17.23 h
INSTRUM spect
PROBHD Z119470_0274 (
PULPROG zgdc30
TD 32768
SOLVENT CDCl3
NS 31
DS 4
SWH 31250.000 Hz
FIDRES 1.907349 Hz
AQ 0.5242880 sec
RG 190.86
DW 16.000 usec
DE 6.50 usec
TE 298.1 K
D1 2.00000000 sec
D11 0.03000000 sec
TD0 1
SFO1 125.7681547 MHz
NUC1 13C
P1 10.00 usec
PLW1 81.29100037 W
SFO2 500.1320005 MHz
NUC2 1H
CPDPRG[2 waltz16
PCPD2 80.00 usec
PLW2 23.04999924 W
PLW12 0.36701301 W

F2 - Processing parameters
SI 32768
SF 125.7577723 MHz
WDW EM
SSB 0
LB 1.00 Hz
GB 0
PC 1.40

**4.12-2**

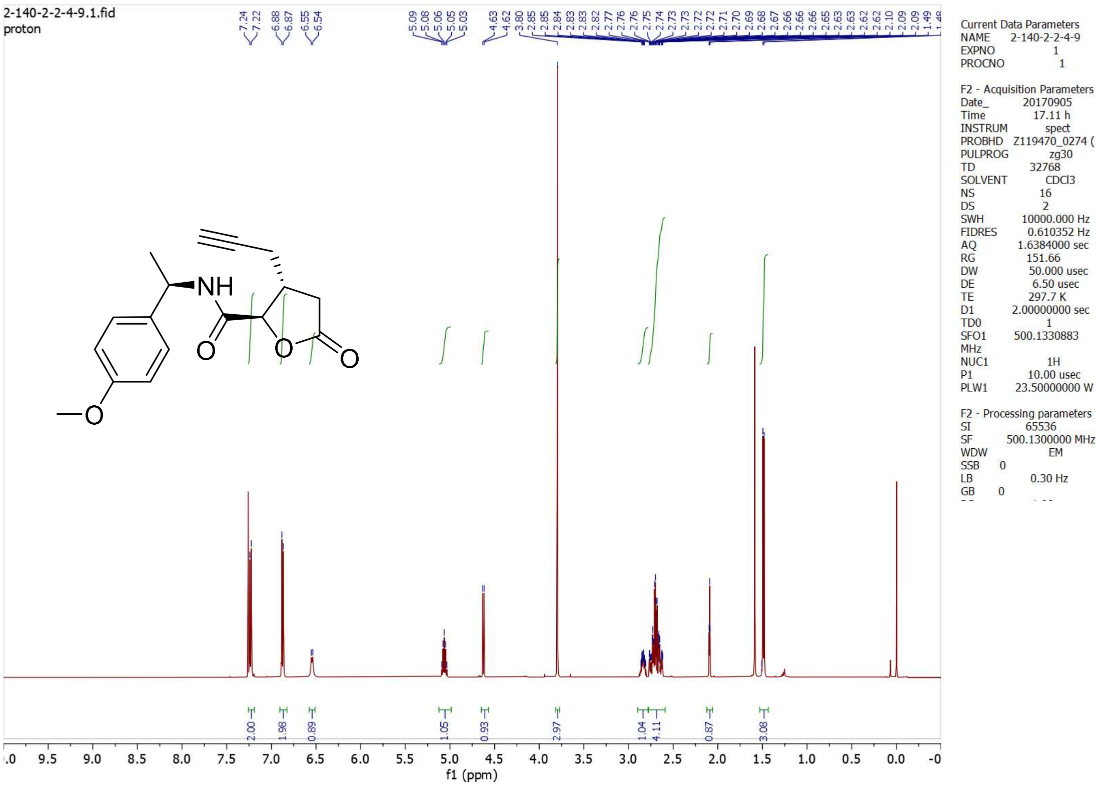

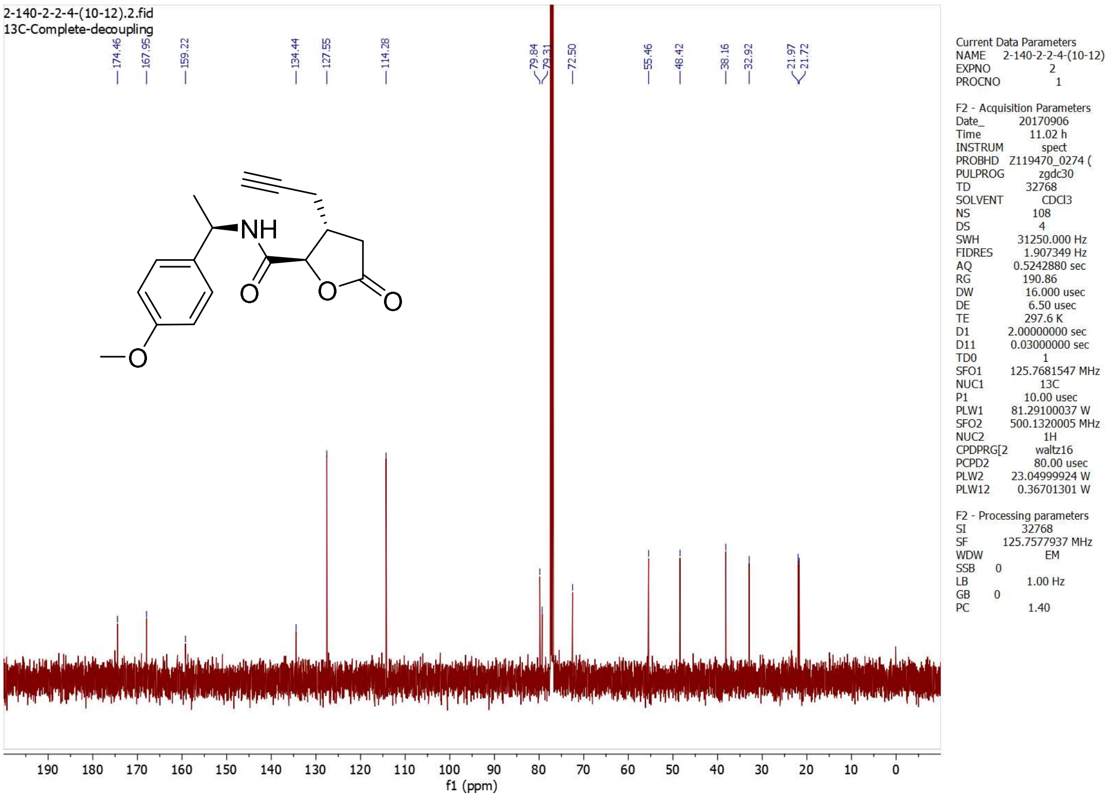

2-140-2-2-4-(10-12).2.fid
13C-Complete-decoupling
174.46
167.95
159.22
134.44
127.55
114.28
79.84
79.31
72.50
55.46
48.42
38.16
32.92
21.97
21.72
NH
O
O
O
O
190 180 170 160 150 140 130 120 110 100 90 80 70 60 50 40 30 20 10 0
f1 (ppm)
Current Data Parameters
NAME 2-140-2-2-4-(10-12)
EXPNO 2
PROCNO 1
F2 - Acquisition Parameters
Date_ 20170906
Time 11.02 h
INSTRUM spect
PROBHD Z119470_0274 (
PULPROG zgdc30
TD 32768
SOLVENT CDCl3
NS 108
DS 4
SWH 31250.000 Hz
FIDRES 1.907349 Hz
AQ 0.5242880 sec
RG 190.86
DW 16.000 usec
DE 6.50 usec
TE 297.6 K
D1 2.00000000 sec
D11 0.03000000 sec
TD0 1
SFO1 125.7681547 MHz
NUC1 13C
P1 10.00 usec
PLW1 81.29100037 W
SFO2 500.1320005 MHz
NUC2 1H
CPDPRG[2 waltz16
PCPD2 80.00 usec
PLW2 23.04999924 W
PLW12 0.36701301 W
F2 - Processing parameters
SI 32768
SF 125.7577937 MHz
WDW EM
SSB 0
LB 1.00 Hz
GB 0
PC 1.40

**4.13-1**

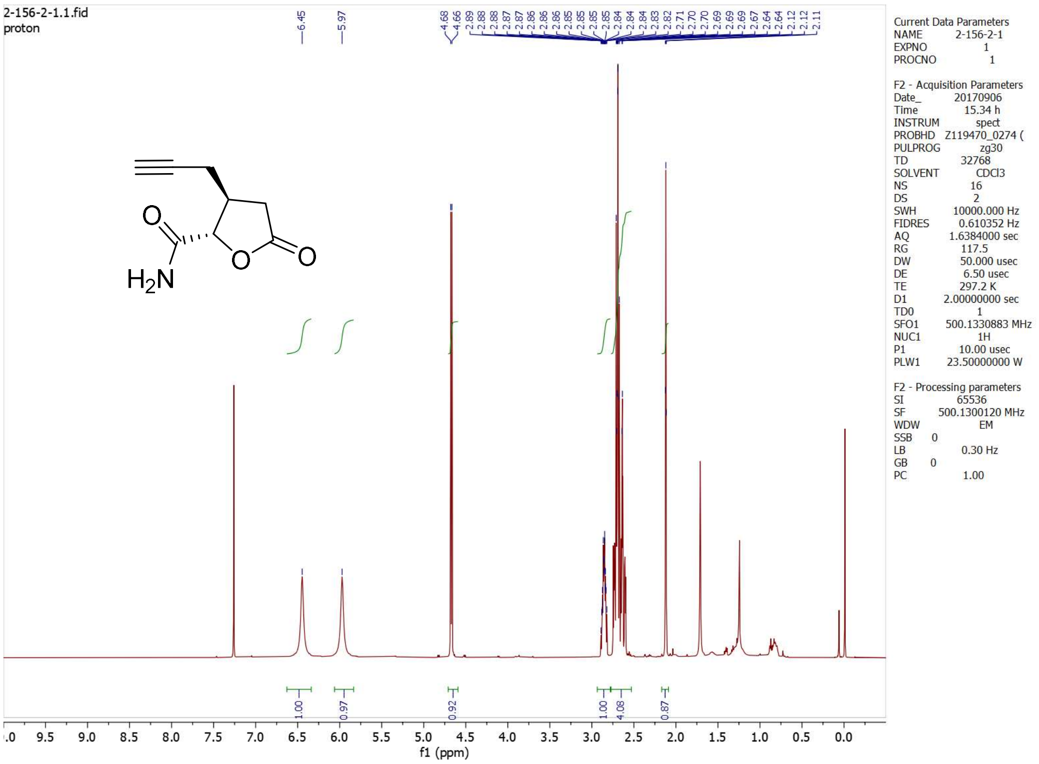

2-156-2-1.1.fid
proton
6.45
5.97
4.68
4.66
2.89
2.88
2.88
2.87
2.87
2.86
2.86
2.86
2.85
2.85
2.85
2.85
2.84
2.84
2.84
2.84
2.83
2.82
2.71
2.70
2.69
2.69
2.69
2.67
2.64
2.12
2.12
2.11
H2N
O
O
O
1.00
0.97
0.92
1.00
4.08
0.87
f1 (ppm)
Current Data Parameters
NAME 2-156-2-1
EXPNO 1
PROCNO 1
F2 - Acquisition Parameters
Date_ 20170906
Time 15.34 h
INSTRUM spect
PROBHD Z119470_0274 (
PULPROG zg30
TD 32768
SOLVENT CDCl3
NS 16
DS 2
SWH 10000.000 Hz
FIDRES 0.610352 Hz
AQ 1.6384000 sec
RG 117.5
DW 50.000 usec
DE 6.50 usec
TE 297.2 K
D1 2.00000000 sec
TD0 1
SFO1 500.1330883 MHz
NUC1 1H
P1 10.00 usec
PLW1 23.50000000 W
F2 - Processing parameters
SI 65536
SF 500.1300120 MHz
WDW EM
SSB 0
LB 0.30 Hz
GB 0
PC 1.00

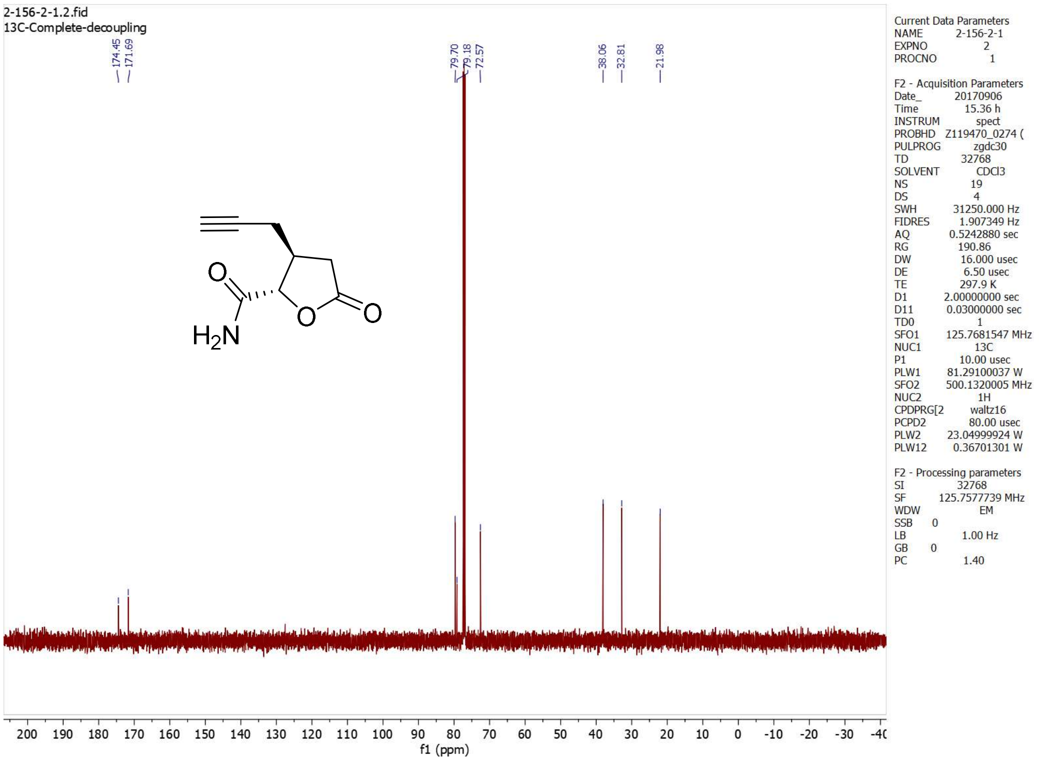

2-156-2-1.2.fid
13C-Complete-decoupling
174.45
171.69
79.70
79.18
72.57
38.06
32.81
21.98
H2N
O
O
O
f1 (ppm)
Current Data Parameters
NAME 2-156-2-1
EXPNO 2
PROCNO 1
F2 - Acquisition Parameters
Date_ 20170906
Time 15.36 h
INSTRUM spect
PROBHD Z119470_0274 (
PULPROG zgdc30
TD 32768
SOLVENT CDCl3
NS 19
DS 4
SWH 31250.000 Hz
FIDRES 1.907349 Hz
AQ 0.5242880 sec
RG 190.86
DW 16.000 usec
DE 6.50 usec
TE 297.9 K
D1 2.00000000 sec
D11 0.03000000 sec
TD0 1
SFO1 125.7681547 MHz
NUC1 13C
P1 10.00 usec
PLW1 81.29100037 W
SFO2 500.1320005 MHz
NUC2 1H
CPDPRG[2 waltz16
PCPD2 80.00 usec
PLW2 23.04999924 W
PLW12 0.36701301 W
F2 - Processing parameters
SI 32768
SF 125.7577739 MHz
WDW EM
SSB 0
LB 1.00 Hz
GB 0
PC 1.40

**4.13-2**

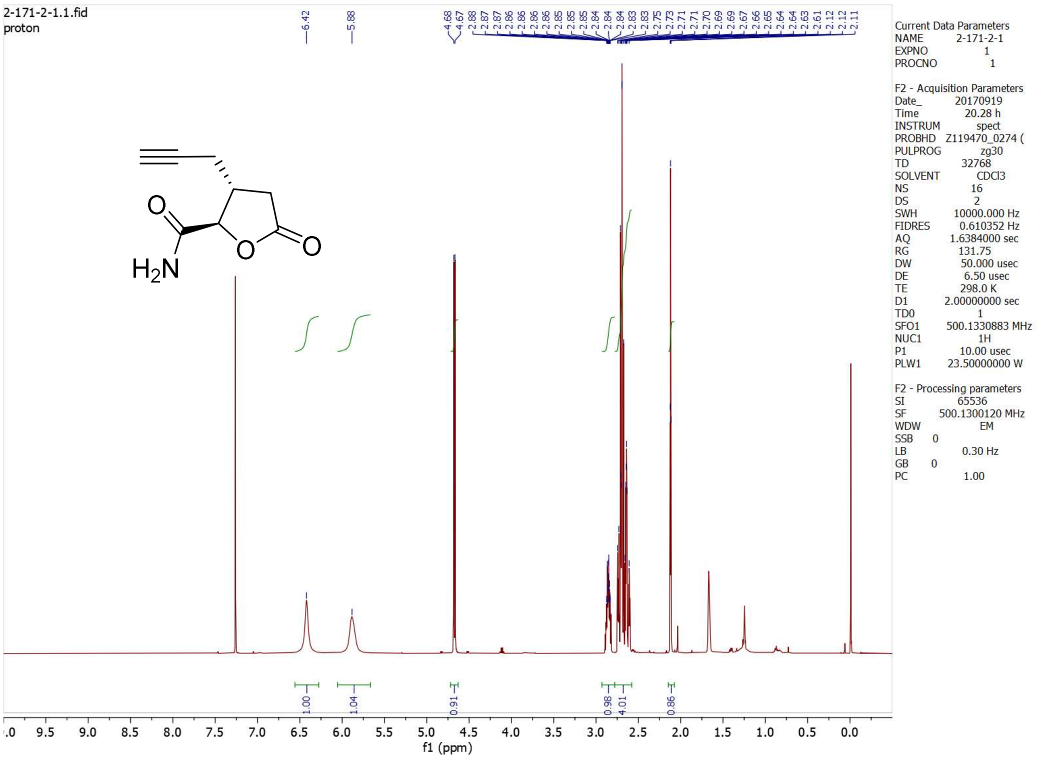
2-171-2-1.1.fid
proton
6.42
5.88
4.68
4.67
H2N
O
O
O
1.00
1.04
0.91
0.98
4.01
0.86
f1 (ppm)
Current Data Parameters
NAME 2-171-2-1
EXPNO 1
PROCNO 1
F2 - Acquisition Parameters
Date_ 20170919
Time 20.28 h
INSTRUM spect
PROBHD Z119470_0274 (
PULPROG zg30
TD 32768
SOLVENT CDCl3
NS 16
DS 2
SWH 10000.000 Hz
FIDRES 0.610352 Hz
AQ 1.6384000 sec
RG 131.75
DW 50.000 usec
DE 6.50 usec
TE 298.0 K
D1 2.00000000 sec
TD0 1
SFO1 500.1330883 MHz
NUC1 1H
P1 10.00 usec
PLW1 23.50000000 W
F2 - Processing parameters
SI 65536
SF 500.1300120 MHz
WDW EM
SSB 0
LB 0.30 Hz
GB 0
PC 1.00

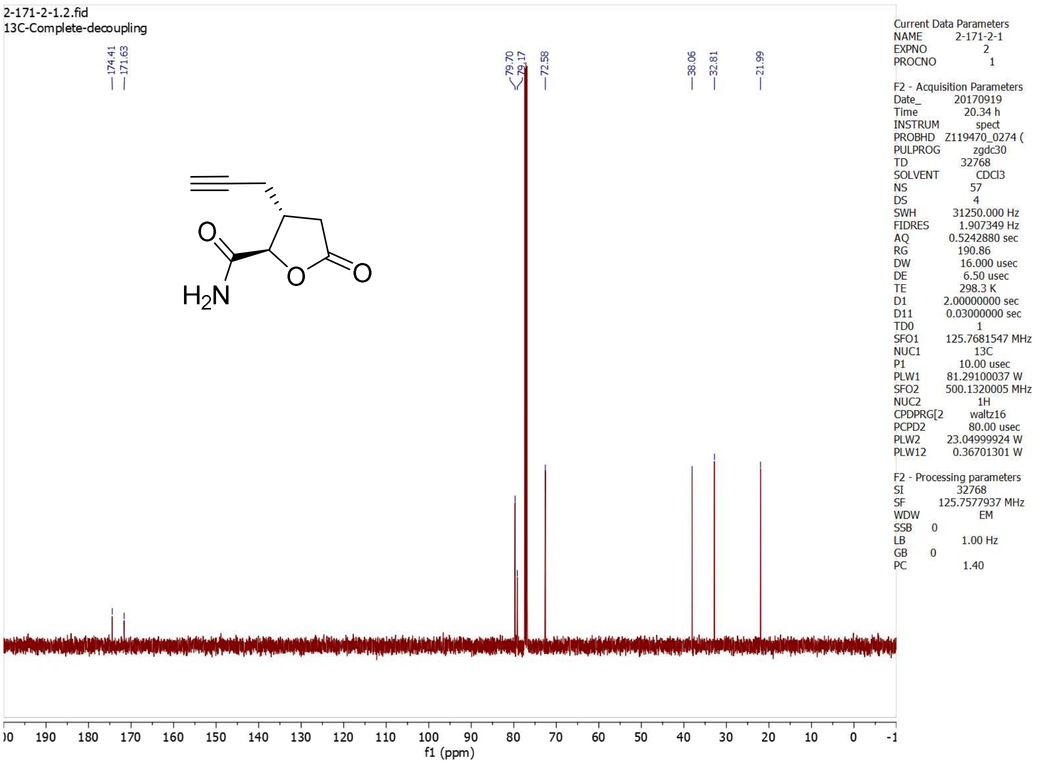
2-171-2-1.2.fid
13C-Complete-decoupling
174.41
171.63
79.70
79.17
72.58
38.06
32.81
21.99
H2N
O
O
O
f1 (ppm)
Current Data Parameters
NAME 2-171-2-1
EXPNO 2
PROCNO 1
F2 - Acquisition Parameters
Date_ 20170919
Time 20.34 h
INSTRUM spect
PROBHD Z119470_0274 (
PULPROG zgdc30
TD 32768
SOLVENT CDCl3
NS 57
DS 4
SWH 31250.000 Hz
FIDRES 1.907349 Hz
AQ 0.5242880 sec
RG 190.86
DW 16.000 usec
DE 6.50 usec
TE 298.3 K
D1 2.00000000 sec
D11 0.03000000 sec
TD0 1
SFO1 125.7681547 MHz
NUC1 13C
P1 10.00 usec
PLW1 81.29100037 W
SFO2 500.1320005 MHz
NUC2 1H
CPDPRG[2 waltz16
PCPD2 80.00 usec
PLW2 23.04999924 W
PLW12 0.36701301 W
F2 - Processing parameters
SI 32768
SF 125.7577937 MHz
WDW EM
SSB 0
LB 1.00 Hz
GB 0
PC 1.40

**4.14-1**

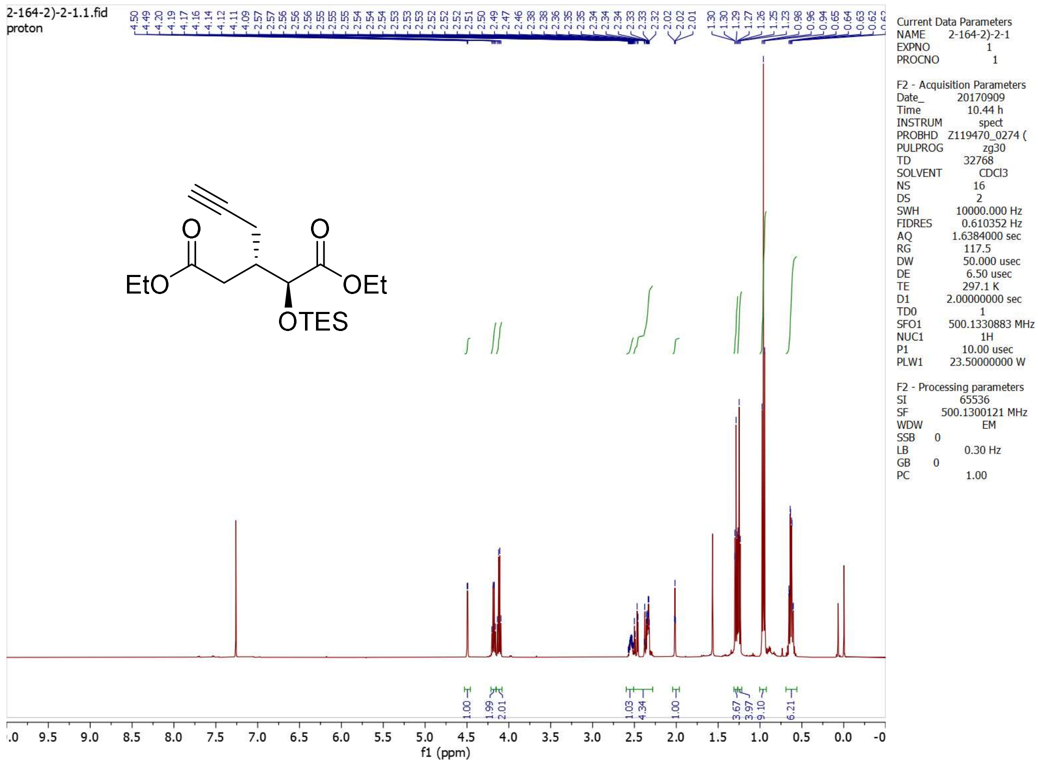

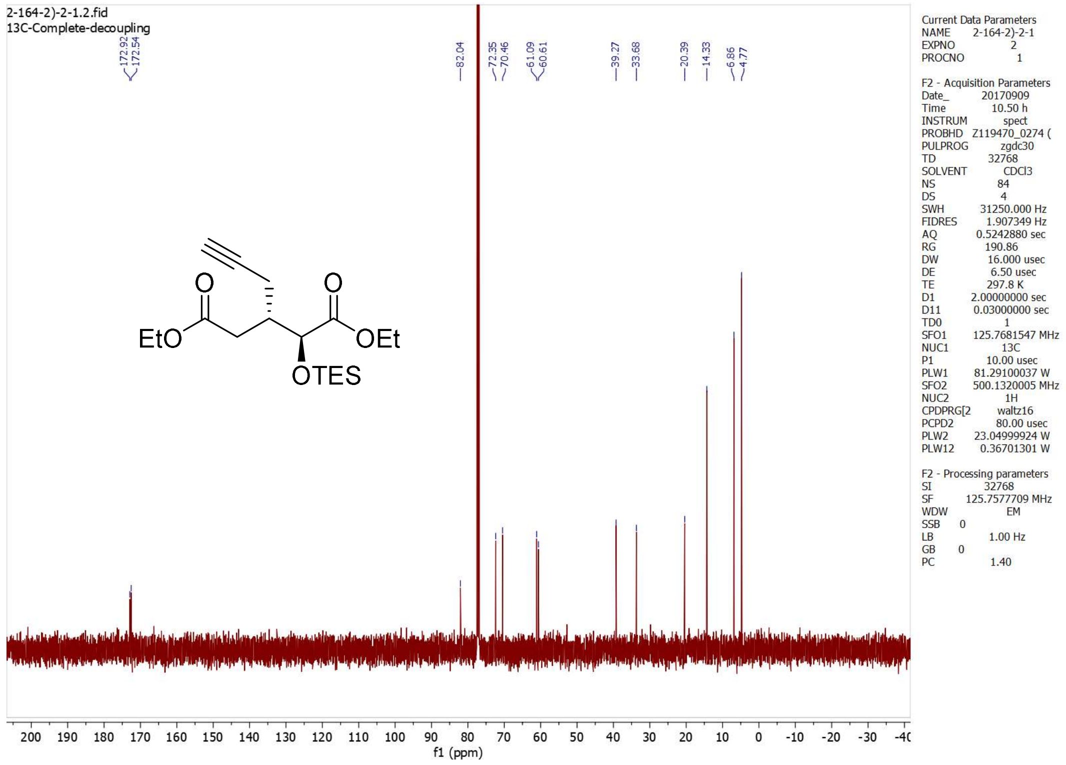

**4-1 (4*SS*)**

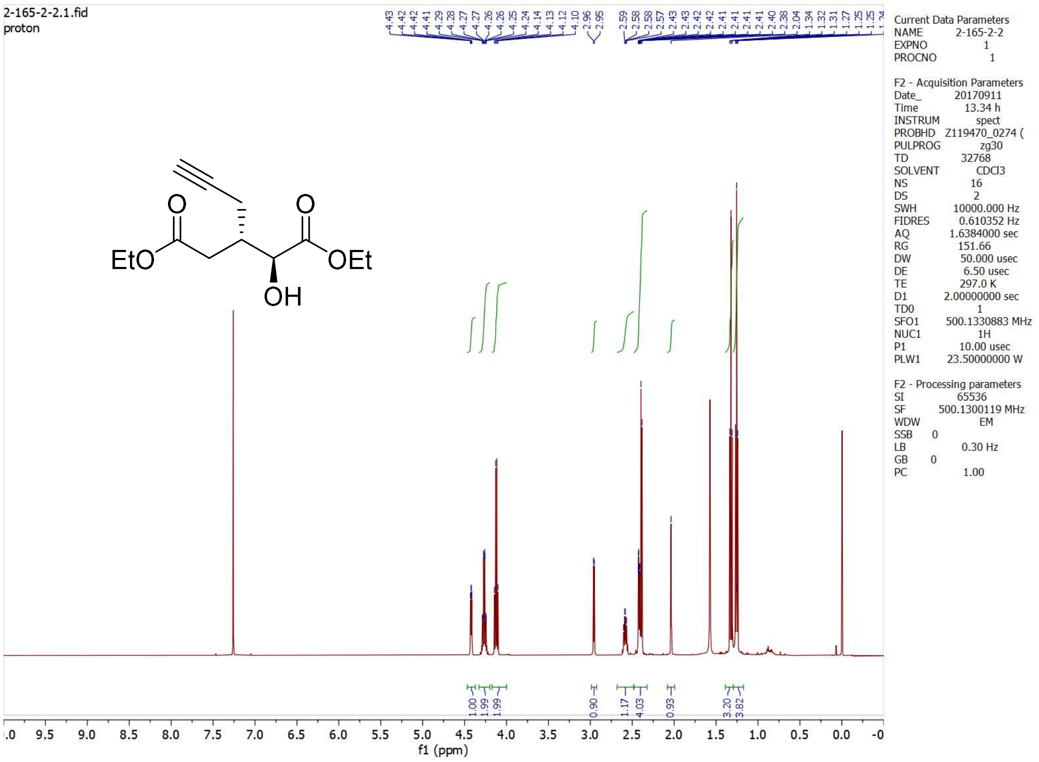

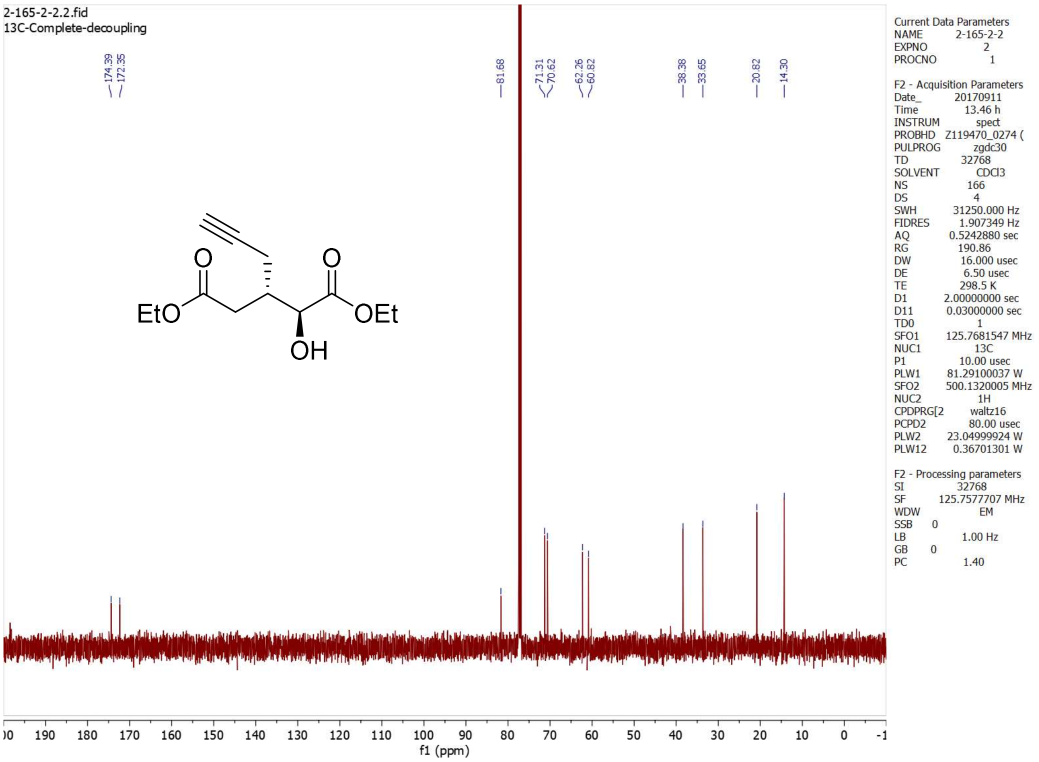

**4-2 (4*RR*)**

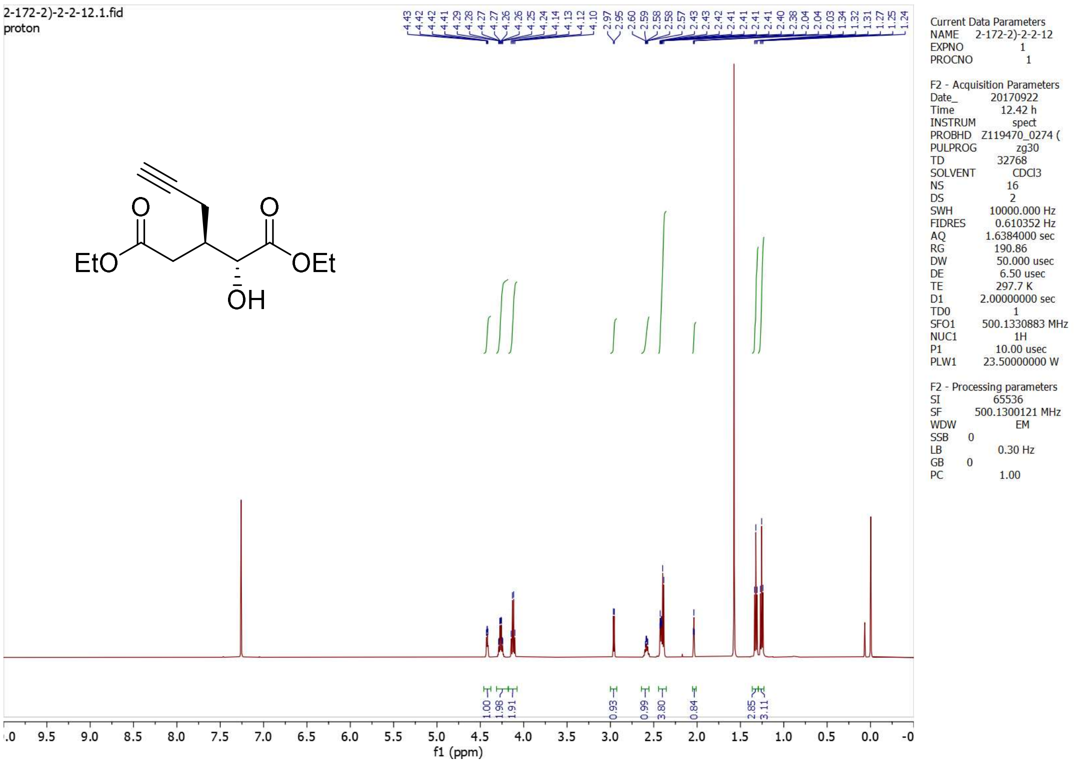

2-172-2)-2-2-12.1.fid
proton
EtO
OEt
OH
f1 (ppm)
Current Data Parameters
NAME 2-172-2)-2-2-12
EXPNO 1
PROCNO 1
F2 - Acquisition Parameters
Date_ 20170922
Time 12.42 h
INSTRUM spect
PROBHD Z119470_0274 (
PULPROG zg30
TD 32768
SOLVENT CDCl3
NS 16
DS 2
SWH 10000.000 Hz
FIDRES 0.610352 Hz
AQ 1.6384000 sec
RG 190.86
DW 50.000 usec
DE 6.50 usec
TE 297.7 K
D1 2.00000000 sec
TD0 1
SFO1 500.1330883 MHz
NUC1 1H
P1 10.00 usec
PLW1 23.50000000 W
F2 - Processing parameters
SI 65536
SF 500.1300121 MHz
WDW EM
SSB 0
LB 0.30 Hz
GB 0
PC 1.00

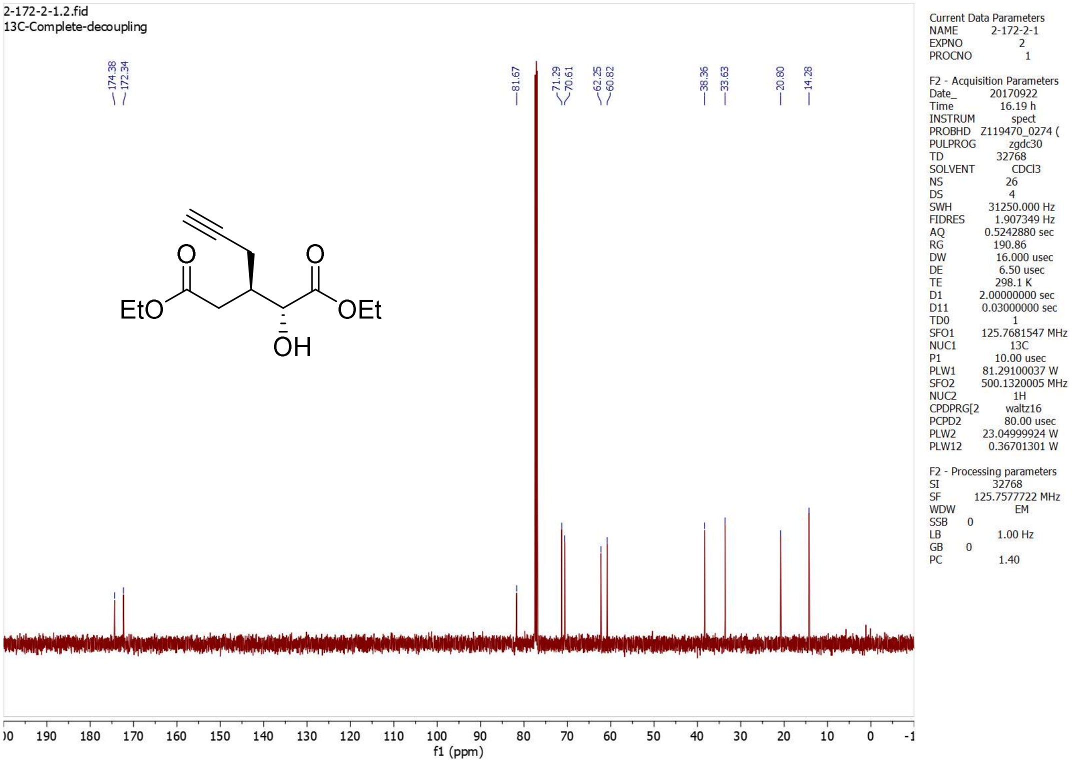

2-172-2-1.2.fid
13C-Complete-decoupling
174.38
172.34
81.67
71.29
70.61
62.25
60.82
38.36
33.63
20.80
14.28
EtO
OEt
OH
f1 (ppm)
Current Data Parameters
NAME 2-172-2-1
EXPNO 2
PROCNO 1
F2 - Acquisition Parameters
Date_ 20170922
Time 16.19 h
INSTRUM spect
PROBHD Z119470_0274 (
PULPROG zgdc30
TD 32768
SOLVENT CDCl3
NS 26
DS 4
SWH 31250.000 Hz
FIDRES 1.907349 Hz
AQ 0.5242880 sec
RG 190.86
DW 16.000 usec
DE 6.50 usec
TE 298.1 K
D1 2.00000000 sec
D11 0.03000000 sec
TD0 1
SFO1 125.7681547 MHz
NUC1 13C
P1 10.00 usec
PLW1 81.29100037 W
SFO2 500.1320005 MHz
NUC2 1H
CPDPRG[2 waltz16
PCPD2 80.00 usec
PLW2 23.04999924 W
PLW12 0.36701301 W
F2 - Processing parameters
SI 32768
SF 125.7577722 MHz
WDW EM
SSB 0
LB 1.00 Hz
GB 0
PC 1.40

**4-3/4 (4*RS*/*SR*)**

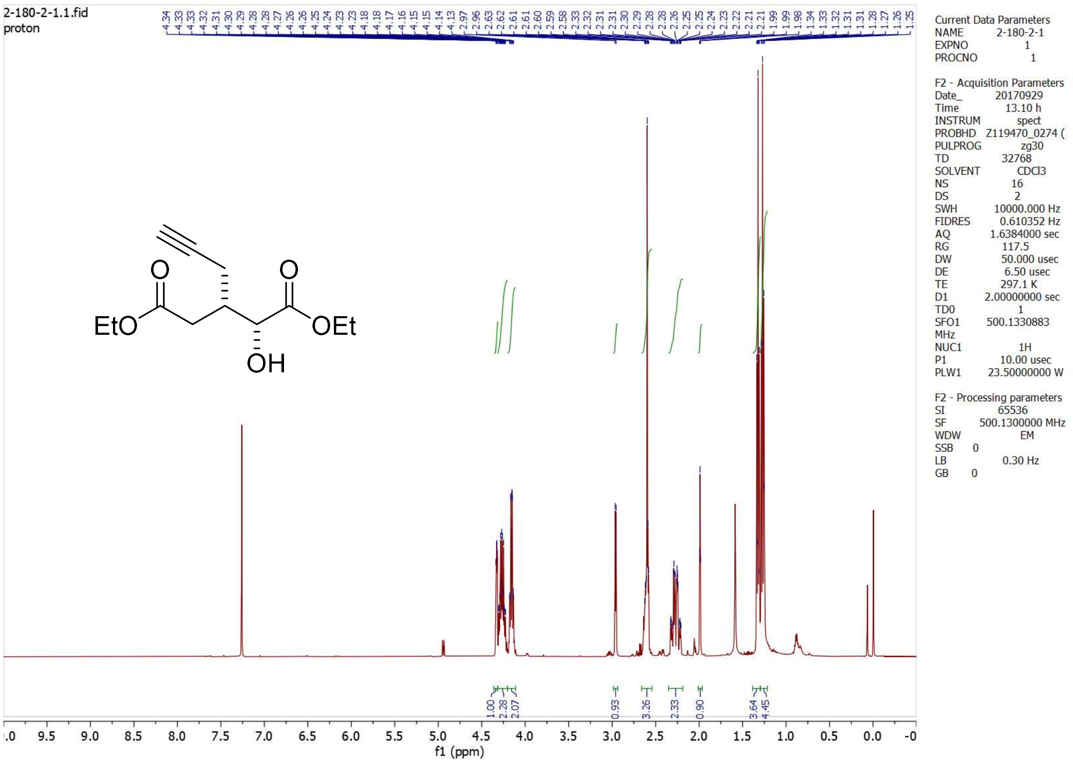

2-180-2-1.1.fid
proton
EtO
OEt
OH
f1 (ppm)
Current Data Parameters
NAME 2-180-2-1
EXPNO 1
PROCNO 1
F2 - Acquisition Parameters
Date_ 20170929
Time 13.10 h
INSTRUM spect
PROBHD Z119470_0274 (
PULPROG zg30
TD 32768
SOLVENT CDCl3
NS 16
DS 2
SWH 10000.000 Hz
FIDRES 0.610352 Hz
AQ 1.6384000 sec
RG 117.5
DW 50.000 usec
DE 6.50 usec
TE 297.1 K
D1 2.00000000 sec
TD0 1
SFO1 500.1330883 MHz
NUC1 1H
P1 10.00 usec
PLW1 23.50000000 W
F2 - Processing parameters
SI 65536
SF 500.1300000 MHz
WDW EM
SSB 0
LB 0.30 Hz
GB 0

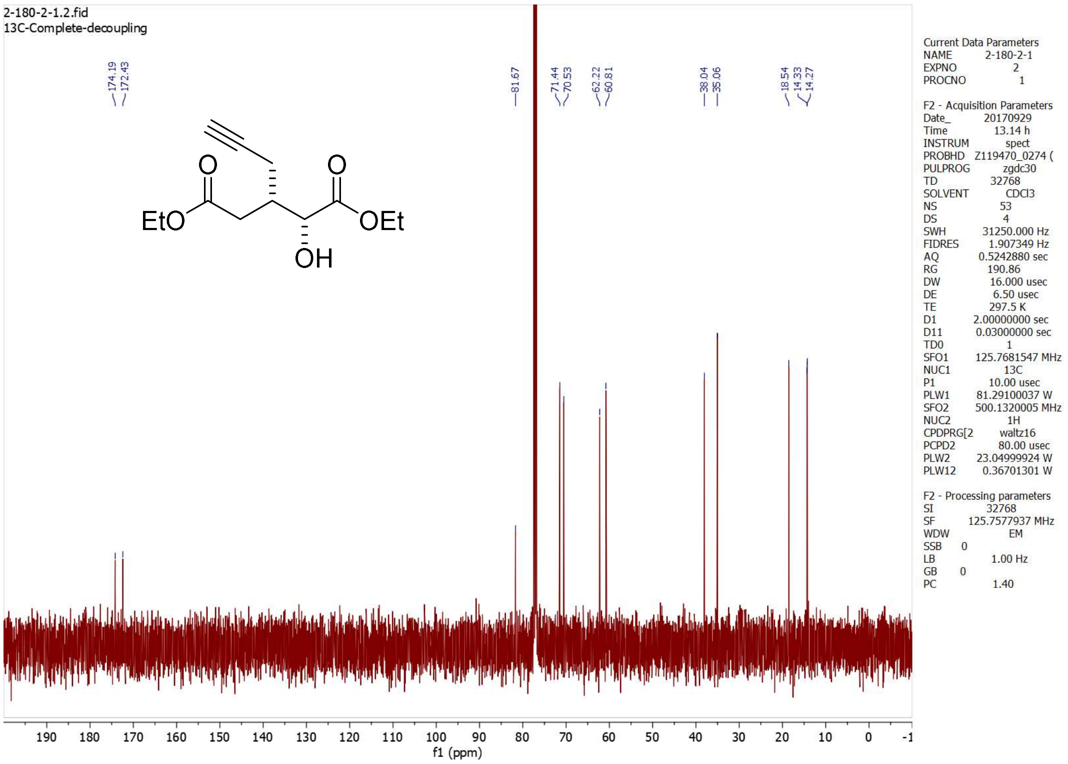

2-180-2-1.2.fid
13C-Complete-decoupling
174.19
172.43
81.67
71.44
70.53
62.22
60.81
38.04
35.06
18.54
14.33
14.27
EtO
OEt
OH
f1 (ppm)
Current Data Parameters
NAME 2-180-2-1
EXPNO 2
PROCNO 1
F2 - Acquisition Parameters
Date_ 20170929
Time 13.14 h
INSTRUM spect
PROBHD Z119470_0274 (
PULPROG zgdc30
TD 32768
SOLVENT CDCl3
NS 53
DS 4
SWH 31250.000 Hz
FIDRES 1.907349 Hz
AQ 0.5242880 sec
RG 190.86
DW 16.000 usec
DE 6.50 usec
TE 297.5 K
D1 2.00000000 sec
D11 0.03000000 sec
TD0 1
SFO1 125.7681547 MHz
NUC1 13C
P1 10.00 usec
PLW1 81.29100037 W
SFO2 500.1320005 MHz
NUC2 1H
CPDPRG[2 waltz16
PCPD2 80.00 usec
PLW2 23.04999924 W
PLW12 0.36701301 W
F2 - Processing parameters
SI 32768
SF 125.7577937 MHz
WDW EM
SSB 0
LB 1.00 Hz
GB 0
PC 1.40

**5.2*S***

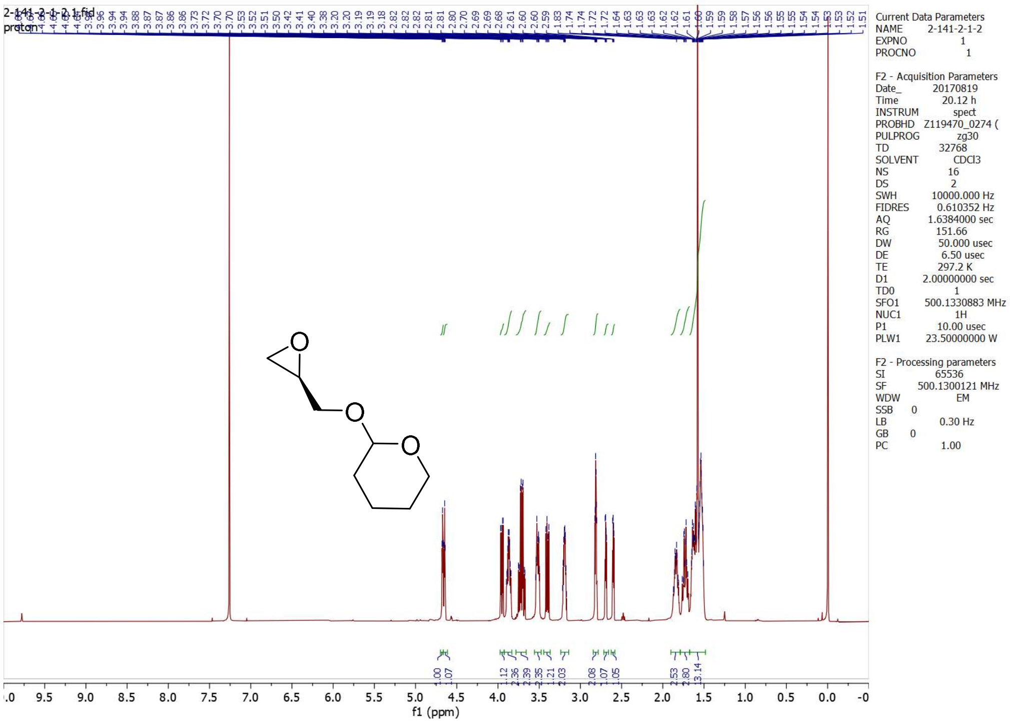

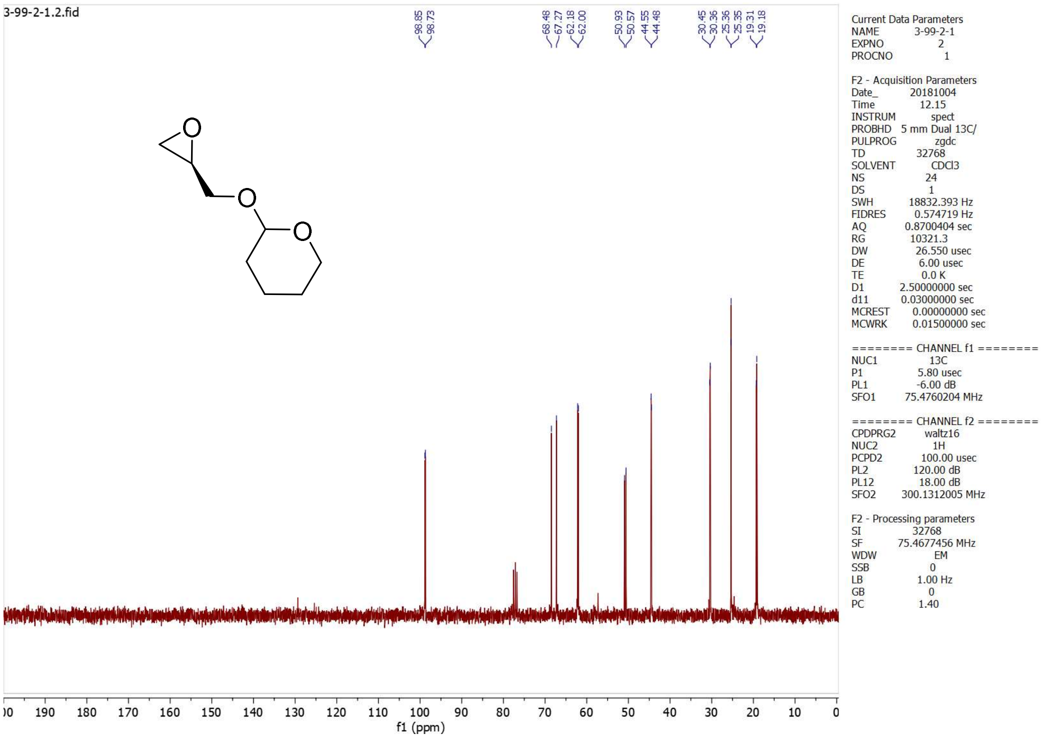

## 5.2*R*

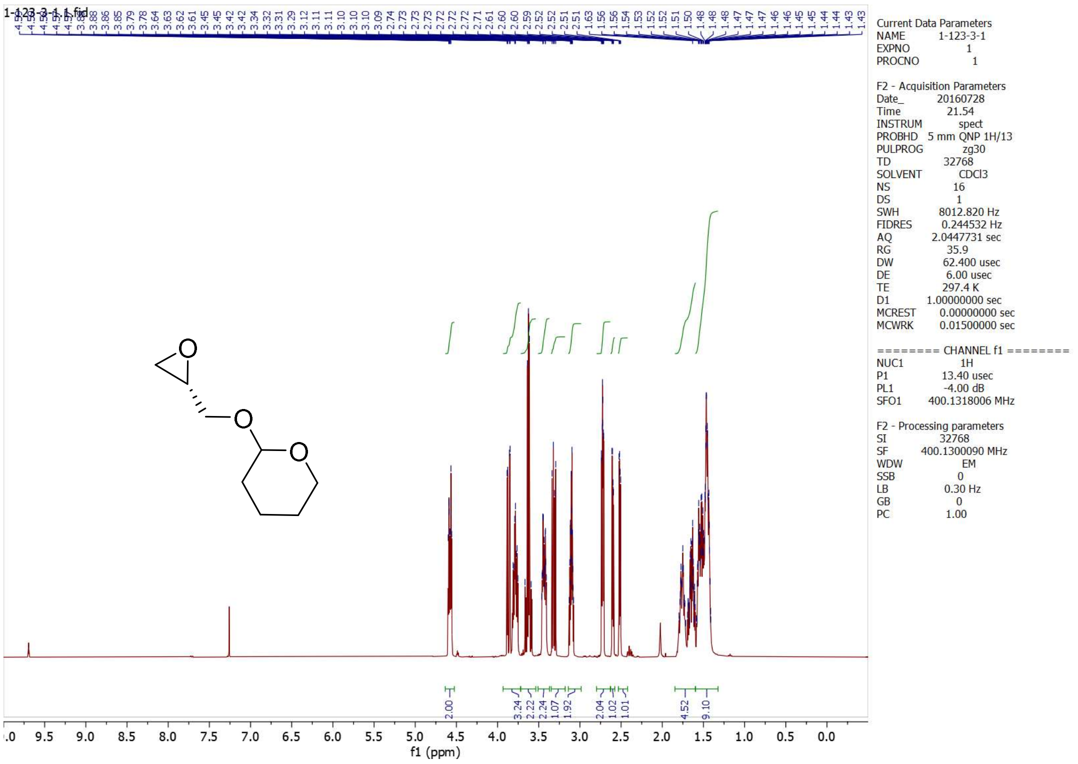
1-123-3-1.1.fid
2.00
3.24
2.22
2.24
1.07
1.92
2.04
1.02
1.01
4.52
9.10
f1 (ppm)
Current Data Parameters
NAME 1-123-3-1
EXPNO 1
PROCNO 1
F2 - Acquisition Parameters
Date_ 20160728
Time 21.54
INSTRUM spect
PROBHD 5 mm QNP 1H/13
PULPROG zg30
TD 32768
SOLVENT CDCl3
NS 16
DS 1
SWH 8012.820 Hz
FIDRES 0.244532 Hz
AQ 2.0447731 sec
RG 35.9
DW 62.400 usec
DE 6.00 usec
TE 297.4 K
D1 1.00000000 sec
MCREST 0.00000000 sec
MCWRK 0.01500000 sec
======== CHANNEL f1 ========
NUC1 1H
P1 13.40 usec
PL1 -4.00 dB
SFO1 400.1318006 MHz
F2 - Processing parameters
SI 32768
SF 400.1300090 MHz
WDW EM
SSB 0
LB 0.30 Hz
GB 0
PC 1.00

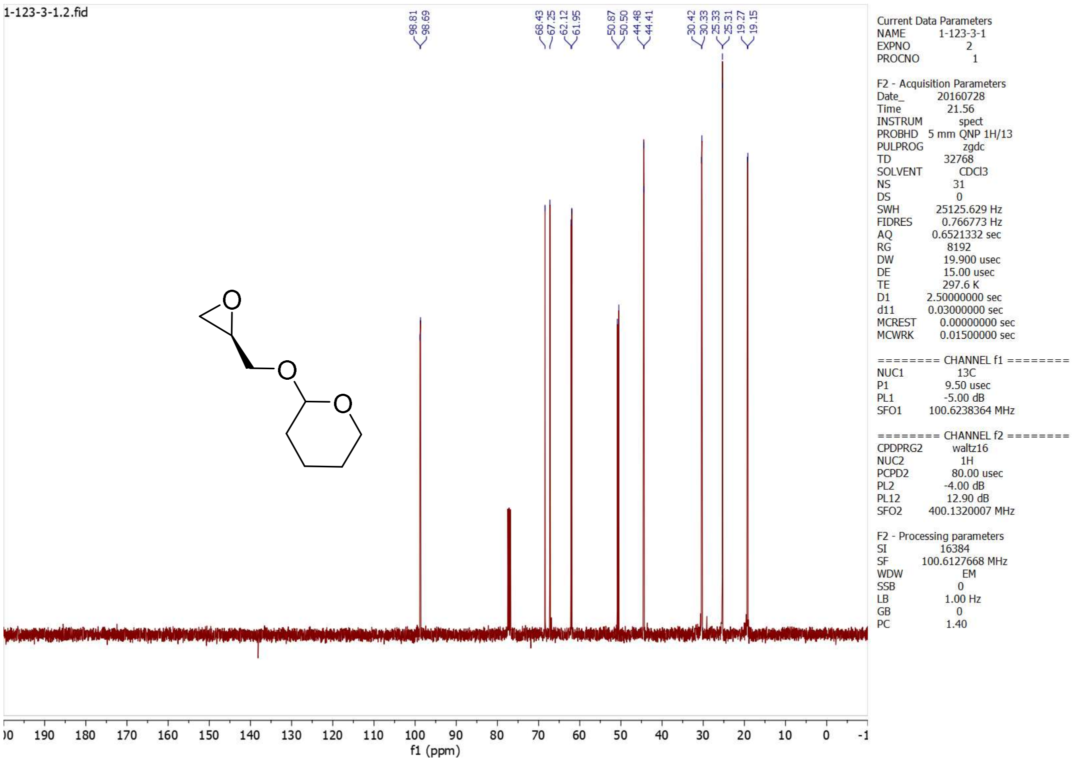
1-123-3-1.2.fid
98.81
98.69
68.43
67.25
62.12
61.95
50.87
50.50
44.48
44.41
30.42
30.33
25.33
25.31
19.27
19.15
f1 (ppm)
Current Data Parameters
NAME 1-123-3-1
EXPNO 2
PROCNO 1
F2 - Acquisition Parameters
Date_ 20160728
Time 21.56
INSTRUM spect
PROBHD 5 mm QNP 1H/13
PULPROG zgdc
TD 32768
SOLVENT CDCl3
NS 31
DS 0
SWH 25125.629 Hz
FIDRES 0.766773 Hz
AQ 0.6521332 sec
RG 8192
DW 19.900 usec
DE 15.00 usec
TE 297.6 K
D1 2.50000000 sec
d11 0.03000000 sec
MCREST 0.00000000 sec
MCWRK 0.01500000 sec
======== CHANNEL f1 ========
NUC1 13C
P1 9.50 usec
PL1 -5.00 dB
SFO1 100.6238364 MHz
======== CHANNEL f2 ========
CPDPRG2 waltz16
NUC2 1H
PCPD2 80.00 usec
PL2 -4.00 dB
PL12 12.90 dB
SFO2 400.1320007 MHz
F2 - Processing parameters
SI 16384
SF 100.6127668 MHz
WDW EM
SSB 0
LB 1.00 Hz
GB 0
PC 1.40

**5.3S**

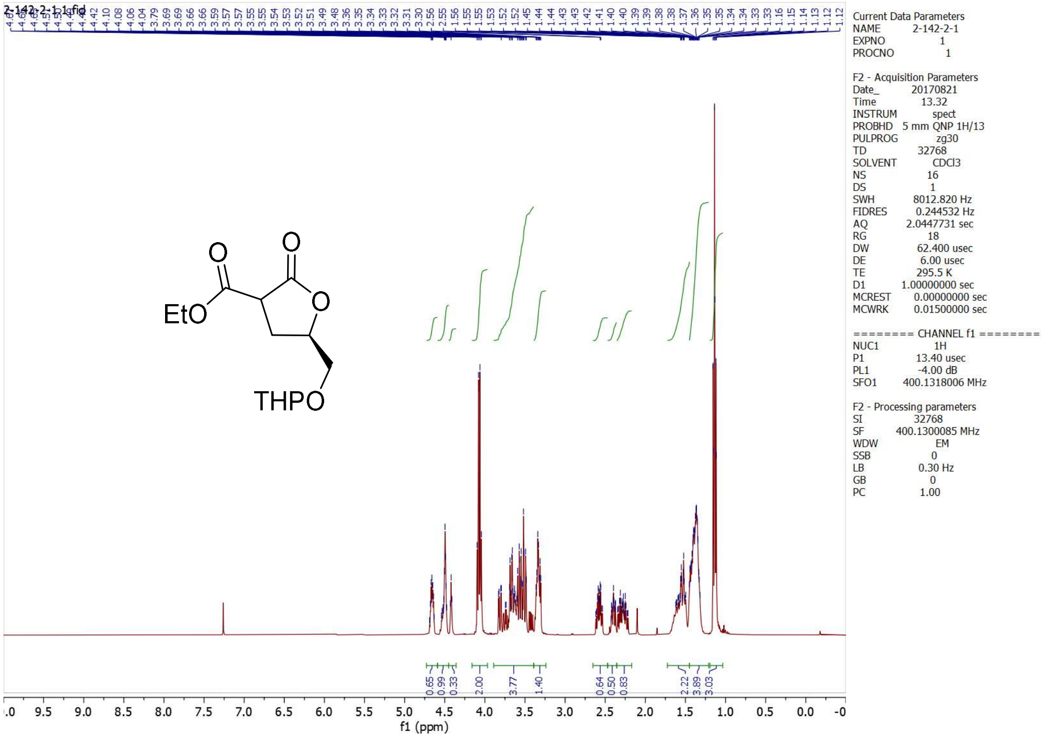

2-142-2-1.1.fid
EtO
O
O
O
THPO
Current Data Parameters
NAME 2-142-2-1
EXPNO 1
PROCNO 1
F2 - Acquisition Parameters
Date_ 20170821
Time 13.32
INSTRUM spect
PROBHD 5 mm QNP 1H/13
PULPROG zg30
TD 32768
SOLVENT CDCl3
NS 16
DS 1
SWH 8012.820 Hz
FIDRES 0.244532 Hz
AQ 2.0447731 sec
RG 18
DW 62.400 usec
DE 6.00 usec
TE 295.5 K
D1 1.00000000 sec
MCREST 0.00000000 sec
MCWRK 0.01500000 sec
======== CHANNEL f1 ========
NUC1 1H
P1 13.40 usec
PL1 -4.00 dB
SFO1 400.1318006 MHz
F2 - Processing parameters
SI 32768
SF 400.1300085 MHz
WDW EM
SSB 0
LB 0.30 Hz
GB 0
PC 1.00
0.65
0.99
0.33
2.00
3.77
1.40
0.64
0.50
0.83
2.22
3.89
3.03
f1 (ppm)

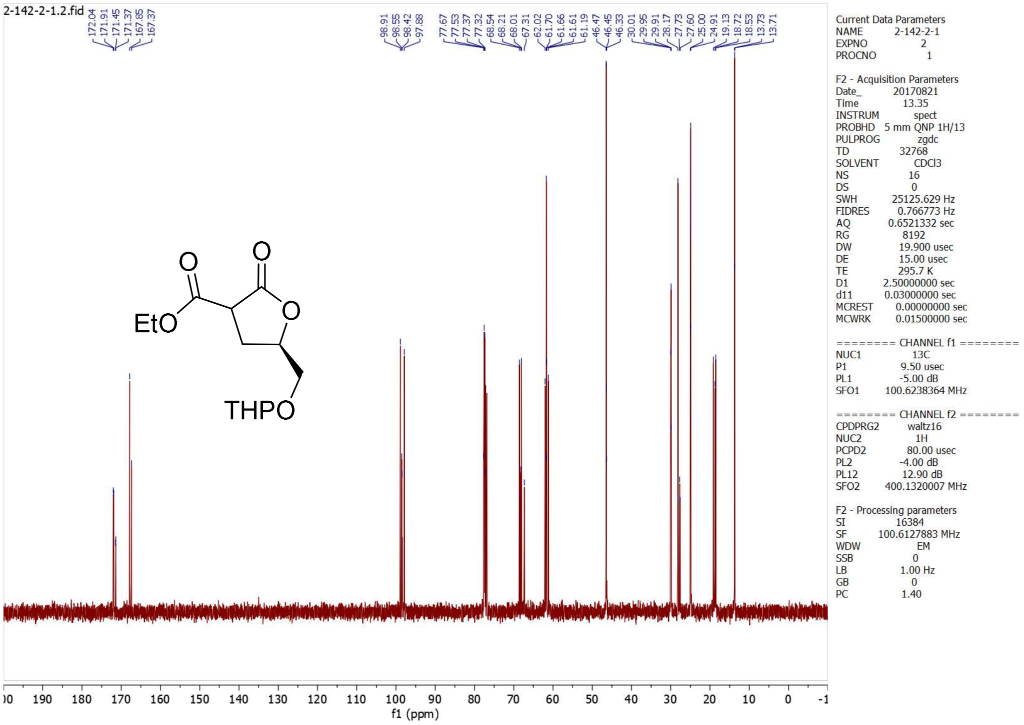

2-142-2-1.2.fid
172.04
171.91
171.45
171.37
167.85
167.37
98.91
98.55
98.42
97.88
77.67
77.53
77.37
77.32
68.54
68.21
68.01
67.31
62.02
61.70
61.66
61.61
61.19
46.47
46.45
46.33
30.01
29.95
29.91
28.17
27.73
27.60
25.00
24.91
19.13
18.72
18.53
13.73
13.71
EtO
O
O
O
THPO
Current Data Parameters
NAME 2-142-2-1
EXPNO 2
PROCNO 1
F2 - Acquisition Parameters
Date_ 20170821
Time 13.35
INSTRUM spect
PROBHD 5 mm QNP 1H/13
PULPROG zgdc
TD 32768
SOLVENT CDCl3
NS 16
DS 0
SWH 25125.629 Hz
FIDRES 0.766773 Hz
AQ 0.6521332 sec
RG 8192
DW 19.900 usec
DE 15.00 usec
TE 295.7 K
D1 2.50000000 sec
d11 0.03000000 sec
MCREST 0.00000000 sec
MCWRK 0.01500000 sec
======== CHANNEL f1 ========
NUC1 13C
P1 9.50 usec
PL1 -5.00 dB
SFO1 100.6238364 MHz
======== CHANNEL f2 ========
CPDPRG2 waltz16
NUC2 1H
PCPD2 80.00 usec
PL2 -4.00 dB
PL12 12.90 dB
SFO2 400.1320007 MHz
F2 - Processing parameters
SI 16384
SF 100.6127883 MHz
WDW EM
SSB 0
LB 1.00 Hz
GB 0
PC 1.40
f1 (ppm)

## 5.3*R*

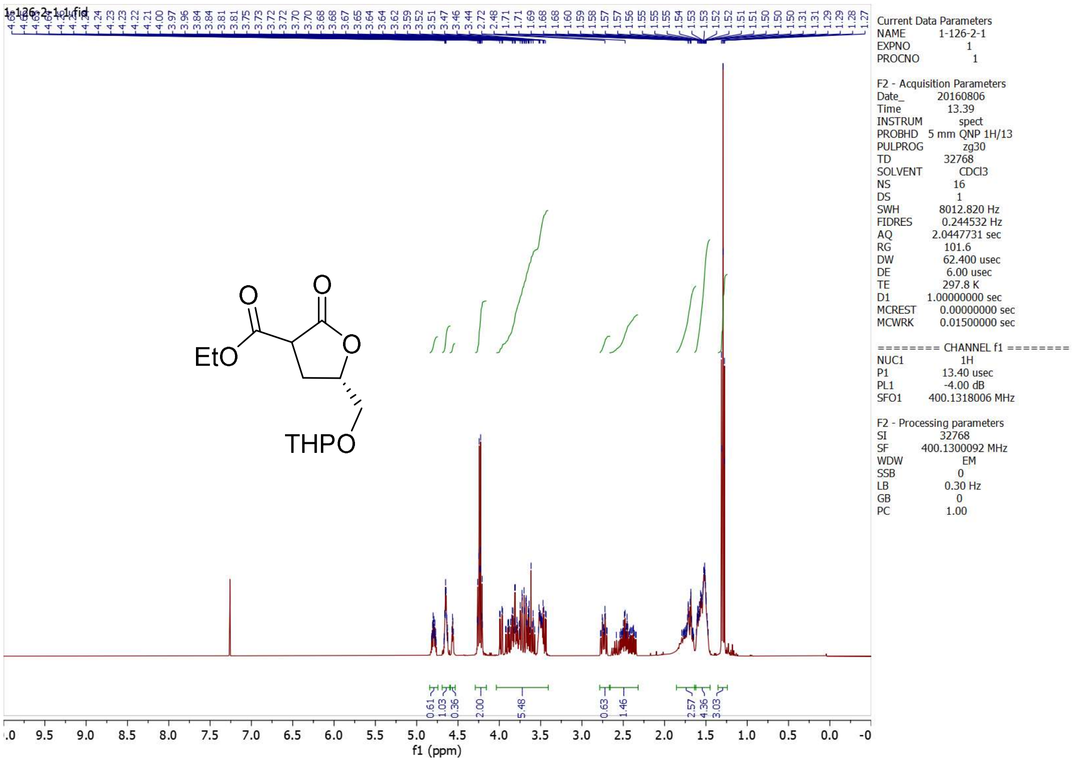
1-126-2-1.1.fid
EtO
O
O
O
THPO
Current Data Parameters
NAME 1-126-2-1
EXPNO 1
PROCNO 1
F2 - Acquisition Parameters
Date_ 20160806
Time 13.39
INSTRUM spect
PROBHD 5 mm QNP 1H/13
PULPROG zg30
TD 32768
SOLVENT CDCl3
NS 16
DS 1
SWH 8012.820 Hz
FIDRES 0.244532 Hz
AQ 2.0447731 sec
RG 101.6
DW 62.400 usec
DE 6.00 usec
TE 297.8 K
D1 1.00000000 sec
MCREST 0.00000000 sec
MCWRK 0.01500000 sec
======== CHANNEL f1 ========
NUC1 1H
P1 13.40 usec
PL1 -4.00 dB
SFO1 400.1318006 MHz
F2 - Processing parameters
SI 32768
SF 400.1300092 MHz
WDW EM
SSB 0
LB 0.30 Hz
GB 0
PC 1.00
f1 (ppm)

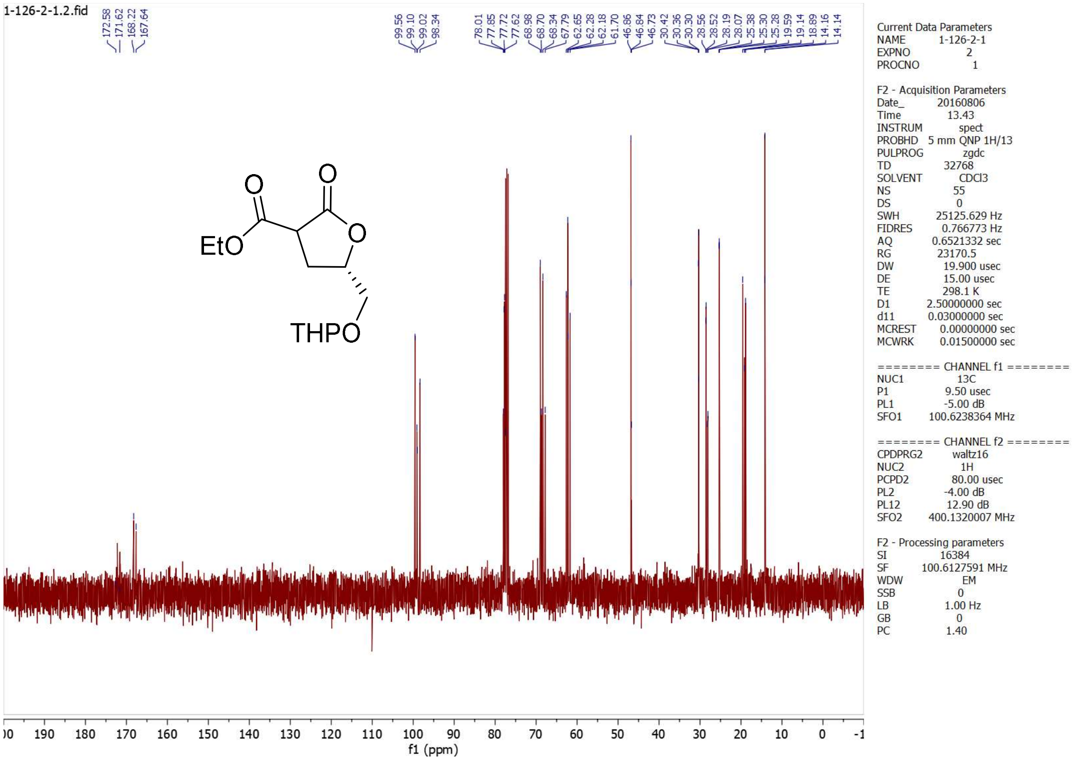
1-126-2-1.2.fid
EtO
O
O
O
THPO
Current Data Parameters
NAME 1-126-2-1
EXPNO 2
PROCNO 1
F2 - Acquisition Parameters
Date_ 20160806
Time 13.43
INSTRUM spect
PROBHD 5 mm QNP 1H/13
PULPROG zgdc
TD 32768
SOLVENT CDCl3
NS 55
DS 0
SWH 25125.629 Hz
FIDRES 0.766773 Hz
AQ 0.6521332 sec
RG 23170.5
DW 19.900 usec
DE 15.00 usec
TE 298.1 K
D1 2.50000000 sec
d11 0.03000000 sec
MCREST 0.00000000 sec
MCWRK 0.01500000 sec
======== CHANNEL f1 ========
NUC1 13C
P1 9.50 usec
PL1 -5.00 dB
SFO1 100.6238364 MHz
======== CHANNEL f2 ========
CPDPRG2 waltz16
NUC2 1H
PCPD2 80.00 usec
PL2 -4.00 dB
PL12 12.90 dB
SFO2 400.1320007 MHz
F2 - Processing parameters
SI 16384
SF 100.6127591 MHz
WDW EM
SSB 0
LB 1.00 Hz
GB 0
PC 1.40
f1 (ppm)

**5.4*S***

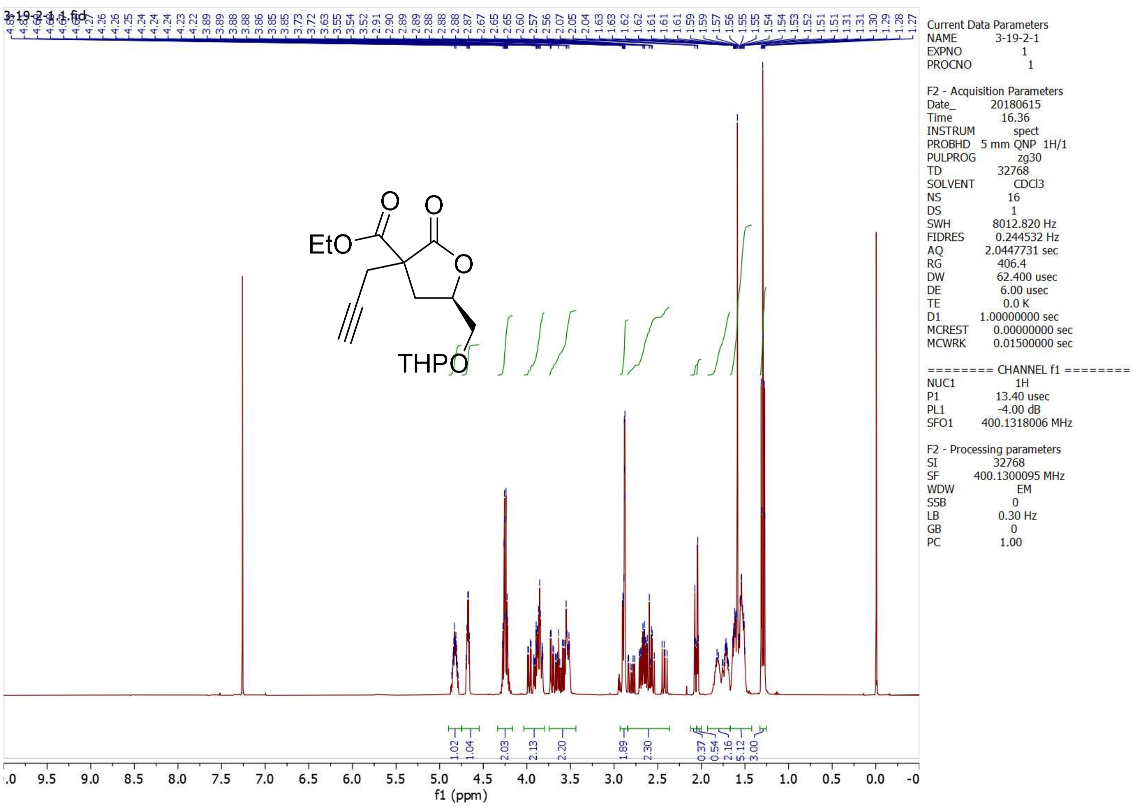

EtO
O
O
O
THPO
Current Data Parameters
NAME 3-19-2-1
EXPNO 1
PROCNO 1
F2 - Acquisition Parameters
Date_ 20180615
Time 16.36
INSTRUM spect
PROBHD 5 mm QNP 1H/1
PULPROG zg30
TD 32768
SOLVENT CDCl3
NS 16
DS 1
SWH 8012.820 Hz
FIDRES 0.244532 Hz
AQ 2.0447731 sec
RG 406.4
DW 62.400 usec
DE 6.00 usec
TE 0.0 K
D1 1.00000000 sec
MCREST 0.00000000 sec
MCWRK 0.01500000 sec
======== CHANNEL f1 ========
NUC1 1H
P1 13.40 usec
PL1 -4.00 dB
SFO1 400.1318006 MHz
F2 - Processing parameters
SI 32768
SF 400.1300095 MHz
WDW EM
SSB 0
LB 0.30 Hz
GB 0
PC 1.00
f1 (ppm)

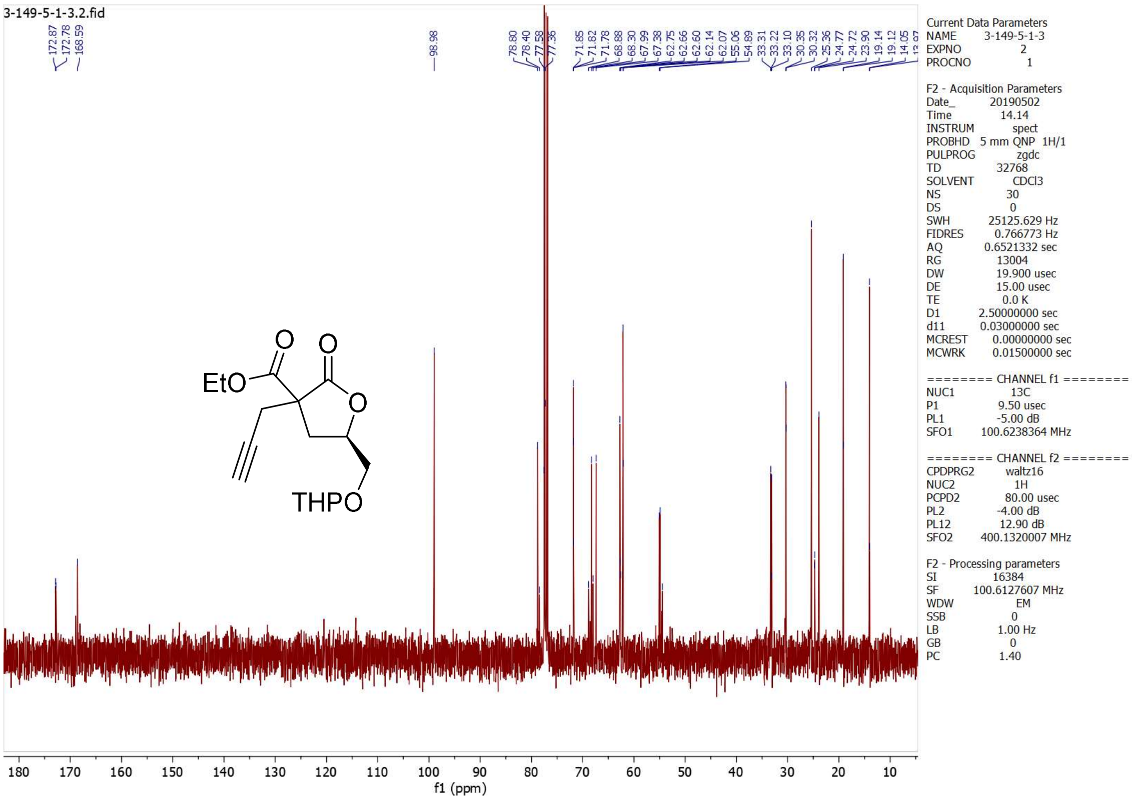
3-149-5-1-3.2.fid
172.87
172.78
168.59
98.98
78.80
78.40
77.58
77.36
71.85
71.82
71.78
68.88
68.30
67.99
67.38
62.75
62.66
62.60
62.14
62.07
55.06
54.89
33.31
33.22
33.10
30.35
30.32
25.36
24.77
24.72
23.90
19.14
19.12
14.05
EtO
O
O
O
THPO
180 170 160 150 140 130 120 110 100 90 80 70 60 50 40 30 20 10
f1 (ppm)
Current Data Parameters
NAME 3-149-5-1-3
EXPNO 2
PROCNO 1
F2 - Acquisition Parameters
Date_ 20190502
Time 14.14
INSTRUM spect
PROBHD 5 mm QNP 1H/1
PULPROG zgdc
TD 32768
SOLVENT CDCl3
NS 30
DS 0
SWH 25125.629 Hz
FIDRES 0.766773 Hz
AQ 0.6521332 sec
RG 13004
DW 19.900 usec
DE 15.00 usec
TE 0.0 K
D1 2.50000000 sec
d11 0.03000000 sec
MCREST 0.00000000 sec
MCWRK 0.01500000 sec
======== CHANNEL f1 ========
NUC1 13C
P1 9.50 usec
PL1 -5.00 dB
SFO1 100.6238364 MHz
======== CHANNEL f2 ========
CPDPRG2 waltz16
NUC2 1H
PCPD2 80.00 usec
PL2 -4.00 dB
PL12 12.90 dB
SFO2 400.1320007 MHz
F2 - Processing parameters
SI 16384
SF 100.6127607 MHz
WDW EM
SSB 0
LB 1.00 Hz
GB 0
PC 1.40

**5.4*R***

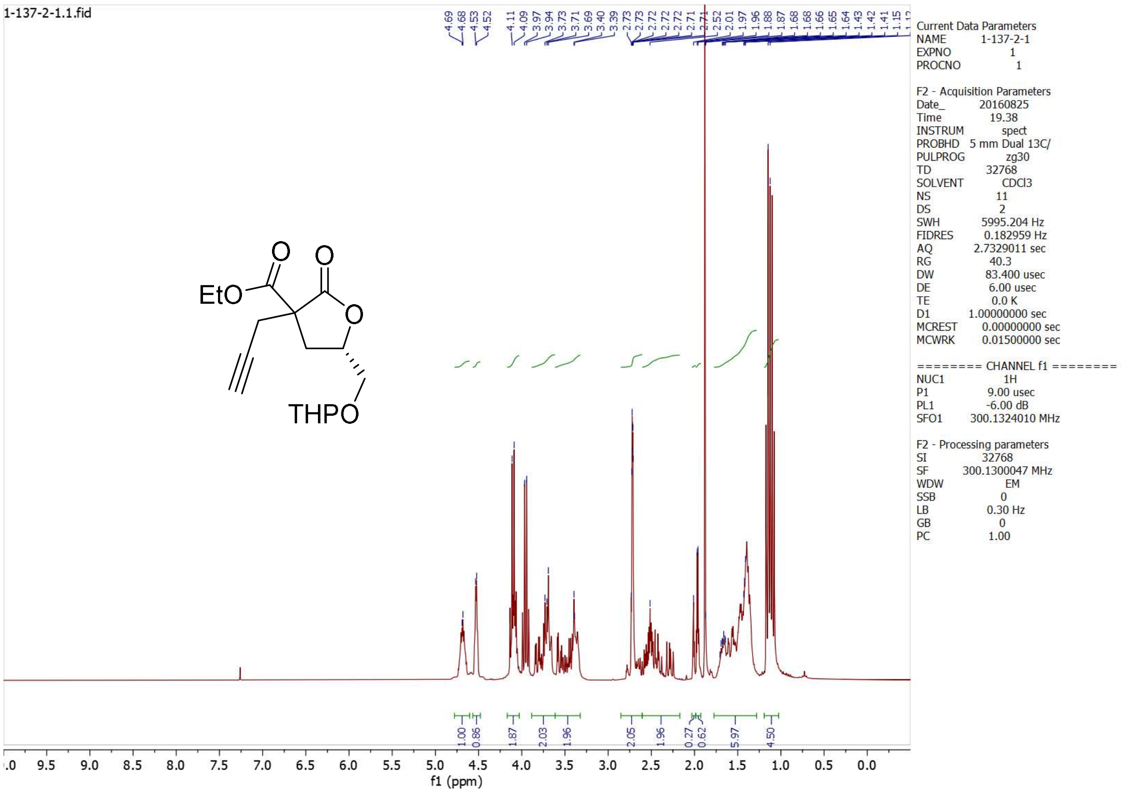

1-137-2-1.1.fid
4.69
4.68
4.53
4.52
4.11
4.09
3.97
3.94
3.73
3.71
3.69
3.40
3.39
2.73
2.73
2.72
2.72
2.72
2.71
2.71
2.52
2.01
1.97
1.96
1.88
1.87
1.68
1.68
1.66
1.65
1.64
1.43
1.42
1.41
1.15
EtO
O
O
O
THPO
1.00
0.86
1.87
2.03
1.96
2.05
1.96
0.27
0.62
5.97
4.50
9.5
9.0
8.5
8.0
7.5
7.0
6.5
6.0
5.5
5.0
4.5
4.0
3.5
3.0
2.5
2.0
1.5
1.0
0.5
0.0
f1 (ppm)
Current Data Parameters
NAME 1-137-2-1
EXPNO 1
PROCNO 1
F2 - Acquisition Parameters
Date_ 20160825
Time 19.38
INSTRUM spect
PROBHD 5 mm Dual 13C/
PULPROG zg30
TD 32768
SOLVENT CDCl3
NS 11
DS 2
SWH 5995.204 Hz
FIDRES 0.182959 Hz
AQ 2.7329011 sec
RG 40.3
DW 83.400 usec
DE 6.00 usec
TE 0.0 K
D1 1.00000000 sec
MCREST 0.00000000 sec
MCWRK 0.01500000 sec
======== CHANNEL f1 ========
NUC1 1H
P1 9.00 usec
PL1 -6.00 dB
SFO1 300.1324010 MHz
F2 - Processing parameters
SI 32768
SF 300.1300047 MHz
WDW EM
SSB 0
LB 0.30 Hz
GB 0
PC 1.00

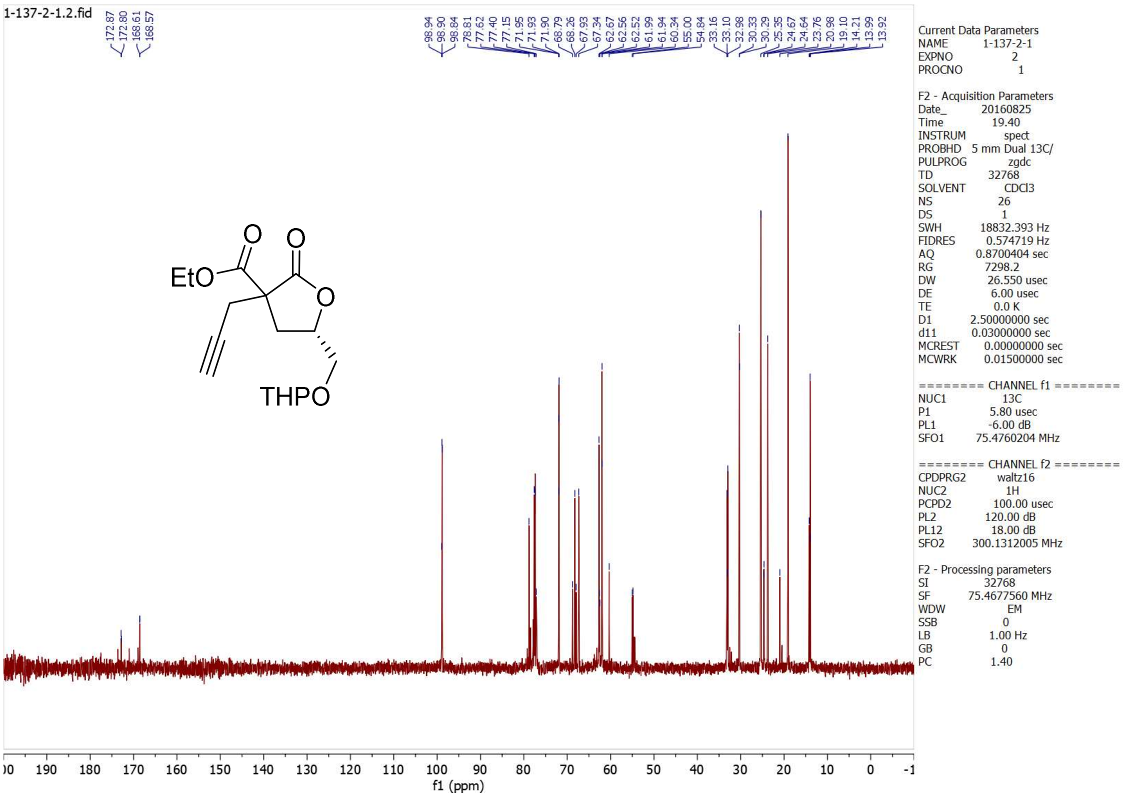
1-137-2-1.2.fid
172.87
172.80
168.61
168.57
98.94
98.90
98.84
78.81
77.62
77.40
77.15
71.95
71.93
71.90
68.79
68.26
67.93
67.94
62.67
62.56
62.52
61.99
61.94
60.34
55.00
54.84
33.16
33.10
32.98
30.33
30.29
25.35
24.67
24.64
23.76
20.98
19.10
14.21
13.99
13.92
EtO
O
O
O
THPO
Current Data Parameters
NAME 1-137-2-1
EXPNO 2
PROCNO 1
F2 - Acquisition Parameters
Date_ 20160825
Time 19.40
INSTRUM spect
PROBHD 5 mm Dual 13C/
PULPROG zgdc
TD 32768
SOLVENT CDCl3
NS 26
DS 1
SWH 18832.393 Hz
FIDRES 0.574719 Hz
AQ 0.8700404 sec
RG 7298.2
DW 26.550 usec
DE 6.00 usec
TE 0.0 K
D1 2.50000000 sec
d11 0.03000000 sec
MCREST 0.00000000 sec
MCWRK 0.01500000 sec
======== CHANNEL f1 ========
NUC1 13C
P1 5.80 usec
PL1 -6.00 dB
SFO1 75.4760204 MHz
======== CHANNEL f2 ========
CPDPRG2 waltz16
NUC2 1H
PCPD2 100.00 usec
PL2 120.00 dB
PL12 18.00 dB
SFO2 300.1312005 MHz
F2 - Processing parameters
SI 32768
SF 75.4677560 MHz
WDW EM
SSB 0
LB 1.00 Hz
GB 0
PC 1.40
190 180 170 160 150 140 130 120 110 100 90 80 70 60 50 40 30 20 10 0
f1 (ppm)

**5.5*S***

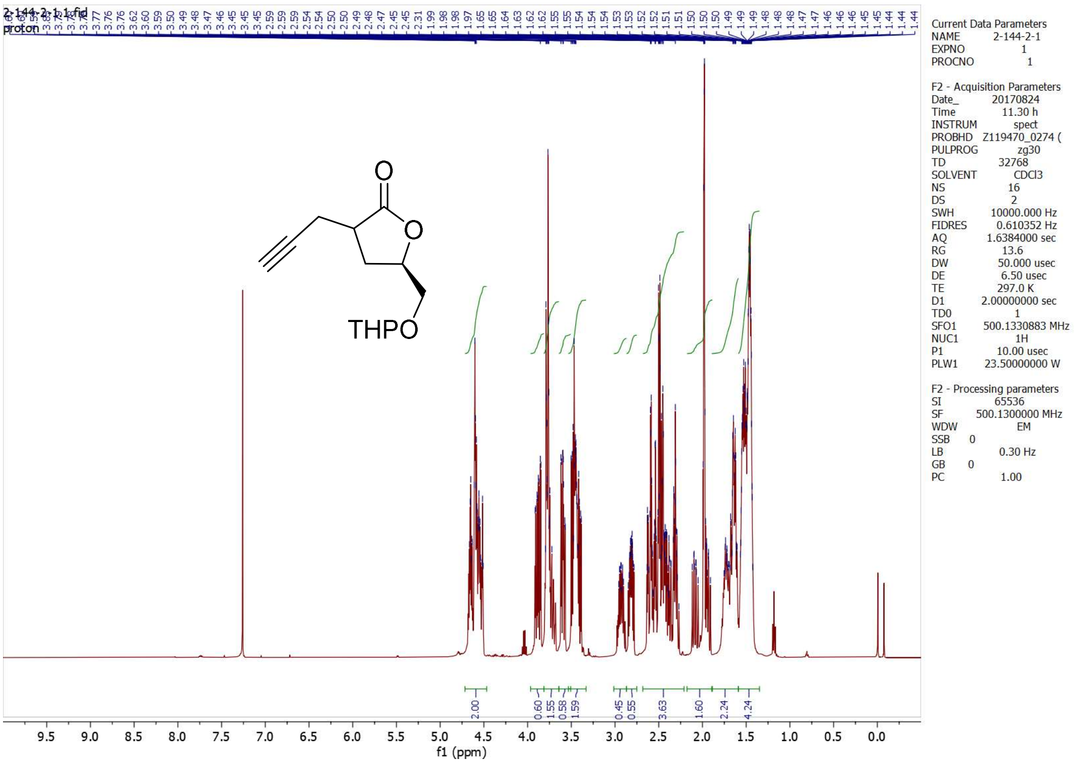
2-144-2-1.1.fid
proton
THPO
O
O
Current Data Parameters
NAME 2-144-2-1
EXPNO 1
PROCNO 1
F2 - Acquisition Parameters
Date_ 20170824
Time 11.30 h
INSTRUM spect
PROBHD Z119470_0274 (
PULPROG zg30
TD 32768
SOLVENT CDCl3
NS 16
DS 2
SWH 10000.000 Hz
FIDRES 0.610352 Hz
AQ 1.6384000 sec
RG 13.6
DW 50.000 usec
DE 6.50 usec
TE 297.0 K
D1 2.00000000 sec
TD0 1
SFO1 500.1330883 MHz
NUC1 1H
P1 10.00 usec
PLW1 23.50000000 W
F2 - Processing parameters
SI 65536
SF 500.1300000 MHz
WDW EM
SSB 0
LB 0.30 Hz
GB 0
PC 1.00
2.00
0.60
1.55
0.58
1.59
0.45
0.55
3.63
1.60
2.24
4.24
f1 (ppm)

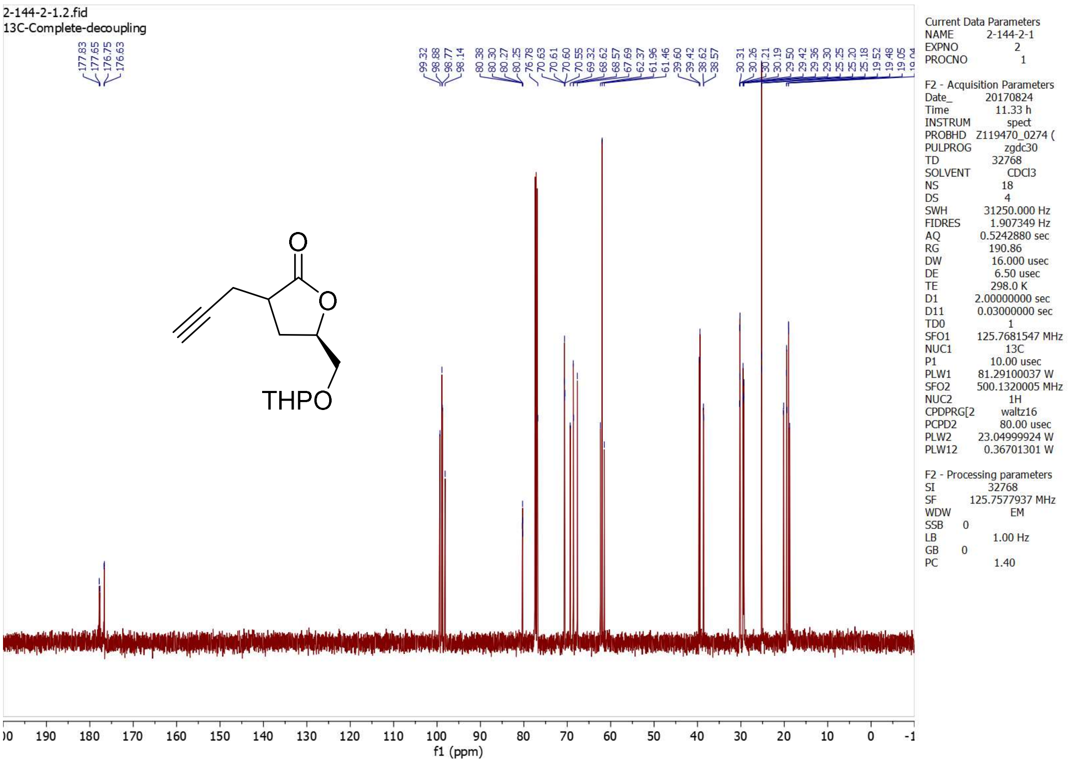
2-144-2-1.2.fid
13C-Complete-decoupling
177.83
177.65
176.75
176.63
99.32
98.88
98.77
98.14
80.38
80.30
80.27
80.25
76.78
70.63
70.61
70.60
70.55
69.32
68.62
68.57
67.69
62.37
61.96
61.46
39.60
39.42
38.62
38.57
30.31
30.26
30.21
30.19
29.50
29.42
29.36
29.30
25.25
25.20
25.18
19.52
19.48
19.05
THPO
O
O
Current Data Parameters
NAME 2-144-2-1
EXPNO 2
PROCNO 1
F2 - Acquisition Parameters
Date_ 20170824
Time 11.33 h
INSTRUM spect
PROBHD Z119470_0274 (
PULPROG zgdc30
TD 32768
SOLVENT CDCl3
NS 18
DS 4
SWH 31250.000 Hz
FIDRES 1.907349 Hz
AQ 0.5242880 sec
RG 190.86
DW 16.000 usec
DE 6.50 usec
TE 298.0 K
D1 2.00000000 sec
D11 0.03000000 sec
TD0 1
SFO1 125.7681547 MHz
NUC1 13C
P1 10.00 usec
PLW1 81.29100037 W
SFO2 500.1320005 MHz
NUC2 1H
CPDPRG[2 waltz16
PCPD2 80.00 usec
PLW2 23.04999924 W
PLW12 0.36701301 W
F2 - Processing parameters
SI 32768
SF 125.7577937 MHz
WDW EM
SSB 0
LB 1.00 Hz
GB 0
PC 1.40
f1 (ppm)

**5.6*S***

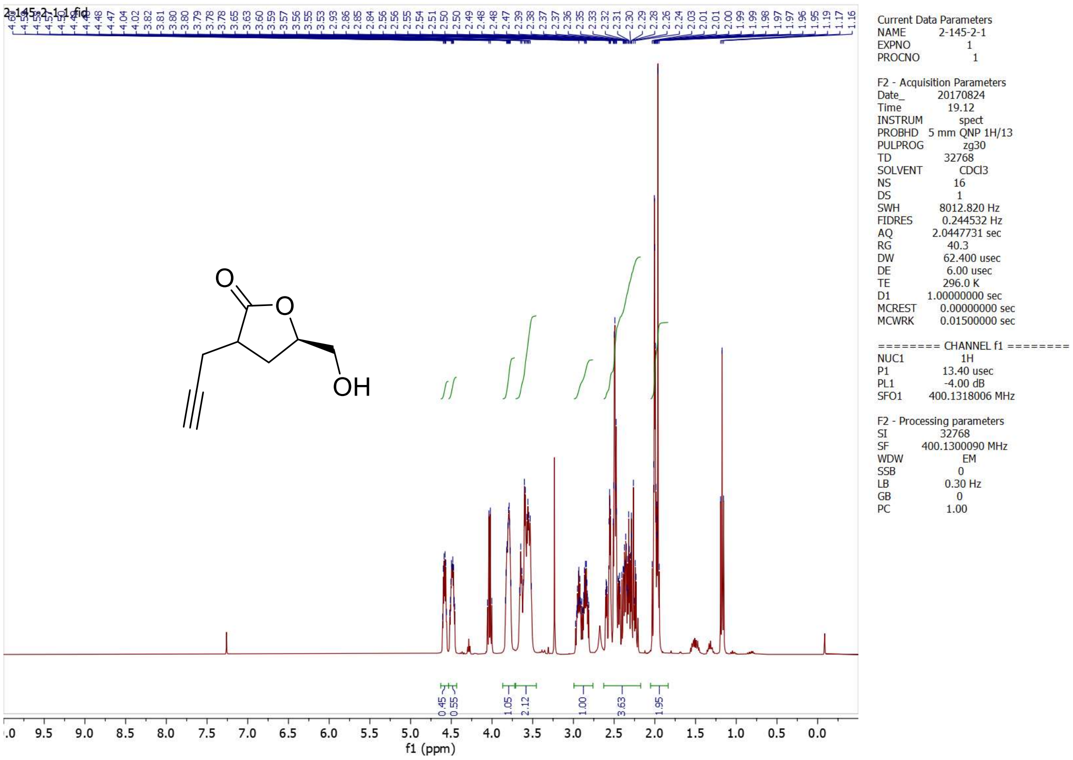
2-145-2-1.1.fid
Current Data Parameters
NAME 2-145-2-1
EXPNO 1
PROCNO 1
F2 - Acquisition Parameters
Date_ 20170824
Time 19.12
INSTRUM spect
PROBHD 5 mm QNP 1H/13
PULPROG zg30
TD 32768
SOLVENT CDCl3
NS 16
DS 1
SWH 8012.820 Hz
FIDRES 0.244532 Hz
AQ 2.0447731 sec
RG 40.3
DW 62.400 usec
DE 6.00 usec
TE 296.0 K
D1 1.00000000 sec
MCREST 0.00000000 sec
MCWRK 0.01500000 sec
======== CHANNEL f1 ========
NUC1 1H
P1 13.40 usec
PL1 -4.00 dB
SFO1 400.1318006 MHz
F2 - Processing parameters
SI 32768
SF 400.1300090 MHz
WDW EM
SSB 0
LB 0.30 Hz
GB 0
PC 1.00
OH
0.45
0.55
1.05
2.12
1.00
3.63
1.95
f1 (ppm)

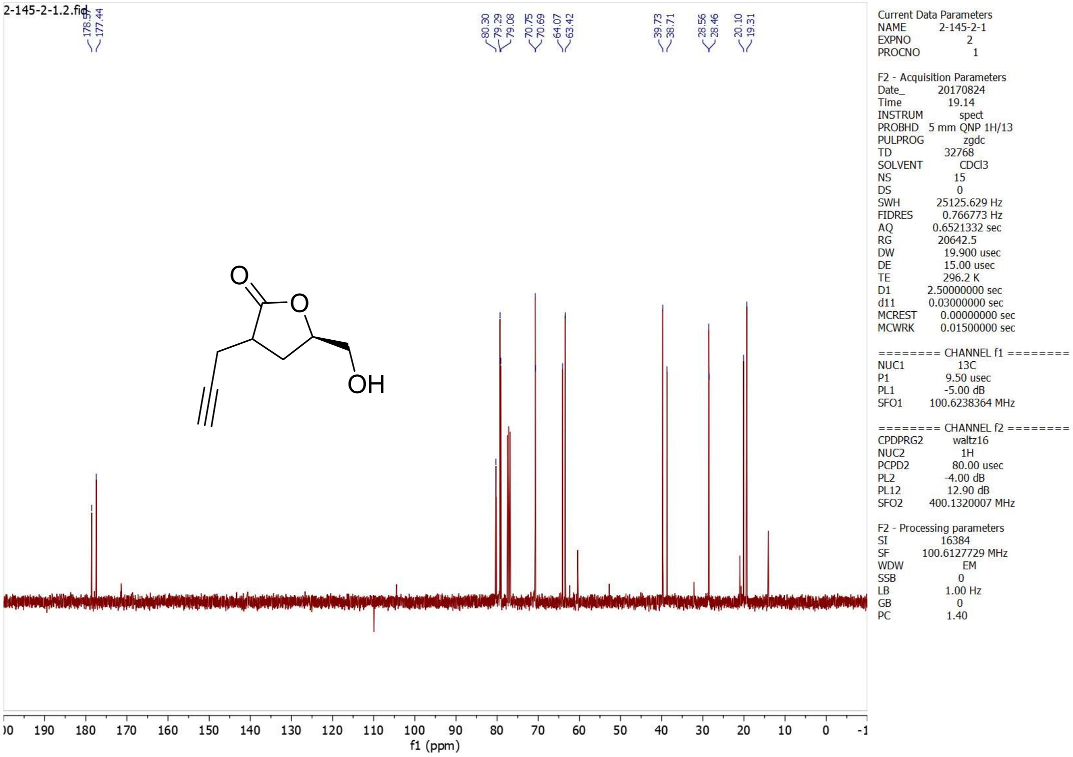
2-145-2-1.2.fid
178.59
177.44
80.30
79.29
79.08
70.75
70.69
64.07
63.42
39.73
38.71
28.56
28.46
20.10
19.31
Current Data Parameters
NAME 2-145-2-1
EXPNO 2
PROCNO 1
F2 - Acquisition Parameters
Date_ 20170824
Time 19.14
INSTRUM spect
PROBHD 5 mm QNP 1H/13
PULPROG zgdc
TD 32768
SOLVENT CDCl3
NS 15
DS 0
SWH 25125.629 Hz
FIDRES 0.766773 Hz
AQ 0.6521332 sec
RG 20642.5
DW 19.900 usec
DE 15.00 usec
TE 296.2 K
D1 2.50000000 sec
d11 0.03000000 sec
MCREST 0.00000000 sec
MCWRK 0.01500000 sec
======== CHANNEL f1 ========
NUC1 13C
P1 9.50 usec
PL1 -5.00 dB
SFO1 100.6238364 MHz
======== CHANNEL f2 ========
CPDPRG2 waltz16
NUC2 1H
PCPD2 80.00 usec
PL2 -4.00 dB
PL12 12.90 dB
SFO2 400.1320007 MHz
F2 - Processing parameters
SI 16384
SF 100.6127729 MHz
WDW EM
SSB 0
LB 1.00 Hz
GB 0
PC 1.40
OH
f1 (ppm)

## 5.6*R*

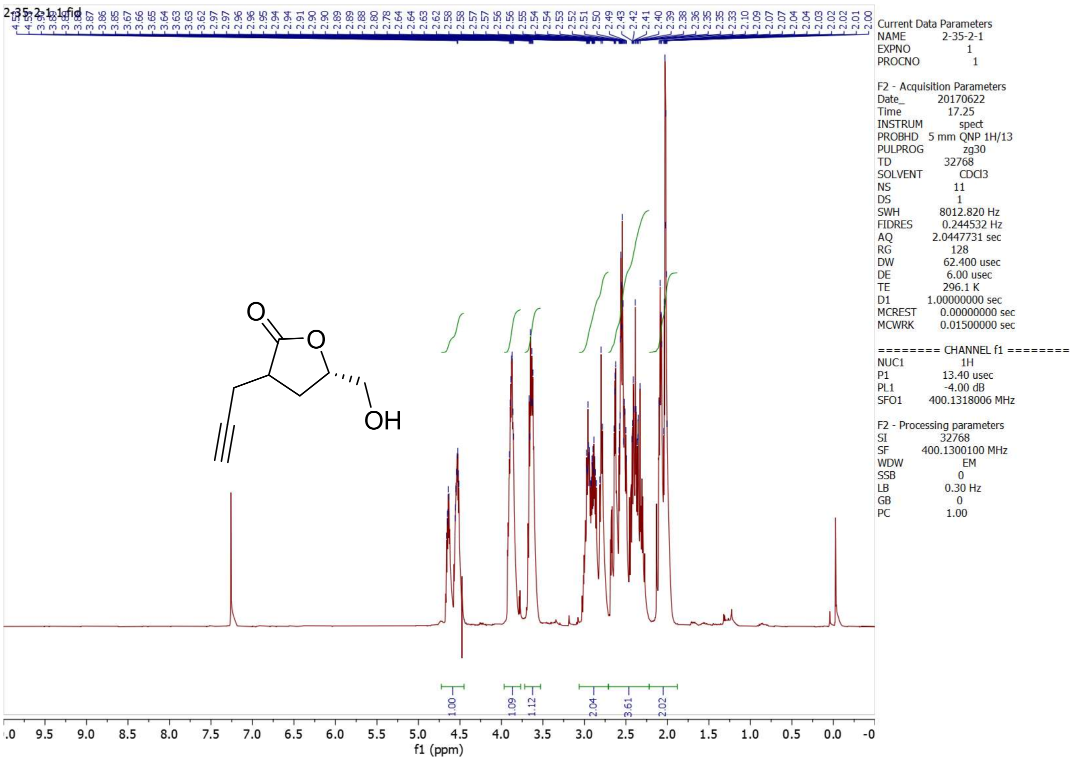
2-35-2-1.1.fid
OH
1.00
1.09
1.12
2.04
3.61
2.02
f1 (ppm)
Current Data Parameters
NAME 2-35-2-1
EXPNO 1
PROCNO 1
F2 - Acquisition Parameters
Date_ 20170622
Time 17.25
INSTRUM spect
PROBHD 5 mm QNP 1H/13
PULPROG zg30
TD 32768
SOLVENT CDCl3
NS 11
DS 1
SWH 8012.820 Hz
FIDRES 0.244532 Hz
AQ 2.0447731 sec
RG 128
DW 62.400 usec
DE 6.00 usec
TE 296.1 K
D1 1.00000000 sec
MCREST 0.00000000 sec
MCWRK 0.01500000 sec
======== CHANNEL f1 ========
NUC1 1H
P1 13.40 usec
PL1 -4.00 dB
SFO1 400.1318006 MHz
F2 - Processing parameters
SI 32768
SF 400.1300100 MHz
WDW EM
SSB 0
LB 0.30 Hz
GB 0
PC 1.00

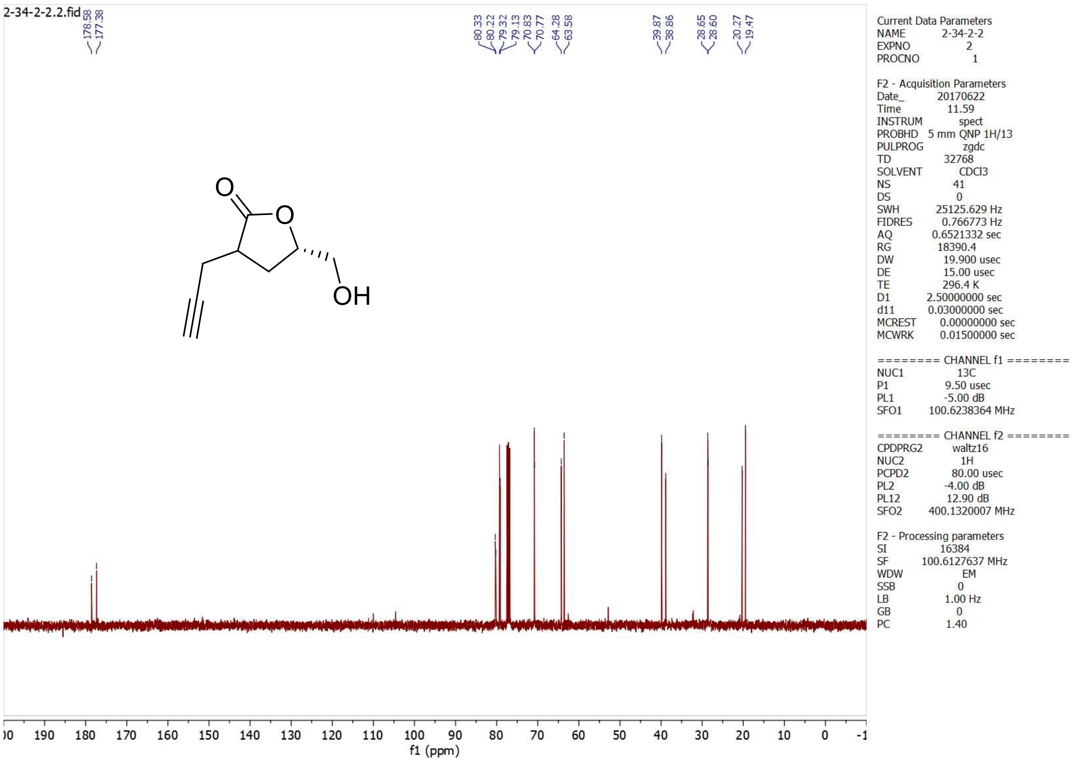
2-34-2-2.2.fid
178.58
177.38
80.33
80.22
79.32
79.13
70.83
70.77
64.28
63.58
39.87
38.86
28.65
28.60
20.27
19.47
OH
f1 (ppm)
Current Data Parameters
NAME 2-34-2-2
EXPNO 2
PROCNO 1
F2 - Acquisition Parameters
Date_ 20170622
Time 11.59
INSTRUM spect
PROBHD 5 mm QNP 1H/13
PULPROG zgdc
TD 32768
SOLVENT CDCl3
NS 41
DS 0
SWH 25125.629 Hz
FIDRES 0.766773 Hz
AQ 0.6521332 sec
RG 18390.4
DW 19.900 usec
DE 15.00 usec
TE 296.4 K
D1 2.50000000 sec
d11 0.03000000 sec
MCREST 0.00000000 sec
MCWRK 0.01500000 sec
======== CHANNEL f1 ========
NUC1 13C
P1 9.50 usec
PL1 -5.00 dB
SFO1 100.6238364 MHz
======== CHANNEL f2 ========
CPDPRG2 waltz16
NUC2 1H
PCPD2 80.00 usec
PL2 -4.00 dB
PL12 12.90 dB
SFO2 400.1320007 MHz
F2 - Processing parameters
SI 16384
SF 100.6127637 MHz
WDW EM
SSB 0
LB 1.00 Hz
GB 0
PC 1.40

## 5.7*S*-1

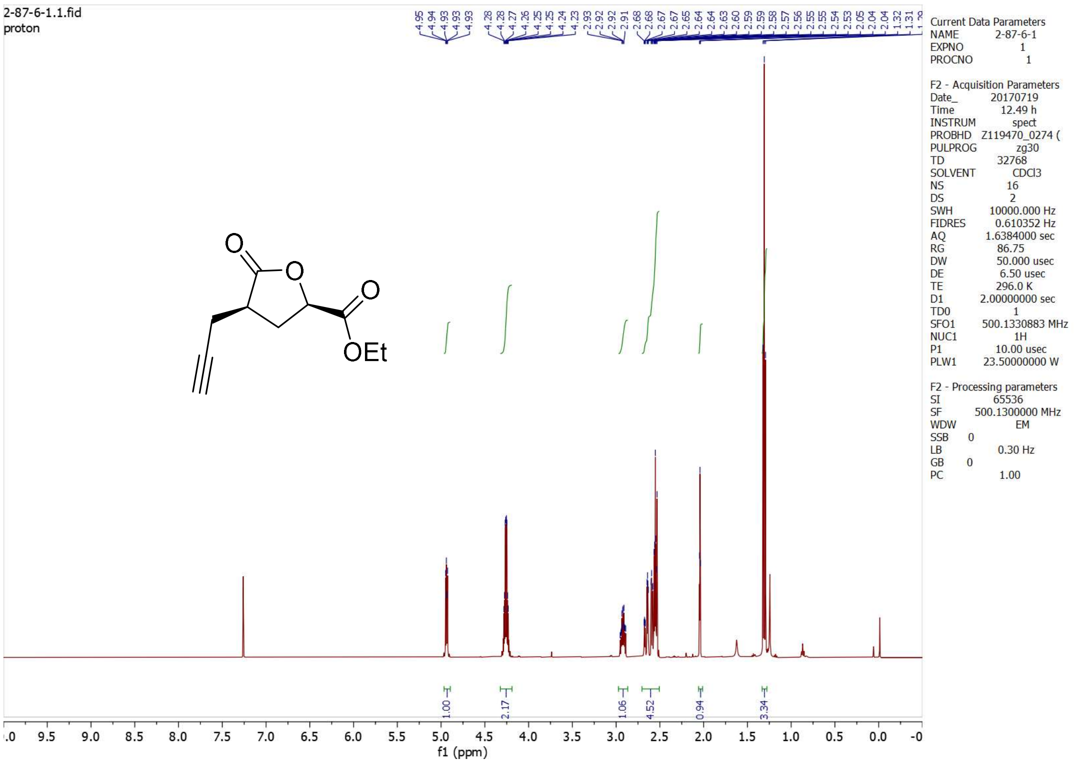

2-87-6-1.1.fid
proton
4.95 4.94 4.93 4.93 4.28 4.28 4.27 4.26 4.25 4.25 4.24 4.23 2.93 2.92 2.92 2.91 2.68 2.68 2.67 2.65 2.64 2.64 2.63 2.60 2.59 2.59 2.58 2.57 2.56 2.55 2.55 2.54 2.53 2.05 2.04 2.04 1.32 1.31
O
O
O
OEt
1.00
2.17
1.06
4.52
0.94
3.34
f1 (ppm)
Current Data Parameters
NAME 2-87-6-1
EXPNO 1
PROCNO 1
F2 - Acquisition Parameters
Date_ 20170719
Time 12.49 h
INSTRUM spect
PROBHD Z119470_0274 (
PULPROG zg30
TD 32768
SOLVENT CDCl3
NS 16
DS 2
SWH 10000.000 Hz
FIDRES 0.610352 Hz
AQ 1.6384000 sec
RG 86.75
DW 50.000 usec
DE 6.50 usec
TE 296.0 K
D1 2.00000000 sec
TD0 1
SFO1 500.1330883 MHz
NUC1 1H
P1 10.00 usec
PLW1 23.50000000 W
F2 - Processing parameters
SI 65536
SF 500.1300000 MHz
WDW EM
SSB 0
LB 0.30 Hz
GB 0
PC 1.00

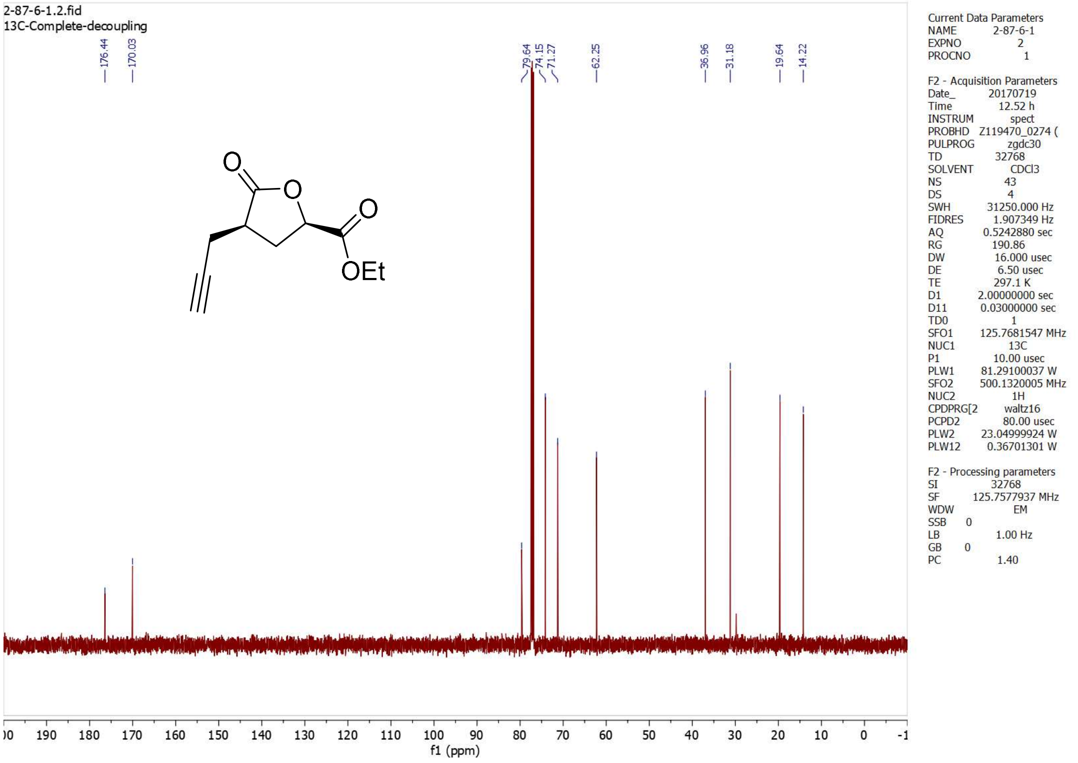

2-87-6-1.2.fid
13C-Complete-decoupling
176.44
170.03
79.64
74.15
71.27
62.25
36.96
31.18
19.64
14.22
O
O
O
OEt
f1 (ppm)
Current Data Parameters
NAME 2-87-6-1
EXPNO 2
PROCNO 1
F2 - Acquisition Parameters
Date_ 20170719
Time 12.52 h
INSTRUM spect
PROBHD Z119470_0274 (
PULPROG zgdc30
TD 32768
SOLVENT CDCl3
NS 43
DS 4
SWH 31250.000 Hz
FIDRES 1.907349 Hz
AQ 0.5242880 sec
RG 190.86
DW 16.000 usec
DE 6.50 usec
TE 297.1 K
D1 2.00000000 sec
D11 0.03000000 sec
TD0 1
SFO1 125.7681547 MHz
NUC1 13C
P1 10.00 usec
PLW1 81.29100037 W
SFO2 500.1320005 MHz
NUC2 1H
CPDPRG[2 waltz16
PCPD2 80.00 usec
PLW2 23.04999924 W
PLW12 0.36701301 W
F2 - Processing parameters
SI 32768
SF 125.7577937 MHz
WDW EM
SSB 0
LB 1.00 Hz
GB 0
PC 1.40

## 5.7*S*-2

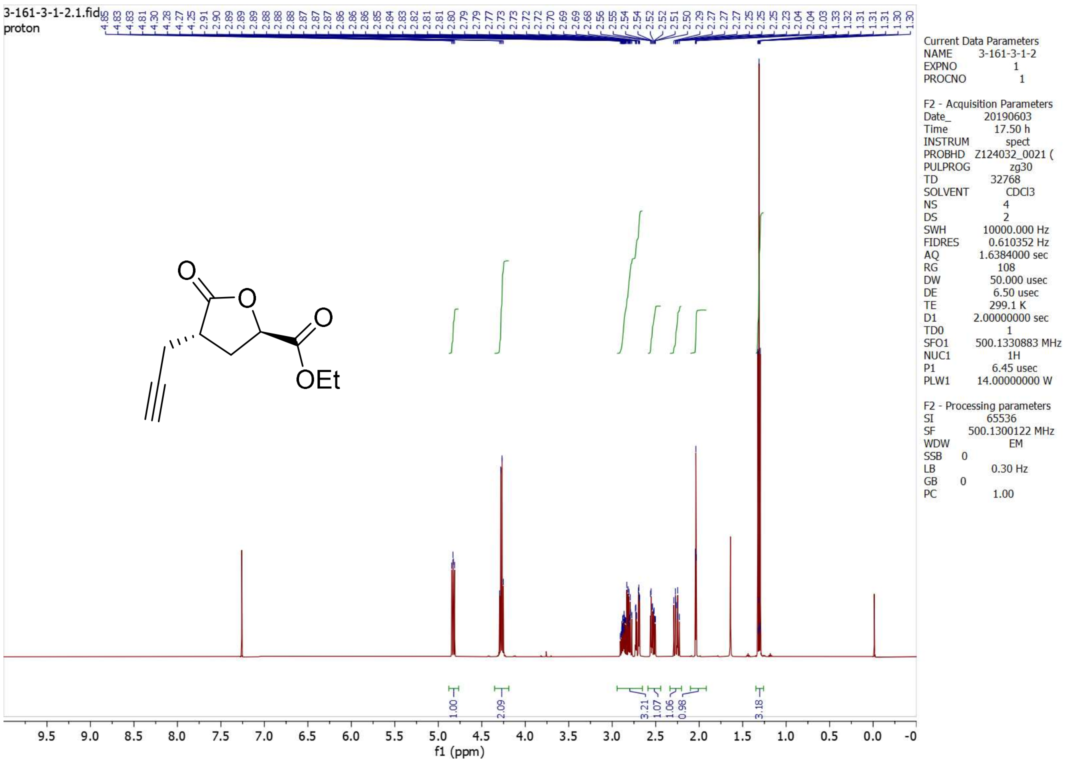
3-161-3-1-2.1.fid
proton
4.85 4.83 4.83 4.81 4.30 4.28 4.27 4.25 2.91 2.90 2.89 2.89 2.89 2.88 2.88 2.88 2.87 2.87 2.87 2.86 2.86 2.86 2.85 2.84 2.83 2.83 2.82 2.81 2.81 2.80 2.79 2.79 2.77 2.77 2.73 2.73 2.72 2.72 2.70 2.69 2.69 2.68 2.56 2.55 2.54 2.54 2.52 2.52 2.51 2.50 2.29 2.27 2.27 2.27 2.25 2.25 2.25 2.23 2.04 2.04 2.03 1.33 1.32 1.32 1.31 1.31 1.31 1.30 1.30
O
O
O
OEt
1.00
2.09
3.21
1.07
1.06
0.98
3.18
9.5 9.0 8.5 8.0 7.5 7.0 6.5 6.0 5.5 5.0 4.5 4.0 3.5 3.0 2.5 2.0 1.5 1.0 0.5 0.0 -0
f1 (ppm)
Current Data Parameters
NAME 3-161-3-1-2
EXPNO 1
PROCNO 1
F2 - Acquisition Parameters
Date_ 20190603
Time 17.50 h
INSTRUM spect
PROBHD Z124032_0021 (
PULPROG zg30
TD 32768
SOLVENT CDCl3
NS 4
DS 2
SWH 10000.000 Hz
FIDRES 0.610352 Hz
AQ 1.6384000 sec
RG 108
DW 50.000 usec
DE 6.50 usec
TE 299.1 K
D1 2.00000000 sec
TD0 1
SFO1 500.1330883 MHz
NUC1 1H
P1 6.45 usec
PLW1 14.00000000 W
F2 - Processing parameters
SI 65536
SF 500.1300122 MHz
WDW EM
SSB 0
LB 0.30 Hz
GB 0
PC 1.00

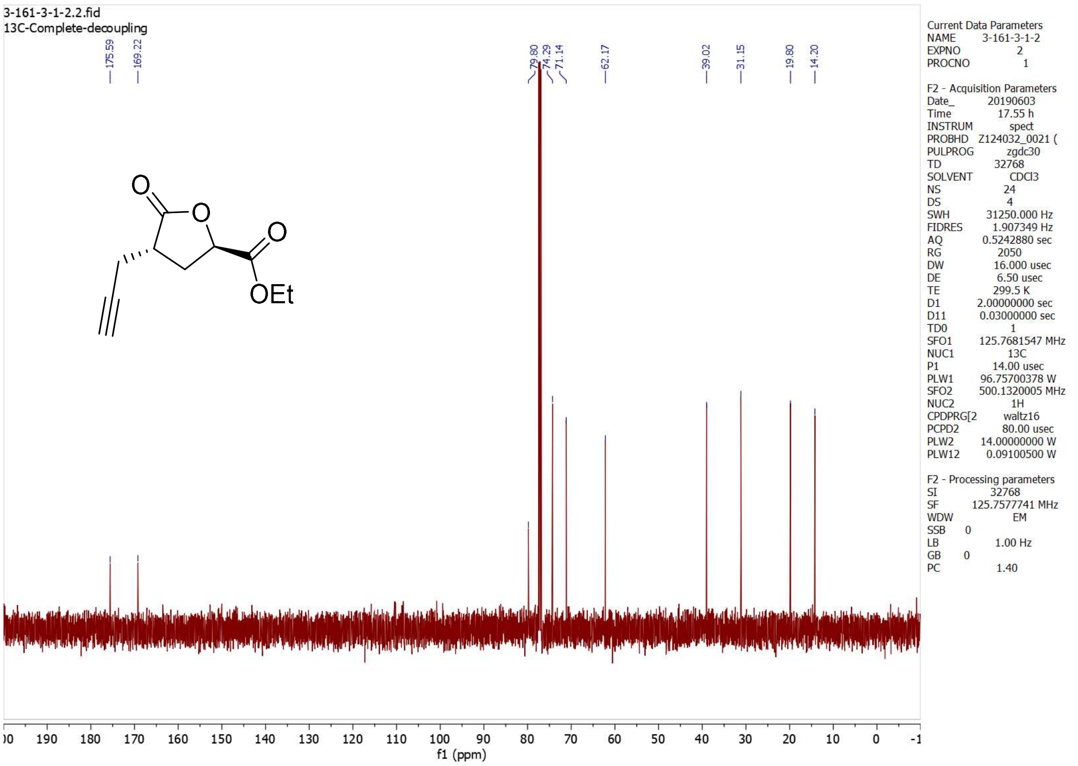
3-161-3-1-2.2.fid
13C-Complete-decoupling
175.59
169.22
79.80
74.29
71.14
62.17
39.02
31.15
19.80
14.20
O
O
O
OEt
00 190 180 170 160 150 140 130 120 110 100 90 80 70 60 50 40 30 20 10 0 -1
f1 (ppm)
Current Data Parameters
NAME 3-161-3-1-2
EXPNO 2
PROCNO 1
F2 - Acquisition Parameters
Date_ 20190603
Time 17.55 h
INSTRUM spect
PROBHD Z124032_0021 (
PULPROG zgdc30
TD 32768
SOLVENT CDCl3
NS 24
DS 4
SWH 31250.000 Hz
FIDRES 1.907349 Hz
AQ 0.5242880 sec
RG 2050
DW 16.000 usec
DE 6.50 usec
TE 299.5 K
D1 2.00000000 sec
D11 0.03000000 sec
TD0 1
SFO1 125.7681547 MHz
NUC1 13C
P1 14.00 usec
PLW1 96.75700378 W
SFO2 500.1320005 MHz
NUC2 1H
CPDPRG[2 waltz16
PCPD2 80.00 usec
PLW2 14.00000000 W
PLW12 0.09100500 W
F2 - Processing parameters
SI 32768
SF 125.7577741 MHz
WDW EM
SSB 0
LB 1.00 Hz
GB 0
PC 1.40

**5.7*R*-1**

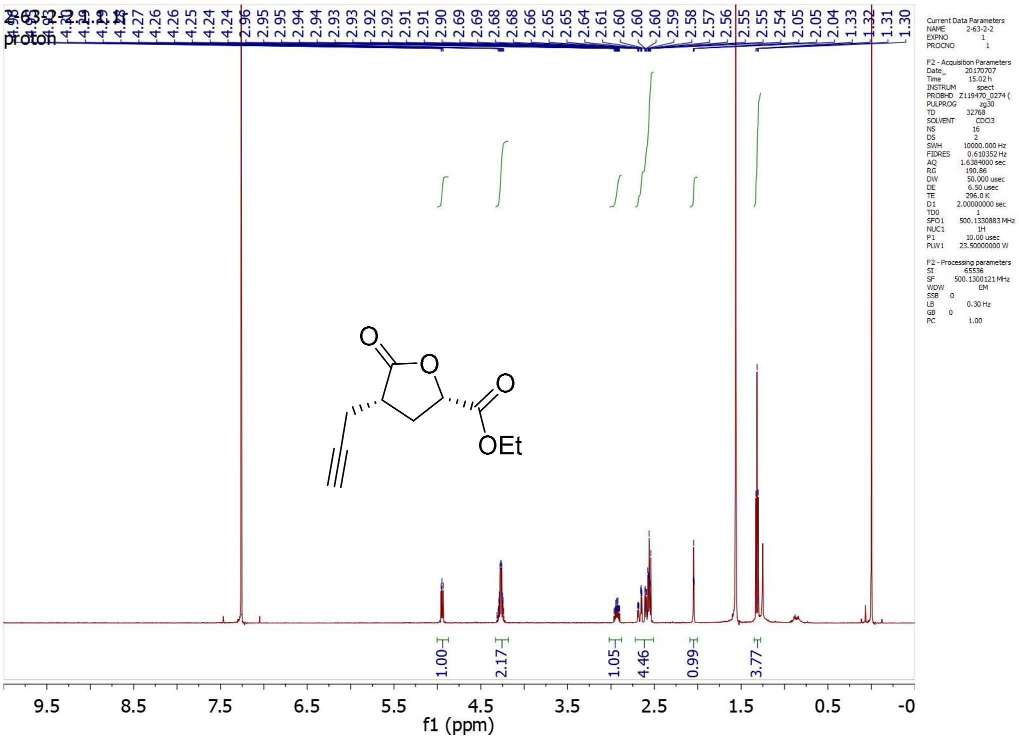
2-63-2-2.1.1.1r
proton
1.00
2.17
1.05
4.46
0.99
3.77
9.5 8.5 7.5 6.5 5.5 4.5 3.5 2.5 1.5 0.5 -0
f1 (ppm)
OEt

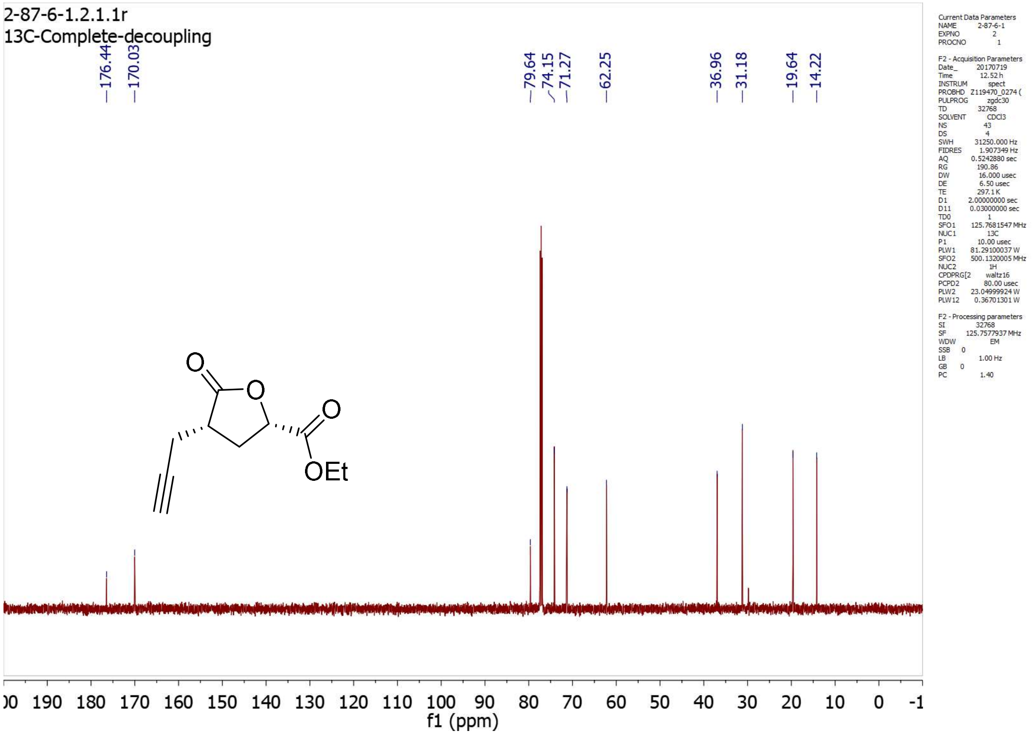
2-87-6-1.2.1.1r
13C-Complete-decoupling
176.44
170.03
79.64
74.15
71.27
62.25
36.96
31.18
19.64
14.22
f1 (ppm)
OEt

**5.7*R*-2**

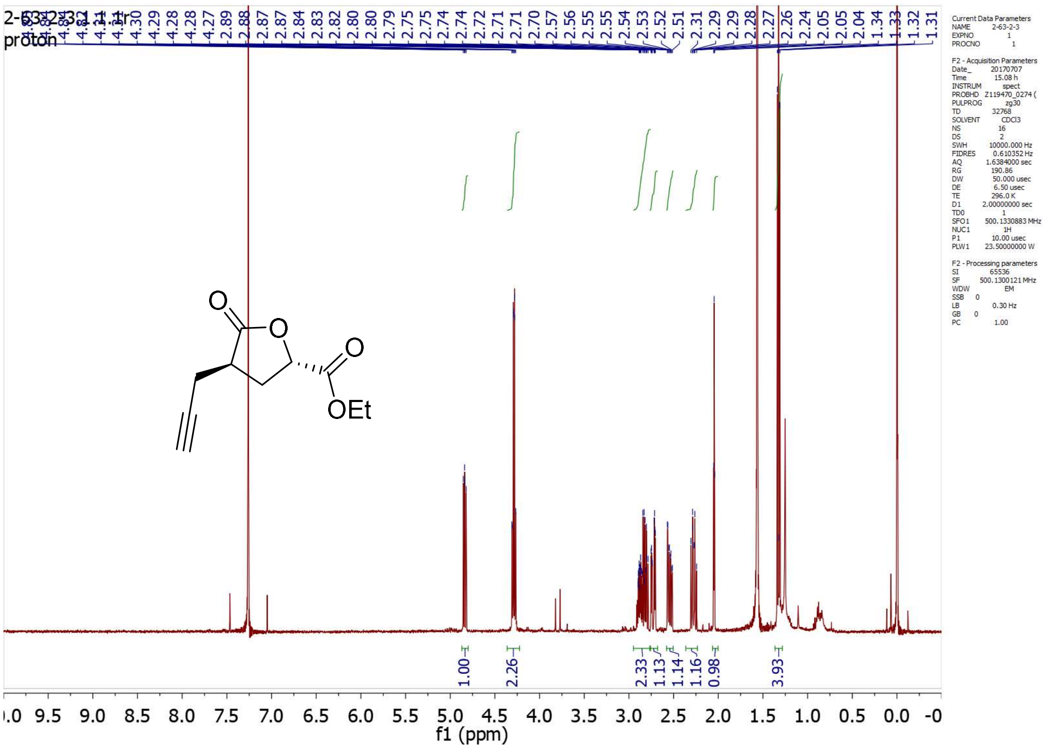

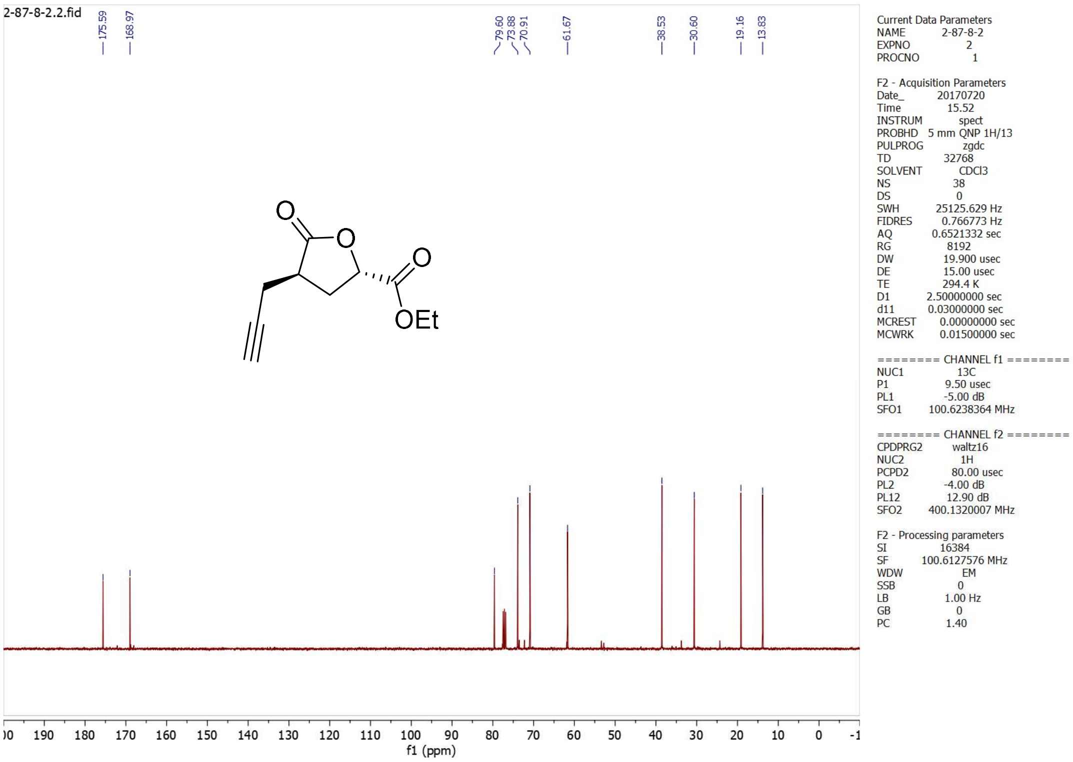

**5.8*S*-1**

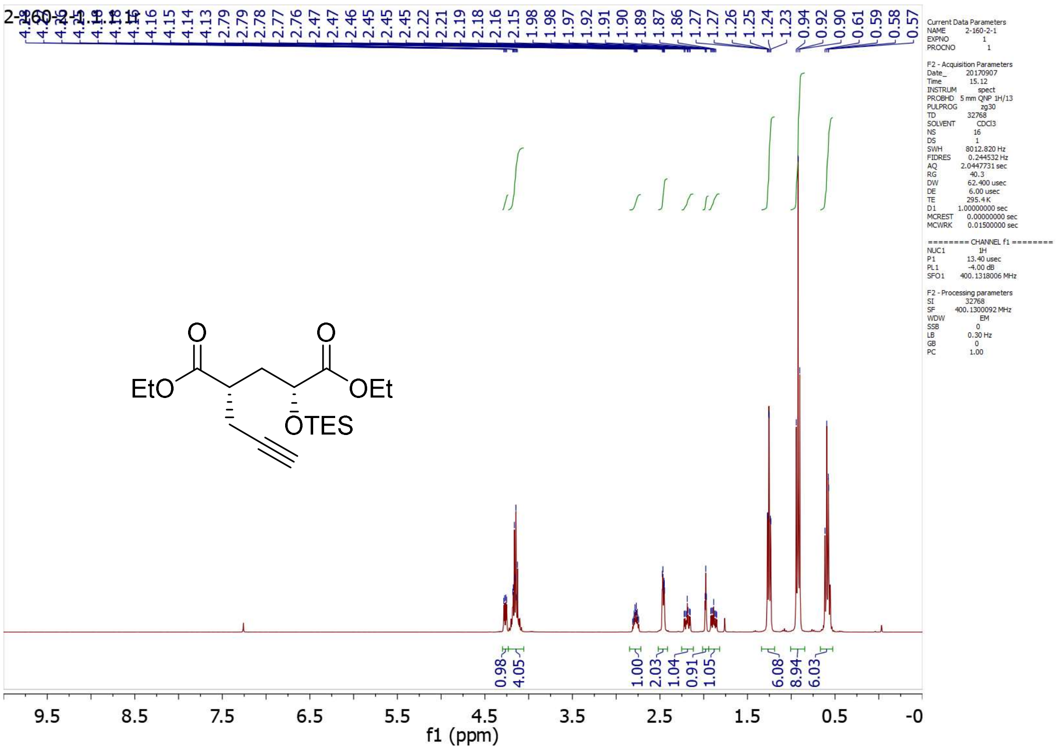

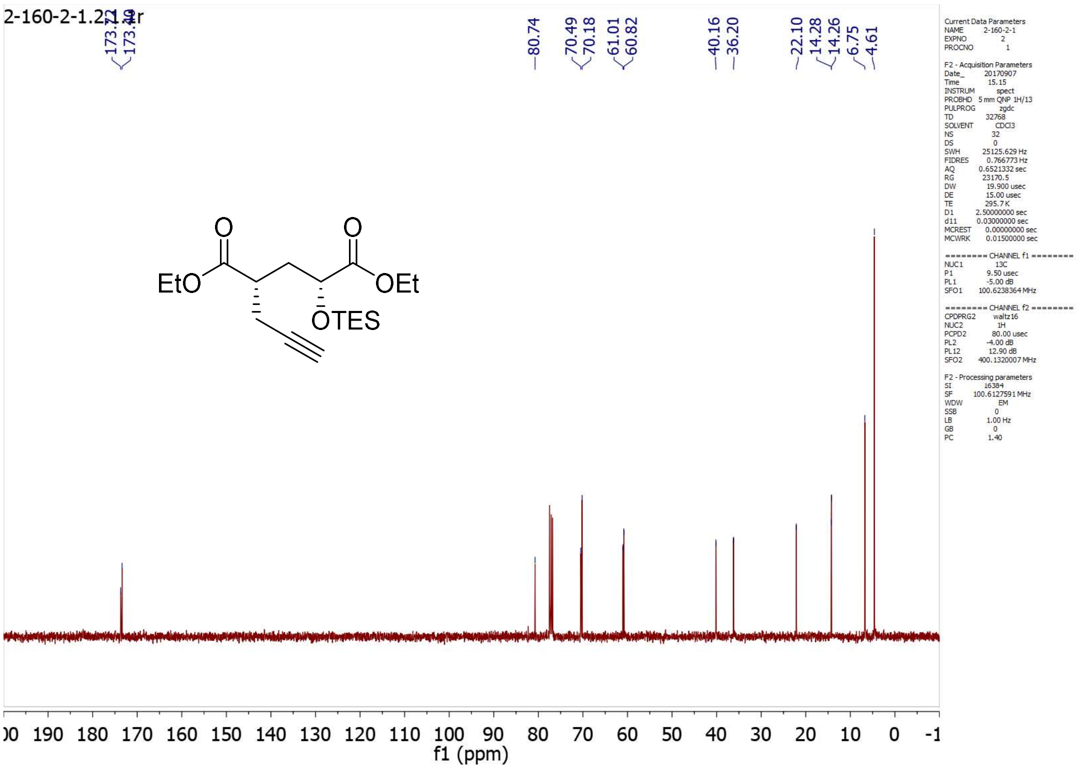

**5.8*S*-2**

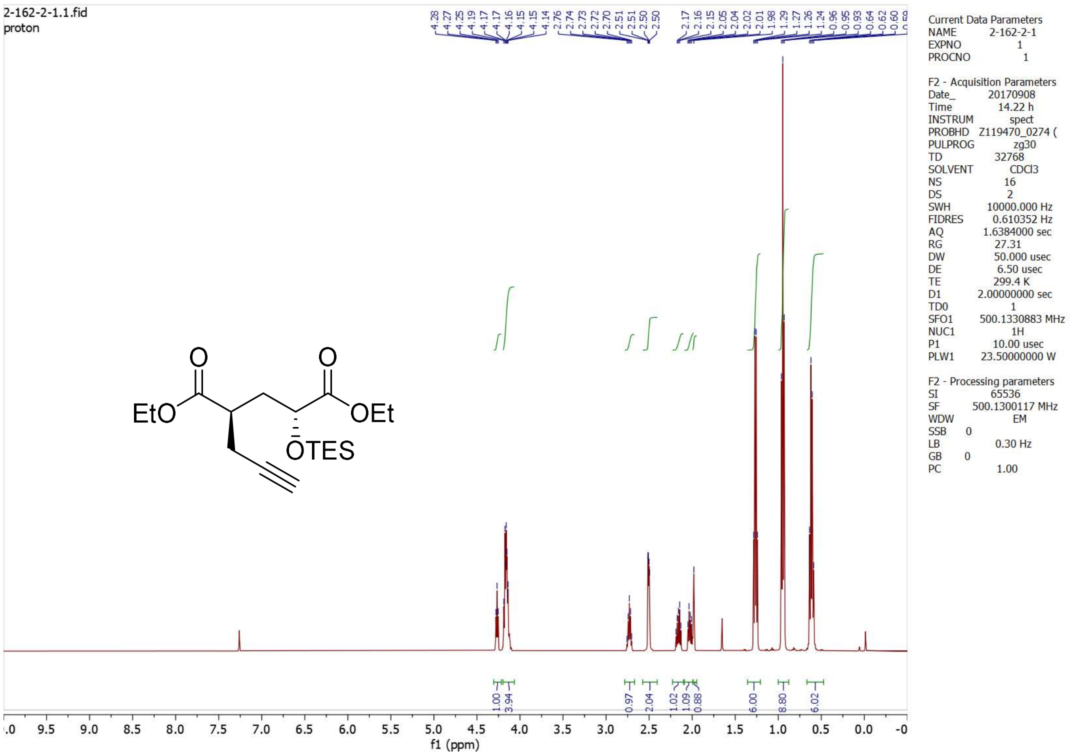
2-162-2-1.1.fid
proton
4.28 4.27 4.25 4.19 4.17 4.17 4.16 4.15 4.15 4.14 2.76 2.74 2.73 2.72 2.70 2.51 2.51 2.50 2.50 2.17 2.16 2.15 2.05 2.04 2.02 2.01 1.98 1.29 1.27 1.26 1.24 0.96 0.95 0.93 0.64 0.62 0.60
EtO
O
O
OEt
OTES
1.00 3.94 0.97 2.04 1.02 1.09 0.88 6.00 8.80 6.02
f1 (ppm)
Current Data Parameters
NAME 2-162-2-1
EXPNO 1
PROCNO 1
F2 - Acquisition Parameters
Date_ 20170908
Time 14.22 h
INSTRUM spect
PROBHD Z119470_0274 (
PULPROG zg30
TD 32768
SOLVENT CDCl3
NS 16
DS 2
SWH 10000.000 Hz
FIDRES 0.610352 Hz
AQ 1.6384000 sec
RG 27.31
DW 50.000 usec
DE 6.50 usec
TE 299.4 K
D1 2.00000000 sec
TD0 1
SFO1 500.1330883 MHz
NUC1 1H
P1 10.00 usec
PLW1 23.50000000 W
F2 - Processing parameters
SI 65536
SF 500.1300117 MHz
WDW EM
SSB 0
LB 0.30 Hz
GB 0
PC 1.00

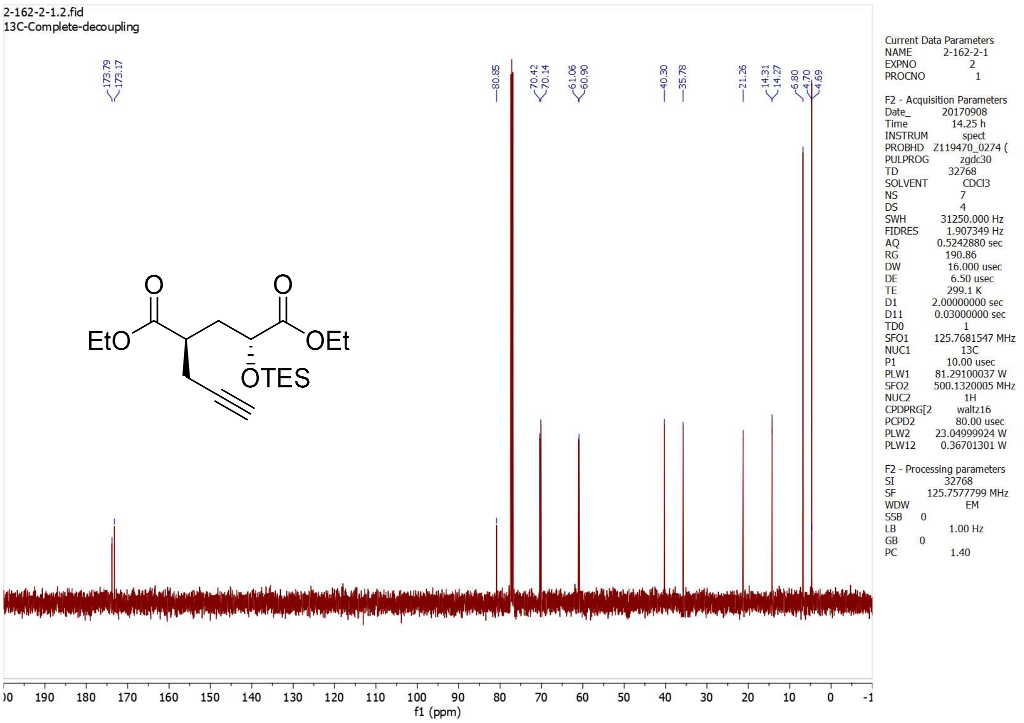
2-162-2-1.2.fid
13C-Complete-decoupling
173.79 173.17 80.85 70.42 70.14 61.06 60.90 40.30 35.78 21.26 14.31 14.27 6.80 4.70 4.69
EtO
O
O
OEt
OTES
f1 (ppm)
Current Data Parameters
NAME 2-162-2-1
EXPNO 2
PROCNO 1
F2 - Acquisition Parameters
Date_ 20170908
Time 14.25 h
INSTRUM spect
PROBHD Z119470_0274 (
PULPROG zgdc30
TD 32768
SOLVENT CDCl3
NS 7
DS 4
SWH 31250.000 Hz
FIDRES 1.907349 Hz
AQ 0.5242880 sec
RG 190.86
DW 16.000 usec
DE 6.50 usec
TE 299.1 K
D1 2.00000000 sec
D11 0.03000000 sec
TD0 1
SFO1 125.7681547 MHz
NUC1 13C
P1 10.00 usec
PLW1 81.29100037 W
SFO2 500.1320005 MHz
NUC2 1H
CPDPRG[2 waltz16
PCPD2 80.00 usec
PLW2 23.04999924 W
PLW12 0.36701301 W
F2 - Processing parameters
SI 32768
SF 125.7577799 MHz
WDW EM
SSB 0
LB 1.00 Hz
GB 0
PC 1.40

**5.8*R*-1**

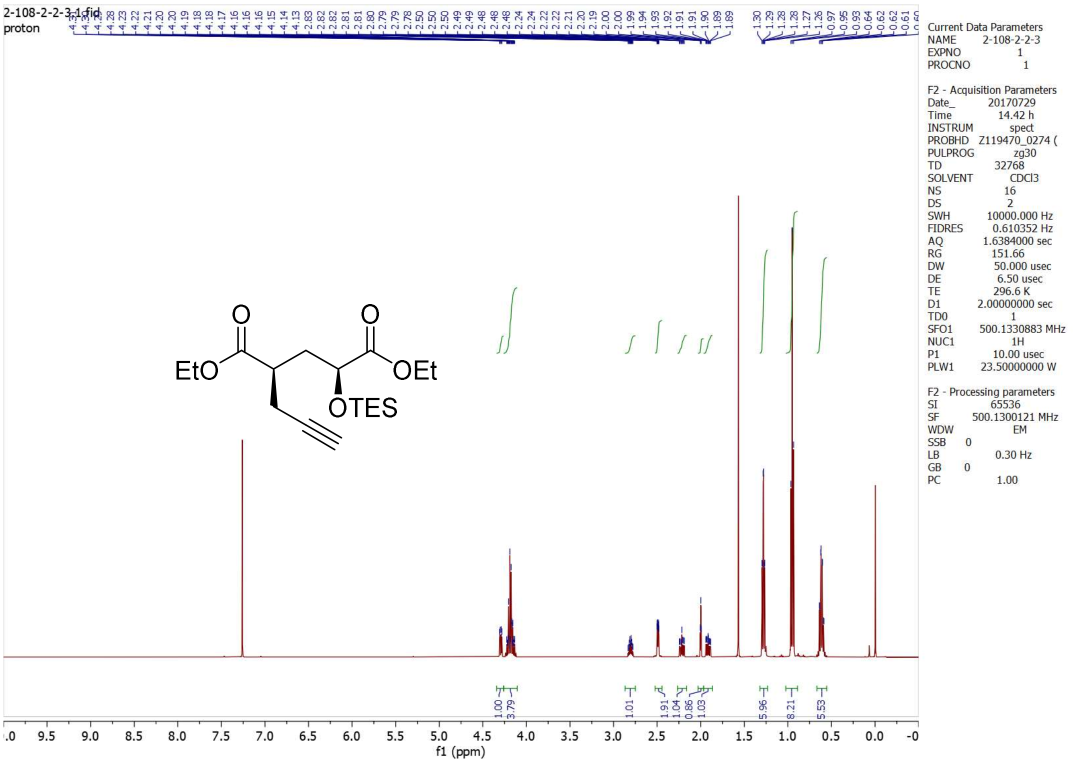

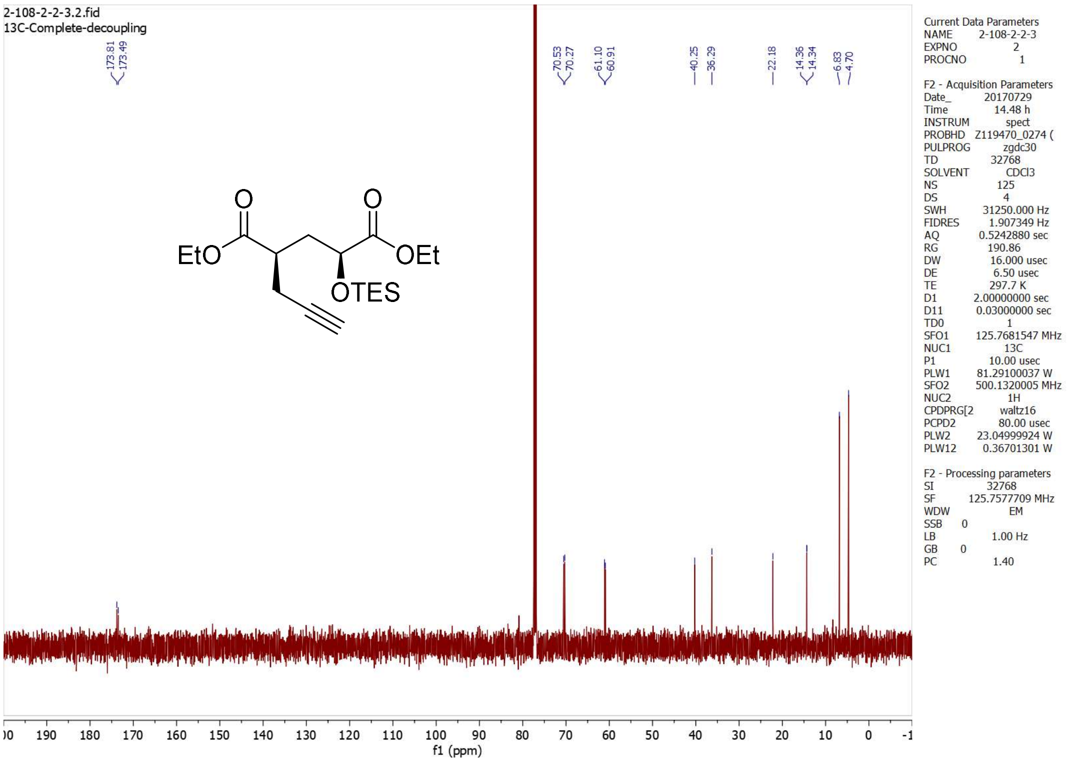

**5.8*R*-2**

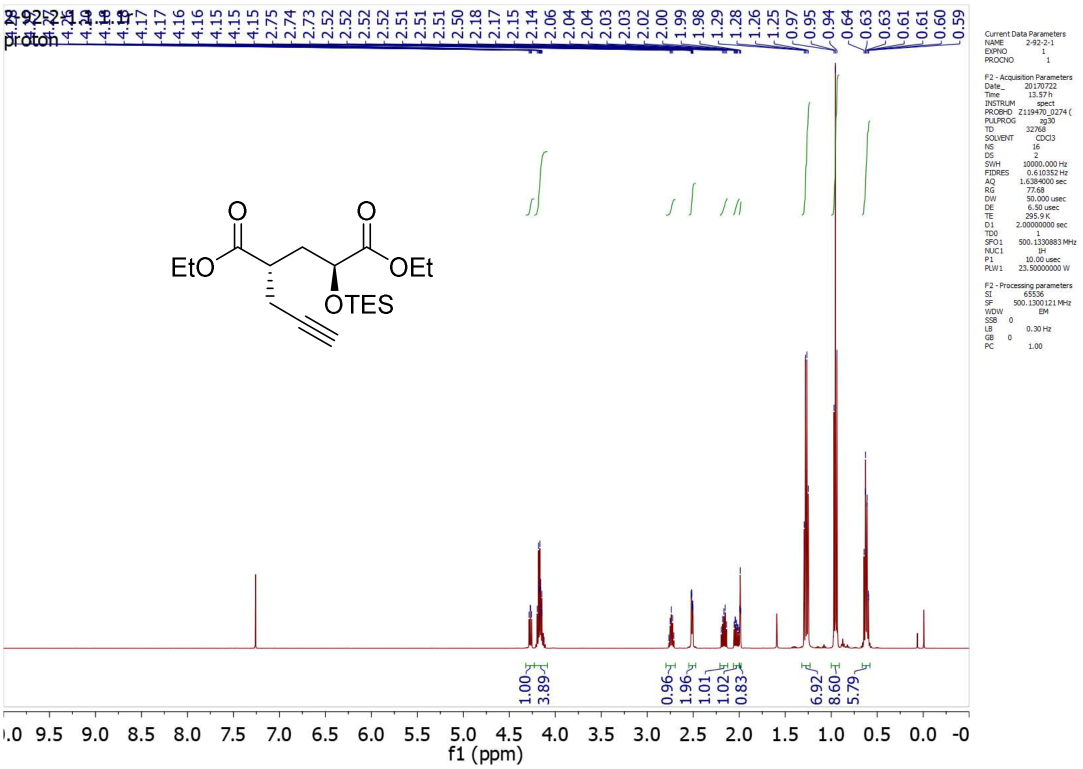
proton
EtO
OEt
OTES
f1 (ppm)

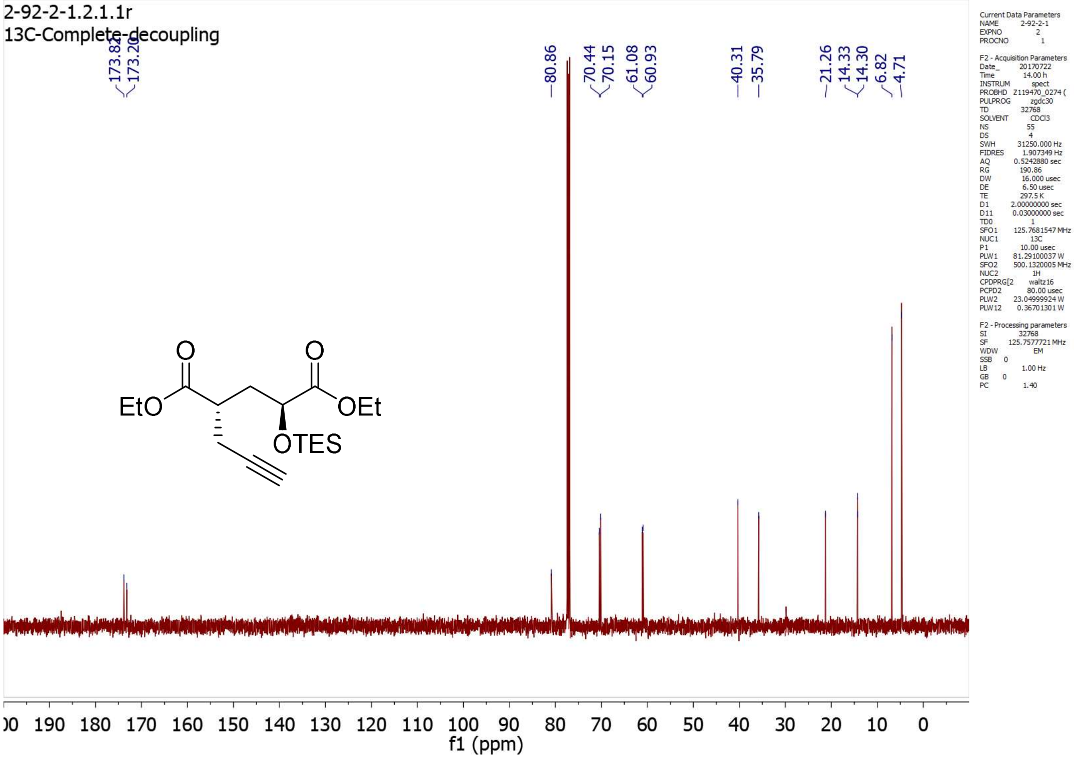
2-92-2-1.2.1.1r
13C-Complete-decoupling
EtO
OEt
OTES
f1 (ppm)

**5*S*-1 (5*RS*)**

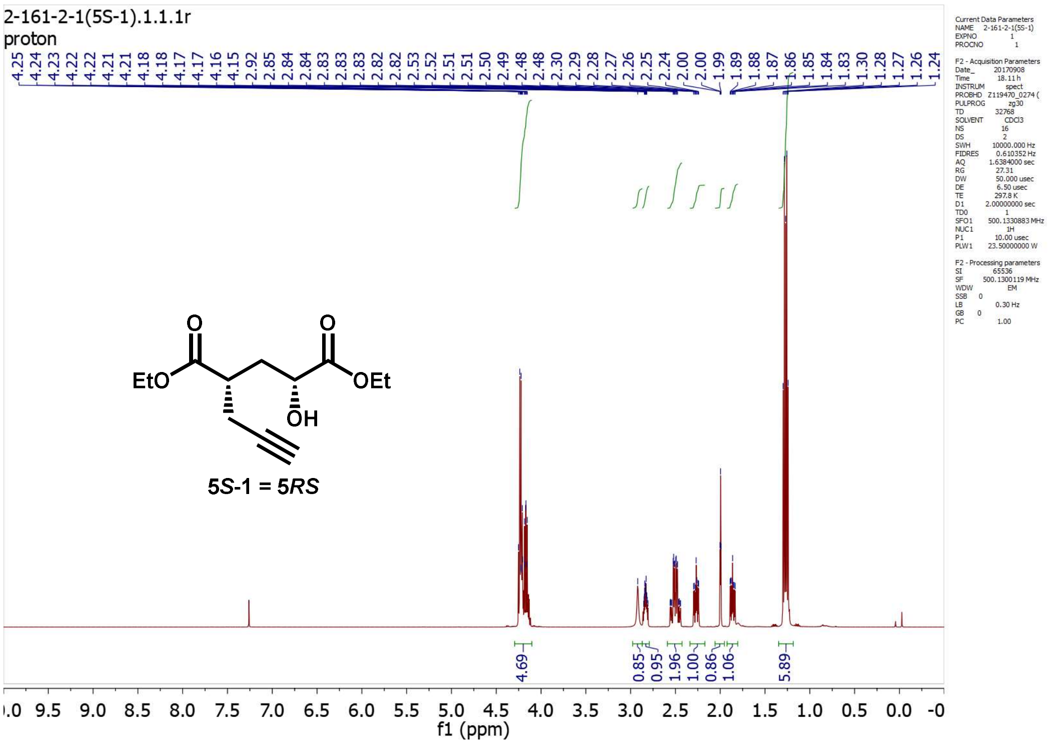

2-161-2-1(5S-1).1.1.1r
proton
4.25 4.24 4.23 4.22 4.22 4.21 4.21 4.18 4.18 4.17 4.17 4.16 4.15 2.92 2.85 2.84 2.84 2.83 2.83 2.82 2.82 2.53 2.52 2.51 2.51 2.50 2.49 2.48 2.48 2.30 2.29 2.28 2.27 2.26 2.25 2.24 2.00 2.00 1.99 1.89 1.88 1.87 1.86 1.85 1.84 1.83 1.30 1.28 1.27 1.26 1.24
EtO
O
O
OEt
OH
5*S*-1 = 5*RS*
4.69
0.85
0.95
1.96
1.00
0.86
1.06
5.89
f1 (ppm)

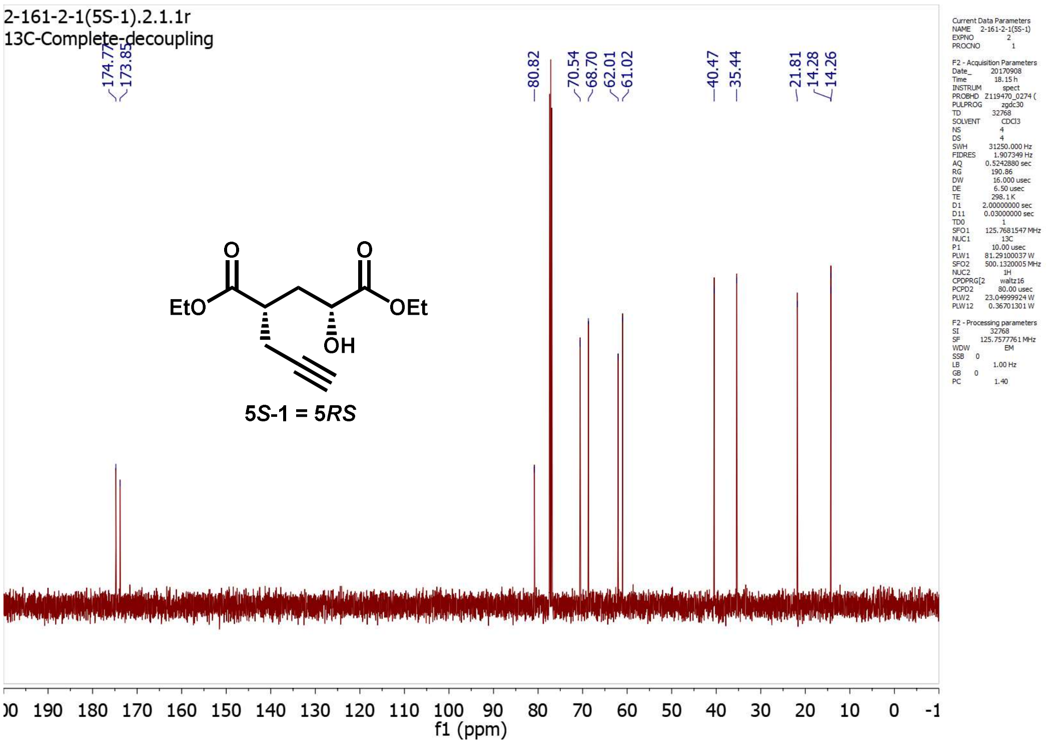

2-161-2-1(5S-1).2.1.1r
13C-Complete-decoupling
174.77 173.85
80.82
70.54 68.70 62.01 61.02
40.47 35.44
21.81 14.28 14.26
EtO
O
O
OEt
OH
5*S*-1 = 5*RS*
f1 (ppm)

**5*S*-2 (5*RR*)**

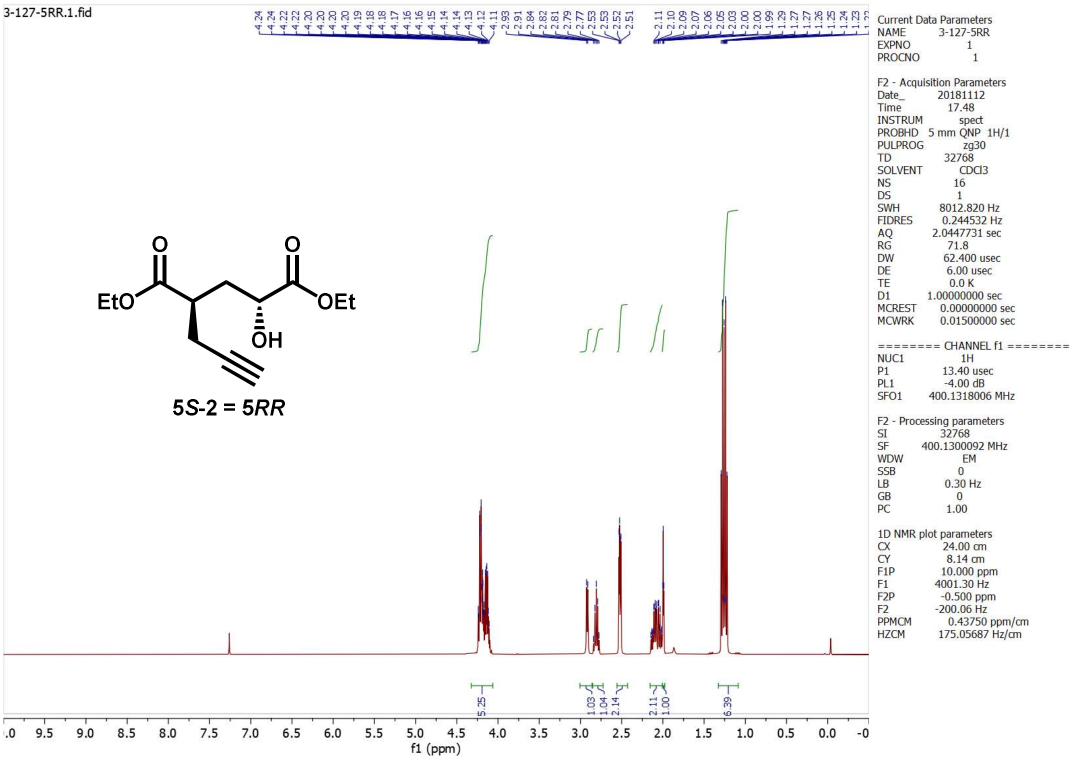
3-127-5RR.1.fid
5S-2 = 5RR
Current Data Parameters
NAME 3-127-5RR
EXPNO 1
PROCNO 1
F2 - Acquisition Parameters
Date_ 20181112
Time 17.48
INSTRUM spect
PROBHD 5 mm QNP 1H/1
PULPROG zg30
TD 32768
SOLVENT CDCl3
NS 16
DS 1
SWH 8012.820 Hz
FIDRES 0.244532 Hz
AQ 2.0447731 sec
RG 71.8
DW 62.400 usec
DE 6.00 usec
TE 0.0 K
D1 1.00000000 sec
MCREST 0.00000000 sec
MCWRK 0.01500000 sec
======== CHANNEL f1 ========
NUC1 1H
P1 13.40 usec
PL1 -4.00 dB
SFO1 400.1318006 MHz
F2 - Processing parameters
SI 32768
SF 400.1300092 MHz
WDW EM
SSB 0
LB 0.30 Hz
GB 0
PC 1.00
1D NMR plot parameters
CX 24.00 cm
CY 8.14 cm
F1P 10.000 ppm
F1 4001.30 Hz
F2P -0.500 ppm
F2 -200.06 Hz
PPMCM 0.43750 ppm/cm
HZCM 175.05687 Hz/cm
f1 (ppm)

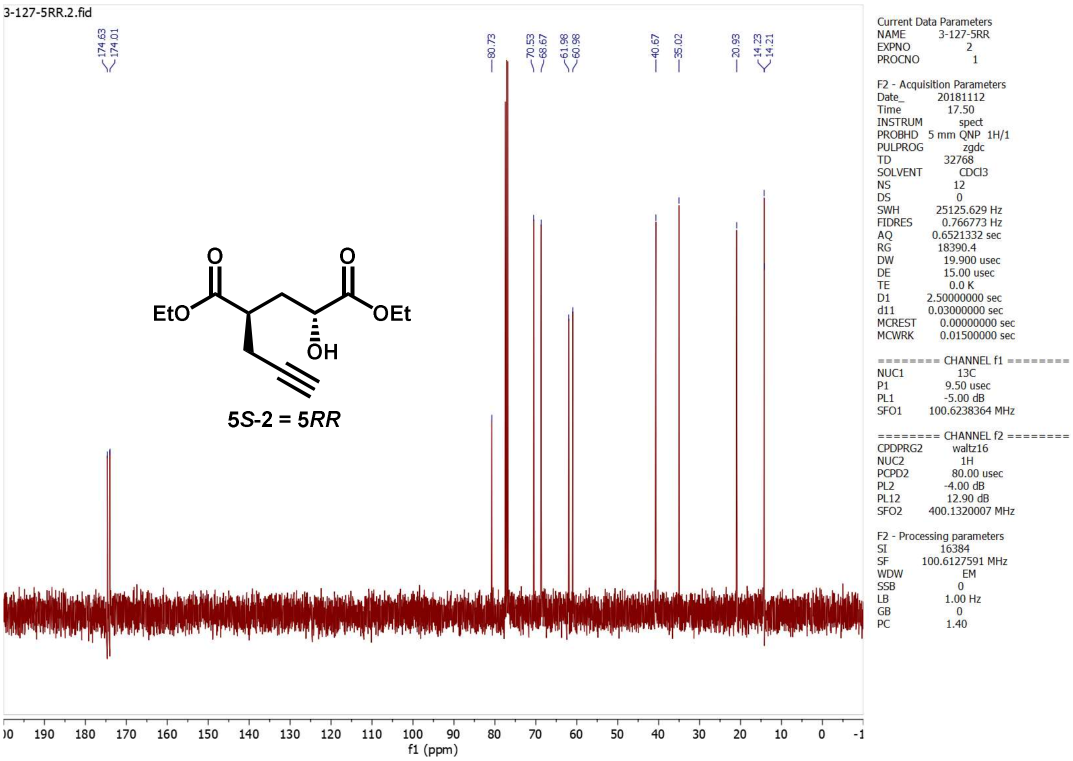
3-127-5RR.2.fid
174.63
174.01
80.73
70.53
68.67
61.98
60.98
40.67
35.02
20.93
14.23
14.21
5S-2 = 5RR
Current Data Parameters
NAME 3-127-5RR
EXPNO 2
PROCNO 1
F2 - Acquisition Parameters
Date_ 20181112
Time 17.50
INSTRUM spect
PROBHD 5 mm QNP 1H/1
PULPROG zgdc
TD 32768
SOLVENT CDCl3
NS 12
DS 0
SWH 25125.629 Hz
FIDRES 0.766773 Hz
AQ 0.6521332 sec
RG 18390.4
DW 19.900 usec
DE 15.00 usec
TE 0.0 K
D1 2.50000000 sec
d11 0.03000000 sec
MCREST 0.00000000 sec
MCWRK 0.01500000 sec
======== CHANNEL f1 ========
NUC1 13C
P1 9.50 usec
PL1 -5.00 dB
SFO1 100.6238364 MHz
======== CHANNEL f2 ========
CPDPRG2 waltz16
NUC2 1H
PCPD2 80.00 usec
PL2 -4.00 dB
PL12 12.90 dB
SFO2 400.1320007 MHz
F2 - Processing parameters
SI 16384
SF 100.6127591 MHz
WDW EM
SSB 0
LB 1.00 Hz
GB 0
PC 1.40
f1 (ppm)

**5*R*-1 (5*SR*)**

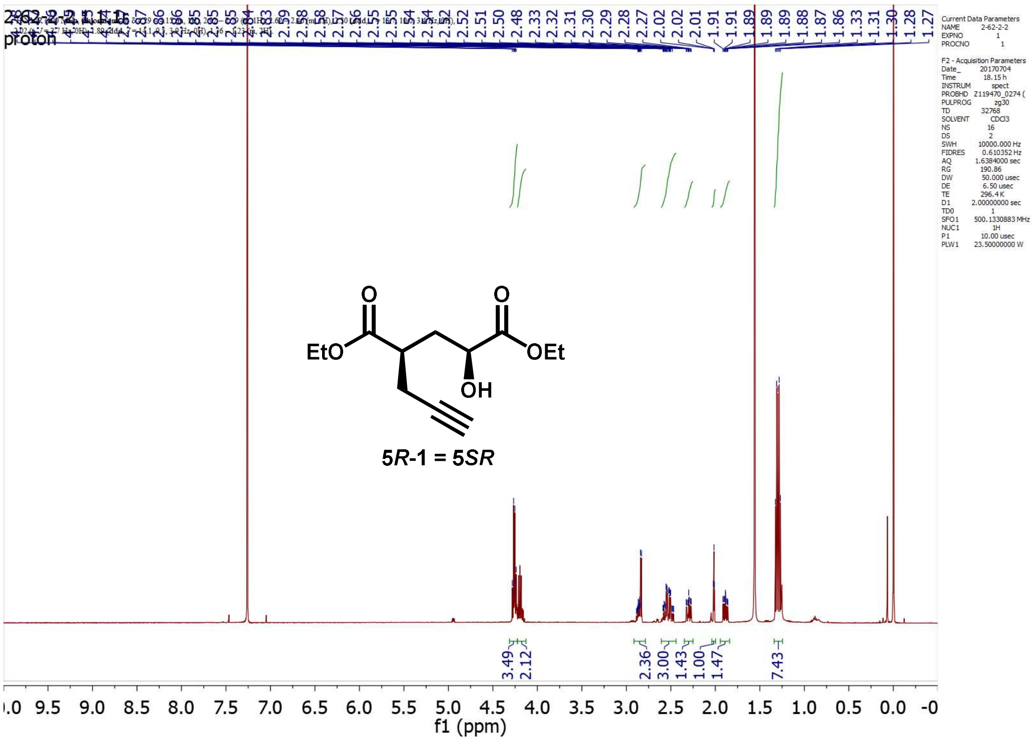
proton
Current Data Parameters
NAME 2-62-2-2
EXPNO 1
PROCNO 1
F2 - Acquisition Parameters
Date_ 20170704
Time 18.15 h
INSTRUM spect
PROBHD Z119470_0274 (
PULPROG zg30
TD 32768
SOLVENT CDCl3
NS 16
DS 2
SWH 10000.000 Hz
FIDRES 0.610352 Hz
AQ 1.6384000 sec
RG 190.86
DW 50.000 usec
DE 6.50 usec
TE 296.4 K
D1 2.00000000 sec
TD0 1
SFO1 500.1330883 MHz
NUC1 1H
P1 10.00 usec
PLW1 23.50000000 W
EtO
OEt
OH
5*R*-1 = 5*SR*
3.49
2.12
2.36
3.00
1.43
1.00
1.47
7.43
f1 (ppm)

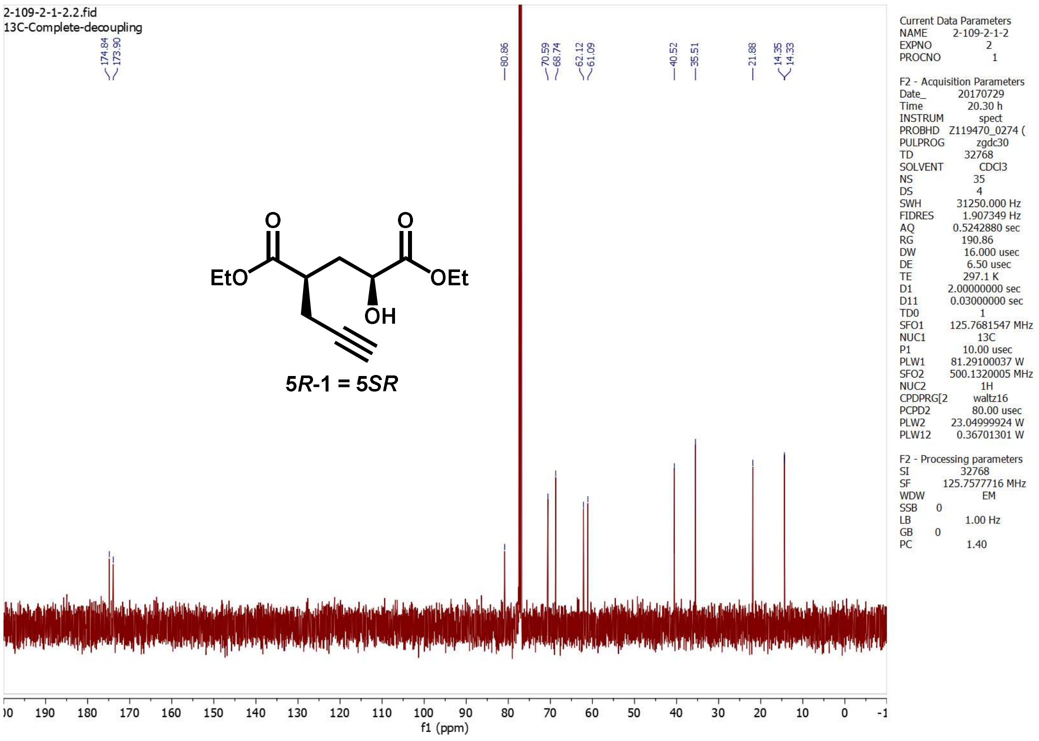
2-109-2-1-2.2.fid
13C-Complete-decoupling
174.84
173.90
80.86
70.59
68.74
62.12
61.09
40.52
35.51
21.88
14.35
14.33
Current Data Parameters
NAME 2-109-2-1-2
EXPNO 2
PROCNO 1
F2 - Acquisition Parameters
Date_ 20170729
Time 20.30 h
INSTRUM spect
PROBHD Z119470_0274 (
PULPROG zgdc30
TD 32768
SOLVENT CDCl3
NS 35
DS 4
SWH 31250.000 Hz
FIDRES 1.907349 Hz
AQ 0.5242880 sec
RG 190.86
DW 16.000 usec
DE 6.50 usec
TE 297.1 K
D1 2.00000000 sec
D11 0.03000000 sec
TD0 1
SFO1 125.7681547 MHz
NUC1 13C
P1 10.00 usec
PLW1 81.29100037 W
SFO2 500.1320005 MHz
NUC2 1H
CPDPRG[2 waltz16
PCPD2 80.00 usec
PLW2 23.04999924 W
PLW12 0.36701301 W
F2 - Processing parameters
SI 32768
SF 125.7577716 MHz
WDW EM
SSB 0
LB 1.00 Hz
GB 0
PC 1.40
EtO
OEt
OH
5*R*-1 = 5*SR*
f1 (ppm)

**5*R*-2 (5*SS*)**

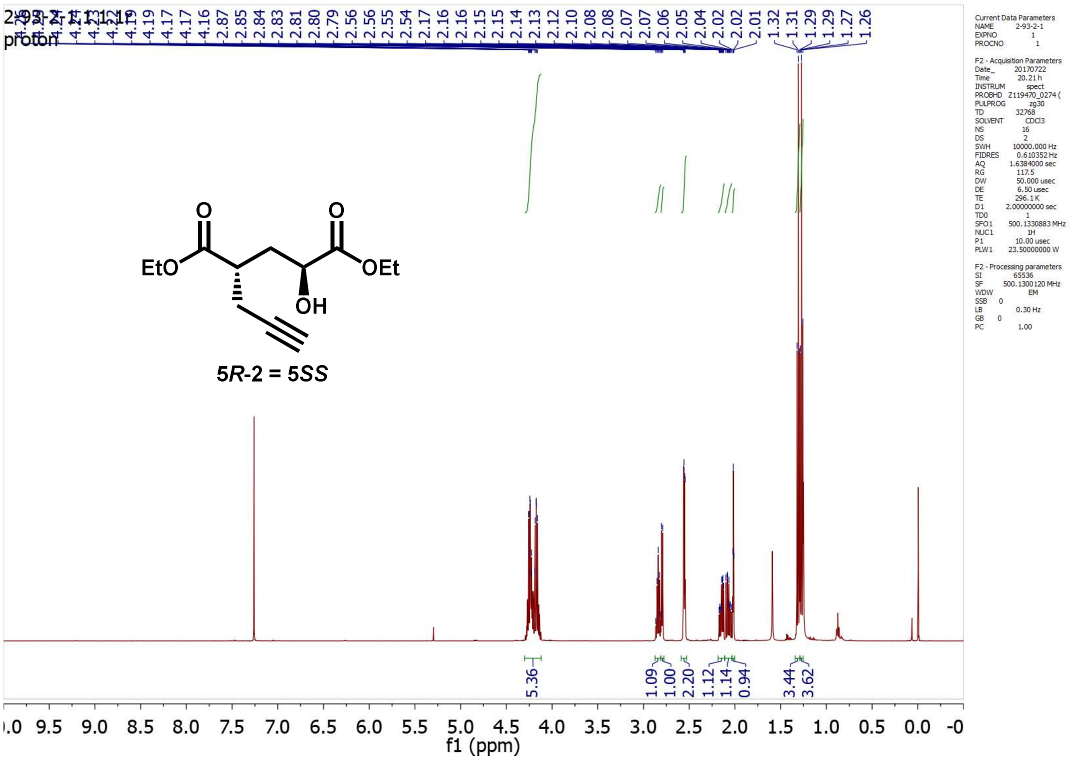

2-93-2-1.1.1r
proton
5.36
1.09
1.00
2.20
1.12
1.14
0.94
3.44
3.62
f1 (ppm)
EtO
O
O
OEt
OH
5R-2 = 5SS

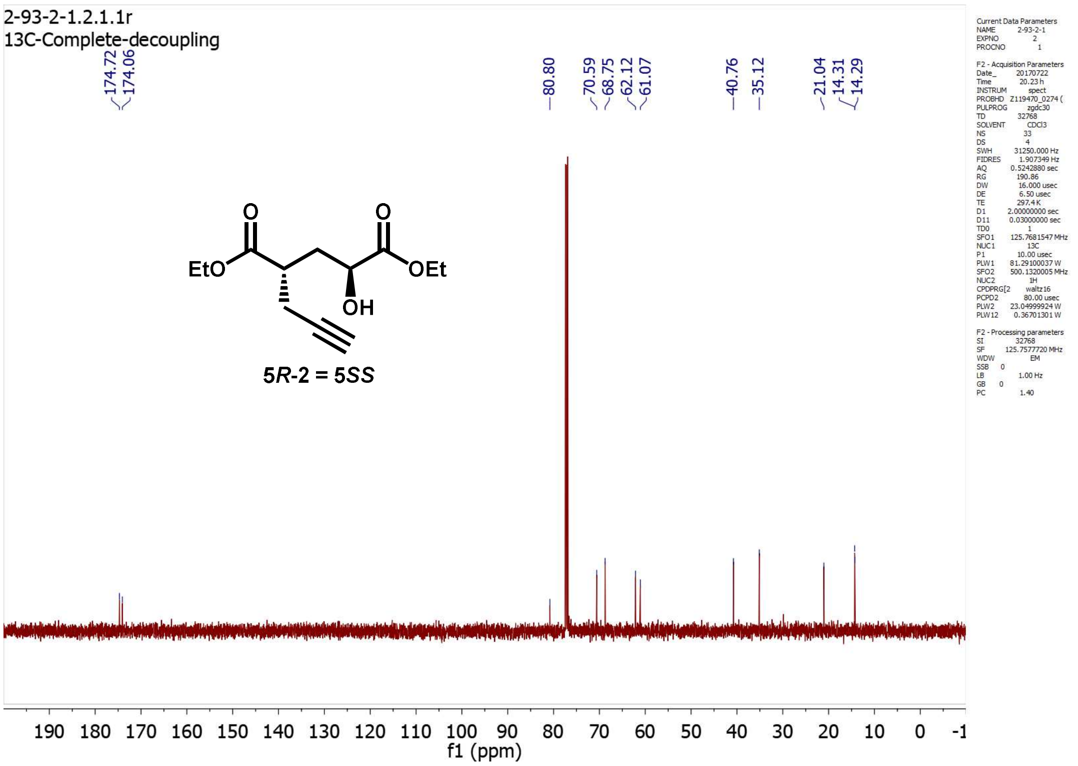

2-93-2-1.2.1.1r
13C-Complete-decoupling
174.72
174.06
80.80
70.59
68.75
62.12
61.07
40.76
35.12
21.04
14.31
14.29
f1 (ppm)
EtO
O
O
OEt
OH
5R-2 = 5SS

**5.9*R*-1**

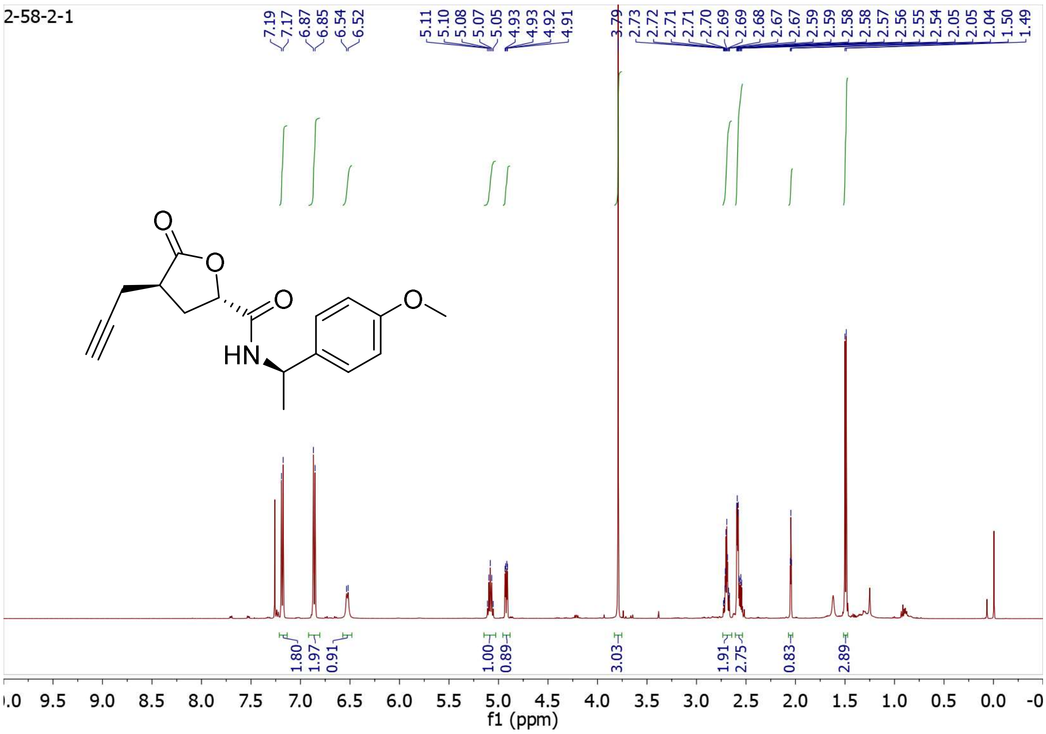
2-58-2-1
7.19
7.17
6.87
6.85
6.54
6.52
5.11
5.10
5.08
5.07
5.05
4.93
4.93
4.92
4.91
3.79
2.73
2.72
2.71
2.71
2.70
2.69
2.69
2.68
2.67
2.67
2.59
2.59
2.58
2.58
2.57
2.56
2.55
2.54
2.05
2.05
2.04
1.50
1.49
O
O
O
HN
O
1.80
1.97
0.91
1.00
0.89
3.03
1.91
2.75
0.83
2.89
f1 (ppm)

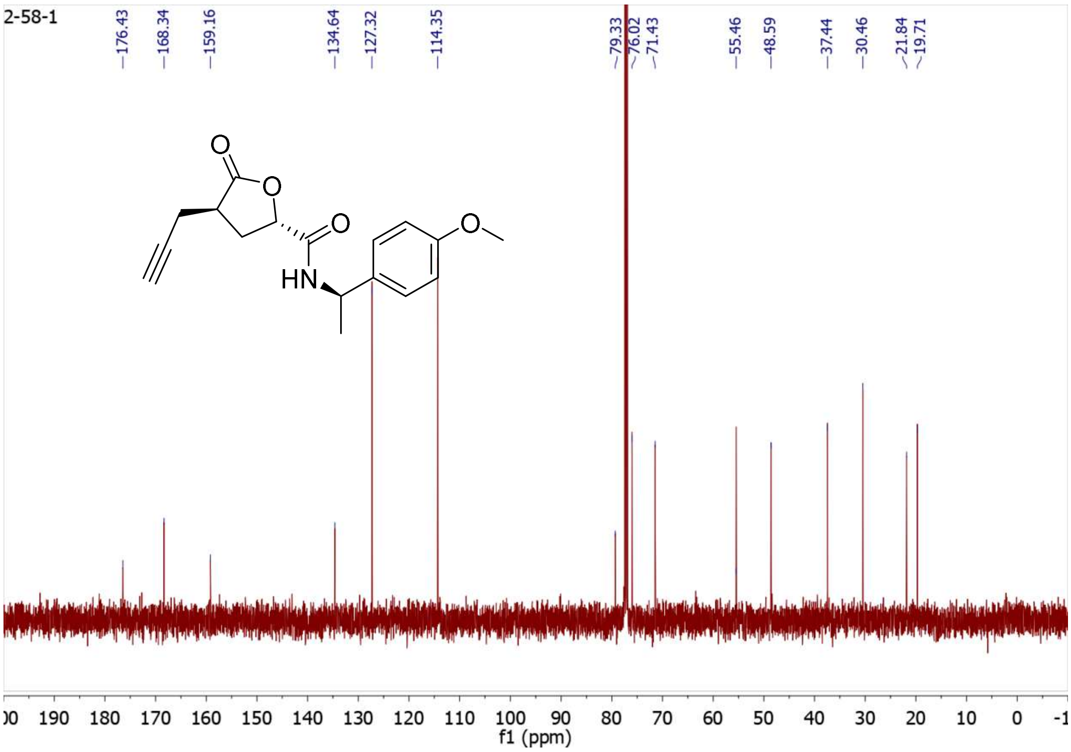
2-58-1
176.43
168.34
159.16
134.64
127.32
114.35
79.33
76.02
71.43
55.46
48.59
37.44
30.46
21.84
19.71
O
O
O
HN
O
f1 (ppm)

**5.9*R*-2**

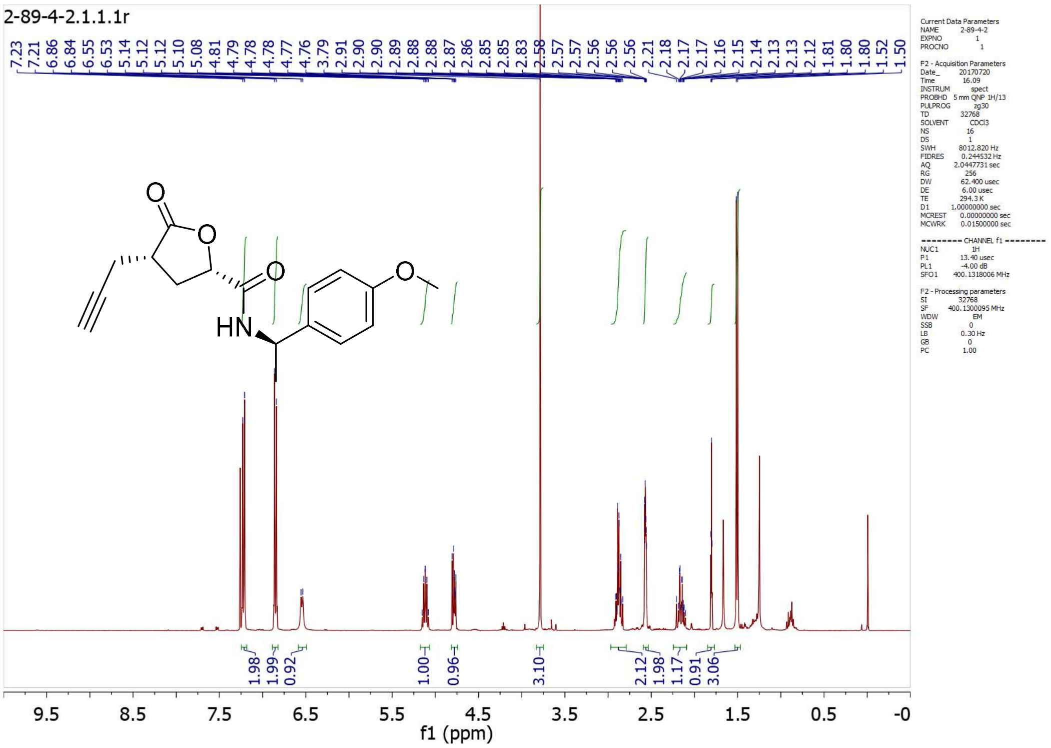

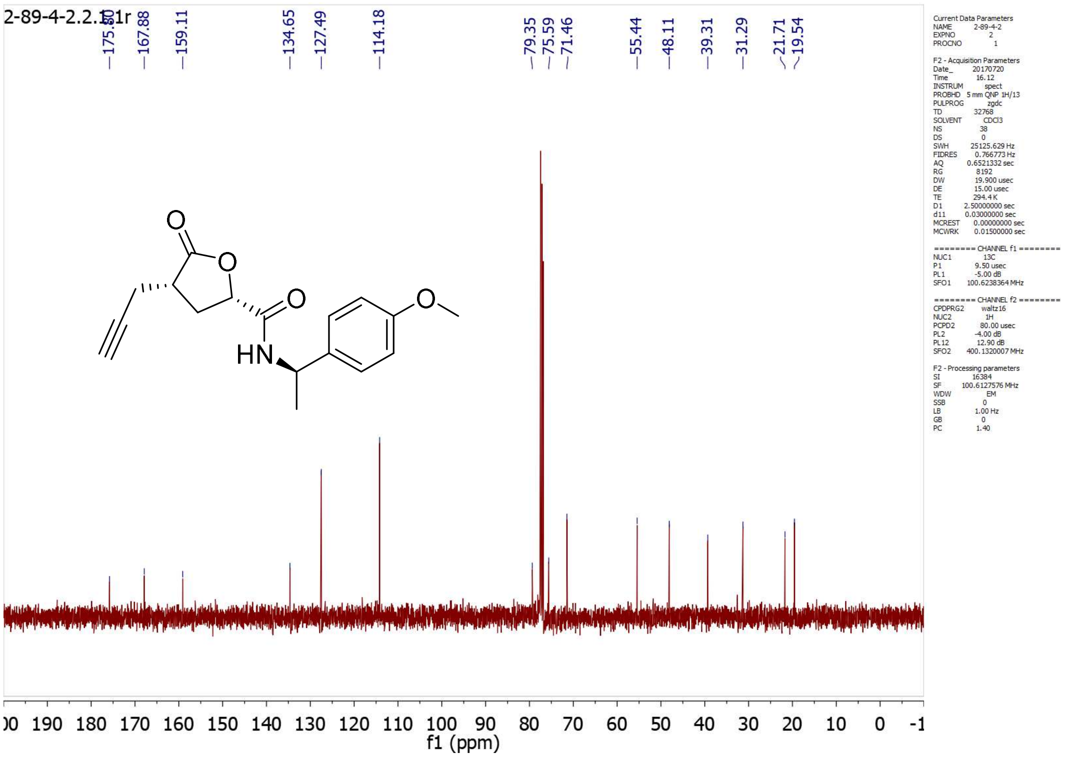

**C5-2**

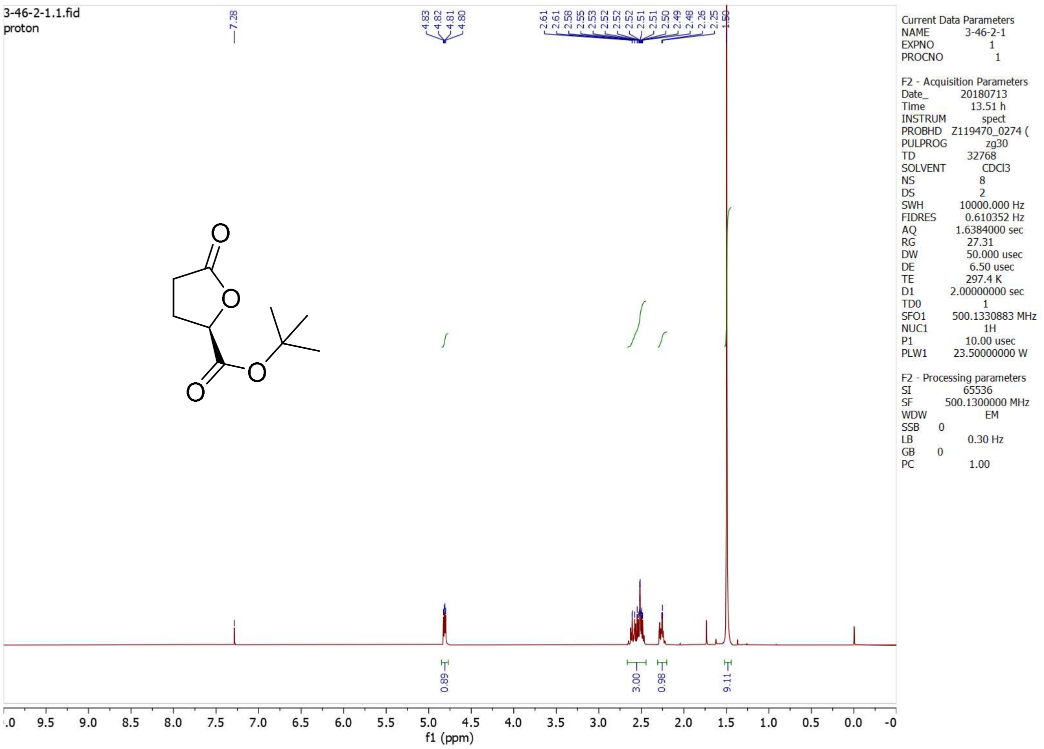

3-46-2-1.1.fid
proton
7.28
4.83
4.82
4.81
4.80
2.61
2.61
2.58
2.55
2.53
2.52
2.52
2.52
2.51
2.51
2.50
2.49
2.48
2.26
2.25
1.50
0.89
3.00
0.98
9.11
f1 (ppm)
Current Data Parameters
NAME 3-46-2-1
EXPNO 1
PROCNO 1
F2 - Acquisition Parameters
Date_ 20180713
Time 13.51 h
INSTRUM spect
PROBHD Z119470_0274 (
PULPROG zg30
TD 32768
SOLVENT CDCl3
NS 8
DS 2
SWH 10000.000 Hz
FIDRES 0.610352 Hz
AQ 1.6384000 sec
RG 27.31
DW 50.000 usec
DE 6.50 usec
TE 297.4 K
D1 2.00000000 sec
TD0 1
SFO1 500.1330883 MHz
NUC1 1H
P1 10.00 usec
PLW1 23.50000000 W
F2 - Processing parameters
SI 65536
SF 500.1300000 MHz
WDW EM
SSB 0
LB 0.30 Hz
GB 0
PC 1.00

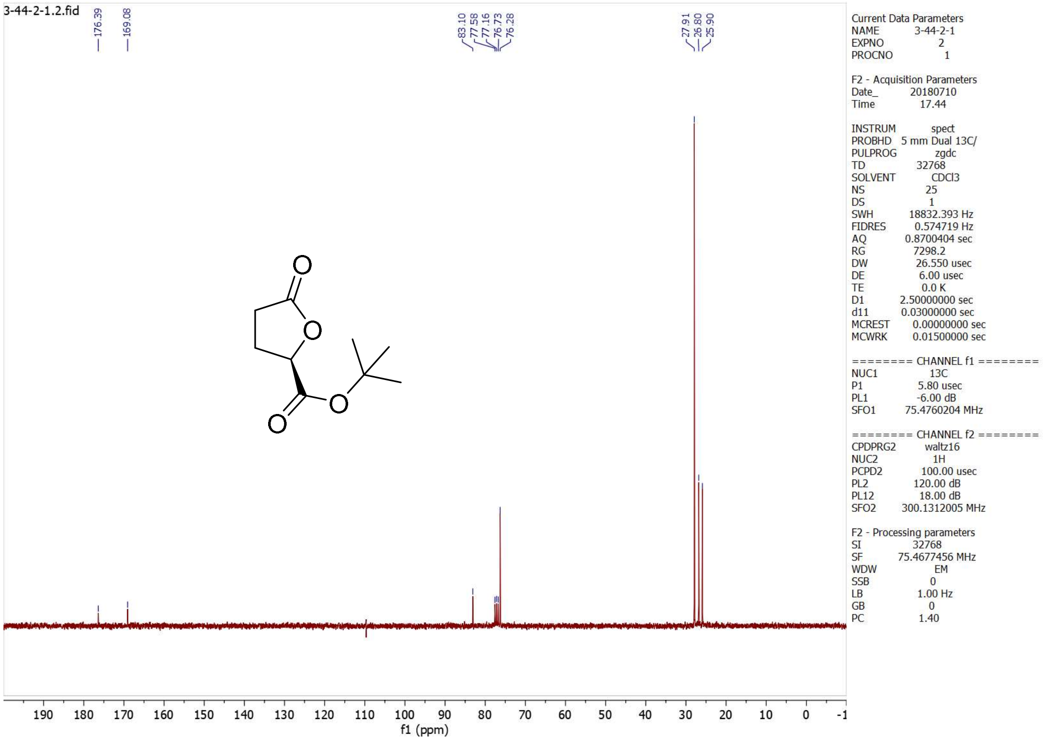

3-44-2-1.1.2.fid
176.39
169.08
83.10
77.58
77.16
76.73
76.28
27.91
26.80
25.90
f1 (ppm)
Current Data Parameters
NAME 3-44-2-1
EXPNO 2
PROCNO 1
F2 - Acquisition Parameters
Date_ 20180710
Time 17.44
INSTRUM spect
PROBHD 5 mm Dual 13C/
PULPROG zgdc
TD 32768
SOLVENT CDCl3
NS 25
DS 1
SWH 18832.393 Hz
FIDRES 0.574719 Hz
AQ 0.8700404 sec
RG 7298.2
DW 26.550 usec
DE 6.00 usec
TE 0.0 K
D1 2.50000000 sec
d11 0.03000000 sec
MCREST 0.00000000 sec
MCWRK 0.01500000 sec
======== CHANNEL f1 ========
NUC1 13C
P1 5.80 usec
PL1 -6.00 dB
SFO1 75.4760204 MHz
======== CHANNEL f2 ========
CPDPRG2 waltz16
NUC2 1H
PCPD2 100.00 usec
PL2 120.00 dB
PL12 18.00 dB
SFO2 300.1312005 MHz
F2 - Processing parameters
SI 32768
SF 75.4677456 MHz
WDW EM
SSB 0
LB 1.00 Hz
GB 0
PC 1.40

**C5-3**

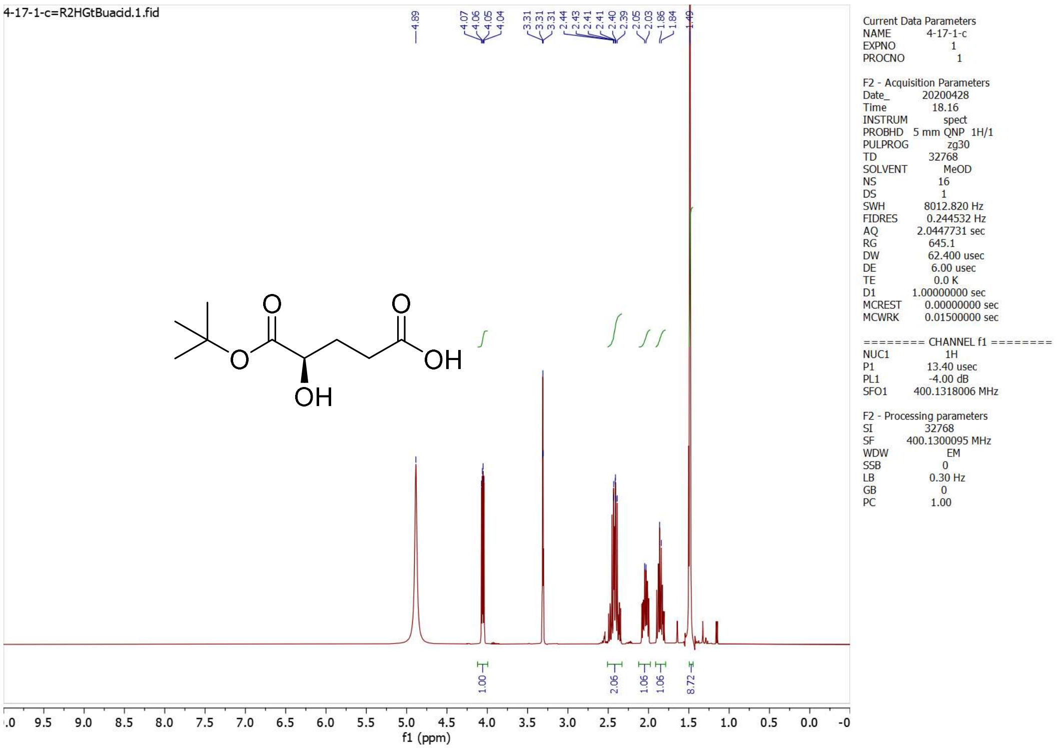
4-17-1-c=R2HGtBuacid.1.fid
4.89
4.07 4.06 4.05 4.04
3.31 3.31 3.31
2.44 2.43 2.41 2.41 2.40 2.39
2.05 2.03
1.86 1.84
1.49
1.00
2.06
1.06
1.06
8.72
f1 (ppm)
O
O
OH
OH
O
Current Data Parameters
NAME 4-17-1-c
EXPNO 1
PROCNO 1
F2 - Acquisition Parameters
Date_ 20200428
Time 18.16
INSTRUM spect
PROBHD 5 mm QNP 1H/1
PULPROG zg30
TD 32768
SOLVENT MeOD
NS 16
DS 1
SWH 8012.820 Hz
FIDRES 0.244532 Hz
AQ 2.0447731 sec
RG 645.1
DW 62.400 usec
DE 6.00 usec
TE 0.0 K
D1 1.00000000 sec
MCREST 0.00000000 sec
MCWRK 0.01500000 sec
======== CHANNEL f1 ========
NUC1 1H
P1 13.40 usec
PL1 -4.00 dB
SFO1 400.1318006 MHz
F2 - Processing parameters
SI 32768
SF 400.1300095 MHz
WDW EM
SSB 0
LB 0.30 Hz
GB 0
PC 1.00

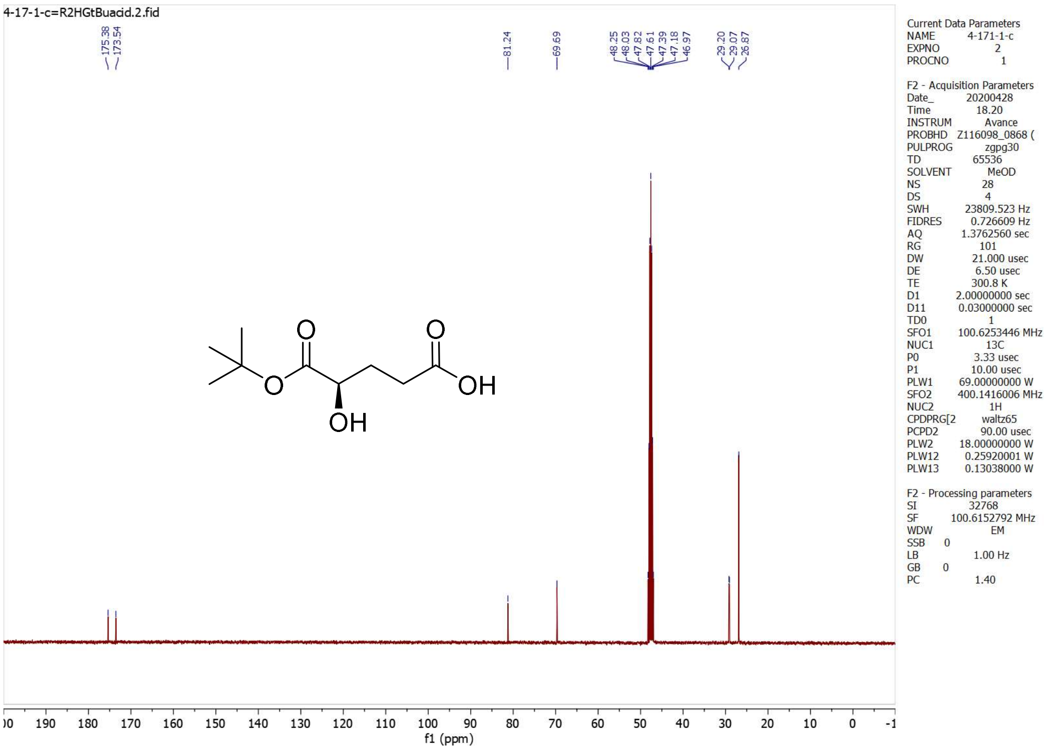
4-17-1-c=R2HGtBuacid.2.fid
175.38 173.54
81.24
69.69
48.25 48.03 47.82 47.61 47.39 47.18 46.97
29.20 29.07 26.87
f1 (ppm)
O
O
OH
OH
O
Current Data Parameters
NAME 4-171-1-c
EXPNO 2
PROCNO 1
F2 - Acquisition Parameters
Date_ 20200428
Time 18.20
INSTRUM Avance
PROBHD Z116098_0868 (
PULPROG zgpg30
TD 65536
SOLVENT MeOD
NS 28
DS 4
SWH 23809.523 Hz
FIDRES 0.726609 Hz
AQ 1.3762560 sec
RG 101
DW 21.000 usec
DE 6.50 usec
TE 300.8 K
D1 2.00000000 sec
D11 0.03000000 sec
TD0 1
SFO1 100.6253446 MHz
NUC1 13C
P0 3.33 usec
P1 10.00 usec
PLW1 69.00000000 W
SFO2 400.1416006 MHz
NUC2 1H
CPDPRG[2 waltz65
PCPD2 90.00 usec
PLW2 18.00000000 W
PLW12 0.25920001 W
PLW13 0.13038000 W
F2 - Processing parameters
SI 32768
SF 100.6152792 MHz
WDW EM
SSB 0
LB 1.00 Hz
GB 0
PC 1.40

**C5-4**

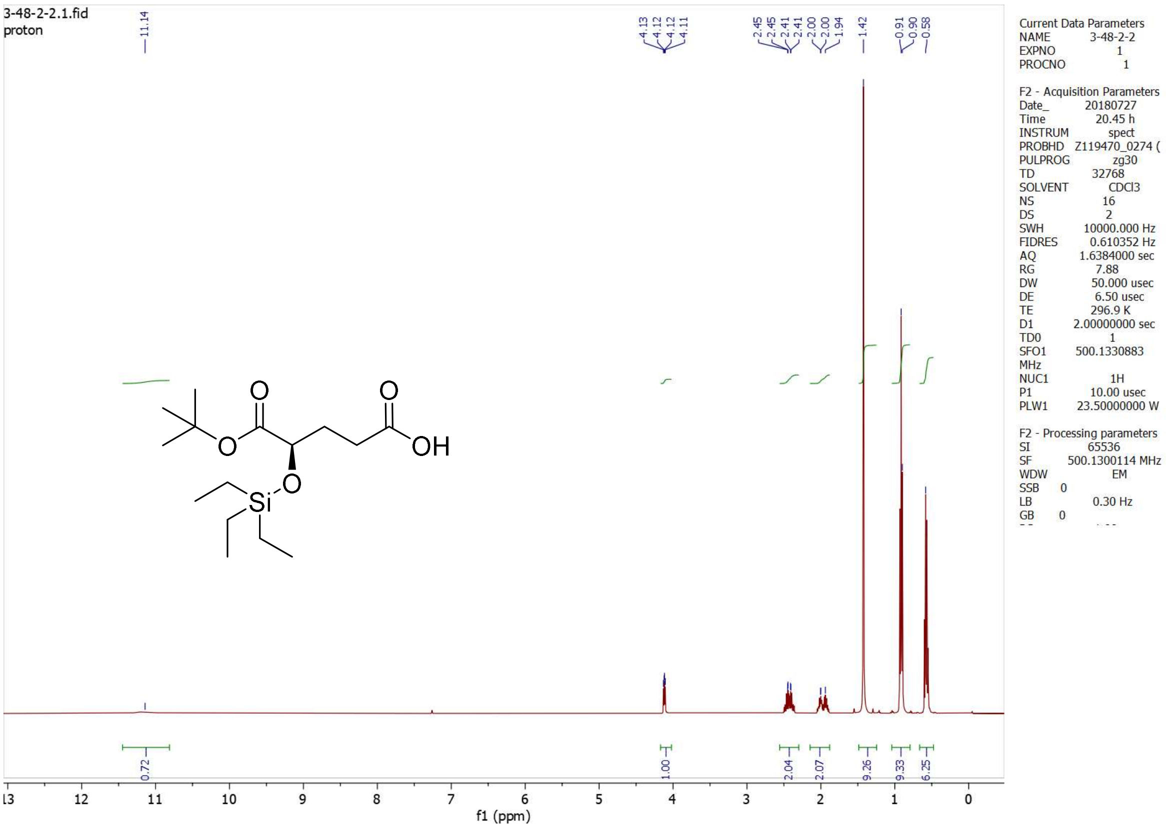
3-48-2-2.1.fid
proton
11.14
4.13
4.12
4.12
4.11
2.45
2.45
2.41
2.41
2.00
2.00
1.94
1.42
0.91
0.90
0.58
O
O
OH
O
Si
O
0.72
1.00
2.04
2.07
9.26
9.33
6.25
13 12 11 10 9 8 7 6 5 4 3 2 1 0
f1 (ppm)
Current Data Parameters
NAME 3-48-2-2
EXPNO 1
PROCNO 1
F2 - Acquisition Parameters
Date_ 20180727
Time 20.45 h
INSTRUM spect
PROBHD Z119470_0274 (
PULPROG zg30
TD 32768
SOLVENT CDCl3
NS 16
DS 2
SWH 10000.000 Hz
FIDRES 0.610352 Hz
AQ 1.6384000 sec
RG 7.88
DW 50.000 usec
DE 6.50 usec
TE 296.9 K
D1 2.00000000 sec
TD0 1
SFO1 500.1330883 MHz
NUC1 1H
P1 10.00 usec
PLW1 23.50000000 W
F2 - Processing parameters
SI 65536
SF 500.1300114 MHz
WDW EM
SSB 0
LB 0.30 Hz
GB 0

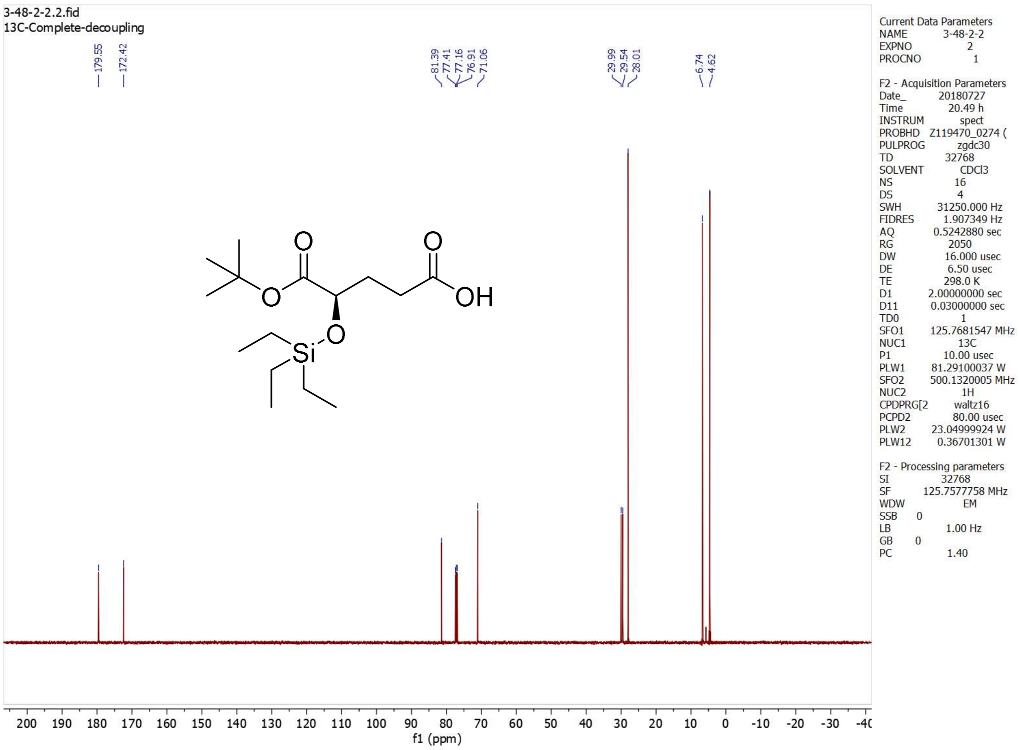
3-48-2-2.2.fid
13C-Complete-decoupling
179.55
172.42
81.39
77.41
77.16
76.91
71.06
29.99
29.54
28.01
6.74
4.62
O
O
O
OH
O
Si
200 190 180 170 160 150 140 130 120 110 100 90 80 70 60 50 40 30 20 10 0 -10 -20 -30 -40
f1 (ppm)
Current Data Parameters
NAME 3-48-2-2
EXPNO 2
PROCNO 1
F2 - Acquisition Parameters
Date_ 20180727
Time 20.49 h
INSTRUM spect
PROBHD Z119470_0274 (
PULPROG zgdc30
TD 32768
SOLVENT CDCl3
NS 16
DS 4
SWH 31250.000 Hz
FIDRES 1.907349 Hz
AQ 0.5242880 sec
RG 2050
DW 16.000 usec
DE 6.50 usec
TE 298.0 K
D1 2.00000000 sec
D11 0.03000000 sec
TD0 1
SFO1 125.7681547 MHz
NUC1 13C
P1 10.00 usec
PLW1 81.29100037 W
SFO2 500.1320005 MHz
NUC2 1H
CPDPRG[2 waltz16
PCPD2 80.00 usec
PLW2 23.04999924 W
PLW12 0.36701301 W
F2 - Processing parameters
SI 32768
SF 125.7577758 MHz
WDW EM
SSB 0
LB 1.00 Hz
GB 0
PC 1.40

#### C5-donor

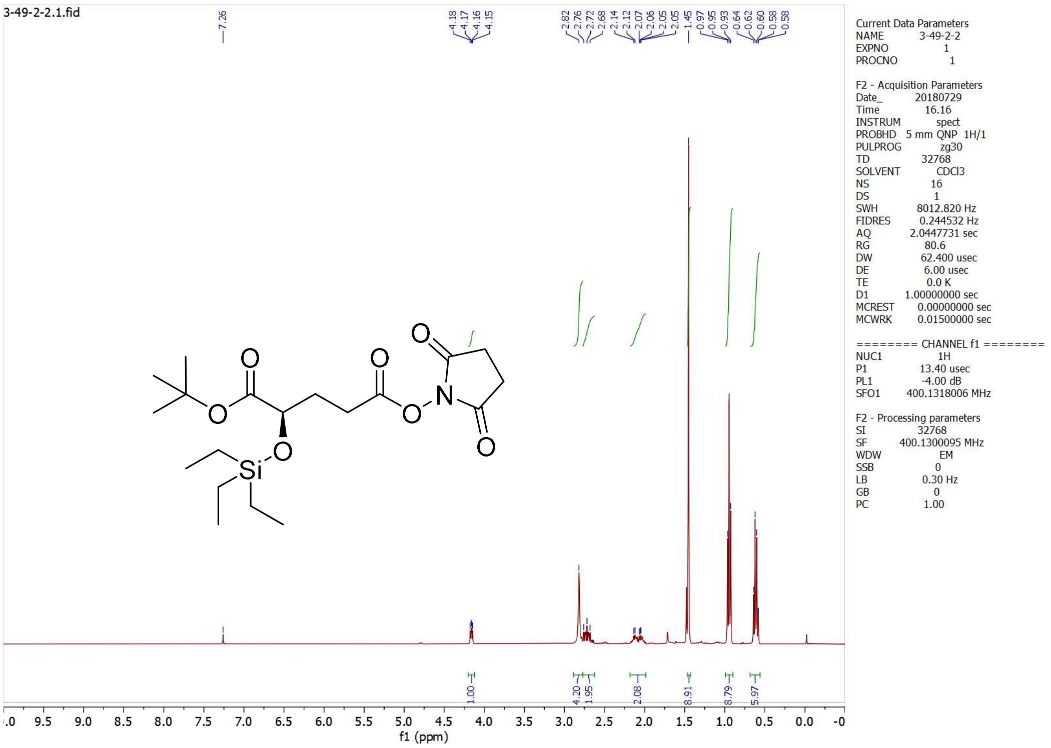

3-49-2-2.1.fid
7.26
4.18
4.17
4.16
4.15
2.82
2.76
2.72
2.68
2.14
2.12
2.07
2.06
2.05
2.05
1.45
0.97
0.95
0.93
0.64
0.62
0.60
0.58
0.58
1.00
4.20
1.95
2.08
8.91
8.79
5.97
O
O
O
O
O
O
N
O
Si
f1 (ppm)
Current Data Parameters
NAME 3-49-2-2
EXPNO 1
PROCNO 1
F2 - Acquisition Parameters
Date_ 20180729
Time 16.16
INSTRUM spect
PROBHD 5 mm QNP 1H/1
PULPROG zg30
TD 32768
SOLVENT CDCl3
NS 16
DS 1
SWH 8012.820 Hz
FIDRES 0.244532 Hz
AQ 2.0447731 sec
RG 80.6
DW 62.400 usec
DE 6.00 usec
TE 0.0 K
D1 1.00000000 sec
MCREST 0.00000000 sec
MCWRK 0.01500000 sec
======== CHANNEL f1 ========
NUC1 1H
P1 13.40 usec
PL1 -4.00 dB
SFO1 400.1318006 MHz
F2 - Processing parameters
SI 32768
SF 400.1300095 MHz
WDW EM
SSB 0
LB 0.30 Hz
GB 0
PC 1.00

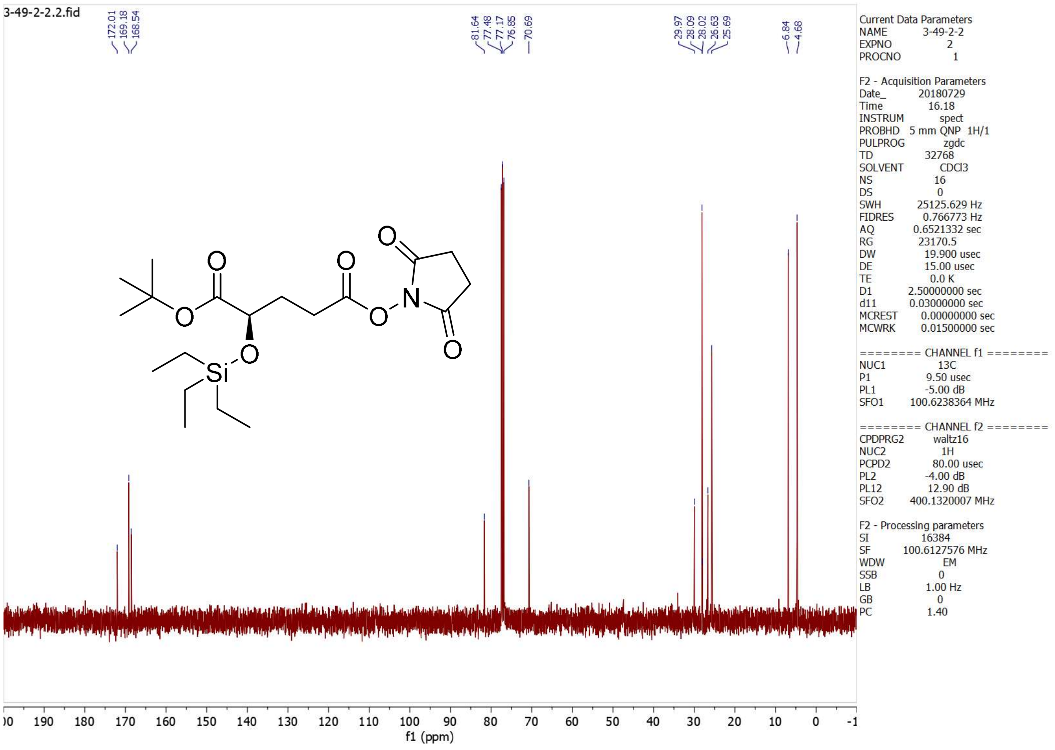

3-49-2-2.2.fid
172.01
169.18
168.54
81.64
77.48
77.17
76.85
70.69
29.97
28.09
28.02
26.63
25.69
6.84
4.68
O
O
O
O
O
O
N
O
Si
f1 (ppm)
Current Data Parameters
NAME 3-49-2-2
EXPNO 2
PROCNO 1
F2 - Acquisition Parameters
Date_ 20180729
Time 16.18
INSTRUM spect
PROBHD 5 mm QNP 1H/1
PULPROG zgdc
TD 32768
SOLVENT CDCl3
NS 16
DS 0
SWH 25125.629 Hz
FIDRES 0.766773 Hz
AQ 0.6521332 sec
RG 23170.5
DW 19.900 usec
DE 15.00 usec
TE 0.0 K
D1 2.50000000 sec
d11 0.03000000 sec
MCREST 0.00000000 sec
MCWRK 0.01500000 sec
======== CHANNEL f1 ========
NUC1 13C
P1 9.50 usec
PL1 -5.00 dB
SFO1 100.6238364 MHz
======== CHANNEL f2 ========
CPDPRG2 waltz16
NUC2 1H
PCPD2 80.00 usec
PL2 -4.00 dB
PL12 12.90 dB
SFO2 400.1320007 MHz
F2 - Processing parameters
SI 16384
SF 100.6127576 MHz
WDW EM
SSB 0
LB 1.00 Hz
GB 0
PC 1.40

**C1-2**

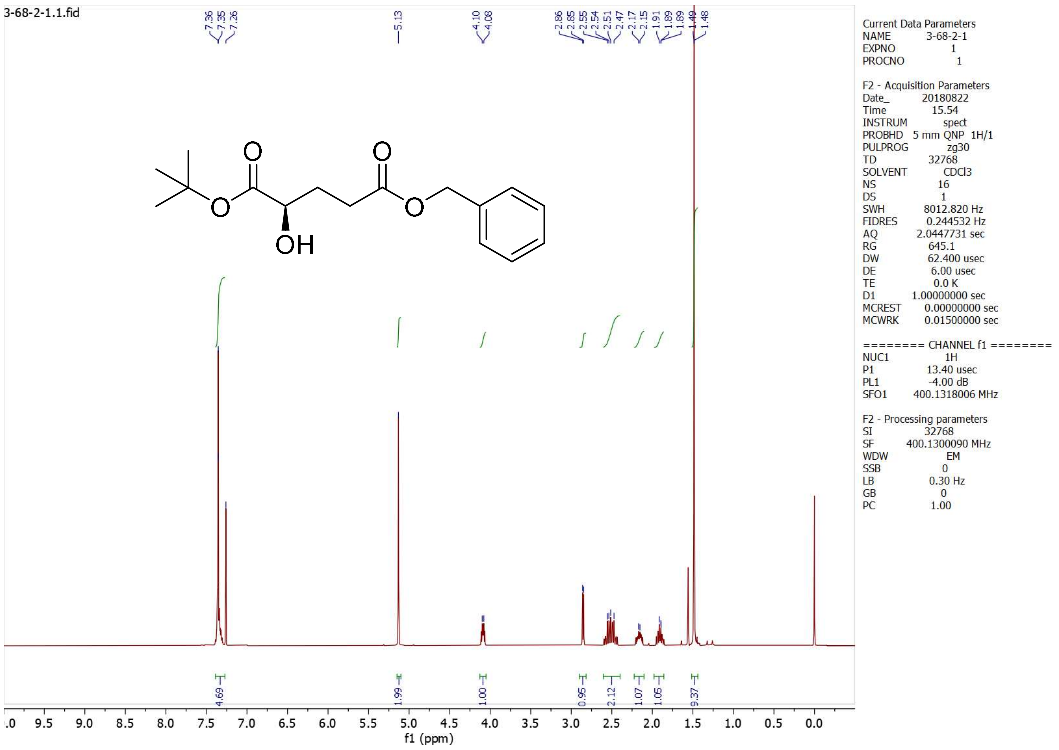
3-68-2-1.1.fid
7.36
7.35
7.26
5.13
4.10
4.08
2.86
2.85
2.55
2.54
2.51
2.47
2.17
2.15
1.91
1.89
1.89
1.49
1.48
OH
O
4.69
1.99
1.00
0.95
2.12
1.07
1.05
9.37
f1 (ppm)
Current Data Parameters
NAME 3-68-2-1
EXPNO 1
PROCNO 1
F2 - Acquisition Parameters
Date_ 20180822
Time 15.54
INSTRUM spect
PROBHD 5 mm QNP 1H/1
PULPROG zg30
TD 32768
SOLVENT CDCl3
NS 16
DS 1
SWH 8012.820 Hz
FIDRES 0.244532 Hz
AQ 2.0447731 sec
RG 645.1
DW 62.400 usec
DE 6.00 usec
TE 0.0 K
D1 1.00000000 sec
MCREST 0.00000000 sec
MCWRK 0.01500000 sec
======== CHANNEL f1 ========
NUC1 1H
P1 13.40 usec
PL1 -4.00 dB
SFO1 400.1318006 MHz
F2 - Processing parameters
SI 32768
SF 400.1300090 MHz
WDW EM
SSB 0
LB 0.30 Hz
GB 0
PC 1.00

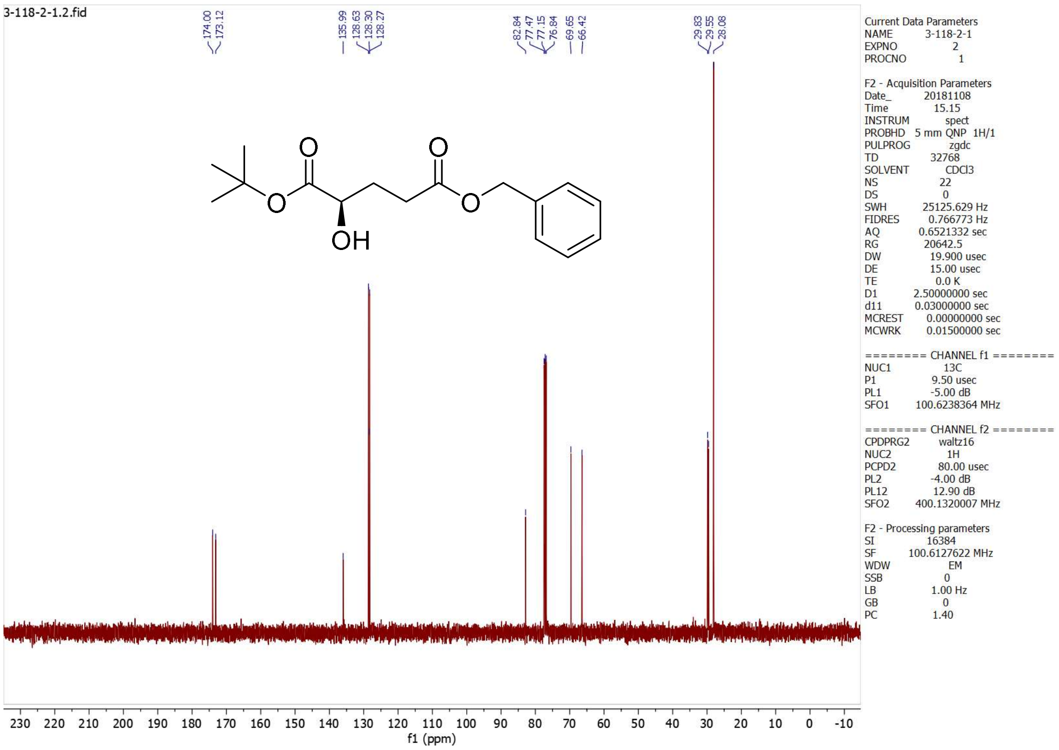
3-118-2-1.2.fid
174.00
173.12
135.99
128.63
128.30
128.27
82.84
77.47
77.15
76.84
69.65
66.42
29.83
29.55
28.08
OH
O
f1 (ppm)
Current Data Parameters
NAME 3-118-2-1
EXPNO 2
PROCNO 1
F2 - Acquisition Parameters
Date_ 20181108
Time 15.15
INSTRUM spect
PROBHD 5 mm QNP 1H/1
PULPROG zgdc
TD 32768
SOLVENT CDCl3
NS 22
DS 0
SWH 25125.629 Hz
FIDRES 0.766773 Hz
AQ 0.6521332 sec
RG 20642.5
DW 19.900 usec
DE 15.00 usec
TE 0.0 K
D1 2.50000000 sec
d11 0.03000000 sec
MCREST 0.00000000 sec
MCWRK 0.01500000 sec
======== CHANNEL f1 ========
NUC1 13C
P1 9.50 usec
PL1 -5.00 dB
SFO1 100.6238364 MHz
======== CHANNEL f2 ========
CPDPRG2 waltz16
NUC2 1H
PCPD2 80.00 usec
PL2 -4.00 dB
PL12 12.90 dB
SFO2 400.1320007 MHz
F2 - Processing parameters
SI 16384
SF 100.6127622 MHz
WDW EM
SSB 0
LB 1.00 Hz
GB 0
PC 1.40

## C1-3

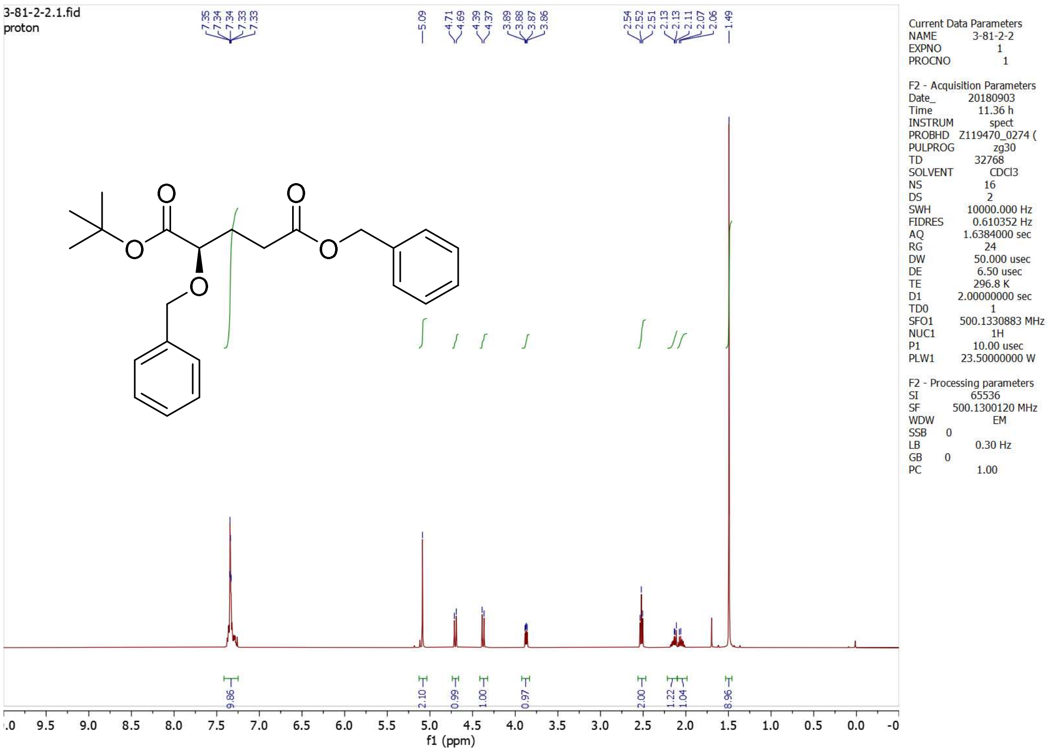
3-81-2-2.1.fid
proton
7.35
7.34
7.34
7.33
7.33
5.09
4.71
4.69
4.39
4.37
3.89
3.88
3.87
3.86
2.54
2.52
2.51
2.13
2.13
2.11
2.07
2.06
1.49
9.86
2.10
0.99
1.00
0.97
2.00
1.22
1.04
8.96
f1 (ppm)
Current Data Parameters
NAME 3-81-2-2
EXPNO 1
PROCNO 1
F2 - Acquisition Parameters
Date_ 20180903
Time 11.36 h
INSTRUM spect
PROBHD Z119470_0274 (
PULPROG zg30
TD 32768
SOLVENT CDCl3
NS 16
DS 2
SWH 10000.000 Hz
FIDRES 0.610352 Hz
AQ 1.6384000 sec
RG 24
DW 50.000 usec
DE 6.50 usec
TE 296.8 K
D1 2.00000000 sec
TD0 1
SFO1 500.1330883 MHz
NUC1 1H
P1 10.00 usec
PLW1 23.50000000 W
F2 - Processing parameters
SI 65536
SF 500.1300120 MHz
WDW EM
SSB 0
LB 0.30 Hz
GB 0
PC 1.00

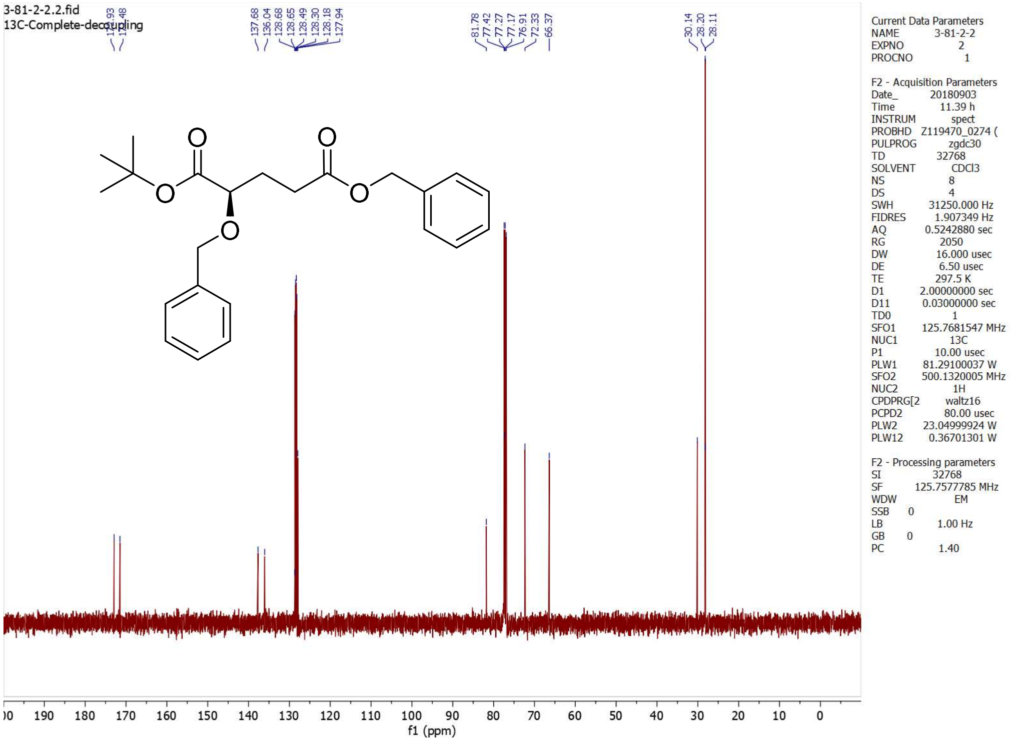
3-81-2-2.2.fid
13C-Complete-decoupling
137.68
136.04
128.68
128.65
128.49
128.30
128.18
127.94
81.78
77.42
77.27
77.17
76.91
72.33
66.37
30.14
28.20
28.11
f1 (ppm)
Current Data Parameters
NAME 3-81-2-2
EXPNO 2
PROCNO 1
F2 - Acquisition Parameters
Date_ 20180903
Time 11.39 h
INSTRUM spect
PROBHD Z119470_0274 (
PULPROG zgdc30
TD 32768
SOLVENT CDCl3
NS 8
DS 4
SWH 31250.000 Hz
FIDRES 1.907349 Hz
AQ 0.5242880 sec
RG 2050
DW 16.000 usec
DE 6.50 usec
TE 297.5 K
D1 2.00000000 sec
D11 0.03000000 sec
TD0 1
SFO1 125.7681547 MHz
NUC1 13C
P1 10.00 usec
PLW1 81.29100037 W
SFO2 500.1320005 MHz
NUC2 1H
CPDPRG[2 waltz16
PCPD2 80.00 usec
PLW2 23.04999924 W
PLW12 0.36701301 W
F2 - Processing parameters
SI 32768
SF 125.7577785 MHz
WDW EM
SSB 0
LB 1.00 Hz
GB 0
PC 1.40

#### C1-donor

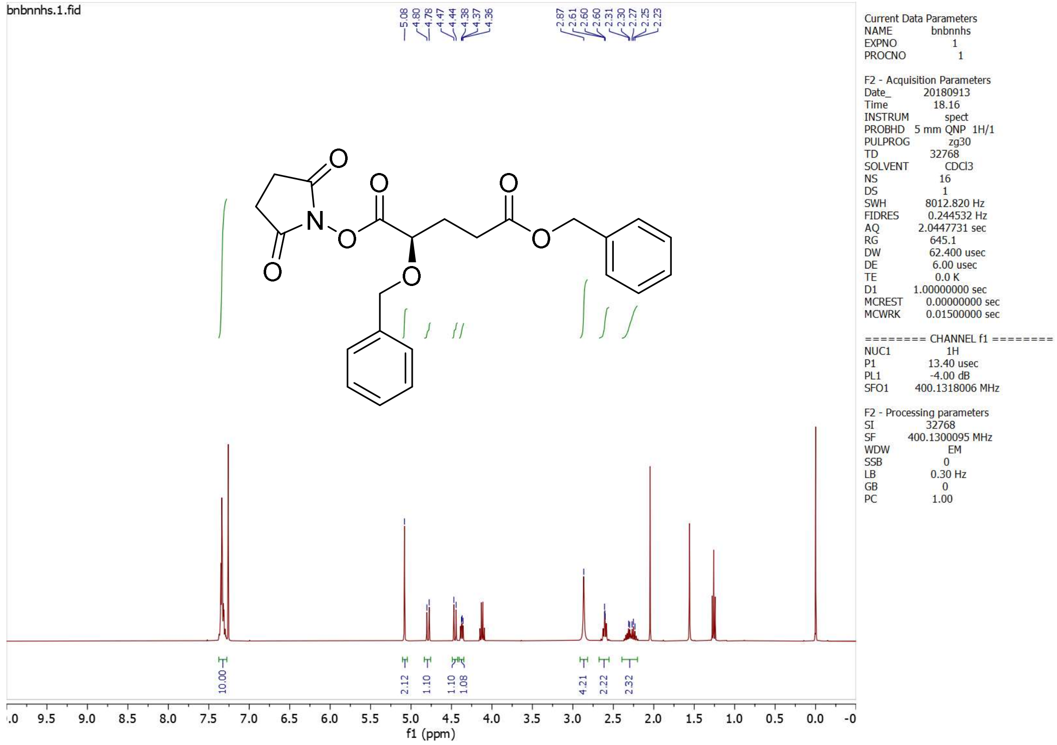
bnbnnhs.1.fid
Current Data Parameters
NAME bnbnnhs
EXPNO 1
PROCNO 1
F2 - Acquisition Parameters
Date_ 20180913
Time 18.16
INSTRUM spect
PROBHD 5 mm QNP 1H/1
PULPROG zg30
TD 32768
SOLVENT CDCl3
NS 16
DS 1
SWH 8012.820 Hz
FIDRES 0.244532 Hz
AQ 2.0447731 sec
RG 645.1
DW 62.400 usec
DE 6.00 usec
TE 0.0 K
D1 1.00000000 sec
MCREST 0.00000000 sec
MCWRK 0.01500000 sec
======== CHANNEL f1 ========
NUC1 1H
P1 13.40 usec
PL1 -4.00 dB
SFO1 400.1318006 MHz
F2 - Processing parameters
SI 32768
SF 400.1300095 MHz
WDW EM
SSB 0
LB 0.30 Hz
GB 0
PC 1.00
f1 (ppm)

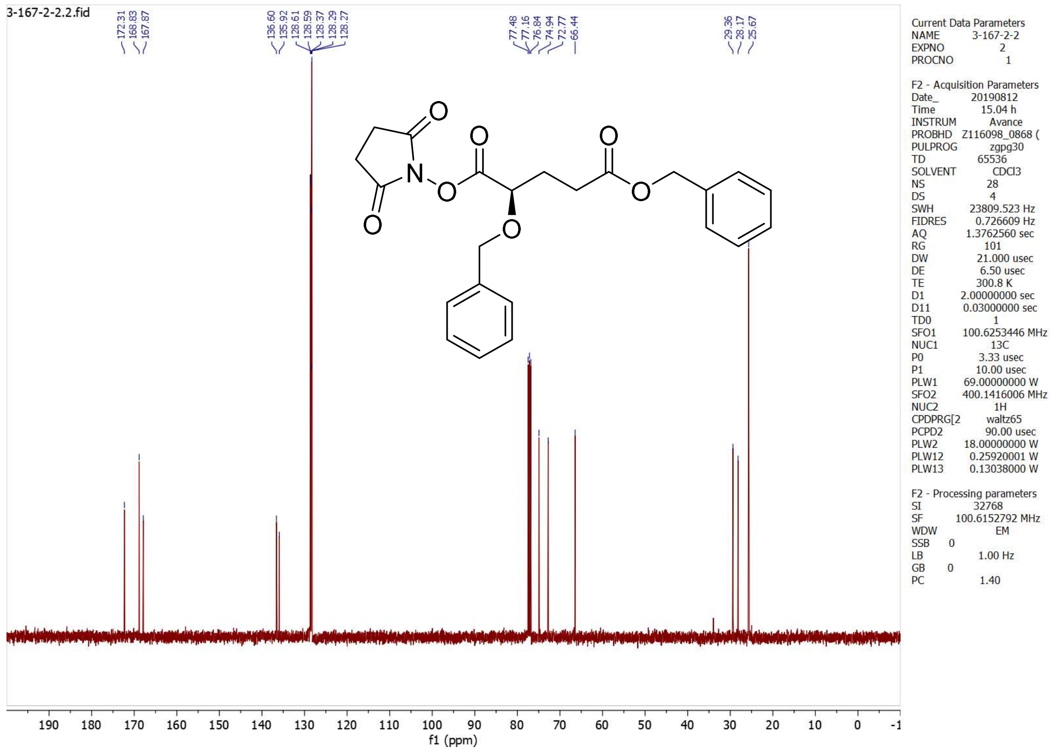
3-167-2-2.2.fid
Current Data Parameters
NAME 3-167-2-2
EXPNO 2
PROCNO 1
F2 - Acquisition Parameters
Date_ 20190812
Time 15.04 h
INSTRUM Avance
PROBHD Z116098_0868 (
PULPROG zgpg30
TD 65536
SOLVENT CDCl3
NS 28
DS 4
SWH 23809.523 Hz
FIDRES 0.726609 Hz
AQ 1.3762560 sec
RG 101
DW 21.000 usec
DE 6.50 usec
TE 300.8 K
D1 2.00000000 sec
D11 0.03000000 sec
TD0 1
SFO1 100.6253446 MHz
NUC1 13C
P0 3.33 usec
P1 10.00 usec
PLW1 69.00000000 W
SFO2 400.1416006 MHz
NUC2 1H
CPDPRG[2 waltz65
PCPD2 90.00 usec
PLW2 18.00000000 W
PLW12 0.25920001 W
PLW13 0.13038000 W
F2 - Processing parameters
SI 32768
SF 100.6152792 MHz
WDW EM
SSB 0
LB 1.00 Hz
GB 0
PC 1.40
f1 (ppm)

**Chiral-HPLC**

**2*R***

Data File C:\HPCHEM\1\DATA\AZQ\1158212R.D    Sample Name: 1158212r
CHIRALCEL OF 076-064-41006
i-PrOH:n-hexane = 30:70, 1 mL/min, 39 bar
1-158-2-1, probe 2r 8.8 mg/1.0 ml, 5 uL

=====================================================================
Injection Date : 10/10/16 4:57:57 PM
Sample Name : 1158212r    Vial : -
Acq. Operator : azq
Acq. Method : C:\HPCHEM\1\METHODS\AZQ.M
Last changed : 10/10/16 2:56:18 PM by kenneth
(modified after loading)
Analysis Method : C:\HPCHEM\1\METHODS\AZQ.M
Last changed : 10/10/16 5:58:16 PM by azq
(modified after loading)
=====================================================================

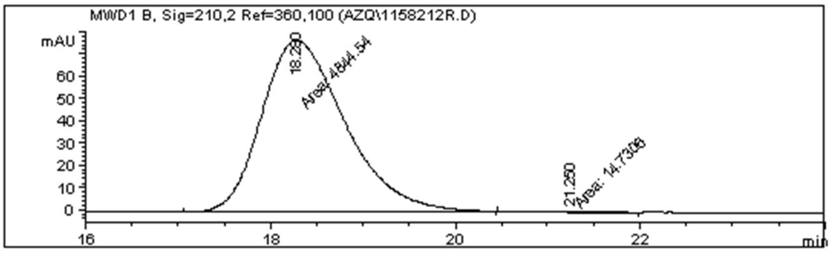

=====================================================================
Area Percent Report
=====================================================================

Sorted By : Signal
Multiplier : 1.0000
Dilution : 1.0000

Signal 1: MWD1 B, Sig=210,2 Ref=360,100

| Peak # | RetTime [min] | Type | Width [min] | Area [mAU*s] | Height [mAU] | Area % |
|---|---|---|---|---|---|---|
| 1 | 18.280 | MM | 1.0455 | 4844.53955 | 77.22689 | 99.6969 |
| 2 | 21.250 | MM | 0.6839 | 14.73057 | 3.58981e-1 | 0.3031 |
| Totals : | | | | 4859.27012 | 77.58587 | |

Results obtained with enhanced integrator!
=====================================================================
*** End of Report ***

Instrument 1 10/10/16 6:01:44 PM azq    Page 1 of 1

**2*S***

```
Data File C:\HPCHEM\1\DATA\AZQ\115921S.D                    Sample Name: 115921s
    CHIRALCEL OF 076-064-41006
    i-PrOH:n-hexane = 30:70, 1 mL/min, 39 bar
    1-159-2-1, probe 2s 11.2 mg/1.0 ml, 5 uL

    =====================================================================
    Injection Date  : 10/10/16 4:20:15 PM
    Sample Name     : 115921s                           Vial :   -
    Acq. Operator   : azq
    Acq. Method     : C:\HPCHEM\1\METHODS\AZQ.M
    Last changed    : 10/10/16 2:56:18 PM by kenneth
                      (modified after loading)
    Analysis Method : C:\HPCHEM\1\METHODS\AZQ.M
    Last changed    : 10/10/16 5:58:16 PM by azq
                      (modified after loading)
    =====================================================================
```

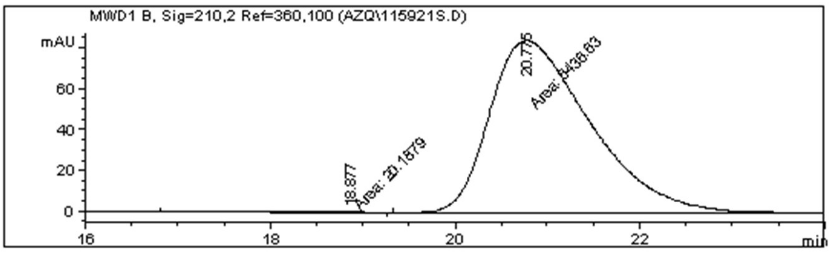

```
    =====================================================================
                             Area Percent Report
    =====================================================================

    Sorted By             :      Signal
    Multiplier            :      1.0000
    Dilution              :      1.0000

    Signal 1: MWD1 B, Sig=210,2 Ref=360,100

    Peak RetTime Type  Width     Area      Height     Area
      #   [min]        [min]   [mAU*s]     [mAU]        %
    ----|-------|----|-------|----------|----------|--------|
       1  18.877 MM    1.1030   20.18793 3.05040e-1   0.3127
       2  20.775 MM    1.2665 6436.62842   84.70600  99.6873

    Totals :                  6456.81635   85.01104

     Results obtained with enhanced integrator!
    =====================================================================
                              *** End of Report ***

Instrument 1 10/10/16 6:03:56 PM azq                              Page 1 of 1
```

Data File C:\HPCHEM\1\DATA\AZQ\115721RA.D Sample Name: 115721rac
CHIRALCEL OF 076-064-41006
i-PrOH:n-hexane = 30:70, 1 mL/min, 39 bar
1-157-2-1, probe 2rac 9.0 mg/1.0 ml, 5 uL

=====================================================================

Injection Date : 10/10/16 5:31:59 PM
Sample Name : 115721rac Vial : -
Acq. Operator : azq
Acq. Method : C:\HPCHEM\1\METHODS\AZQ.M
Last changed : 10/10/16 2:56:18 PM by kenneth
(modified after loading)
Analysis Method : C:\HPCHEM\1\METHODS\AZQ.M
Last changed : 10/10/16 5:58:16 PM by azq
(modified after loading)

=====================================================================

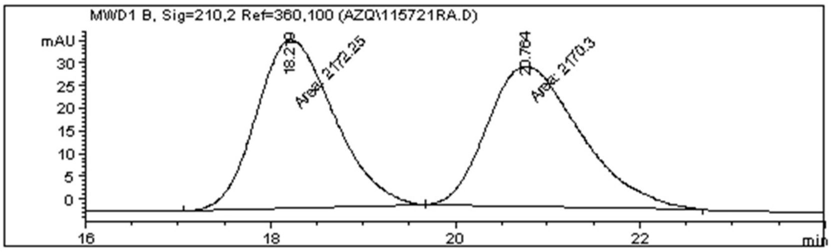

=====================================================================

Area Percent Report

=====================================================================

Sorted By : Signal
Multiplier : 1.0000
Dilution : 1.0000

Signal 1: MWD1 B, Sig=210,2 Ref=360,100

| Peak # | RetTime [min] | Type | Width [min] | Area [mAU*s] | Height [mAU] | Area % |
|---|---|---|---|---|---|---|
| 1 | 18.219 | MM | 0.9755 | 2172.25220 | 37.11288 | 50.0225 |
| 2 | 20.764 | MM | 1.1723 | 2170.29858 | 30.85599 | 49.9775 |
| Totals : | | | | 4342.55078 | 67.96887 | |

Results obtained with enhanced integrator!

=====================================================================

*** End of Report ***

**2*R*ME**

Data File C:\HPCHEM\1\DATA\AZQ\2RME16.D                Sample Name: 2rme16
CHIRALPAK ADH-H ADHOCE MB006
i-PrOH:n-hexane = 17:83, 1 mL/min, 40 bar
2rme, isopropanol, 5 uL

=====================================================================

Injection Date : 6/15/17 5:09:35 PM
Sample Name : 2rme16                Vial : -
Acq. Operator : azq
Acq. Method : C:\HPCHEM\1\METHODS\AZQ.M
Last changed : 6/15/17 3:31:34 PM by azq
(modified after loading)
Analysis Method : C:\HPCHEM\1\METHODS\AZQ.M
Last changed : 6/20/17 11:45:57 AM by azq
(modified after loading)

=====================================================================

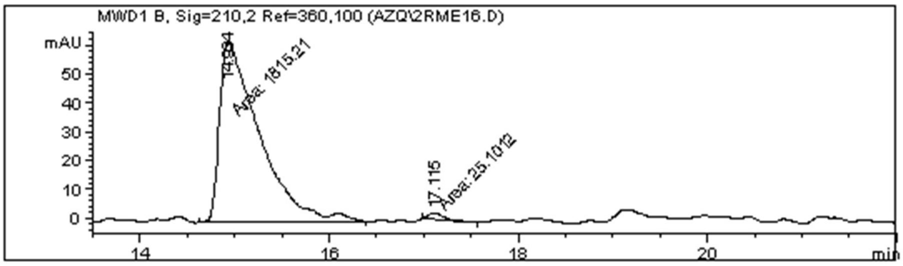

=====================================================================

Area Percent Report

=====================================================================

Sorted By : Signal
Multiplier : 1.0000
Dilution : 1.0000

Signal 1: MWD1 B, Sig=210,2 Ref=360,100

| Peak # | RetTime [min] | Type | Width [min] | Area [mAU*s] | Height [mAU] | Area % |
|---|---|---|---|---|---|---|
| 1 | 14.934 | MM | 0.4850 | 1815.20593 | 62.37236 | 98.6360 |
| 2 | 17.115 | MM | 0.1905 | 25.10120 | 2.19551 | 1.3640 |
| Totals : | | | | 1840.30713 | 64.56787 | |

Results obtained with enhanced integrator!

=====================================================================

*** End of Report ***

Instrument 1 6/20/17 11:54:38 AM azq                Page 1 of 1

**2*S*ME**

```
Data File C:\HPCHEM\1\DATA\AZQ\2SME17.D                         Sample Name: 2sme17
    CHIRALPAK ADH-H ADHOCE MB006
    i-PrOH:n-hexane = 17:83, 1 mL/min, 40 bar
    2sme, isopropanol, 5 uL

    ===================================================================
    Injection Date  : 6/15/17 6:43:56 PM
    Sample Name     : 2sme17                           Vial :   -
    Acq. Operator   : azq
    Acq. Method     : C:\HPCHEM\1\METHODS\AZQ.M
    Last changed    : 6/15/17 3:31:34 PM by azq
                      (modified after loading)
    Analysis Method : C:\HPCHEM\1\METHODS\AZQ.M
    Last changed    : 6/20/17 11:45:57 AM by azq
                      (modified after loading)
    ===================================================================
```

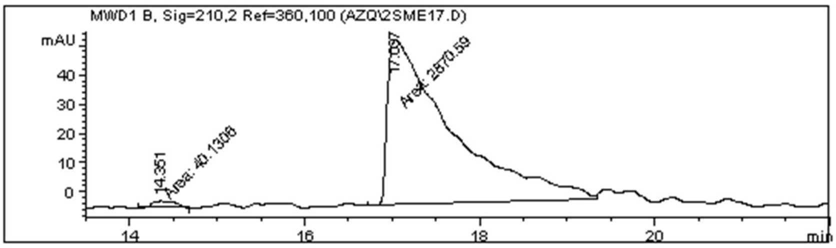

```
    ===================================================================
                             Area Percent Report
    ===================================================================

    Sorted By            :      Signal
    Multiplier           :      1.0000
    Dilution             :      1.0000

    Signal 1: MWD1 B, Sig=210,2 Ref=360,100

    Peak RetTime Type  Width     Area      Height     Area
      #   [min]        [min]   [mAU*s]     [mAU]        %
    ----|-------|----|-------|----------|----------|--------|
       1  14.351 MM    0.2919   40.13060    2.29137   1.3787
       2  17.037 MM    0.8527 2870.58667   56.10592  98.6213

    Totals :                  2910.71727   58.39729

     Results obtained with enhanced integrator!
    ===================================================================
                             *** End of Report ***
```

Data File C:\HPCHEM\1\DATA\AZQ\2RACME17.D                    Sample Name: 2racme17

CHIRALPAK ADH-H ADHOCE MB006

i-PrOH:n-hexane = 17:83, 1 mL/min, 40 bar

2racme, isopropanol, 5 uL

=====================================================================

| | |
|---|---|
| Injection Date | : 6/15/17 7:10:11 PM |
| Sample Name | : 2racme17 Vial : - |
| Acq. Operator | : azq |
| Acq. Method | : C:\HPCHEM\1\METHODS\AZQ.M |
| Last changed | : 6/15/17 3:31:34 PM by azq (modified after loading) |
| Analysis Method | : C:\HPCHEM\1\METHODS\AZQ.M |
| Last changed | : 6/20/17 11:45:57 AM by azq (modified after loading) |

=====================================================================

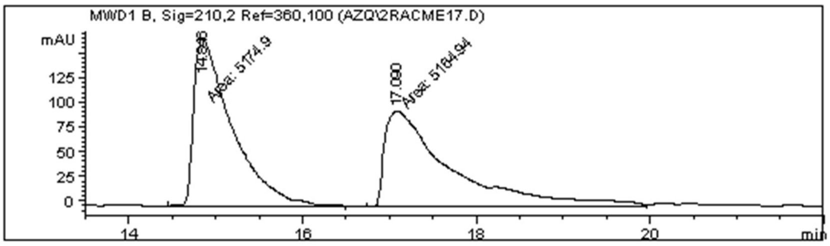

=====================================================================

Area Percent Report

=====================================================================

| | | |
|---|---|---|
| Sorted By | : | Signal |
| Multiplier | : | 1.0000 |
| Dilution | : | 1.0000 |

Signal 1: MWD1 B, Sig=210,2 Ref=360,100

| Peak # | RetTime [min] | Type | Width [min] | Area [mAU*s] | Height [mAU] | Area % |
|---|---|---|---|---|---|---|
| 1 | 14.846 | MM | 0.5108 | 5174.90039 | 168.85767 | 50.0482 |
| 2 | 17.090 | MM | 0.8885 | 5164.93652 | 96.88893 | 49.9518 |
| Totals : | | | | 1.03398e4 | 265.74660 | |

Results obtained with enhanced integrator!

=====================================================================

*** End of Report ***

Instrument 1 6/20/17 11:49:47 AM azq                    Page 1 of 1

**2*R*AM**

```
Data File C:\HPCHEM\1\DATA\AZQ\2RAM7.D                          Sample Name: 2ram7
    CHIRALPAK ADH-H ADHOCE MB006
    i-PrOH:n-hexane = 30:70, 1 mL/min, 40 bar
    2ram, isopropanol, 5 uL

    =====================================================================
    Injection Date  : 6/26/17 2:44:55 PM
    Sample Name     : 2ram7                             Vial :   -
    Acq. Operator   : azq
    Acq. Method     : C:\HPCHEM\1\METHODS\AZQ.M
    Last changed    : 6/26/17 2:16:36 PM by azq
                      (modified after loading)
    Analysis Method : C:\HPCHEM\1\METHODS\AZQ.M
    Last changed    : 6/26/17 4:25:24 PM by azq
                      (modified after loading)
    =====================================================================
```

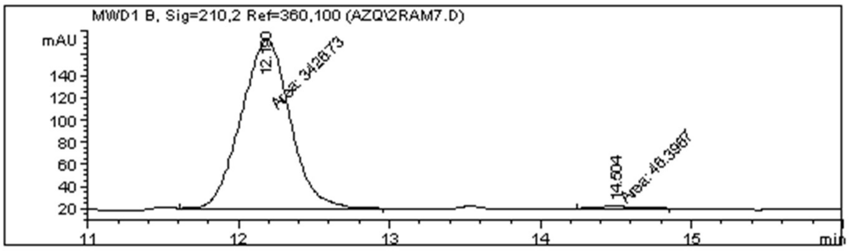

```
    =====================================================================
                             Area Percent Report
    =====================================================================

    Sorted By             :      Signal
    Multiplier            :      1.0000
    Dilution              :      1.0000

    Signal 1: MWD1 B, Sig=210,2 Ref=360,100

    Peak RetTime Type  Width     Area      Height     Area
      #   [min]        [min]   [mAU*s]     [mAU]        %
    ----|-------|----|-------|----------|----------|--------|
       1  12.190 MM    0.3750 3426.73218  152.30231  98.6641
       2  14.504 MM    0.4289   46.39666    1.80289   1.3359

    Totals :                  3473.12884  154.10519

     Results obtained with enhanced integrator!
    =====================================================================
                             *** End of Report ***
```

Instrument 1 6/26/17 4:30:12 PM azq                                Page 1 of 1

**2*S*AM**

```
Data File C:\HPCHEM\1\DATA\AZQ\2SAM7.D                                Sample Name: 2sam7
    CHIRALPAK ADH-H ADHOCE MB006
    i-PrOH:n-hexane = 30:70, 1 mL/min, 40 bar
    2sam, isopropanol, 5 uL

    =====================================================================
    Injection Date  : 6/26/17 3:39:27 PM
    Sample Name     : 2sam7                                Vial :   -
    Acq. Operator   : azq
    Acq. Method     : C:\HPCHEM\1\METHODS\AZQ.M
    Last changed    : 6/26/17 2:16:36 PM by azq
                      (modified after loading)
    Analysis Method : C:\HPCHEM\1\METHODS\AZQ.M
    Last changed    : 6/26/17 4:25:24 PM by azq
                      (modified after loading)
    =====================================================================
```

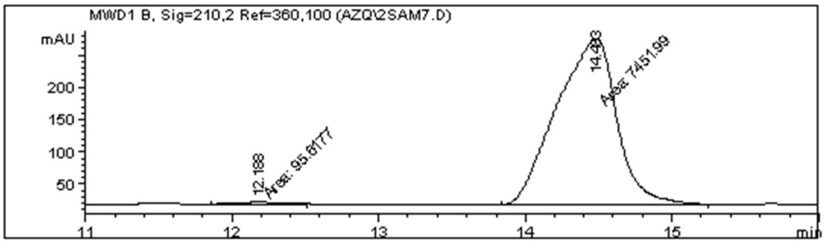

```
    =====================================================================
                             Area Percent Report
    =====================================================================

    Sorted By             :      Signal
    Multiplier            :      1.0000
    Dilution              :      1.0000

    Signal 1: MWD1 B, Sig=210,2 Ref=360,100

    Peak RetTime Type  Width     Area      Height     Area
      #   [min]        [min]   [mAU*s]     [mAU]        %
    ----|-------|----|-------|----------|----------|--------|
       1  12.188 MM    0.3620   95.61771    4.40217   1.2669
       2  14.483 MM    0.4799 7451.99268  258.80771  98.7331

    Totals :                  7547.61038  263.20988

     Results obtained with enhanced integrator!
    =====================================================================
                             *** End of Report ***
```

```
Instrument 1 6/26/17 4:31:21 PM azq                                    Page 1 of 1
```

Data File C:\HPCHEM\1\DATA\AZQ\2RACAM7.D    Sample Name: 2racam7

CHIRALPAK ADH-H ADHOCE MB006
i-PrOH:n-hexane = 30:70, 1 mL/min, 40 bar
2racam, isopropanol, 5 uL

=====================================================================

Injection Date : 6/26/17 2:17:53 PM
Sample Name : 2racam7    Vial : -
Acq. Operator : azq
Acq. Method : C:\HPCHEM\1\METHODS\AZQ.M
Last changed : 6/26/17 2:16:36 PM by azq
(modified after loading)
Analysis Method : C:\HPCHEM\1\METHODS\AZQ.M
Last changed : 6/26/17 4:25:24 PM by azq
(modified after loading)

=====================================================================

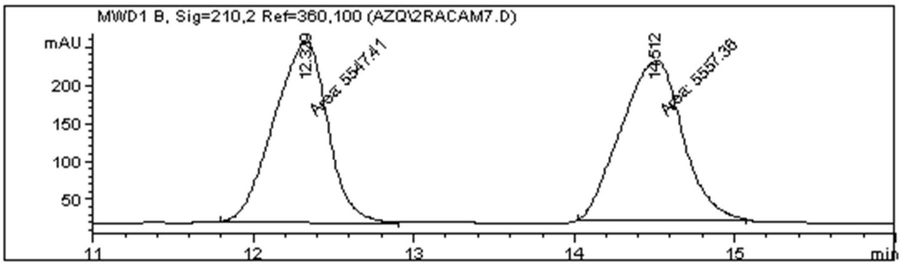

=====================================================================

Area Percent Report

=====================================================================

Sorted By : Signal
Multiplier : 1.0000
Dilution : 1.0000

Signal 1: MWD1 B, Sig=210,2 Ref=360,100

| Peak # | RetTime [min] | Type | Width [min] | Area [mAU*s] | Height [mAU] | Area % |
|---|---|---|---|---|---|---|
| 1 | 12.329 | MM | 0.3899 | 5547.40625 | 237.15739 | 49.9552 |
| 2 | 14.512 | MM | 0.4426 | 5557.35645 | 209.26860 | 50.0448 |

Totals : 1.11048e4 446.42599

Results obtained with enhanced integrator!

=====================================================================

*** End of Report ***

Instrument 1 6/26/17 4:27:39 PM azq    Page 1 of 1

**3-1 (3*R*)**

```
Data File C:\HPCHEM\1\DATA\AZQ\P312.D                         Sample Name: p3-1-2
    CHIRALPAK AD-H ADHOCE MB006
    i-PrOH:n-hexane = 10:90, 1 ml/min, 40 bar
    1-probe3-1 for 3-1,  mg/1 mL isopropanol, 5 ul

    =====================================================================
    Injection Date  : 10/28/16 12:11:22 PM
    Sample Name     : p3-1-2                              Vial :   -
    Acq. Operator   : azq
    Acq. Method     : C:\HPCHEM\1\METHODS\AZQ.M
    Last changed    : 10/22/16 12:39:56 PM by azq
    Analysis Method : C:\HPCHEM\1\METHODS\AZQ.M
    Last changed    : 10/31/16 6:54:17 PM by azq
                      (modified after loading)
    =====================================================================
```

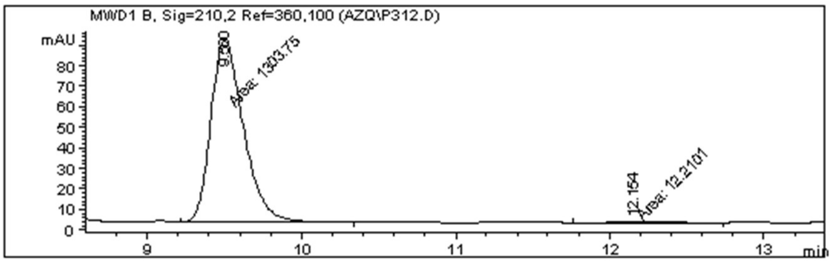

```
    =====================================================================
                             Area Percent Report
    =====================================================================

    Sorted By             :      Signal
    Multiplier            :      1.0000
    Dilution              :      1.0000

    Signal 1: MWD1 B, Sig=210,2 Ref=360,100

    Peak RetTime Type  Width     Area      Height     Area
      #   [min]        [min]   [mAU*s]     [mAU]        %
    ----|-------|----|-------|----------|----------|--------|
       1   9.500 MM    0.2453 1303.75488   88.58266 99.0722
       2  12.154 MM    0.3640   12.21007 5.59051e-1  0.9278

    Totals :                  1315.96495   89.14171

     Results obtained with enhanced integrator!
    =====================================================================
                             *** End of Report ***
```

Instrument 1 10/31/16 7:00:04 PM azq                                    Page 1 of 1

**3-2 (3*S*)**

Data File C:\HPCHEM\1\DATA\AZQ\P3-2.D Sample Name: p3-2
CHIRALPAK AD-H ADHOCE MB006
i-PrOH:n-hexane = 10:90, 1 ml/min, 40 bar
1-probe3-2rac, mg/1 mL isopropanol, 5 ul

=====================================================================

Injection Date : 10/25/16 5:13:32 PM
Sample Name : p3-2 Vial : -
Acq. Operator : azq
Acq. Method : C:\HPCHEM\1\METHODS\AZQ.M
Last changed : 10/22/16 12:39:56 PM by azq
Analysis Method : C:\HPCHEM\1\METHODS\AZQ.M
Last changed : 10/31/16 6:54:17 PM by azq
(modified after loading)

=====================================================================

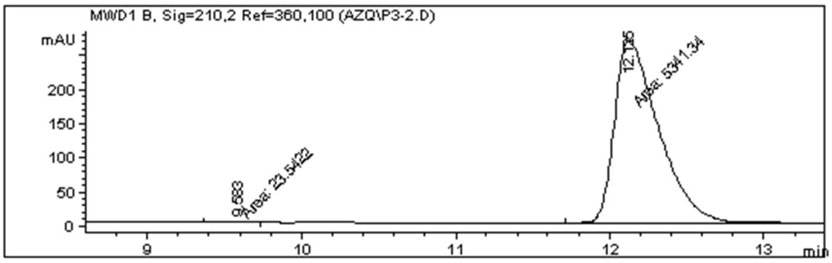

=====================================================================

Area Percent Report

=====================================================================

Sorted By : Signal
Multiplier : 1.0000
Dilution : 1.0000

Signal 1: MWD1 B, Sig=210,2 Ref=360,100

| Peak # | RetTime [min] | Type | Width [min] | Area [mAU*s] | Height [mAU] | Area % |
|---|---|---|---|---|---|---|
| 1 | 9.583 | MM | 0.1878 | 23.54221 | 2.08924 | 0.4388 |
| 2 | 12.125 | MM | 0.3321 | 5341.34033 | 268.05963 | 99.5612 |
| Totals : | | | | 5364.88254 | 270.14887 | |

Results obtained with enhanced integrator!

=====================================================================

*** End of Report ***

Instrument 1 10/31/16 6:58:37 PM azq Page 1 of 1

Data File C:\HPCHEM\1\DATA\AZQ\P3RAC2.D    Sample Name: p3rac2

CHIRALPAK AD-H ADHOCE MB006
i-PrOH:n-hexane = 10:90, 1 ml/min, 40 bar
1-probe3rac for 3-1,  mg/1 mL isopropanol, 5 ul

=====================================================================

Injection Date  : 10/28/16 11:29:37 AM
Sample Name     : p3rac2                              Vial :   -
Acq. Operator   : azq
Acq. Method     : C:\HPCHEM\1\METHODS\AZQ.M
Last changed    : 10/22/16 12:39:56 PM by azq
Analysis Method : C:\HPCHEM\1\METHODS\AZQ.M
Last changed    : 10/31/16 6:54:17 PM by azq
(modified after loading)

=====================================================================

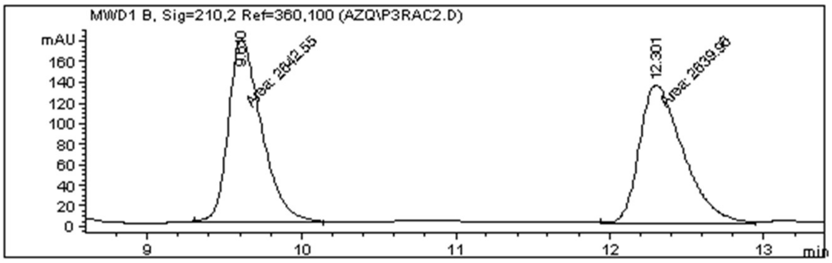

=====================================================================

Area Percent Report

=====================================================================

Sorted By : Signal
Multiplier : 1.0000
Dilution : 1.0000

Signal 1: MWD1 B, Sig=210,2 Ref=360,100

| Peak # | RetTime [min] | Type | Width [min] | Area [mAU*s] | Height [mAU] | Area % |
|---|---|---|---|---|---|---|
| 1 | 9.610 | MM | 0.2493 | 2642.55420 | 176.67404 | 50.0245 |
| 2 | 12.301 | MF | 0.3282 | 2639.96265 | 134.04523 | 49.9755 |

Totals : 5282.51685 310.71927

Results obtained with enhanced integrator!

=====================================================================

*** End of Report ***

Instrument 1 10/31/16 7:03:46 PM azq    Page 1 of 1

**4-1 (4*SS*)**

```
Data File C:\Users\P...eckout 2021-07-17 16-21-37\4ss YKQ 2021-07-17 17-03-50id-95-5-1.0.D
Sample Name: 4ss

    =====================================================================
    Acq. Operator   : SYSTEM                       Seq. Line :   3
    Sample Operator : SYSTEM
    Acq. Instrument : HPLC                          Location :   P2-A-03
    Injection Date  : 7/17/2021 5:04:46 PM               Inj :   1
                                                  Inj Volume : 20.000 µl
    Acq. Method     : C:\Users\Public\Documents\ChemStation\1\Data\angela\checkout 2021-07-17 16-
                      21-37\YKQ.M
    Last changed    : 7/17/2021 4:21:34 PM by SYSTEM
    Analysis Method : C:\Users\Public\Documents\ChemStation\1\Data\angela\checkout 2021-07-17 16-
                      21-37\YKQ.M (Sequence Method)
    Last changed    : 7/17/2021 5:22:14 PM by SYSTEM
                      (modified after loading)
    Additional Info : Peak(s) manually integrated
```

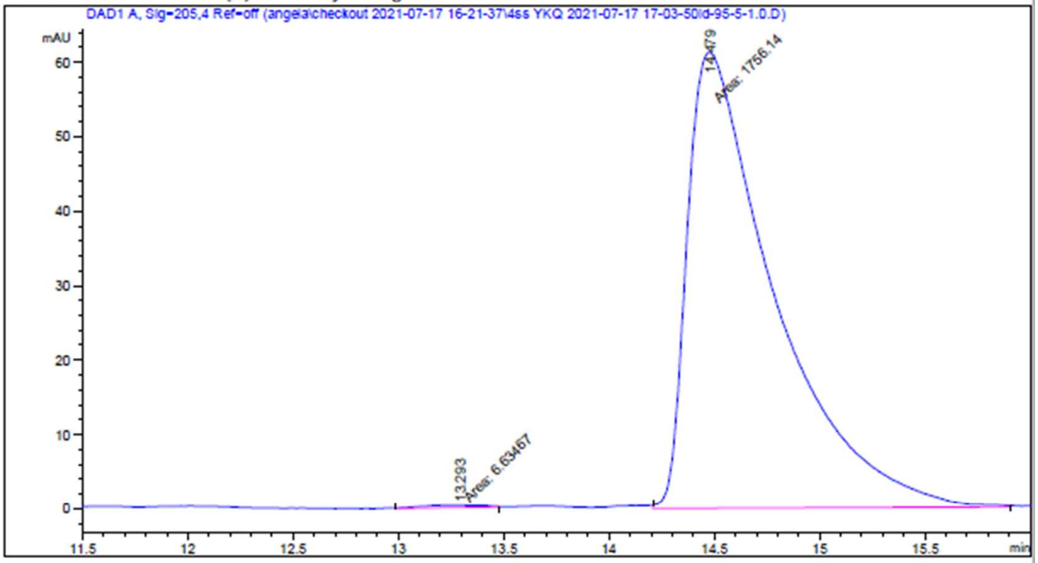

```
    =====================================================================
                             Area Percent Report
    =====================================================================

    Sorted By             :      Signal
    Multiplier            :      1.0000
    Dilution              :      1.0000
    Use Multiplier & Dilution Factor with ISTDs

    Signal 1: DAD1 A, Sig=205,4 Ref=off

    Peak RetTime Type  Width     Area      Height     Area
      #   [min]        [min]   [mAU*s]     [mAU]       %
    ----|-------|----|-------|----------|----------|--------|
       1  13.293 MM    0.3288    6.63467 3.36332e-1   0.3764
       2  14.479 MM    0.4776 1756.13916   61.28517  99.6236

    Totals :                  1762.77383   61.62150

HPLC 7/17/2021 5:25:29 PM SYSTEM                                              Page    1 of 2
```

**4-2 (4*RR*)**

Data File C:\Users\P...eckout 2021-07-17 16-21-37\4rr YKQ 2021-07-17 16-42-45id-95-5-1.0.D
Sample Name: 4rr

=====================================================================

Acq. Operator : SYSTEM Seq. Line : 2
Sample Operator : SYSTEM
Acq. Instrument : HPLC Location : P2-A-02
Injection Date : 7/17/2021 4:43:42 PM Inj : 1
Inj Volume : 20.000 µl
Acq. Method : C:\Users\Public\Documents\ChemStation\1\Data\angela\checkout 2021-07-17 16-21-37\YKQ.M
Last changed : 7/17/2021 4:21:34 PM by SYSTEM
Analysis Method : C:\Users\Public\Documents\ChemStation\1\Data\angela\checkout 2021-07-17 16-21-37\YKQ.M (Sequence Method)
Last changed : 7/17/2021 5:22:14 PM by SYSTEM
(modified after loading)
Additional Info : Peak(s) manually integrated

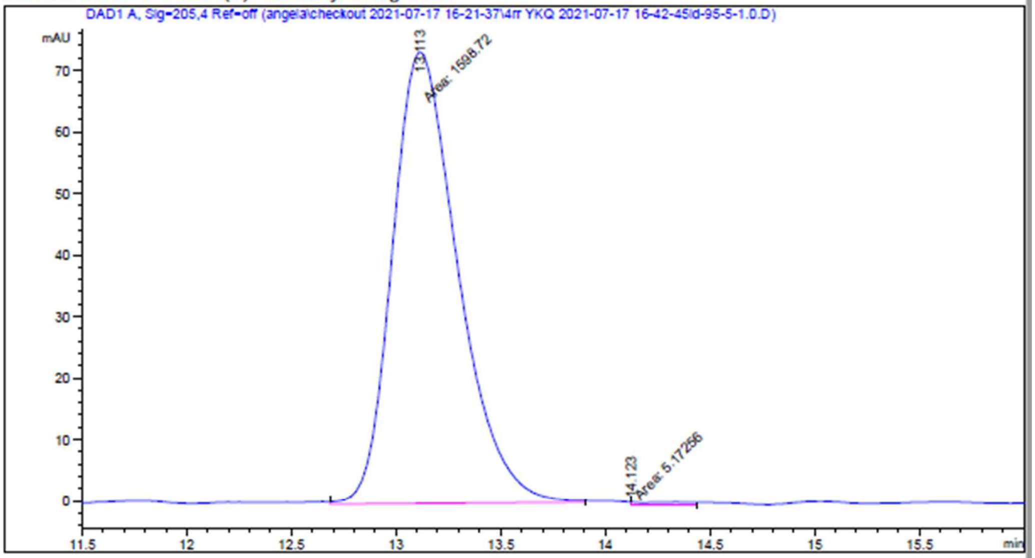

=====================================================================
Area Percent Report
=====================================================================

Sorted By : Signal
Multiplier : 1.0000
Dilution : 1.0000
Use Multiplier & Dilution Factor with ISTDs

Signal 1: DAD1 A, Sig=205,4 Ref=off

| Peak # | RetTime [min] | Type | Width [min] | Area [mAU*s] | Height [mAU] | Area % |
|---|---|---|---|---|---|---|
| 1 | 13.113 | MM | 0.3630 | 1598.71875 | 73.41035 | 99.6775 |
| 2 | 14.123 | MM | 0.2597 | 5.17256 | 3.31980e-1 | 0.3225 |

Totals : 1603.89131 73.74233

HPLC 7/17/2021 5:24:13 PM SYSTEM Page 1 of 2

Data File C:\Users\P...kout 2021-07-17 16-21-37\4rrss YKQ 2021-07-17 16-21-38id-95-5-1.0.D
Sample Name: 4rrss

=====================================================================

Acq. Operator : SYSTEM Seq. Line : 1
Sample Operator : SYSTEM
Acq. Instrument : HPLC Location : P2-A-01
Injection Date : 7/17/2021 4:22:37 PM Inj : 1
Inj Volume : 20.000 µl
Acq. Method : C:\Users\Public\Documents\ChemStation\1\Data\angela\checkout 2021-07-17 16-21-37\YKQ.M
Last changed : 7/17/2021 4:21:34 PM by SYSTEM
Analysis Method : C:\Users\Public\Documents\ChemStation\1\Data\angela\checkout 2021-07-17 16-21-37\YKQ.M (Sequence Method)
Last changed : 7/17/2021 5:22:14 PM by SYSTEM
(modified after loading)
Additional Info : Peak(s) manually integrated

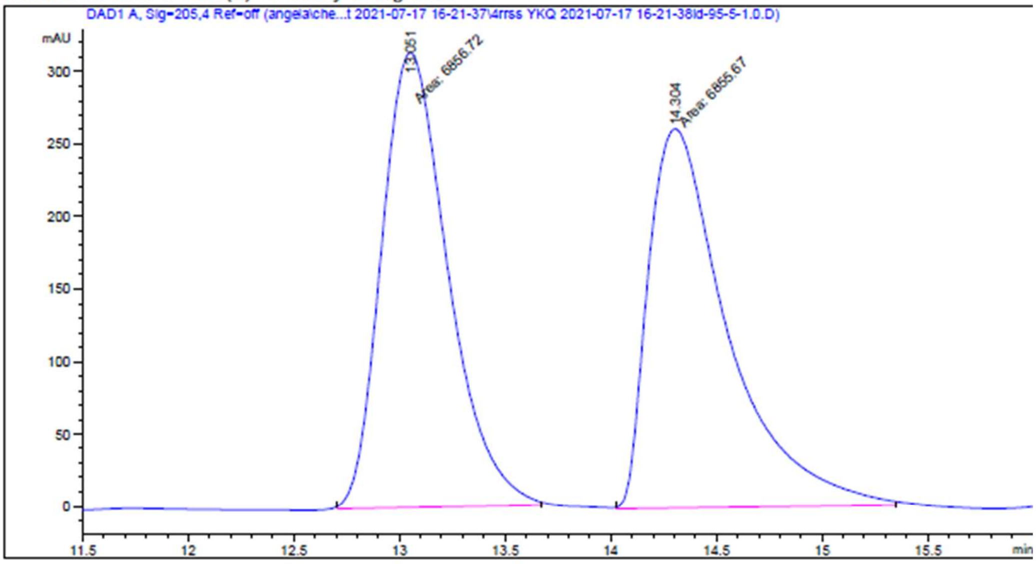

=====================================================================

Area Percent Report

=====================================================================

Sorted By : Signal
Multiplier : 1.0000
Dilution : 1.0000
Use Multiplier & Dilution Factor with ISTDs

Signal 1: DAD1 A, Sig=205,4 Ref=off

| Peak # | RetTime [min] | Type | Width [min] | Area [mAU*s] | Height [mAU] | Area % |
|---|---|---|---|---|---|---|
| 1 | 13.051 | MM | 0.3641 | 6856.71777 | 313.85315 | 50.0038 |
| 2 | 14.304 | MM | 0.4367 | 6855.66943 | 261.62888 | 49.9962 |

Totals : 1.37124e4 575.48203

**5S-1 (5*RS*)**

```
Data File C:\Users\P...eckout 2021-07-16 17-34-06\5rs YKQ 2021-07-16 17-55-15id-95-5-1.0.D
Sample Name: 5rs

    =====================================================================
    Acq. Operator   : SYSTEM                          Seq. Line :   2
    Sample Operator : SYSTEM
    Acq. Instrument : HPLC                             Location :   P2-A-05
    Injection Date  : 7/16/2021 5:56:12 PM                  Inj :   1
                                                     Inj Volume : 20.000 µl
    Acq. Method     : C:\Users\Public\Documents\ChemStation\1\Data\angela\checkout 2021-07-16 17-
                      34-06\YKQ.M
    Last changed    : 7/16/2021 5:33:59 PM by SYSTEM
    Analysis Method : C:\Users\Public\Documents\ChemStation\1\Data\angela\checkout 2021-07-16 17-
                      34-06\YKQ.M (Sequence Method)
    Last changed    : 7/17/2021 5:18:33 PM by SYSTEM
                      (modified after loading)
    Additional Info : Peak(s) manually integrated
```

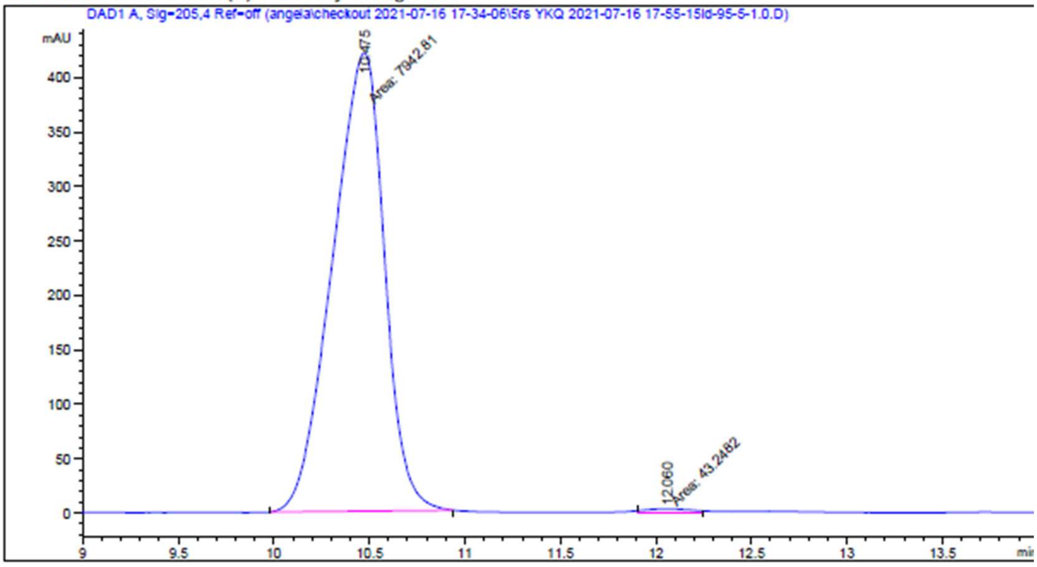

```
    =====================================================================
                         Area Percent Report
    =====================================================================

    Sorted By             :      Signal
    Multiplier            :      1.0000
    Dilution              :      1.0000
    Use Multiplier & Dilution Factor with ISTDs

    Signal 1: DAD1 A, Sig=205,4 Ref=off

    Peak RetTime Type  Width     Area      Height     Area
      #   [min]        [min]   [mAU*s]     [mAU]        %
    ----|-------|----|-------|----------|----------|--------|
       1  10.475 MM    0.3147 7942.80908  420.70035  99.4585
       2  12.060 MM    0.2518   43.24819    2.86264   0.5415

    Totals :                  7986.05727  423.56299

HPLC 7/17/2021 5:20:45 PM SYSTEM                                         Page    1 of 2
```

**5R-1 (5*SR*)**

```
Data File C:\Users\P...eckout 2021-07-16 17-34-06\5sr YKQ 2021-07-16 18-16-19id-95-5-1.0.D
Sample Name: 5sr

    =====================================================================
    Acq. Operator   : SYSTEM                         Seq. Line :   3
    Sample Operator : SYSTEM
    Acq. Instrument : HPLC                            Location :   P2-A-06
    Injection Date  : 7/16/2021 6:17:16 PM                 Inj :   1
                                                    Inj Volume : 20.000 µl
    Acq. Method     : C:\Users\Public\Documents\ChemStation\1\Data\angela\checkout 2021-07-16 17-
                      34-06\YKQ.M
    Last changed    : 7/16/2021 5:33:59 PM by SYSTEM
    Analysis Method : C:\Users\Public\Documents\ChemStation\1\Data\angela\checkout 2021-07-16 17-
                      34-06\YKQ.M (Sequence Method)
    Last changed    : 7/17/2021 5:18:33 PM by SYSTEM
                      (modified after loading)
    Additional Info : Peak(s) manually integrated
```

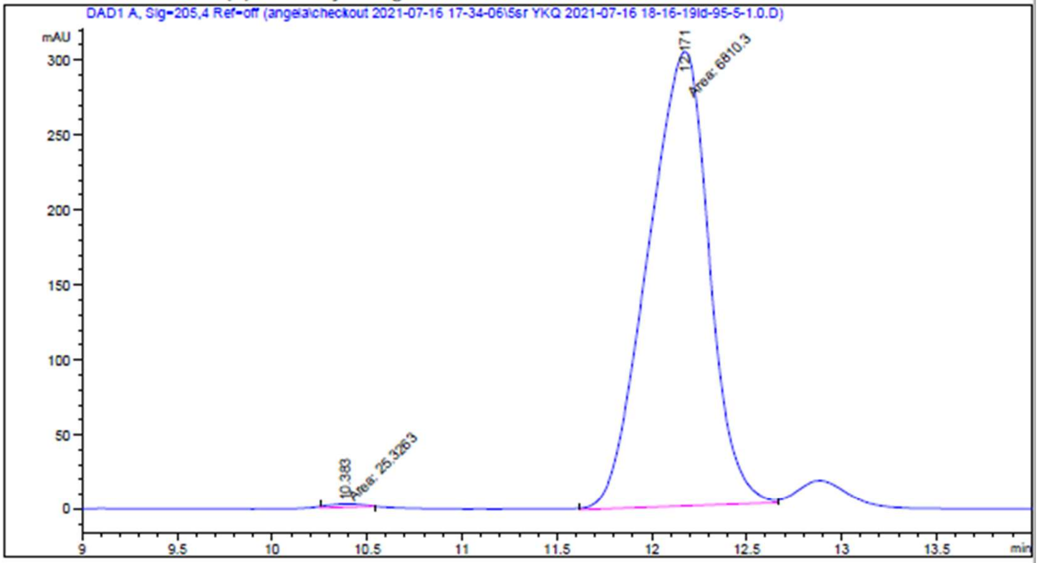

```
    =====================================================================
                         Area Percent Report
    =====================================================================

    Sorted By             :      Signal
    Multiplier            :      1.0000
    Dilution              :      1.0000
    Use Multiplier & Dilution Factor with ISTDs

    Signal 1: DAD1 A, Sig=205,4 Ref=off
```

| Peak # | RetTime [min] | Type | Width [min] | Area [mAU*s] | Height [mAU] | Area % |
|---|---|---|---|---|---|---|
| 1 | 10.383 | MM | 0.1929 | 25.32630 | 2.18859 | 0.3705 |
| 2 | 12.171 | MM | 0.3737 | 6810.29834 | 303.70422 | 99.6295 |

Totals : 6835.62464 305.89281

HPLC 7/17/2021 5:21:31 PM SYSTEM Page 1 of 2

Data File C:\Users\P...kout 2021-07-16 17-34-06\5rssr YKQ 2021-07-16 17-34-07id-95-5-1.0.D
Sample Name: 5rssr

=====================================================================

Acq. Operator : SYSTEM Seq. Line : 1
Sample Operator : SYSTEM
Acq. Instrument : HPLC Location : P2-A-04
Injection Date : 7/16/2021 5:35:07 PM Inj : 1
Inj Volume : 20.000 µl
Acq. Method : C:\Users\Public\Documents\ChemStation\1\Data\angela\checkout 2021-07-16 17-34-06\YKQ.M
Last changed : 7/16/2021 5:33:59 PM by SYSTEM
Analysis Method : C:\Users\Public\Documents\ChemStation\1\Data\angela\checkout 2021-07-16 17-34-06\YKQ.M (Sequence Method)
Last changed : 7/17/2021 5:18:33 PM by SYSTEM
(modified after loading)
Additional Info : Peak(s) manually integrated

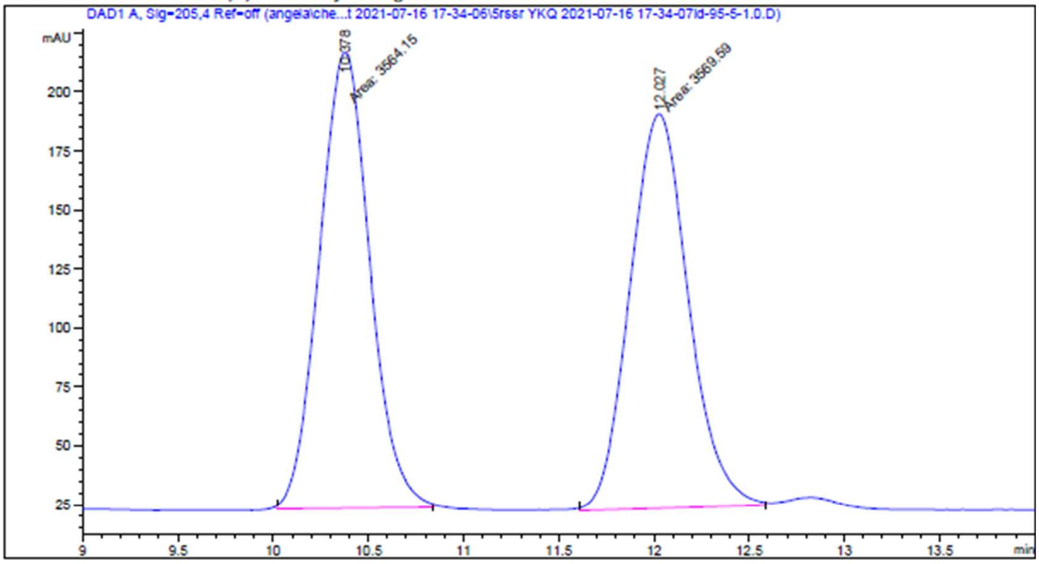

=====================================================================

Area Percent Report

=====================================================================

Sorted By : Signal
Multiplier : 1.0000
Dilution : 1.0000
Use Multiplier & Dilution Factor with ISTDs

Signal 1: DAD1 A, Sig=205,4 Ref=off

| Peak # | RetTime [min] | Type | Width [min] | Area [mAU*s] | Height [mAU] | Area % |
|---|---|---|---|---|---|---|
| 1 | 10.378 | MM | 0.3075 | 3564.14551 | 193.19498 | 49.9618 |
| 2 | 12.027 | MM | 0.3556 | 3569.59131 | 167.29362 | 50.0382 |

Totals : 7133.73682 360.48860

HPLC 7/17/2021 5:19:47 PM SYSTEM Page 1 of 2

**5S-2 (5*RR*)**

```
Data File C:\Users\P...eckout 2021-07-16 15-57-01\5rr YKQ 2021-07-16 16-13-10id-95-5-1.0.D
Sample Name: 5rr

    =====================================================================
    Acq. Operator   : SYSTEM                          Seq. Line :    2
    Sample Operator : SYSTEM
    Acq. Instrument : HPLC                             Location :   P2-A-02
    Injection Date  : 7/16/2021 4:14:07 PM                  Inj :    1
                                                    Inj Volume : 20.000 µl
    Acq. Method     : C:\Users\Public\Documents\ChemStation\1\Data\angela\checkout 2021-07-16 15-
                      57-01\YKQ.M
    Last changed    : 7/16/2021 4:10:03 PM by SYSTEM
    Analysis Method : C:\Users\Public\Documents\ChemStation\1\Data\angela\checkout 2021-07-16 15-
                      57-01\YKQ.M (Sequence Method)
    Last changed    : 7/17/2021 5:14:15 PM by SYSTEM
                      (modified after loading)
    Additional Info : Peak(s) manually integrated
```

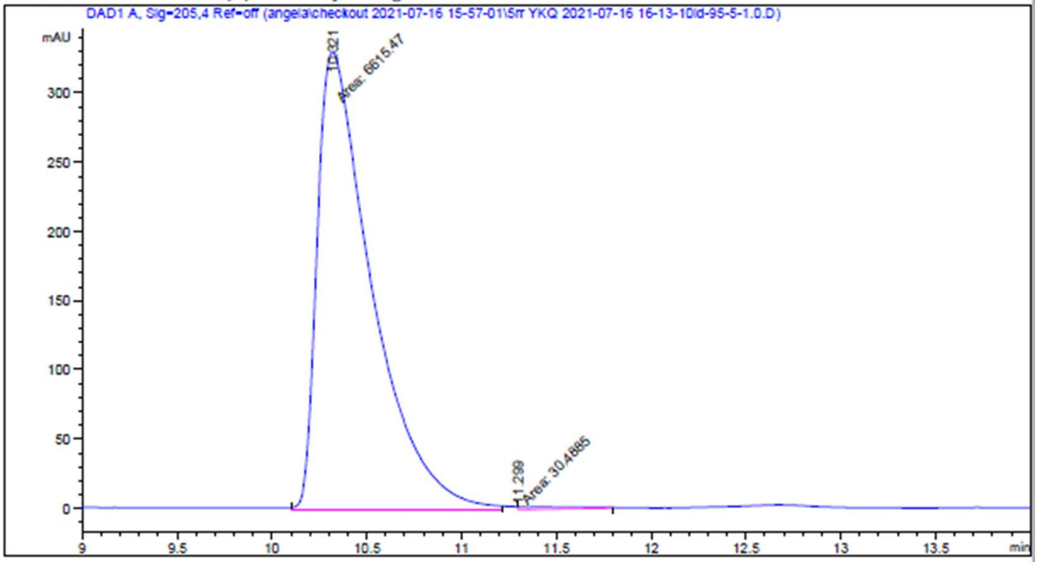

```
    =====================================================================
                            Area Percent Report
    =====================================================================

    Sorted By             :      Signal
    Multiplier            :      1.0000
    Dilution              :      1.0000
    Use Multiplier & Dilution Factor with ISTDs

    Signal 1: DAD1 A, Sig=205,4 Ref=off

    Peak RetTime Type  Width     Area      Height     Area
      #   [min]        [min]   [mAU*s]     [mAU]       %
    ----|-------|----|-------|----------|----------|--------|
       1  10.321 MM    0.3334 6615.46680  330.73581  99.5412
       2  11.299 MM    0.2490   30.48853    2.04062   0.4588

    Totals :                  6645.95533  332.77642
```

HPLC 7/17/2021 5:14:21 PM SYSTEM                                     Page    1 of 2

**5R-2 (5*SS*)**

```
Data File C:\Users\P...eckout 2021-07-16 15-57-01\5ss YKQ 2021-07-16 16-29-14id-95-5-1.0.D
Sample Name: 5ss

    ===================================================================
    Acq. Operator   : SYSTEM                        Seq. Line :   3
    Sample Operator : SYSTEM
    Acq. Instrument : HPLC                           Location :   P2-A-03
    Injection Date  : 7/16/2021 4:30:11 PM                Inj :   1
                                                Inj Volume : 20.000 µl
    Acq. Method     : C:\Users\Public\Documents\ChemStation\1\Data\angela\checkout 2021-07-16 15-
                      57-01\YKQ.M
    Last changed    : 7/16/2021 4:10:03 PM by SYSTEM
    Analysis Method : C:\Users\Public\Documents\ChemStation\1\Data\angela\checkout 2021-07-16 15-
                      57-01\YKQ.M (Sequence Method)
    Last changed    : 7/17/2021 5:15:49 PM by SYSTEM
                      (modified after loading)
    Additional Info : Peak(s) manually integrated
```

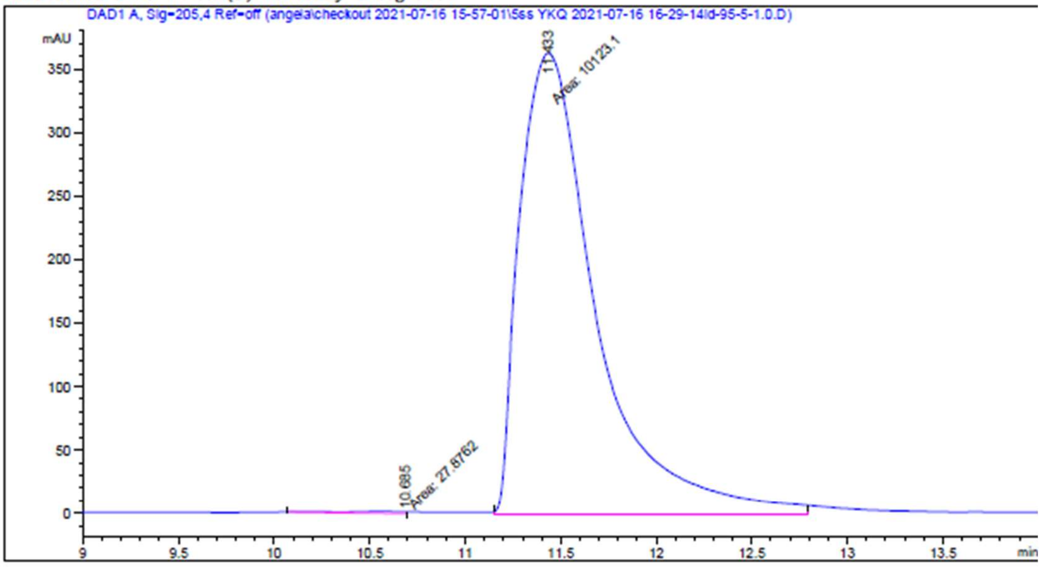

```
    ===================================================================
                         Area Percent Report
    ===================================================================

    Sorted By             :      Signal
    Multiplier            :      1.0000
    Dilution              :      1.0000
    Use Multiplier & Dilution Factor with ISTDs

    Signal 1: DAD1 A, Sig=205,4 Ref=off

    Peak RetTime Type  Width     Area      Height     Area
      #   [min]        [min]   [mAU*s]     [mAU]        %
    ----|-------|----|-------|----------|----------|--------|
       1  10.685 MM    0.3027   27.87616    1.53502   0.2746
       2  11.433 MM    0.4646 1.01231e4  363.13770  99.7254

    Totals :                  1.01509e4  364.67271
```

Data File C:\Users\P...kout 2021-07-16 15-57-01\5rrss YKQ 2021-07-16 15-57-02id-95-5-1.0.D
Sample Name: 5rrss

=====================================================================

Acq. Operator : SYSTEM    Seq. Line : 1
Sample Operator : SYSTEM
Acq. Instrument : HPLC    Location : P2-A-01
Injection Date : 7/16/2021 3:58:02 PM    Inj : 1
Inj Volume : 20.000 µl
Acq. Method : C:\Users\Public\Documents\ChemStation\1\Data\angela\checkout 2021-07-16 15-57-01\YKQ.M
Last changed : 7/16/2021 4:10:03 PM by SYSTEM
(modified after loading)
Analysis Method : C:\Users\Public\Documents\ChemStation\1\Data\angela\checkout 2021-07-16 15-57-01\YKQ.M (Sequence Method)
Last changed : 7/17/2021 5:15:49 PM by SYSTEM
(modified after loading)
Additional Info : Peak(s) manually integrated

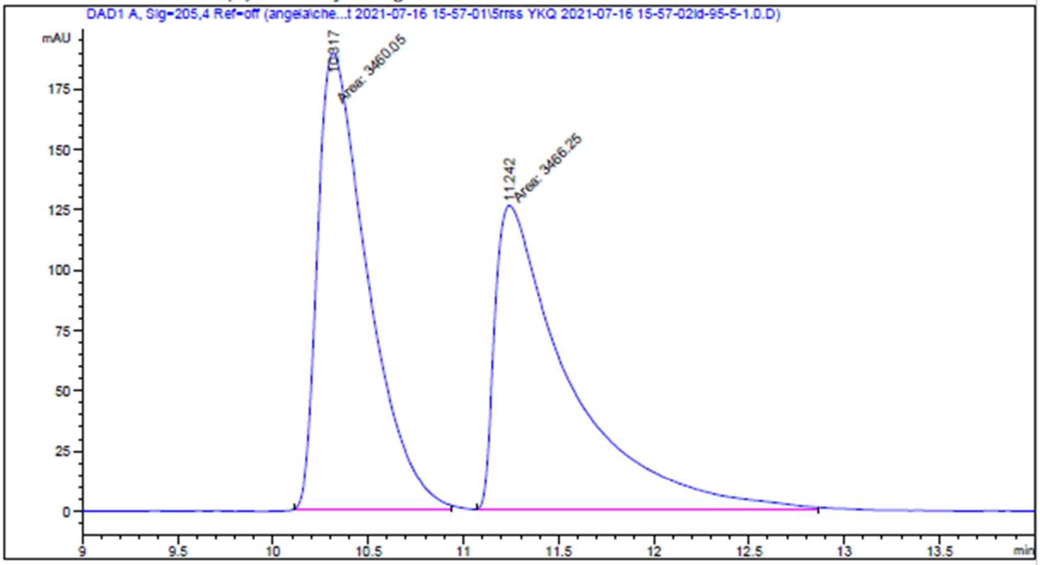

=====================================================================

Area Percent Report

=====================================================================

Sorted By : Signal
Multiplier : 1.0000
Dilution : 1.0000
Use Multiplier & Dilution Factor with ISTDs

Signal 1: DAD1 A, Sig=205,4 Ref=off

| Peak # | RetTime [min] | Type | Width [min] | Area [mAU*s] | Height [mAU] | Area % |
|---|---|---|---|---|---|---|
| 1 | 10.317 | MM | 0.3042 | 3460.04956 | 189.57672 | 49.9552 |
| 2 | 11.242 | MM | 0.4574 | 3466.25146 | 126.30481 | 50.0448 |

#### X-Ray crystallography

### 3.19-2

**Table 1**
Experimental details

| | |
|---|---|
| Crystal data | |
| Chemical formula | $C_{17}H_{19}NO_4$ |
| $M_r$ | 301.33 |
| Crystal system, space group | Orthorhombic, $P2_12_12_1$ |
| Temperature (K) | 173 |
| *a*, *b*, *c* (Å) | 6.0081 (2), 9.5689 (4), 26.818 (1) |
| *V* (Å$^3$) | 1541.79 (10) |
| *Z* | 4 |
| Radiation type | Cu *Kα* |
| *μ* (mm$^{-1}$) | 0.76 |
| Crystal size (mm) | 0.46 × 0.33 × 0.27 |
| Data collection | |
| Diffractometer | Bruker D8 VENTURE FIXED-CHI PHOTON 100 CMOS |
| Absorption correction | Multi-scan<br>*SADABS* 2014/5 (Sheldrick, 2014) |
| $T_{min}$, $T_{max}$ | 0.580, 0.753 |
| No. of measured, independent and observed [$I > 2\sigma(I)$] reflections | 6237, 2656, 2564 |
| $R_{int}$ | 0.052 |
| $(\sin \theta/\lambda)_{max}$ (Å$^{-1}$) | 0.595 |
| Refinement | |
| $R[F^2 > 2\sigma(F^2)]$, $wR(F^2)$, *S* | 0.053, 0.157, 1.15 |
| No. of reflections | 2656 |
| No. of parameters | 206 |
| H-atom treatment | H atoms treated by a mixture of independent and constrained refinement |
| $\Delta\rho_{max}$, $\Delta\rho_{min}$ (e Å$^{-3}$) | 0.23, −0.28 |
| Absolute structure | Flack x determined using 982 quotients [(I+)-(I-)]/[(I+)+(I-)] (Parsons, Flack and Wagner, Acta Cryst. B69 (2013) 249-259). |
| Absolute structure parameter | 0.05 (12) |

Computer programs: *APEX3* v2015.5-2 (Bruker-AXS, 2015), *SAINT* v8.34A (Bruker-AXS, 2013), *SHELXT2014/5* (Sheldrick, 2015), *SHELXL2014/7* (Sheldrick, 2014), Bruker *SHELXTL*.

**Acknowledgements**

The authors thank The University of Hong Kong's University Development Fund and the Dr. Hui Wai Haan Fund for funding the Bruker D8 VENTURE Photon100 CMOS X-Ray Diffractometer.

##### Funding information

##### Figure 1

Fig. 1. *ORTEP* diagram of <your molecule> with thermal ellipsoids at 50% probability level.

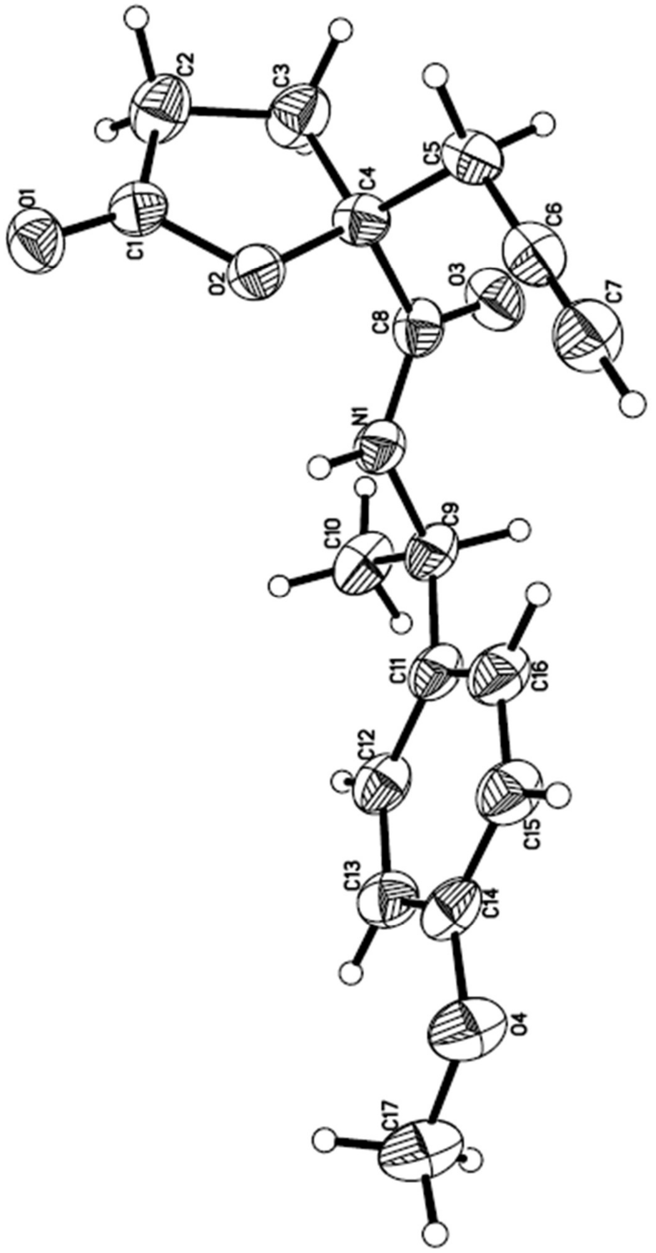

#### Title

##### Computing details

Data collection: *APEX3* v2015.5-2 (Bruker-AXS, 2015); cell refinement: *SAINT* v8.34A (Bruker-AXS, 2013); data reduction: *SAINT* v8.34A (Bruker-AXS, 2013); program(s) used to solve structure: *SHELXT2014/5* (Sheldrick, 2015); program(s) used to refine structure: *SHELXL2014/7* (Sheldrick, 2014); molecular graphics: Bruker *SHELXTL*; software used to prepare material for publication: Bruker *SHELXTL*.

##### (cu_CHEM0371)

*Crystal data*

$C_{17}H_{19}NO_4$
$M_r$ = 301.33
Orthorhombic, $P2_12_12_1$
$a$ = 6.0081 (2) Å
$b$ = 9.5689 (4) Å
$c$ = 26.818 (1) Å
$V$ = 1541.79 (10) Å$^3$
$Z$ = 4
$F$(000) = 640
$D_x$ = 1.298 Mg m$^{-3}$
Cu $K\alpha$ radiation, $\lambda$ = 1.54178 Å
Cell parameters from 174 reflections
$\theta$ = 3–45°
$\mu$ = 0.76 mm$^{-1}$
$T$ = 173 K
Block, colorless
0.46 × 0.33 × 0.27 mm

*Data collection*

Bruker D8 VENTURE FIXED-CHI PHOTON 100 CMOS diffractometer
Radiation source: I$\mu$S HB micro-focus sealed tube
Detector resolution: 10.24 pixels mm$^{-1}$
$\varphi$ and $\omega$ scans
Absorption correction: multi-scan *SADABS* 2014/5 (Sheldrick, 2014)
$T_{min}$ = 0.580, $T_{max}$ = 0.753
6237 measured reflections
2656 independent reflections
2564 reflections with $I > 2\sigma(I)$
$R_{int}$ = 0.052
$\theta_{max}$ = 66.7°, $\theta_{min}$ = 3.3°
$h$ = −6→7
$k$ = −10→11
$l$ = −27→31

*Refinement*

Refinement on $F^2$
Least-squares matrix: full
$R[F^2 > 2\sigma(F^2)]$ = 0.053
$wR(F^2)$ = 0.157
$S$ = 1.15
2656 reflections
206 parameters
0 restraints
Hydrogen site location: mixed
H atoms treated by a mixture of independent and constrained refinement
$w = 1/[\sigma^2(F_o^2) + (0.0744P)^2 + 0.7712P]$ where $P = (F_o^2 + 2F_c^2)/3$
$(\Delta/\sigma)_{max}$ < 0.001
$\Delta\rho_{max}$ = 0.23 e Å$^{-3}$
$\Delta\rho_{min}$ = −0.28 e Å$^{-3}$
Extinction correction: *SHELXL2014/7* (Sheldrick 2014, $Fc^* = kFc[1+0.001xFc^2\lambda^3/\sin(2\theta)]^{-1/4}$
Extinction coefficient: 0.023 (3)
Absolute structure: Flack x determined using 982 quotients [(I+)-(I-)]/[(I+)+(I-)] (Parsons, Flack and Wagner, Acta Cryst. B69 (2013) 249-259).
Absolute structure parameter: 0.05 (12)

*Special details*

*Geometry*. All e.s.d.'s (except the e.s.d. in the dihedral angle between two l.s. planes) are estimated using the full covariance matrix. The cell e.s.d.'s are taken into account individually in the estimation of e.s.d.'s in distances, angles and torsion angles; correlations between e.s.d.'s in cell parameters are only used when they are defined by crystal symmetry. An approximate (isotropic) treatment of cell e.s.d.'s is used for estimating e.s.d.'s involving l.s. planes.

*Fractional atomic coordinates and isotropic or equivalent isotropic displacement parameters ($Å^2$)*

| | $x$ | $y$ | $z$ | $U_{iso}$*/$U_{eq}$ |
|---|---|---|---|---|
| O1 | 0.0663 (5) | 0.6534 (3) | 0.23756 (10) | 0.0400 (7) |
| O2 | 0.2980 (4) | 0.5346 (2) | 0.28549 (9) | 0.0346 (6) |
| O3 | 0.4535 (5) | 0.1970 (3) | 0.32957 (11) | 0.0431 (7) |
| O4 | 0.0429 (6) | 0.6670 (3) | 0.56504 (10) | 0.0489 (8) |
| N1 | 0.2241 (6) | 0.3634 (3) | 0.36024 (11) | 0.0333 (7) |
| C1 | 0.1841 (6) | 0.5542 (4) | 0.24226 (13) | 0.0323 (8) |
| C2 | 0.2323 (8) | 0.4382 (4) | 0.20710 (15) | 0.0453 (10) |
| H2A | 0.3019 | 0.4750 | 0.1763 | 0.054* |
| H2B | 0.0929 | 0.3894 | 0.1979 | 0.054* |
| C3 | 0.3871 (9) | 0.3404 (4) | 0.23293 (14) | 0.0455 (11) |
| H3A | 0.5271 | 0.3297 | 0.2138 | 0.055* |
| H3B | 0.3177 | 0.2473 | 0.2371 | 0.055* |
| C4 | 0.4319 (6) | 0.4089 (4) | 0.28411 (13) | 0.0314 (8) |
| C5 | 0.6780 (6) | 0.4507 (4) | 0.28962 (14) | 0.0340 (8) |
| H5A | 0.7704 | 0.3651 | 0.2893 | 0.041* |
| H5B | 0.7217 | 0.5080 | 0.2605 | 0.041* |
| C6 | 0.7254 (7) | 0.5292 (4) | 0.33513 (15) | 0.0364 (9) |
| C7 | 0.7660 (8) | 0.5932 (4) | 0.37185 (16) | 0.0449 (10) |
| H7 | 0.7986 | 0.6445 | 0.4013 | 0.054* |
| C8 | 0.3673 (6) | 0.3122 (4) | 0.32704 (13) | 0.0324 (8) |
| C9 | 0.1595 (7) | 0.2838 (4) | 0.40484 (13) | 0.0367 (9) |
| H9 | 0.2895 | 0.2244 | 0.4145 | 0.044* |
| C10 | −0.0334 (8) | 0.1858 (5) | 0.39303 (15) | 0.0455 (10) |
| H10A | −0.1652 | 0.2408 | 0.3842 | 0.068* |
| H10B | −0.0665 | 0.1282 | 0.4223 | 0.068* |
| H10C | 0.0072 | 0.1252 | 0.3650 | 0.068* |
| C11 | 0.1179 (7) | 0.3857 (4) | 0.44743 (13) | 0.0353 (9) |
| C12 | −0.0791 (7) | 0.3848 (4) | 0.47521 (14) | 0.0389 (9) |
| H12 | −0.1933 | 0.3199 | 0.4672 | 0.047* |
| C13 | −0.1100 (7) | 0.4780 (4) | 0.51452 (14) | 0.0405 (9) |
| H13 | −0.2448 | 0.4766 | 0.5330 | 0.049* |
| C14 | 0.0550 (8) | 0.5722 (4) | 0.52659 (13) | 0.0398 (9) |
| C15 | 0.2524 (8) | 0.5752 (4) | 0.49891 (15) | 0.0420 (10) |
| H15 | 0.3661 | 0.6404 | 0.5069 | 0.050* |
| C16 | 0.2809 (7) | 0.4829 (4) | 0.45994 (14) | 0.0410 (9) |
| H16 | 0.4150 | 0.4857 | 0.4412 | 0.049* |
| C17 | −0.1460 (10) | 0.6594 (5) | 0.59696 (16) | 0.0568 (12) |
| H17A | −0.2819 | 0.6717 | 0.5772 | 0.085* |
| H17B | −0.1368 | 0.7333 | 0.6222 | 0.085* |
| H17C | −0.1494 | 0.5680 | 0.6135 | 0.085* |
| H1 | 0.165 (6) | 0.450 (4) | 0.3571 (12) | 0.010 (7)* |

*Atomic displacement parameters (Å²)*

| | $U^{11}$ | $U^{22}$ | $U^{33}$ | $U^{12}$ | $U^{13}$ | $U^{23}$ |
|---|---|---|---|---|---|---|
| O1 | 0.0348 (14) | 0.0369 (15) | 0.0483 (15) | 0.0027 (12) | 0.0028 (12) | 0.0129 (12) |
| O2 | 0.0432 (15) | 0.0290 (12) | 0.0316 (12) | 0.0077 (11) | 0.0015 (11) | 0.0010 (10) |
| O3 | 0.0412 (15) | 0.0328 (13) | 0.0554 (16) | 0.0058 (12) | 0.0060 (14) | 0.0115 (13) |
| O4 | 0.0554 (19) | 0.0545 (19) | 0.0369 (14) | −0.0105 (15) | 0.0047 (13) | −0.0022 (13) |
| N1 | 0.0380 (18) | 0.0318 (16) | 0.0303 (15) | 0.0034 (15) | −0.0009 (13) | 0.0060 (12) |
| C1 | 0.0328 (18) | 0.0302 (18) | 0.0340 (18) | −0.0054 (16) | 0.0040 (15) | 0.0087 (14) |
| C2 | 0.054 (2) | 0.041 (2) | 0.041 (2) | 0.000 (2) | −0.010 (2) | −0.0026 (16) |
| C3 | 0.063 (3) | 0.038 (2) | 0.0356 (19) | 0.008 (2) | −0.005 (2) | −0.0065 (16) |
| C4 | 0.0352 (19) | 0.0264 (16) | 0.0325 (17) | −0.0002 (15) | 0.0040 (16) | 0.0024 (14) |
| C5 | 0.0339 (19) | 0.0297 (17) | 0.0385 (19) | 0.0017 (15) | 0.0080 (16) | 0.0017 (14) |
| C6 | 0.0338 (19) | 0.0297 (17) | 0.046 (2) | −0.0017 (16) | 0.0032 (17) | 0.0041 (16) |
| C7 | 0.043 (2) | 0.044 (2) | 0.048 (2) | −0.0027 (19) | −0.003 (2) | −0.0055 (18) |
| C8 | 0.0330 (18) | 0.0282 (17) | 0.0360 (17) | −0.0024 (15) | −0.0041 (16) | 0.0053 (14) |
| C9 | 0.043 (2) | 0.0360 (19) | 0.0312 (18) | −0.0028 (17) | −0.0036 (17) | 0.0087 (15) |
| C10 | 0.053 (3) | 0.046 (2) | 0.0380 (19) | −0.013 (2) | −0.0082 (19) | 0.0059 (17) |
| C11 | 0.042 (2) | 0.0365 (19) | 0.0276 (17) | −0.0024 (16) | −0.0028 (16) | 0.0103 (14) |
| C12 | 0.042 (2) | 0.041 (2) | 0.0342 (18) | −0.0101 (17) | −0.0038 (17) | 0.0073 (16) |
| C13 | 0.042 (2) | 0.047 (2) | 0.0331 (19) | −0.0027 (19) | 0.0044 (17) | 0.0096 (16) |
| C14 | 0.052 (2) | 0.039 (2) | 0.0282 (17) | −0.0008 (19) | −0.0063 (17) | 0.0074 (15) |
| C15 | 0.043 (2) | 0.047 (2) | 0.0357 (19) | −0.010 (2) | −0.0025 (18) | 0.0032 (16) |
| C16 | 0.043 (2) | 0.048 (2) | 0.0322 (19) | −0.0079 (19) | −0.0014 (17) | 0.0056 (16) |
| C17 | 0.069 (3) | 0.065 (3) | 0.036 (2) | −0.008 (3) | 0.009 (2) | −0.003 (2) |

*Geometric parameters (Å, º)*

| | | | |
|---|---|---|---|
| O1—C1 | 1.191 (4) | C6—C7 | 1.185 (6) |
| O2—C1 | 1.359 (4) | C7—H7 | 0.9500 |
| O2—C4 | 1.447 (4) | C9—C11 | 1.522 (5) |
| O3—C8 | 1.219 (5) | C9—C10 | 1.524 (6) |
| O4—C14 | 1.375 (5) | C9—H9 | 1.0000 |
| O4—C17 | 1.424 (6) | C10—H10A | 0.9800 |
| N1—C8 | 1.332 (5) | C10—H10B | 0.9800 |
| N1—C9 | 1.470 (5) | C10—H10C | 0.9800 |
| N1—H1 | 0.91 (4) | C11—C16 | 1.392 (6) |
| C1—C2 | 1.485 (5) | C11—C12 | 1.399 (6) |
| C2—C3 | 1.491 (6) | C12—C13 | 1.393 (6) |
| C2—H2A | 0.9900 | C12—H12 | 0.9500 |
| C2—H2B | 0.9900 | C13—C14 | 1.379 (6) |
| C3—C4 | 1.545 (5) | C13—H13 | 0.9500 |
| C3—H3A | 0.9900 | C14—C15 | 1.400 (6) |
| C3—H3B | 0.9900 | C15—C16 | 1.379 (6) |
| C4—C8 | 1.528 (5) | C15—H15 | 0.9500 |
| C4—C5 | 1.539 (5) | C16—H16 | 0.9500 |
| C5—C6 | 1.461 (5) | C17—H17A | 0.9800 |
| C5—H5A | 0.9900 | C17—H17B | 0.9800 |
| C5—H5B | 0.9900 | C17—H17C | 0.9800 |
| C1—O2—C4 | 111.9 (3) | N1—C9—C11 | 108.8 (3) |
| C14—O4—C17 | 117.4 (4) | N1—C9—C10 | 110.5 (3) |

| | | | |
|---|---|---|---|
| C8—N1—C9 | 121.6 (3) | C11—C9—C10 | 115.2 (4) |
| C8—N1—H1 | 122 (2) | N1—C9—H9 | 107.3 |
| C9—N1—H1 | 116 (2) | C11—C9—H9 | 107.3 |
| O1—C1—O2 | 120.0 (3) | C10—C9—H9 | 107.3 |
| O1—C1—C2 | 130.1 (3) | C9—C10—H10A | 109.5 |
| O2—C1—C2 | 109.9 (3) | C9—C10—H10B | 109.5 |
| C1—C2—C3 | 107.2 (3) | H10A—C10—H10B | 109.5 |
| C1—C2—H2A | 110.3 | C9—C10—H10C | 109.5 |
| C3—C2—H2A | 110.3 | H10A—C10—H10C | 109.5 |
| C1—C2—H2B | 110.3 | H10B—C10—H10C | 109.5 |
| C3—C2—H2B | 110.3 | C16—C11—C12 | 118.1 (4) |
| H2A—C2—H2B | 108.5 | C16—C11—C9 | 119.6 (4) |
| C2—C3—C4 | 104.8 (3) | C12—C11—C9 | 122.3 (4) |
| C2—C3—H3A | 110.8 | C13—C12—C11 | 120.8 (4) |
| C4—C3—H3A | 110.8 | C13—C12—H12 | 119.6 |
| C2—C3—H3B | 110.8 | C11—C12—H12 | 119.6 |
| C4—C3—H3B | 110.8 | C14—C13—C12 | 120.0 (4) |
| H3A—C3—H3B | 108.9 | C14—C13—H13 | 120.0 |
| O2—C4—C8 | 110.1 (3) | C12—C13—H13 | 120.0 |
| O2—C4—C5 | 108.4 (3) | O4—C14—C13 | 124.7 (4) |
| C8—C4—C5 | 109.2 (3) | O4—C14—C15 | 115.4 (4) |
| O2—C4—C3 | 106.2 (3) | C13—C14—C15 | 119.9 (4) |
| C8—C4—C3 | 111.6 (3) | C16—C15—C14 | 119.6 (4) |
| C5—C4—C3 | 111.3 (3) | C16—C15—H15 | 120.2 |
| C6—C5—C4 | 113.7 (3) | C14—C15—H15 | 120.2 |
| C6—C5—H5A | 108.8 | C15—C16—C11 | 121.6 (4) |
| C4—C5—H5A | 108.8 | C15—C16—H16 | 119.2 |
| C6—C5—H5B | 108.8 | C11—C16—H16 | 119.2 |
| C4—C5—H5B | 108.8 | O4—C17—H17A | 109.5 |
| H5A—C5—H5B | 107.7 | O4—C17—H17B | 109.5 |
| C7—C6—C5 | 179.3 (4) | H17A—C17—H17B | 109.5 |
| C6—C7—H7 | 180.0 | O4—C17—H17C | 109.5 |
| O3—C8—N1 | 124.7 (3) | H17A—C17—H17C | 109.5 |
| O3—C8—C4 | 118.8 (3) | H17B—C17—H17C | 109.5 |
| N1—C8—C4 | 116.4 (3) | | |

| | | | |
|---|---|---|---|
| C4—O2—C1—O1 | −180.0 (3) | C5—C4—C8—N1 | 111.8 (4) |
| C4—O2—C1—C2 | 0.5 (4) | C3—C4—C8—N1 | −124.7 (4) |
| O1—C1—C2—C3 | −178.5 (4) | C8—N1—C9—C11 | 146.8 (4) |
| O2—C1—C2—C3 | 1.0 (5) | C8—N1—C9—C10 | −85.8 (5) |
| C1—C2—C3—C4 | −1.9 (5) | N1—C9—C11—C16 | −53.0 (5) |
| C1—O2—C4—C8 | −122.7 (3) | C10—C9—C11—C16 | −177.7 (3) |
| C1—O2—C4—C5 | 117.9 (3) | N1—C9—C11—C12 | 127.6 (4) |
| C1—O2—C4—C3 | −1.8 (4) | C10—C9—C11—C12 | 2.9 (5) |
| C2—C3—C4—O2 | 2.2 (4) | C16—C11—C12—C13 | −0.5 (5) |
| C2—C3—C4—C8 | 122.2 (4) | C9—C11—C12—C13 | 178.9 (4) |
| C2—C3—C4—C5 | −115.6 (4) | C11—C12—C13—C14 | −0.2 (6) |
| O2—C4—C5—C6 | 57.5 (4) | C17—O4—C14—C13 | 5.2 (6) |
| C8—C4—C5—C6 | −62.4 (4) | C17—O4—C14—C15 | −174.5 (4) |
| C3—C4—C5—C6 | 173.9 (3) | C12—C13—C14—O4 | −179.0 (4) |
| C9—N1—C8—O3 | 1.4 (6) | C12—C13—C14—C15 | 0.8 (6) |
| C9—N1—C8—C4 | −176.3 (3) | O4—C14—C15—C16 | 179.3 (3) |
| O2—C4—C8—O3 | 175.1 (3) | C13—C14—C15—C16 | −0.5 (6) |
| C5—C4—C8—O3 | −66.0 (4) | C14—C15—C16—C11 | −0.3 (6) |
| C3—C4—C8—O3 | 57.5 (5) | C12—C11—C16—C15 | 0.8 (6) |
| O2—C4—C8—N1 | −7.0 (5) | C9—C11—C16—C15 | −178.6 (4) |

**4.12-1**

**Table 1**

Experimental details

| | |
|---|---|
| Crystal data | |
| Chemical formula | $2(C_{17}H_{19}NO_4)\cdot H_2O$ |
| $M_r$ | 620.68 |
| Crystal system, space group | Monoclinic, *C*2 |
| Temperature (K) | 150 |
| *a*, *b*, *c* (Å) | 21.0408 (11), 5.4841 (3), 13.5744 (9) |
| $\beta$ (°) | 101.620 (3) |
| *V* (Å$^3$) | 1534.25 (16) |
| *Z* | 2 |
| Radiation type | Cu $K\alpha$ |
| $\mu$ (mm$^{-1}$) | 0.80 |
| Crystal size (mm) | 0.36 × 0.25 × 0.20 |
| Data collection | |
| Diffractometer | Bruker D8 VENTURE FIXED-CHI PHOTON 100 CMOS |
| Absorption correction | Multi-scan<br>*SADABS* 2014/5 (Sheldrick, 2014) |
| $T_{min}$, $T_{max}$ | 0.632, 0.753 |
| No. of measured, independent and observed [$I > 2\sigma(I)$] reflections | 5794, 2547, 2413 |
| $R_{int}$ | 0.038 |
| $(\sin \theta/\lambda)_{max}$ (Å$^{-1}$) | 0.595 |
| Refinement | |
| $R[F^2 > 2\sigma(F^2)]$, $wR(F^2)$, $S$ | 0.031, 0.079, 1.06 |
| No. of reflections | 2547 |
| No. of parameters | 215 |
| No. of restraints | 1 |
| H-atom treatment | H atoms treated by a mixture of independent and constrained refinement |
| $\Delta\rho_{max}$, $\Delta\rho_{min}$ (e Å$^{-3}$) | 0.17, −0.19 |
| Absolute structure | Flack x determined using 940 quotients [(I+)-(I-)]/[(I+)+(I-)] (Parsons, Flack and Wagner, Acta Cryst. B69 (2013) 249-259). |
| Absolute structure parameter | 0.09 (13) |

Computer programs: *APEX3* v2015.5-2 (Bruker-AXS, 2015), *SAINT* v8.34A (Bruker-AXS, 2007), *SHELXT2014/5* (Sheldrick, 2015), *SHELXL2014/7* (Sheldrick, 2014), Bruker *SHELXTL*.

**Acknowledgements**

The authors thank The University of Hong Kong's University Development Fund and the Dr. Hui Wai Haan Fund for funding the Bruker D8 VENTURE Photon100 CMOS X-Ray Diffractometer.

**Funding information**

**Figure 1**

Fig. 1. *ORTEP* diagram of the molecule with thermal ellipsoids at 50% probability level. co-crystallized water molecule is omitted for clarity.

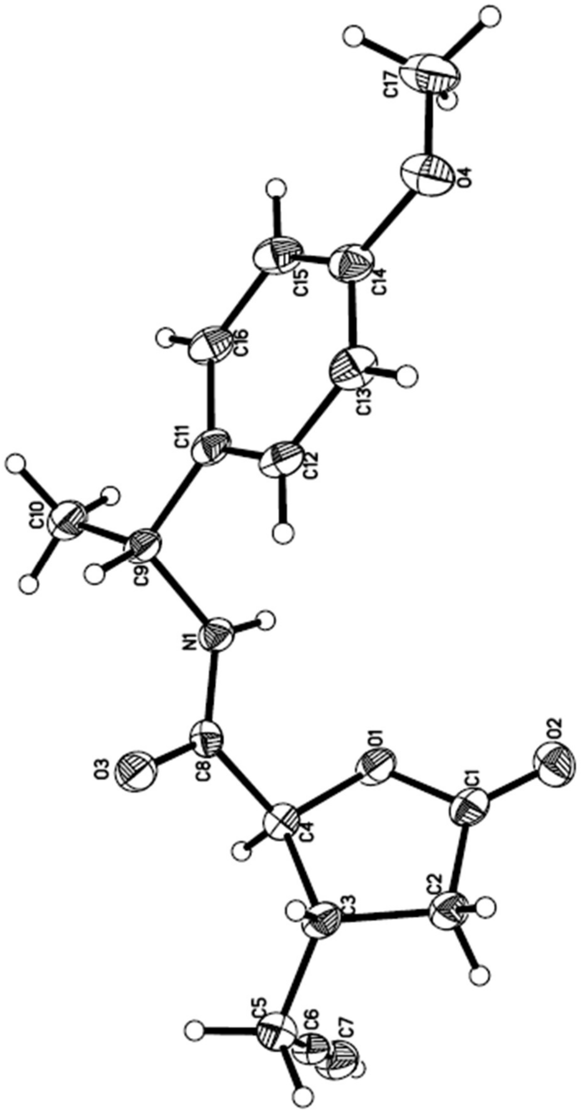
C17
O4
C15
C14
C16
C13
C11
C12
C10
C9
N1
O3
C8
O1
C1
O2
C4
C2
C3
C5
C6
C7

### Title

#### Computing details

Data collection: *APEX3* v2015.5-2 (Bruker-AXS, 2015); cell refinement: *SAINT* v8.34A (Bruker-AXS, 2007); data reduction: *SAINT* v8.34A (Bruker-AXS, 2007); program(s) used to solve structure: *SHELXT2014/5* (Sheldrick, 2015); program(s) used to refine structure: *SHELXL2014/7* (Sheldrick, 2014); molecular graphics: Bruker *SHELXTL*; software used to prepare material for publication: Bruker *SHELXTL*.

#### (cu_CHEM0415)

##### *Crystal data*

$2(C_{17}H_{19}NO_4)\cdot H_2O$
$M_r = 620.68$
Monoclinic, $C2$
$a = 21.0408$ (11) Å
$b = 5.4841$ (3) Å
$c = 13.5744$ (9) Å
$\beta = 101.620$ (3)°
$V = 1534.25$ (16) Å$^3$
$Z = 2$

$F(000) = 660$
$D_x = 1.344$ Mg m$^{-3}$
Cu $K\alpha$ radiation, $\lambda = 1.54178$ Å
Cell parameters from 223 reflections
$\theta = 3–45°$
$\mu = 0.80$ mm$^{-1}$
$T = 150$ K
Rod, colourless
0.36 × 0.25 × 0.20 mm

##### *Data collection*

Bruker D8 VENTURE FIXED-CHI PHOTON 100 CMOS diffractometer
Radiation source: I$\mu$S HB micro-focus sealed tube
Detector resolution: 10.24 pixels mm$^{-1}$
$\varphi$ and $\omega$ scans
Absorption correction: multi-scan *SADABS* 2014/5 (Sheldrick, 2014)
$T_{min} = 0.632$, $T_{max} = 0.753$

5794 measured reflections
2547 independent reflections
2413 reflections with $I > 2\sigma(I)$
$R_{int} = 0.038$
$\theta_{max} = 66.6°$, $\theta_{min} = 3.3°$
$h = -24 \rightarrow 22$
$k = -6 \rightarrow 6$
$l = -16 \rightarrow 16$

##### *Refinement*

Refinement on $F^2$
Least-squares matrix: full
$R[F^2 > 2\sigma(F^2)] = 0.031$
$wR(F^2) = 0.079$
$S = 1.06$
2547 reflections
215 parameters
1 restraint
Hydrogen site location: mixed
H atoms treated by a mixture of independent and constrained refinement

$w = 1/[\sigma^2(F_o^2) + (0.0303P)^2 + 0.8254P]$ where $P = (F_o^2 + 2F_c^2)/3$
$(\Delta/\sigma)_{max} < 0.001$
$\Delta\rho_{max} = 0.17$ e Å$^{-3}$
$\Delta\rho_{min} = -0.19$ e Å$^{-3}$
Extinction correction: *SHELXL2014/7* (Sheldrick 2014, $Fc^* = kFc[1+0.001xFc^2\lambda^3/\sin(2\theta)]^{-1/4}$
Extinction coefficient: 0.0027 (3)
Absolute structure: Flack x determined using 940 quotients [(I+)-(I-)]/[(I+)+(I-)] (Parsons, Flack and Wagner, Acta Cryst. B69 (2013) 249-259).
Absolute structure parameter: 0.09 (13)

*Special details*

*Geometry*. All e.s.d.'s (except the e.s.d. in the dihedral angle between two l.s. planes) are estimated using the full covariance matrix. The cell e.s.d.'s are taken into account individually in the estimation of e.s.d.'s in distances, angles and torsion angles; correlations between e.s.d.'s in cell parameters are only used when they are defined by crystal symmetry. An approximate (isotropic) treatment of cell e.s.d.'s is used for estimating e.s.d.'s involving l.s. planes.

*Fractional atomic coordinates and isotropic or equivalent isotropic displacement parameters (Å²)*

| | *x* | *y* | *z* | $U_{iso}$*/$U_{eq}$ |
|---|---|---|---|---|
| O1 | 0.67566 (8) | 0.7695 (3) | 0.45223 (13) | 0.0251 (4) |
| O2 | 0.77011 (8) | 0.7057 (4) | 0.40681 (13) | 0.0298 (5) |
| O3 | 0.57687 (9) | 0.4329 (4) | 0.58630 (13) | 0.0289 (4) |
| O4 | 0.89063 (9) | 0.6776 (4) | 0.97384 (14) | 0.0326 (5) |
| O5 | 0.5000 | 0.0265 (5) | 0.5000 | 0.0314 (6) |
| H2 | 0.5279 (17) | 0.110 (8) | 0.533 (3) | 0.058 (12)* |
| N1 | 0.64128 (9) | 0.7609 (4) | 0.63164 (15) | 0.0194 (4) |
| H1 | 0.6643 (13) | 0.880 (6) | 0.611 (2) | 0.020 (7)* |
| C1 | 0.71596 (12) | 0.6355 (5) | 0.40761 (17) | 0.0222 (5) |
| C2 | 0.68243 (12) | 0.4103 (5) | 0.36192 (19) | 0.0243 (6) |
| H2A | 0.7089 | 0.2641 | 0.3851 | 0.029* |
| H2B | 0.6753 | 0.4181 | 0.2876 | 0.029* |
| C3 | 0.61780 (12) | 0.3991 (5) | 0.39608 (18) | 0.0221 (5) |
| H3 | 0.6184 | 0.2578 | 0.4428 | 0.026* |
| C4 | 0.61638 (11) | 0.6379 (5) | 0.45544 (18) | 0.0209 (5) |
| H4 | 0.5787 | 0.7380 | 0.4210 | 0.025* |
| C5 | 0.55862 (12) | 0.3752 (5) | 0.3093 (2) | 0.0288 (6) |
| H5A | 0.5611 | 0.2171 | 0.2750 | 0.035* |
| H5B | 0.5187 | 0.3743 | 0.3374 | 0.035* |
| C6 | 0.55400 (12) | 0.5714 (6) | 0.23527 (19) | 0.0280 (6) |
| C7 | 0.55087 (13) | 0.7333 (6) | 0.1767 (2) | 0.0342 (6) |
| H7 | 0.5484 | 0.8631 | 0.1297 | 0.041* |
| C8 | 0.61052 (11) | 0.5993 (5) | 0.56504 (18) | 0.0197 (5) |
| C9 | 0.63525 (11) | 0.7574 (5) | 0.73747 (17) | 0.0208 (5) |
| H9 | 0.6121 | 0.6038 | 0.7485 | 0.025* |
| C10 | 0.59353 (13) | 0.9690 (5) | 0.7581 (2) | 0.0283 (6) |
| H10A | 0.6147 | 1.1229 | 0.7471 | 0.042* |
| H10B | 0.5878 | 0.9611 | 0.8280 | 0.042* |
| H10C | 0.5510 | 0.9598 | 0.7126 | 0.042* |
| C11 | 0.70251 (11) | 0.7462 (5) | 0.80469 (17) | 0.0204 (5) |
| C12 | 0.74144 (12) | 0.5435 (5) | 0.79722 (17) | 0.0229 (5) |
| H12 | 0.7250 | 0.4157 | 0.7520 | 0.028* |
| C13 | 0.80338 (13) | 0.5258 (5) | 0.85440 (19) | 0.0256 (6) |
| H13 | 0.8291 | 0.3864 | 0.8484 | 0.031* |
| C14 | 0.82827 (12) | 0.7123 (5) | 0.92110 (18) | 0.0245 (5) |
| C15 | 0.79014 (13) | 0.9110 (5) | 0.93107 (19) | 0.0263 (6) |
| H15 | 0.8064 | 1.0367 | 0.9775 | 0.032* |
| C16 | 0.72748 (13) | 0.9275 (5) | 0.87283 (18) | 0.0240 (5) |
| H16 | 0.7015 | 1.0655 | 0.8800 | 0.029* |
| C17 | 0.91838 (14) | 0.8710 (6) | 1.0388 (2) | 0.0379 (7) |
| H17A | 0.9180 | 1.0214 | 0.9998 | 0.057* |
| H17B | 0.9632 | 0.8293 | 1.0702 | 0.057* |
| H17C | 0.8930 | 0.8950 | 1.0912 | 0.057* |

*Atomic displacement parameters (Å²)*

| | $U^{11}$ | $U^{22}$ | $U^{33}$ | $U^{12}$ | $U^{13}$ | $U^{23}$ |
|---|---|---|---|---|---|---|
| O1 | 0.0337 (9) | 0.0223 (9) | 0.0217 (9) | −0.0066 (8) | 0.0112 (7) | −0.0038 (7) |
| O2 | 0.0273 (9) | 0.0361 (11) | 0.0266 (9) | −0.0048 (8) | 0.0071 (7) | 0.0066 (8) |
| O3 | 0.0331 (9) | 0.0298 (10) | 0.0262 (10) | −0.0127 (9) | 0.0115 (8) | −0.0067 (8) |
| O4 | 0.0304 (9) | 0.0364 (12) | 0.0276 (10) | −0.0027 (9) | −0.0026 (7) | −0.0001 (9) |
| O5 | 0.0300 (14) | 0.0214 (14) | 0.0384 (16) | 0.000 | −0.0035 (12) | 0.000 |
| N1 | 0.0219 (9) | 0.0205 (11) | 0.0162 (10) | −0.0026 (9) | 0.0050 (8) | −0.0001 (9) |
| C1 | 0.0284 (13) | 0.0242 (14) | 0.0144 (11) | 0.0034 (11) | 0.0054 (9) | 0.0052 (10) |
| C2 | 0.0245 (12) | 0.0235 (13) | 0.0249 (13) | 0.0023 (11) | 0.0048 (10) | −0.0046 (11) |
| C3 | 0.0272 (13) | 0.0198 (14) | 0.0203 (12) | −0.0012 (11) | 0.0075 (10) | −0.0034 (10) |
| C4 | 0.0220 (11) | 0.0211 (13) | 0.0201 (12) | 0.0006 (10) | 0.0054 (9) | −0.0004 (10) |
| C5 | 0.0265 (13) | 0.0328 (16) | 0.0279 (14) | −0.0067 (12) | 0.0076 (11) | −0.0072 (12) |
| C6 | 0.0209 (12) | 0.0416 (18) | 0.0207 (12) | −0.0025 (12) | 0.0021 (9) | −0.0117 (13) |
| C7 | 0.0315 (13) | 0.0446 (18) | 0.0250 (14) | 0.0028 (14) | 0.0020 (10) | −0.0054 (14) |
| C8 | 0.0163 (10) | 0.0217 (13) | 0.0217 (12) | 0.0008 (10) | 0.0049 (9) | −0.0008 (10) |
| C9 | 0.0264 (12) | 0.0213 (13) | 0.0158 (11) | −0.0031 (11) | 0.0069 (9) | −0.0010 (10) |
| C10 | 0.0281 (13) | 0.0338 (17) | 0.0236 (13) | 0.0029 (11) | 0.0067 (10) | −0.0034 (11) |
| C11 | 0.0278 (12) | 0.0197 (12) | 0.0149 (11) | −0.0026 (11) | 0.0071 (9) | 0.0021 (10) |
| C12 | 0.0318 (13) | 0.0215 (13) | 0.0156 (12) | −0.0037 (11) | 0.0052 (10) | −0.0021 (10) |
| C13 | 0.0326 (13) | 0.0208 (13) | 0.0235 (13) | 0.0037 (11) | 0.0058 (10) | −0.0002 (11) |
| C14 | 0.0288 (12) | 0.0269 (14) | 0.0171 (12) | −0.0024 (11) | 0.0026 (9) | 0.0045 (10) |
| C15 | 0.0364 (14) | 0.0195 (13) | 0.0219 (13) | −0.0049 (12) | 0.0031 (11) | −0.0030 (11) |
| C16 | 0.0340 (13) | 0.0185 (12) | 0.0195 (12) | 0.0013 (11) | 0.0054 (10) | −0.0007 (10) |
| C17 | 0.0392 (16) | 0.0432 (19) | 0.0265 (15) | −0.0149 (14) | −0.0050 (12) | 0.0038 (13) |

*Geometric parameters (Å, °)*

| | | | |
|---|---|---|---|
| O1—C1 | 1.354 (3) | C6—C7 | 1.185 (5) |
| O1—C4 | 1.449 (3) | C7—H7 | 0.9500 |
| O2—C1 | 1.205 (3) | C9—C10 | 1.515 (4) |
| O3—C8 | 1.225 (3) | C9—C11 | 1.523 (3) |
| O4—C14 | 1.375 (3) | C9—H9 | 1.0000 |
| O4—C17 | 1.427 (4) | C10—H10A | 0.9800 |
| O5—H2 | 0.81 (4) | C10—H10B | 0.9800 |
| N1—C8 | 1.336 (3) | C10—H10C | 0.9800 |
| N1—C9 | 1.468 (3) | C11—C16 | 1.387 (4) |
| N1—H1 | 0.89 (3) | C11—C12 | 1.397 (4) |
| C1—C2 | 1.494 (4) | C12—C13 | 1.379 (3) |
| C2—C3 | 1.524 (3) | C12—H12 | 0.9500 |
| C2—H2A | 0.9900 | C13—C14 | 1.396 (4) |
| C2—H2B | 0.9900 | C13—H13 | 0.9500 |
| C3—C5 | 1.538 (3) | C14—C15 | 1.376 (4) |
| C3—C4 | 1.541 (3) | C15—C16 | 1.397 (4) |
| C3—H3 | 1.0000 | C15—H15 | 0.9500 |
| C4—C8 | 1.532 (3) | C16—H16 | 0.9500 |
| C4—H4 | 1.0000 | C17—H17A | 0.9800 |
| C5—C6 | 1.462 (4) | C17—H17B | 0.9800 |
| C5—H5A | 0.9900 | C17—H17C | 0.9800 |
| C5—H5B | 0.9900 | | |

| | | | |
|---|---|---|---|
| C1—O1—C4 | 111.31 (19) | N1—C9—C10 | 109.8 (2) |
| C14—O4—C17 | 116.6 (2) | N1—C9—C11 | 109.51 (18) |
| C8—N1—C9 | 121.9 (2) | C10—C9—C11 | 115.0 (2) |
| C8—N1—H1 | 119.3 (17) | N1—C9—H9 | 107.4 |
| C9—N1—H1 | 118.7 (17) | C10—C9—H9 | 107.4 |
| O2—C1—O1 | 120.6 (2) | C11—C9—H9 | 107.4 |
| O2—C1—C2 | 129.0 (2) | C9—C10—H10A | 109.5 |
| O1—C1—C2 | 110.4 (2) | C9—C10—H10B | 109.5 |
| C1—C2—C3 | 106.4 (2) | H10A—C10—H10B | 109.5 |
| C1—C2—H2A | 110.4 | C9—C10—H10C | 109.5 |
| C3—C2—H2A | 110.4 | H10A—C10—H10C | 109.5 |
| C1—C2—H2B | 110.4 | H10B—C10—H10C | 109.5 |
| C3—C2—H2B | 110.4 | C16—C11—C12 | 118.0 (2) |
| H2A—C2—H2B | 108.6 | C16—C11—C9 | 123.5 (2) |
| C2—C3—C5 | 113.9 (2) | C12—C11—C9 | 118.4 (2) |
| C2—C3—C4 | 103.7 (2) | C13—C12—C11 | 121.1 (2) |
| C5—C3—C4 | 112.0 (2) | C13—C12—H12 | 119.4 |
| C2—C3—H3 | 109.0 | C11—C12—H12 | 119.4 |
| C5—C3—H3 | 109.0 | C12—C13—C14 | 120.1 (3) |
| C4—C3—H3 | 109.0 | C12—C13—H13 | 119.9 |
| O1—C4—C8 | 109.63 (18) | C14—C13—H13 | 119.9 |
| O1—C4—C3 | 107.46 (19) | O4—C14—C15 | 124.9 (2) |
| C8—C4—C3 | 113.9 (2) | O4—C14—C13 | 115.5 (2) |
| O1—C4—H4 | 108.6 | C15—C14—C13 | 119.6 (2) |
| C8—C4—H4 | 108.6 | C14—C15—C16 | 119.9 (2) |
| C3—C4—H4 | 108.6 | C14—C15—H15 | 120.1 |
| C6—C5—C3 | 113.3 (2) | C16—C15—H15 | 120.1 |
| C6—C5—H5A | 108.9 | C11—C16—C15 | 121.2 (3) |
| C3—C5—H5A | 108.9 | C11—C16—H16 | 119.4 |
| C6—C5—H5B | 108.9 | C15—C16—H16 | 119.4 |
| C3—C5—H5B | 108.9 | O4—C17—H17A | 109.5 |
| H5A—C5—H5B | 107.7 | O4—C17—H17B | 109.5 |
| C7—C6—C5 | 178.7 (3) | H17A—C17—H17B | 109.5 |
| C6—C7—H7 | 180.0 | O4—C17—H17C | 109.5 |
| O3—C8—N1 | 124.0 (2) | H17A—C17—H17C | 109.5 |
| O3—C8—C4 | 119.7 (2) | H17B—C17—H17C | 109.5 |
| N1—C8—C4 | 116.3 (2) | | |

| | | | |
|---|---|---|---|
| C4—O1—C1—O2 | −172.8 (2) | C3—C4—C8—N1 | 146.1 (2) |
| C4—O1—C1—C2 | 8.6 (3) | C8—N1—C9—C10 | −107.1 (3) |
| O2—C1—C2—C3 | 173.8 (2) | C8—N1—C9—C11 | 125.7 (2) |
| O1—C1—C2—C3 | −7.9 (3) | N1—C9—C11—C16 | 117.5 (3) |
| C1—C2—C3—C5 | 126.0 (2) | C10—C9—C11—C16 | −6.7 (3) |
| C1—C2—C3—C4 | 4.0 (3) | N1—C9—C11—C12 | −62.1 (3) |
| C1—O1—C4—C8 | 118.4 (2) | C10—C9—C11—C12 | 173.7 (2) |
| C1—O1—C4—C3 | −5.8 (3) | C16—C11—C12—C13 | −1.2 (4) |
| C2—C3—C4—O1 | 0.7 (2) | C9—C11—C12—C13 | 178.4 (2) |
| C5—C3—C4—O1 | −122.6 (2) | C11—C12—C13—C14 | −0.1 (4) |
| C2—C3—C4—C8 | −120.9 (2) | C17—O4—C14—C15 | −4.0 (4) |
| C5—C3—C4—C8 | 115.8 (2) | C17—O4—C14—C13 | 177.0 (2) |
| C2—C3—C5—C6 | −56.9 (3) | C12—C13—C14—O4 | −179.5 (2) |
| C4—C3—C5—C6 | 60.4 (3) | C12—C13—C14—C15 | 1.5 (4) |
| C9—N1—C8—O3 | −2.3 (4) | O4—C14—C15—C16 | 179.5 (2) |
| C9—N1—C8—C4 | 174.8 (2) | C13—C14—C15—C16 | −1.5 (4) |
| O1—C4—C8—O3 | −157.1 (2) | C12—C11—C16—C15 | 1.2 (4) |
| C3—C4—C8—O3 | −36.7 (3) | C9—C11—C16—C15 | −178.5 (2) |
| O1—C4—C8—N1 | 25.7 (3) | C14—C15—C16—C11 | 0.2 (4) |

**5.9*R*-2**

### Title

Enter author details here

#### Abstract

##### Table 1

Experimental details

| | |
|---|---|
| Crystal data | |
| Chemical formula | $C_{17}H_{19}NO_4$ |
| $M_r$ | 301.33 |
| Crystal system, space group | Monoclinic, *C*2 |
| Temperature (K) | 173 |
| *a*, *b*, *c* (Å) | 21.788 (2), 5.0518 (6), 14.1121 (16) |
| β (°) | 90.606 (3) |
| *V* (Å$^3$) | 1553.2 (3) |
| *Z* | 4 |
| Radiation type | Mo *K*α |
| μ (mm$^{-1}$) | 0.09 |
| Crystal size (mm) | 0.33 × 0.08 × 0.05 |
| Data collection | |
| Diffractometer | Bruker D8 VENTURE FIXED-CHI PHOTON 100 CMOS |
| Absorption correction | Multi-scan<br>*SADABS* 2014/5 (Sheldrick, 2014) |
| $T_{min}$, $T_{max}$ | 0.369, 0.745 |
| No. of measured, independent and observed [$I > 2\sigma(I)$] reflections | 13005, 2725, 2152 |
| $R_{int}$ | 0.093 |
| $(\sin \theta/\lambda)_{max}$ (Å$^{-1}$) | 0.597 |
| Refinement | |
| $R[F^2 > 2\sigma(F^2)]$, $wR(F^2)$, $S$ | 0.045, 0.105, 1.06 |
| No. of reflections | 2725 |
| No. of parameters | 205 |
| No. of restraints | 13 |
| H-atom treatment | H atoms treated by a mixture of independent and constrained refinement |
| $\Delta\rho_{max}$, $\Delta\rho_{min}$ (e Å$^{-3}$) | 0.18, −0.25 |
| Absolute structure | Flack x determined using 802 quotients [(I+)-(I-)]/[(I+)+(I-)] (Parsons, Flack and Wagner, Acta Cryst. B69 (2013) 249-259). |
| Absolute structure parameter | 1.9 (10) |

Computer programs: *APEX3* v2015.5-2 (Bruker-AXS, 2015), *SAINT* v8.34A (Bruker-AXS, 2013), *SHELXT2014*/5 (Sheldrick, 2015), *SHELXL2014*/7 (Sheldrick, 2014), Bruker *SHELXTL*.

##### Acknowledgements

The authors thank The University of Hong Kong's University Development Fund and the Dr. Hui Wai Haan Fund for funding the Bruker D8 VENTURE Photon100 CMOS X-Ray Diffractometer.

##### Funding information

##### Figure 1

Fig. 1. *ORTEP* diagram of <your molecule> with thermal ellipsoids at 50% probability level.

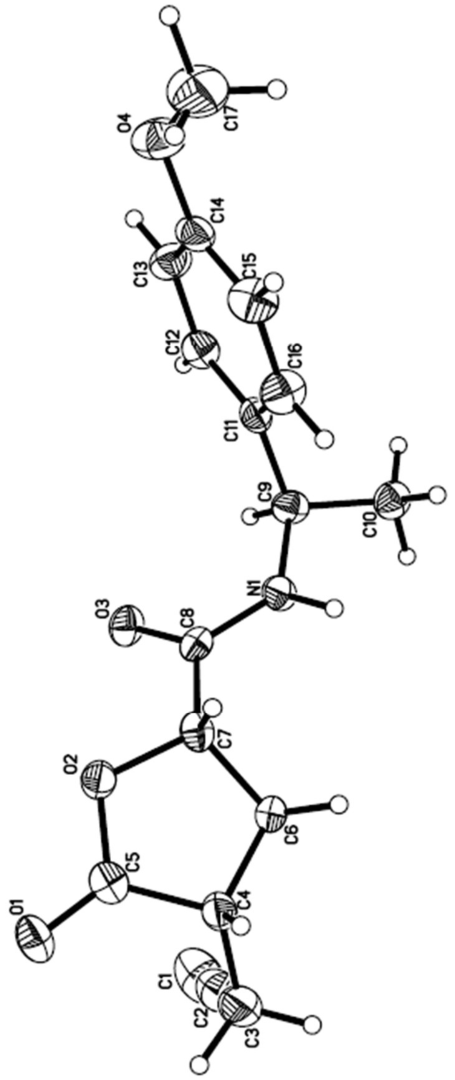
O4
C17
C14
C13
C15
C12
C16
C11
C9
C10
N1
O3
C8
C7
O2
C6
C5
C4
O1
C1
C2
C3

### Title

##### Computing details

Data collection: *APEX3* v2015.5-2 (Bruker-AXS, 2015); cell refinement: *SAINT* v8.34A (Bruker-AXS, 2013); data reduction: *SAINT* v8.34A (Bruker-AXS, 2013); program(s) used to solve structure: *SHELXT2014/5* (Sheldrick, 2015); program(s) used to refine structure: *SHELXL2014/7* (Sheldrick, 2014); molecular graphics: Bruker *SHELXTL*; software used to prepare material for publication: Bruker *SHELXTL*.

##### (mo_CHEM0363)

*Crystal data*

$C_{17}H_{19}NO_4$
$M_r = 301.33$
Monoclinic, *C*2
$a = 21.788$ (2) Å
$b = 5.0518$ (6) Å
$c = 14.1121$ (16) Å
$\beta = 90.606$ (3)°
$V = 1553.2$ (3) Å$^3$
$Z = 4$
$F(000) = 640$
$D_x = 1.289$ Mg m$^{-3}$
Mo $K\alpha$ radiation, $\lambda = 0.71073$ Å
Cell parameters from 153 reflections
$\theta = 2.8$–22.5°
$\mu = 0.09$ mm$^{-1}$
$T = 173$ K
Rod, colorless
0.33 × 0.08 × 0.05 mm

*Data collection*

Bruker D8 VENTURE FIXED-CHI PHOTON 100 CMOS diffractometer
Radiation source: I$\mu$S HB micro-focus sealed tube
Detector resolution: 10.24 pixels mm$^{-1}$
$\varphi$ and $\omega$ scans
Absorption correction: multi-scan *SADABS* 2014/5 (Sheldrick, 2014)
$T_{min} = 0.369$, $T_{max} = 0.745$
13005 measured reflections
2725 independent reflections
2152 reflections with $I > 2\sigma(I)$
$R_{int} = 0.093$
$\theta_{max} = 25.1°$, $\theta_{min} = 2.4°$
$h = -25 \rightarrow 25$
$k = -6 \rightarrow 6$
$l = -16 \rightarrow 16$

*Refinement*

Refinement on $F^2$
Least-squares matrix: full
$R[F^2 > 2\sigma(F^2)] = 0.045$
$wR(F^2) = 0.105$
$S = 1.06$
2725 reflections
205 parameters
13 restraints
Hydrogen site location: mixed
H atoms treated by a mixture of independent and constrained refinement
$w = 1/[\sigma^2(F_o^2) + (0.0207P)^2 + 0.1195P]$ where $P = (F_o^2 + 2F_c^2)/3$
$(\Delta/\sigma)_{max} < 0.001$
$\Delta\rho_{max} = 0.18$ e Å$^{-3}$
$\Delta\rho_{min} = -0.25$ e Å$^{-3}$
Absolute structure: Flack x determined using 802 quotients [(I+)-(I-)]/[(I+)+(I-)] (Parsons, Flack and Wagner, Acta Cryst. B69 (2013) 249-259).
Absolute structure parameter: 1.9 (10)

*Special details*

*Geometry*. All e.s.d.'s (except the e.s.d. in the dihedral angle between two l.s. planes) are estimated using the full covariance matrix. The cell e.s.d.'s are taken into account individually in the estimation of e.s.d.'s in distances, angles and torsion angles; correlations between e.s.d.'s in cell parameters are only used when they are defined by crystal symmetry. An approximate (isotropic) treatment of cell e.s.d.'s is used for estimating e.s.d.'s involving l.s. planes.

*Fractional atomic coordinates and isotropic or equivalent isotropic displacement parameters ($Å^2$)*

| | *x* | *y* | *z* | $U_{iso}$*/$U_{eq}$ |
|---|---|---|---|---|
| O1 | 0.71008 (10) | 0.8046 (5) | 1.07290 (17) | 0.0327 (6) |
| O2 | 0.68115 (9) | 0.6896 (4) | 0.92712 (15) | 0.0253 (6) |
| O3 | 0.60774 (10) | 0.7962 (4) | 0.77922 (15) | 0.0273 (6) |
| O4 | 0.70708 (10) | 0.4441 (6) | 0.33816 (16) | 0.0422 (7) |
| N1 | 0.58919 (13) | 0.3645 (5) | 0.74827 (18) | 0.0256 (7) |
| C1 | 0.52901 (19) | 0.7558 (10) | 1.1327 (3) | 0.0560 (12) |
| H1 | 0.4965 | 0.8807 | 1.1287 | 0.067* |
| C2 | 0.56931 (19) | 0.6012 (9) | 1.1376 (3) | 0.0431 (9) |
| C3 | 0.61984 (16) | 0.4124 (9) | 1.1403 (2) | 0.0410 (9) |
| H3A | 0.6035 | 0.2354 | 1.1566 | 0.049* |
| H3B | 0.6491 | 0.4648 | 1.1911 | 0.049* |
| C4 | 0.65405 (14) | 0.3935 (7) | 1.0474 (2) | 0.0263 (8) |
| H4 | 0.6865 | 0.2547 | 1.0544 | 0.032* |
| C5 | 0.68451 (14) | 0.6481 (6) | 1.0216 (2) | 0.0245 (8) |
| C6 | 0.61569 (14) | 0.3295 (7) | 0.9594 (2) | 0.0247 (8) |
| H6A | 0.6128 | 0.1362 | 0.9489 | 0.030* |
| H6B | 0.5738 | 0.4046 | 0.9638 | 0.030* |
| C7 | 0.65240 (13) | 0.4640 (6) | 0.8817 (2) | 0.0224 (7) |
| H7 | 0.6849 | 0.3401 | 0.8590 | 0.027* |
| C8 | 0.61449 (14) | 0.5591 (6) | 0.7980 (2) | 0.0195 (7) |
| C9 | 0.55407 (14) | 0.4086 (7) | 0.6607 (2) | 0.0251 (8) |
| H9 | 0.5330 | 0.5838 | 0.6658 | 0.030* |
| C10 | 0.50518 (15) | 0.1967 (7) | 0.6510 (2) | 0.0289 (8) |
| H10A | 0.4773 | 0.2072 | 0.7049 | 0.043* |
| H10B | 0.5247 | 0.0219 | 0.6500 | 0.043* |
| H10C | 0.4820 | 0.2238 | 0.5920 | 0.043* |
| C11 | 0.59636 (13) | 0.4191 (7) | 0.5757 (2) | 0.0239 (7) |
| C12 | 0.58937 (15) | 0.6078 (7) | 0.5060 (2) | 0.0279 (8) |
| H12 | 0.5584 | 0.7388 | 0.5125 | 0.034* |
| C13 | 0.62615 (15) | 0.6116 (7) | 0.4271 (2) | 0.0314 (8) |
| H13 | 0.6201 | 0.7424 | 0.3795 | 0.038* |
| C14 | 0.67219 (14) | 0.4224 (7) | 0.4175 (2) | 0.0287 (8) |
| C15 | 0.68019 (15) | 0.2328 (8) | 0.4866 (3) | 0.0359 (9) |
| H15 | 0.7115 | 0.1032 | 0.4808 | 0.043* |
| C16 | 0.64217 (16) | 0.2328 (7) | 0.5645 (2) | 0.0340 (9) |
| H16 | 0.6477 | 0.1007 | 0.6118 | 0.041* |
| C17 | 0.74758 (19) | 0.2307 (10) | 0.3167 (3) | 0.0549 (12) |
| H17A | 0.7243 | 0.0649 | 0.3137 | 0.082* |
| H17B | 0.7793 | 0.2175 | 0.3661 | 0.082* |
| H17C | 0.7669 | 0.2632 | 0.2554 | 0.082* |
| H1H | 0.5970 (14) | 0.182 (8) | 0.764 (2) | 0.028 (9)* |

*Atomic displacement parameters (Å²)*

| | $U^{11}$ | $U^{22}$ | $U^{33}$ | $U^{12}$ | $U^{13}$ | $U^{23}$ |
|---|---|---|---|---|---|---|
| O1 | 0.0283 (12) | 0.0290 (13) | 0.0408 (14) | −0.0031 (11) | −0.0086 (11) | −0.0051 (12) |
| O2 | 0.0244 (12) | 0.0241 (13) | 0.0275 (13) | −0.0024 (10) | −0.0027 (9) | 0.0032 (10) |
| O3 | 0.0303 (12) | 0.0180 (13) | 0.0334 (14) | 0.0038 (10) | −0.0014 (10) | 0.0042 (11) |
| O4 | 0.0349 (13) | 0.0588 (18) | 0.0330 (14) | 0.0001 (14) | 0.0083 (11) | −0.0016 (14) |
| N1 | 0.0388 (16) | 0.0145 (16) | 0.0234 (15) | 0.0043 (13) | −0.0048 (13) | 0.0014 (12) |
| C1 | 0.034 (2) | 0.070 (3) | 0.064 (3) | −0.006 (2) | 0.008 (2) | −0.023 (2) |
| C2 | 0.0375 (18) | 0.057 (2) | 0.0350 (17) | −0.0166 (17) | 0.0080 (16) | −0.0104 (18) |
| C3 | 0.0420 (18) | 0.050 (2) | 0.0305 (17) | −0.0148 (17) | 0.0000 (15) | −0.0023 (18) |
| C4 | 0.0292 (17) | 0.0221 (18) | 0.0276 (17) | −0.0020 (15) | −0.0048 (14) | 0.0008 (15) |
| C5 | 0.0180 (15) | 0.0234 (19) | 0.032 (2) | 0.0068 (15) | −0.0038 (14) | −0.0020 (17) |
| C6 | 0.0299 (17) | 0.0198 (18) | 0.0243 (18) | −0.0048 (15) | 0.0001 (14) | 0.0014 (15) |
| C7 | 0.0203 (16) | 0.0172 (16) | 0.0296 (18) | 0.0036 (14) | 0.0021 (14) | −0.0039 (15) |
| C8 | 0.0182 (16) | 0.0173 (19) | 0.0233 (18) | 0.0044 (14) | 0.0072 (14) | 0.0013 (14) |
| C9 | 0.0299 (17) | 0.0230 (18) | 0.0224 (17) | 0.0059 (15) | −0.0018 (14) | 0.0014 (15) |
| C10 | 0.0310 (18) | 0.0254 (18) | 0.0304 (19) | 0.0030 (16) | 0.0001 (15) | 0.0031 (16) |
| C11 | 0.0264 (17) | 0.0214 (18) | 0.0237 (18) | −0.0032 (16) | −0.0041 (14) | −0.0017 (16) |
| C12 | 0.0284 (18) | 0.0264 (19) | 0.0288 (19) | 0.0024 (15) | −0.0053 (15) | 0.0000 (16) |
| C13 | 0.0335 (19) | 0.036 (2) | 0.0250 (19) | −0.0006 (18) | −0.0007 (15) | 0.0058 (16) |
| C14 | 0.0240 (17) | 0.038 (2) | 0.0242 (19) | −0.0076 (17) | 0.0020 (15) | −0.0077 (18) |
| C15 | 0.0287 (18) | 0.041 (2) | 0.038 (2) | 0.0067 (17) | 0.0015 (17) | −0.0031 (19) |
| C16 | 0.0358 (19) | 0.030 (2) | 0.036 (2) | 0.0078 (17) | 0.0022 (17) | 0.0032 (17) |
| C17 | 0.039 (2) | 0.071 (3) | 0.055 (3) | −0.001 (2) | 0.019 (2) | −0.013 (2) |

*Geometric parameters (Å, º)*

| | | | |
|---|---|---|---|
| O1—C5 | 1.205 (4) | C7—C8 | 1.512 (5) |
| O2—C5 | 1.351 (4) | C7—H7 | 1.0000 |
| O2—C7 | 1.447 (4) | C9—C10 | 1.515 (5) |
| O3—C8 | 1.235 (4) | C9—C11 | 1.520 (4) |
| O4—C14 | 1.365 (4) | C9—H9 | 1.0000 |
| O4—C17 | 1.428 (5) | C10—H10A | 0.9800 |
| N1—C8 | 1.325 (4) | C10—H10B | 0.9800 |
| N1—C9 | 1.464 (4) | C10—H10C | 0.9800 |
| N1—H1H | 0.96 (4) | C11—C12 | 1.377 (5) |
| C1—C2 | 1.177 (6) | C11—C16 | 1.382 (5) |
| C1—H1 | 0.9500 | C12—C13 | 1.380 (4) |
| C2—C3 | 1.457 (6) | C12—H12 | 0.9500 |
| C3—C4 | 1.518 (4) | C13—C14 | 1.393 (5) |
| C3—H3A | 0.9900 | C13—H13 | 0.9500 |
| C3—H3B | 0.9900 | C14—C15 | 1.376 (5) |
| C4—C5 | 1.494 (5) | C15—C16 | 1.384 (5) |
| C4—C6 | 1.525 (5) | C15—H15 | 0.9500 |
| C4—H4 | 1.0000 | C16—H16 | 0.9500 |
| C6—C7 | 1.524 (4) | C17—H17A | 0.9800 |
| C6—H6A | 0.9900 | C17—H17B | 0.9800 |
| C6—H6B | 0.9900 | C17—H17C | 0.9800 |
| C5—O2—C7 | 109.5 (2) | N1—C9—C10 | 109.2 (3) |
| C14—O4—C17 | 117.7 (3) | N1—C9—C11 | 110.8 (2) |

| | | | |
|---|---|---|---|
| C8—N1—C9 | 123.1 (3) | C10—C9—C11 | 112.7 (3) |
| C8—N1—H1H | 121 (2) | N1—C9—H9 | 108.0 |
| C9—N1—H1H | 116 (2) | C10—C9—H9 | 108.0 |
| C2—C1—H1 | 180.0 | C11—C9—H9 | 108.0 |
| C1—C2—C3 | 177.9 (4) | C9—C10—H10A | 109.5 |
| C2—C3—C4 | 113.3 (3) | C9—C10—H10B | 109.5 |
| C2—C3—H3A | 108.9 | H10A—C10—H10B | 109.5 |
| C4—C3—H3A | 108.9 | C9—C10—H10C | 109.5 |
| C2—C3—H3B | 108.9 | H10A—C10—H10C | 109.5 |
| C4—C3—H3B | 108.9 | H10B—C10—H10C | 109.5 |
| H3A—C3—H3B | 107.7 | C12—C11—C16 | 117.7 (3) |
| C5—C4—C3 | 112.4 (3) | C12—C11—C9 | 121.6 (3) |
| C5—C4—C6 | 103.0 (3) | C16—C11—C9 | 120.7 (3) |
| C3—C4—C6 | 116.6 (3) | C11—C12—C13 | 121.7 (3) |
| C5—C4—H4 | 108.2 | C11—C12—H12 | 119.1 |
| C3—C4—H4 | 108.2 | C13—C12—H12 | 119.1 |
| C6—C4—H4 | 108.2 | C12—C13—C14 | 119.6 (3) |
| O1—C5—O2 | 120.7 (3) | C12—C13—H13 | 120.2 |
| O1—C5—C4 | 128.5 (3) | C14—C13—H13 | 120.2 |
| O2—C5—C4 | 110.8 (3) | O4—C14—C15 | 124.7 (3) |
| C7—C6—C4 | 101.8 (3) | O4—C14—C13 | 115.6 (3) |
| C7—C6—H6A | 111.4 | C15—C14—C13 | 119.7 (3) |
| C4—C6—H6A | 111.4 | C14—C15—C16 | 119.4 (3) |
| C7—C6—H6B | 111.4 | C14—C15—H15 | 120.3 |
| C4—C6—H6B | 111.4 | C16—C15—H15 | 120.3 |
| H6A—C6—H6B | 109.3 | C11—C16—C15 | 122.0 (3) |
| O2—C7—C8 | 109.0 (3) | C11—C16—H16 | 119.0 |
| O2—C7—C6 | 105.1 (2) | C15—C16—H16 | 119.0 |
| C8—C7—C6 | 114.6 (2) | O4—C17—H17A | 109.5 |
| O2—C7—H7 | 109.3 | O4—C17—H17B | 109.5 |
| C8—C7—H7 | 109.3 | H17A—C17—H17B | 109.5 |
| C6—C7—H7 | 109.3 | O4—C17—H17C | 109.5 |
| O3—C8—N1 | 123.9 (3) | H17A—C17—H17C | 109.5 |
| O3—C8—C7 | 122.6 (3) | H17B—C17—H17C | 109.5 |
| N1—C8—C7 | 113.5 (3) | | |

| | | | |
|---|---|---|---|
| C2—C3—C4—C5 | 63.2 (4) | C6—C7—C8—N1 | −66.3 (3) |
| C2—C3—C4—C6 | −55.3 (4) | C8—N1—C9—C10 | −149.3 (3) |
| C7—O2—C5—O1 | 175.7 (3) | C8—N1—C9—C11 | 86.0 (4) |
| C7—O2—C5—C4 | −3.6 (3) | N1—C9—C11—C12 | −136.2 (3) |
| C3—C4—C5—O1 | 38.3 (5) | C10—C9—C11—C12 | 101.1 (4) |
| C6—C4—C5—O1 | 164.5 (3) | N1—C9—C11—C16 | 45.8 (4) |
| C3—C4—C5—O2 | −142.5 (3) | C10—C9—C11—C16 | −76.9 (4) |
| C6—C4—C5—O2 | −16.2 (3) | C16—C11—C12—C13 | 0.6 (5) |
| C5—C4—C6—C7 | 27.9 (3) | C9—C11—C12—C13 | −177.5 (3) |
| C3—C4—C6—C7 | 151.3 (3) | C11—C12—C13—C14 | −0.8 (5) |
| C5—O2—C7—C8 | 145.5 (2) | C17—O4—C14—C15 | 11.3 (5) |
| C5—O2—C7—C6 | 22.2 (3) | C17—O4—C14—C13 | −169.4 (3) |
| C4—C6—C7—O2 | −30.7 (3) | C12—C13—C14—O4 | −178.8 (3) |
| C4—C6—C7—C8 | −150.5 (3) | C12—C13—C14—C15 | 0.5 (5) |
| C9—N1—C8—O3 | 4.9 (5) | O4—C14—C15—C16 | 179.3 (3) |
| C9—N1—C8—C7 | −175.7 (3) | C13—C14—C15—C16 | 0.1 (5) |
| O2—C7—C8—O3 | −4.4 (4) | C12—C11—C16—C15 | 0.0 (5) |
| C6—C7—C8—O3 | 113.1 (3) | C9—C11—C16—C15 | 178.1 (3) |
| O2—C7—C8—N1 | 176.2 (2) | C14—C15—C16—C11 | −0.4 (5) |

1. Cox, J.; Mann, M., MaxQuant enables high peptide identification rates, individualized ppb-range mass accuracies and proteome-wide protein quantification. *Nature biotechnology* **2008,** *26* (12), 1367.

2. Li, D.; Fu, Y.; Sun, R.; Ling, C. X.; Wei, Y.; Zhou, H.; Zeng, R.; Yang, Q.; He, S.; Gao, W., pFind: a novel database-searching software system for automated peptide and protein identification via tandem mass spectrometry. *Bioinformatics* **2005,** *21* (13), 3049-3050.

3. Boersema, P. J.; Raijmakers, R.; Lemeer, S.; Mohammed, S.; Heck, A. J., Multiplex peptide stable isotope dimethyl labeling for quantitative proteomics. *Nature protocols* **2009,** *4* (4), 484-494.
